# Adaptive Introgression Promotes Fast Adaptation In Oaks Marginal Populations

**DOI:** 10.1101/731919

**Authors:** Pablo G Goicoechea, Laura Guillardín, Lierni Fernández-Ibarrodo, Maria Valbuena-Carabaña, Santiago C González-Martínez, Ricardo Alía, Antoine Kremer

## Abstract

Range shifts and species range limits are two fundamental, related processes in population and evolutionary genetics that have received much attention since a large impact of climate change in species’ distributions was predicted. In general, there is a broad consensus on the effects of abiotic interactions on range limits, but comprehensive evidence supporting/rejecting the impact of biotic interactions is lacking. Hybridization has long been recognized as a biotic interaction favoring marginal populations establishment and range expansion through transgressive segregation or adaptive introgression, but recently new roles have been claimed for hybridization, such as the trigger of adaptive radiations, or indirect effects on population sizes that would allow persistence until new mutations arises or the environment changes. In this work, we selected two Mediterranean oak species with ecological discrimination based on soil pH, and intensively sampled three interspecific pairs of marginal populations from taxon-extreme environments under heterogeneous climate conditions. We genotyped 110 EST-SSR markers evenly distributed across their genomes and applied a variety of population and landscape genetics models to validate candidate genes for local adaptation. Then, several introgression screens on shared candidates showed that the three inter-specific population pairs contain evidences of adaptive introgression and that events occur in both directions. Other significant findings from our work are: (i) Aproximate Bayesian Computation coupled to coalescent simulations supports small hybridization rates since recent secondary contact in two population pairs affected by Quaternary climatic oscillations but continuous old interspecific gene flow in the pair less affected by climate, (ii) introgression at loci involved in local adaptations leads to strong geographic structure of marginal oak populations when sampling is large enough, and (iii) sampling efforts can be targeted to reveal different components of populations structure. Finally, we review evidences that support our conclusions and discuss some evolutionary implications of adaptive introgression on range expansion.

## Introduction

Limits to species distribution areas and range shifts (expansion, contraction or both) are related processes in ecology, population and evolutionary genetics that have received renewed attention since large impacts on species distributions were predicted as a consequence of climate change (Alexander *et al.*, 2015; Urban *et al.*, 2016). Understanding how species establish limits to their distribution areas and under which circumstances marginal populations will be able to survive or shift their ranges without noticeable loss of diversity is difficult because of the myriad biotic and abiotic interactions species/populations face (Sexton *et al.*, 2009; Gilman *et al.*, 2010). However, whereas a broad consensus on the importance of abiotic interactions has been reached (Gaston, 2009; Atkins and Travis, 2010), there is not general evidence to support/reject the importance of biotic interactions on niche limits and range dynamics (Godsoe *et al.*, 2017).

Marginal populations, those at the range edges, have played an important role in studies of species’ distributions and range shifts, as their individuals are the most likely to disperse beyond the range, even if they are not the best fit to colonize the new habitats (McLane and Aitken, 2012). Indeed, they are also prone to extinction, when environmental and/or biotic interactions increment population stress beyond a limit (Aspin *et al.*, 2019). Establishment of marginal populations, at a certain point along environmental gradients, is thought to occur via local adaptation at range edge conditions (Levin, 2000, Hargreaves *et al.*, 2015), which should confer marginal populations an advantage beyond the range relative to central populations (Hargreaves and Eckert, 2019; but see McLane and Aitken, 2012). However, fitness and population sizes are reduced towards the range edges (Pironon *et al.*, 2017), thus bringing undesirable negative effects to offspring quality that could wreck their adaptation and colonization ability (Hargreaves *et al.*, 2015). In the presence of gene flow from the central population, marginal populations are demographic sinks (Dias, 1996) and the balance between gene flow (m) and selection (s) is the single most important factor influencing the fate of adaptive alleles. They will be maintained and their frequencies will rise if migration is small compared to selection [specifically if 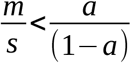; where *a* is the ratio of selection coefficients between two demes], or they will be lost by gene swamping if migration is large compared to selection (specifically if the left term from the equation is larger than the right term), and maladaptation will occur (Lenormand, 2002; Kawecki, 2008). Other aspects influencing the establishment and maintenance of adaptive alleles are traits architecture, the size of allele effects, linkage with already selected gene(s), recombination, or the landscape grain coarseness (Gillespie, 1974; Endler, 1977; Barton, 1983; Le Corre and Kremer, 2012; Samuk *et al.*, 2017).

Over the last two decades, the paradigm shift from a ‘coherent’ to a ‘porous’ genome (Wu, 2001), advances in gene flow estimates (Hey, 2006, Sousa and Hey, 2013) and in genome scan (GS) and genetic-environmental association (GEA) models (Foll and Gaggiotti, 2008; Eckert *et al*., 2010), together with the shift of focus from humans to non-model organisms, have revolutionized with a wealth of data studies of adaptation (and speciation) with gene flow (Haasl and Payseur, 2016; Tigano and Friesen, 2016). However, non-addressed sources of confounding and severe but accurate criticisms of some studies conclusions and of *F_ST-_* based GS and GEA models, have largely discredited the use of these methods in adaptation-speciation genetics (reviewed by Hoban *et al.*, 2016; Ravinet *et al.*, 2017; Wolf and Ellegren, 2017). Conflicting issues include (1) the statistical properties, the significance and the low information content of *F_ST_*-related differentiation coefficients (Whitlock and McCauley, 1999; Jost, 2008; Buerkle *et al*., 2011; Meirmans and Hedrick, 2011; Bierne *et al.*, 2013; Haasl *et al.*, 2014; Buerkle, 2017); (2) heterogeneity of mutation and recombination rates (Noor and Bennet, 2009; Nachman and Payseur, 2012; Cruickshank and Hahn, 2014; Haasl and Payseur, 2016); (3) linked selection, background selection, selection at non-targeted loci and selection from standing variation (Schluter and Conte, 2009; Charlesworth, 2012; Cutter and Payseur, 2013; Burri *et al.*, 2015; Christe *et al.*, 2017); (4) demographic and evolutionary histories, including ancient introgression and introgression from other sister species (Fraïsse *et al.*, 2014, 2016; Nadeau *et al.*, 2016; Ma *et al.*, 2018); (5) gene density (Nordborg *et al.*, 2005; Yeaman, 2013); or (6) the genetic architecture of adaptive and/or barrier loci (Le Corre and Kremer, 2012; Flaxman *et al.*, 2014; Conte *et al.*, 2015; Yeaman, 2015). GEA models have evolved with the aim to reduce false-positives from diverse sources of confounding, such as patterns of isolation-by-distance (IBD), alignment of environmental and genetic gradients, or cryptic relatedness (Frichot *et al.*, 2013; Frichot *et al.*, 2015; Rellstab *et al.*, 2015; de Villemereuil *et al.*, 2015) and they probably outperform GS models in identifying environmentally-driven local adaptation (De Mita *et al.*, 2013; Payseur and Rieseberg, 2016). However, a number of pitfalls have been also identified in GEA models, leading to ‘roadmaps’ and ‘good practices’ recommendations (Tiffin and Ross-Ibarra, 2014; Ćalić *et al.*, 2016; Hoban *et al.*, 2016; Richardson *et al.*, 2016; Li *et al.*, 2017). In spite of criticism, GSs and GEAs have claimed the identification of genes under natural selection for local adaptation in a large number of species (2,757 PUBMED references for “local adaptation genes”, peaking at 288 in 2016).

Standing variation, new mutations and hybridization are the raw materials for adaptation to novel environments (Barret and Schlutter, 2008; Hedrick, 2013). They leave different footprints in the genomic signatures of selection imposed by the environment and they promote local adaptation at different rates: rare hybridization events result in intermediate evolutionary rates, although recurrent introgression might increase initial frequencies leading to faster evolution than standing variation (Hedrick, 2013). Traditionally, scientists have considered two main beneficial effects of hybridization on adaptation and range expansion: (i) transgressive segregations, including homoployds and alloployds (Rieseberg *et al*., 2003; Gerstein *et al*., 2011; Zhang *et al.*, 2017), and (ii) adaptive introgression of large effect local alleles (Suarez-Gonzalez *et al.*, 2018; Leroy *et al*., 2019). In addition, hybridization could have other evolutionary benefits, such as promoting adaptive radiations (Meier *et al*., 2017), or increasing effective population size of edge populations to allow survival until new mutations are available or biotic/abiotic interactions become more favorable (Pfenning *et al.*, 2016; Pierce *et al*., 2017).

The European white oaks represent the newest radiation (1.5 – 5 MY) within the genus *Quercus*, which underwent older radiations in Asia and North America (Hubert *et al.*, 2014). The sympatric temperate species *Q. robur* L. and *Q. petraea* (Matt.) Liebl. have the largest distribution areas and economic importance, which explains the long-term interests in their study and the recent development of genomic assets (Lesur *et al.*, 2015; Bodénès *et al.*, 2016; Plomion *et al.*, 2016) to complement traditional genetic resources (provenance trials, intra- and inter-specific crosses, etc.). Long-term provenance trials established during the last century have discovered large amounts of genetic and phenotypic variance for adaptive (e.g, phenology) and growth traits (Ducousso *et al*., 1996; Jensen and Hansen, 2006), whereas recent molecular analyses have identified several of the genes involved in adaptation, adaptive introgression and speciation (Rellstab *et al.*, 2016; Leroy *et al.*, 2018; 2019).

The Mediterranean basin contains over a dozen, perhaps 20 little known oak species with sometimes unclear taxonomic status, low economic value but a high relevance to the regional ecology of the three southern Mediterranean peninsulas (Webb, 2010). In Iberia, *Q. faginea* Lam. (‘quejigo’) and *Q. pyrenaica* Willd. (‘marojo’) are the white oaks with largest distribution areas (insets in Figure 1a). These oaks show ecological exclusion by soil pH, the first one inhabiting the limy soils of the south and the east, the second growing preferentially in the acidic soils of the north-west (Blanco-Castro *et al.*, 1997). Although there is no provenance trials of these oaks, and thus local adaptation of their ecotypes is just an assumption, chances are that strong climatic differences between the north/northwest and the south/east of Iberia have favored local adaptations to aridity/temperate climate and/or other ecological conditions. Furthermore, previous genetic studies demonstrated the existence of intermediate genotypes that were cataloged as hybrids/admixed trees (Valbuena-Carabaña *et al.*, 2006; Goicoechea *et al.*, 2015), which opens up the possibility of adaptive introgression at least in some populations.

**Figure 1:**
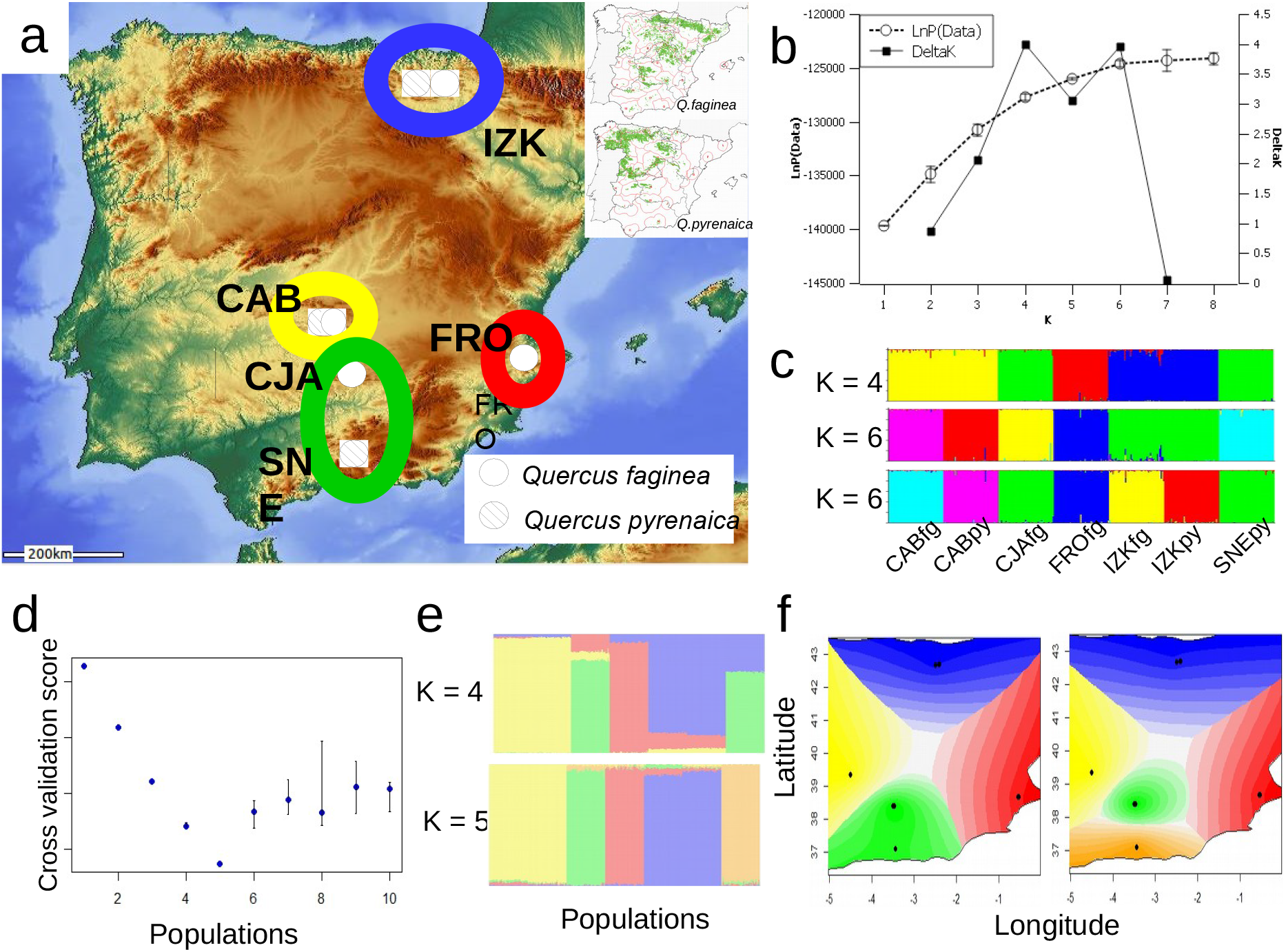
Sampling locations and inferred ancestral relationships for the seven populations analyzed in this study (a) Inference according to Structure, the inset shows approximate current distribution of both species (https://especiesforestales.com/D_Qpyrenaica.html). (b) probabilities of Structure models with different numbers of ancestor populations and Evano’s inference of the “true” number of ancestors; (c) individual ancestry proportions for models with *K*=4 and *K*=6 ancestors. Cross-validation scores, individual ancestry proportions for *K*=4 and *K*=5, and landscape plots of the inferred ancestries using Tess3r are shown in (d), (e) and (f), respectively.

In the present study we focus on inter-specific pairs of populations at the distribution edges of both Mediterranean oaks and analyze their genetic diversity using 110 EST-SSRs. We apply an array of GS and GEA models ‘to find the needle in the hay’ and to build a set of strong local adaptation candidates, which we use to elucidate the roles of hybridization and adaptive introgression in marginal population dynamics. Specifically, we aimed to investigate whether hybridization has a direct effect on adaptation at the range limits through adaptive introgression of large effect alleles, or whether hybridization’s indirect effect over population size is of importance in these species. We reasoned that under the direct introgression hypothesis both species should have the same alleles responsible for local adaptations, whereas adaptive alleles could and should be different in both species under the hypothesis of indirect hybridization effects. Thus, the main objectives of our study are: (i) to test the suitability of allele-frequency based GSs and GEAs in the characterization of local adaptation signatures using EST-SSRs markers, and (ii) to search adaptive introgression footprints in a reduced set of strong candidates for local adaptation. Additional objectives derived from the results obtained were (iii) to model population dynamics at the range edges using coalescent simulations coupled to model selection by approximate Bayesian computation (ABC), and (iv) to analyze the effects of local adaptation and introgression in the genetic structure of marginal populations.

## Materials and methods

A total of 336 trees (48 x 7) were sampled from 5 Natural Parks in Spain to get three inter-specific population pairs plus one extra *Q. faginea* population (Figure 1a). The first pair was collected from ‘El Estrecho’ place within Cabañeros National Park, which contains a parapatric oak forest with a sharp contact area between the two target species (populations CABfg and CABpy). The second pair was sampled from Izki Natural Park, which is best known by its large *Q. pyrenaica* forest (population IZKpy), but it also contains a hybrid zone ending into a clay basset that contains an apparently pure *Q. faginea* stand (population IZKfg). Distance between these populations is approximately 5 km. The remaining forests contain one of the analyzed white oaks in sympatry with evergreen holm oaks. The third interspecific population pair was sampled from Collado de los Jardines Natural Park and from Sierra Nevada National Park (populations CJAfg and SNEpy, respectively) and it was meant to represent allopatry (distance between the two mountain locations from south Iberia is larger than 100 km) as well as altitudinal constraints (the CJAfg population is located at the highest altitude of the mountain range). The last *Q. faginea* population was sampled from another mountain location at the Font Roja Natural Park (population FROfg), which is located about 300 km from Sierra Nevada, because biogeography relates both locations. The three population pairs represent contrasting environmental conditions favoring heterogeneous natural selection (arid-Mediterranean, near temperate and high altitude-Mediterranean, respectively) and increasing distances between the two populations from each pair, which could have implications on interspecific gene flow and thus in introgression.

### Environmental Variables

Individual latitude and longitude coordinates from each sampled tree were obtained with a portable GPS and they are provided within the file deposited at (to be specified on publication decision). Climatic raw data for each sampling site (Supplemental File 1, Tables S1-1a-S1-1d; note the same environmental values were used for the two CAB populations) were spatially interpolated (200 meters spatial resolution) based on historic climate records at a representative network of 2,285 weather stations (Ninyerola et al., 2005). The lengths of climate records vary from 15 to 50 years for the air temperature and solar radiation data, whereas precipitation records contain 20-50 years during the period 1951-1999. Monthly means of the 4 climate variables at the sampling sites were obtained from the open Geographic Information System (GIS) map server ‘Atlas Climático Digital de la Península Ibérica’ (http://opengis.uab.es/wms/iberia/index.htm). We used this local system because it has a larger resolution within the Iberian Peninsula than other global systems (e.g., WorldClim).

Raw climate data were transformed into 27 Bioclim variables that represent annual trends, seasonality and extreme or limiting environmental factors (Hutchinson *et al*., 2009), which are suitable for species distribution modeling (SDM; Xu and Hutchinson, 2011, Booth *et al.*, 2014) but their performance on GEA models has not been tested to date. Table S1-2 provides the definitions and the centered, standardized values of the first 27 Bioclim variables for the 7 studied populations. Highly correlated variables (> 0.85) were dropped to obtain a reduced dataset of 10 ‘uncorrelated’ Bioclim variables in order to test effects of correlated Bioclim variables in GEA models. We used the R libray ‘dismo’ to obtain Bioclims 1-19, whereas Bioclims 20-27 were calculated in LibreOffice spreadsheets (https://www.libreoffice.org/).

### Genetic Markers

One-hundred and ten microsatellite markers (103 EST-SSRs and 7 gSSRs) were used to analyze the genome-wide architecture of genetic diversity in the seven populations. Seventy-two markers had been previously used and described by Goicoechea *et al*. (2015); the rest were selected from available European oak resources to fill the gaps in the composite consensus genetic map (*https://w3.pierroton.inra.fr/QuercusPortal/*). Supplemental File 2, Table S2-1 provides a full description of the 110 SSRs used in this study, including the linkage group (LG) they map to, the repeat motifs, the forward and reverse primer sequences and the annealing temperatures.

Details regarding DNA isolation and purification, PCR amplifications and electrophoresis conditions, as well as the method for binning raw allele sizes have been described elsewhere (Goicoechea *et al.*, 2015), with slight modifications. In particular, the concentration of the M13 5’-marked primer in the PCR mixes were cut to half at the cost of a small increase in the number of PCR cycles. Furthermore, the construction of a composite consensus map, using the R library LPmerge (Endelman and Plomion, 2014), was carried out with all available SSR markers from each linkage group. Then, internal markers not used in this study were dropped from the respective LGs to draw the composite consensus map shown in Figure 2

**Figure 2:**
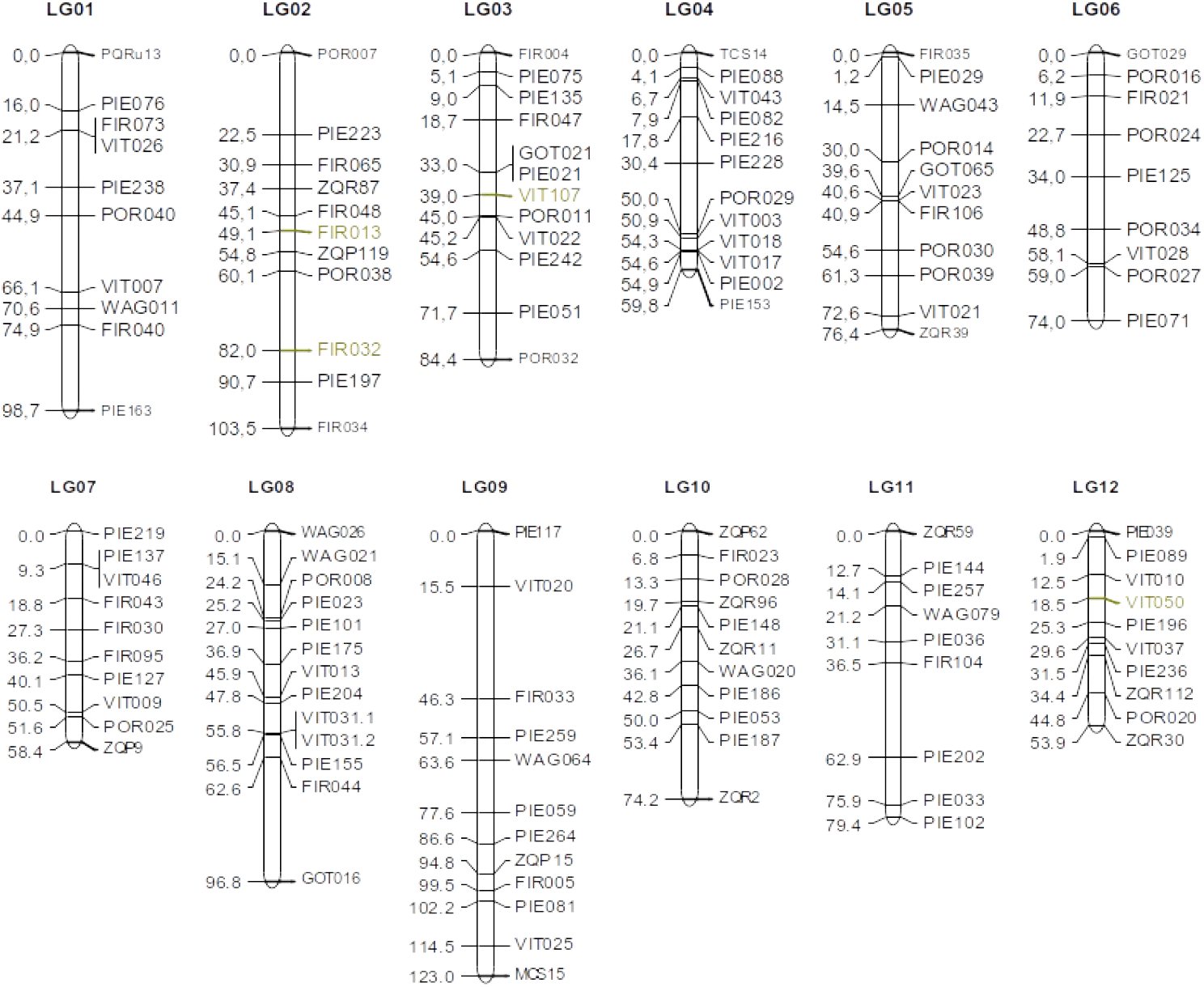
Composite linkage map of the 110 markers used in this study. Several markers have been added to the end of the respective linkage groups for reference. Colored markers have been assigned approximate positions within the respective LGs.

### Population Structure

The genetic structure of the samples was studied with the model-based Bayesian clustering method of Structure v.2.3.3 (Pritchard *et al.*, 2000; Falush *et al.*, 2003) and with the spatially-explicit, model-free method implemented in the R library Tess3r (Caye *et al.*, 2016), which provides geographically constrained least-squares estimates of ancestry coefficients. The analyses were carried out for the complete dataset, for each population pair and for each species separately. We also analyzed sub-sampled datasets with n=8, 12, 24, 32 individuals per population, and a ‘neutral’ dataset with 70 markers (see below).

Most Structure models inferred independent ancestry for each of the seven populations. Only the ‘Linkage’ model with independent allele frequencies and *λ* estimated from the data (*λ* is the parameter of the Dirichlet distribution used to draw allele frequencies) was able to infer some genetic structure. Thus, we run 5 independent chain replicates of 5×10^5^ iterations following a burn-in period of 2.5×10^5^ iterations, for a fixed number of populations (*K*=1-8). Convergence of the Bayesian chains posterior distributions was surveyed with plots of the parameter likelihoods across iterations. The most likely number of genetic groups was inferred according to Evanno *et. al.* (2005) with the web-based software Structure-Harvester (Earl and vonHoldt, 2012). Then, a long run was set up, with 2×10^6^ iterations following a burn-in period of 5’10^5^ iterations, to gain confidence in admixture proportions. The group membership values for each individual, at the most likely *K*, were visualized with Structure plots.

The main idea implemented in Tess3 (Caye *et al.*, 2016) is that the probability an individual *i* carries the genotype *j* at locus *l* is determined by the law of total probability:

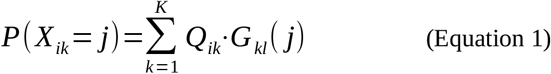

where *Q_ik_* and *G_kl_* are elements of the Q-matrix of individual ancestry coefficients and the *G*-matrix of ancestral genotypic frequencies, respectively. The above formula establishes that each individual genotype is sampled from *K* pools of ancestral genotypes with sampling probabilities corresponding to their admixture coefficients. The formula is also equivalent to the factorization of the genotype probability matrix *P*, using matrices *Q* and *G* as factors (Frichot *et al*., 2014). Contrary to Structure, this method makes no assumptions about HWE, which makes it appropriate to deal with situations of inbreeding or geographically restricted mating.

Ancestry coefficients were estimated from the genotype matrix and the geographical coordinates for *K*=1 to 10 populations, using 30 repetitions of the algorithm up to a maximum of 10^4^ iterations. Root mean-squared errors, obtained from a subset of loci used for crossvalidation, were plotted for each *K* to estimate the number of ancestral populations. Individual admixture coefficients from the *Q*-matrix with lowest cross-validation scores were visualized using standard R plotting commands, which are available in the software vignette. Finally, the *Q*-matrix values were interpolated on a geographic map using the Tess3r command ‘plot’ and the R library ‘gglot2’.

The existence of a nested phylogeographic structure was tested by analyzing separately the geographic and phylogenetic components, i.e., analyses within species and within locations. To further investigate the stability of the sample structure, we analyzed 3-5 resampled datasets of 8, 12, 24 and 36 individuals from each population. Finally, the effect of non-neutral markers on population structure estimates was analyzed by means of a ‘neutral’ dataset of 70 markers, which was obtained after eliminating 40 markers from the tails of the global *F*_ST_ distribution values (20 markers from each side).

### Linkage Disequilibrium

Linkage disequilibrium (LD) between consecutive pairs of loci was tested within populations to avoid structure-related spurious LD, i.e., isolate breaking (Nei and Li, 1973). We used the multi-allelic specific test *T*2 (Zaykin *et al.*, 2008), which does not require haplotype inferences and it is robust to departures from HWE. Calculations were performed with the R library ‘pegas’ (Paradis, 2010). Significant T2 values were obtained by Chi-square tests, which render similar results to the permutation-based ‘exact’ tests (Zaykin *et al*., 2008). Corrections for multiple, non-independent comparisons were carried out according to Benjamini and Yekutieli (2001) using the ‘p.adjust’ function from the ‘R-stats’ package (R Core Team, 2015), with a false discovery rate of *q* = 0.1.

### Genome Scans

Within-species GSs should allow the discovery of genes involved in adaptation, specially in species with large distribution areas under contrasting environmental conditions. For this task, we used two GS models related to the GEA analysis described in the next section. BayeScan (Foll and Gaggiotti, 2008) shares its conceptual framework with BayeScEnv, but it is resilient to departures from the assumption of a large number of populations. The settings for these analysis were: 50 pilot runs of length 50,000 each, followed by the main chain consisting of 7.25×10^5^ iterations, with a thining interval of 25, after 10^5^ burning iterations. The false discovery rate (FDR) for all analyses was 10%. Di-nucleotide EST-SSRs were analyzed separately from 3/6-bp EST-SSRs to avoid false positives caused by differences in mutation rates (as recommended by the software authors).

Within-species genome scans were also performed with Tess3r, following the strategy described by Martins *et al.* (2016). For this task, the ‘tess3’ function was run for 5 values of the number of ancestral populations surrounding the *K* value previously estimated, and the number of iterations were increased up to a maximum of 150,000. This function returns, together with ancestry coefficients, an *F_ST_* statistic for each locus that can be tested against the null hypothesis of selective neutrality with the function ‘pvalue’. Adjusted *p*-values frequencies of the best run (i.e., the run with lowest root mean squared error between the observed and fitted genotype matrixes) were plotted for each *K* value (*K*=2-6) and the histogram shapes guided the selection of the best *K* to warrant efficient control of FDR, according to software authors instructions (François *et al.*, 2016). Manhattan plots of outliers for several FDR rates (*q* = 0.1, 0,01, 0,001, 0,0001), obtained with the Benjamini-Hochberg algorithm, were used for a final visual inspection of the reported outliers.

### Genomic-Environmental Associations

We estimated intra-specific GEAs using two models conceptually related to the above GSs: the latent factors mixed model (Lfmm; Frichot *et al.*, 2013) and BayeScEnv (de Villemereuil and Gaggiotti, 2015). We are aware that GEA models are most appropriate for large number of populations and clinal environmental variation (Endler, 1977). Therefore, our tests with a few populations from radically different environments will be an indicator of resilience against this type of model violations.

BayeScEnv is an extension to the outliers searching method of BayeScan (Foll and Gaggiotti, 2008) that allows testing the likelihood of a local adaptation effect (*g*) linked to an environmental variable (*E_j_*), in addition to the population (β) and locus specific (α) effects estimated by the original model. Thus, the three tests used to infer the genetic structure at individual loci were:

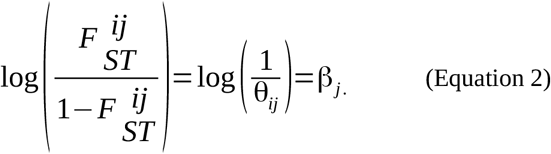

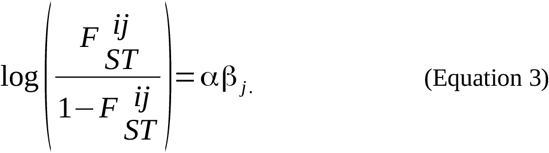

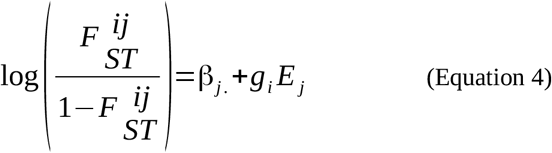

where *β_j_*. is the population-specific differentiation, *α* is the locus-specific effect, and *g_i_* is the local adaptation effect linked to environmental variable *E_j_*. Note that equations (2) and (3) constitute the original BayeScan test (Foll and Gaggiotti, 2008), whereas equations (3 and (4) allow to distinguish outliers driven by local adaptations from those due to unknown reasons (deviant mutation rates, etc). Contrary to other genomic-environmental association models that use the raw allele frequencies, BayeScEnv relies on the relation between the population-specific *F_ST_*(s) and the environmental differentiation (usually, but not necessarily, to the mean environment). Most importantly, BayeScEnv does not need covariates to account for population differentiation, as population structure is integrated into *β_j_*. Similarly to BayeScan, *q*-values statistical support was maintained at 10% FDR. The analysis, which was carried out separately for EST-SSRs with 2 and 3-6 bp repeat motifs, included 50 pilot runs of length 2.5×10^4^ iterations to approximate the posterior distributions of the parameters *g* and *α*, and a final run of 10^5^ iterations, with a thinning interval of 10, after 1.5×10^4^ burn-in iterations. The convergence of the posterior distributions and the effective size of the sample used to estimate the unknown parameters were tested for all significant GEAs using the R library ‘Coda’ (Plummer *et al*., 2006).

Environmentally-driven positive selection was further analyzed using Lfmm, as implemented in the R/Bioconductor library LEA (Frichot and François, 2015). Briefly, Lfmm fits a regression mixed model to the matrix of allele frequencies (G). The environmental covariates are introduced as fixed effects while population structure is modeled using latent factors. Mathematically,

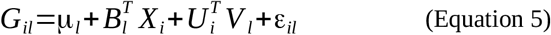

where *G_il_* is the matrix of alleles, *μ*_l_ is a locus specific effect, *B_l_* is a vector of regression coefficients, *X_i_* is a vector of environmental variables for each individual, *U_i_ V_l_* are scalar vectors with *K* dimensions that model latent factors and their scores, and ε_*il*_ are the residuals.

Fitting the Bioclim variables to the genotypic frequencies with the latent-factors mixed model was carried out in two different ways. First, each Bioclim variable was fit independently, similarly to the BayeScEnv model. Furthermore, all Bioclim variables were fit at the same time with Lfmm ‘all=TRUE’ option (we called this model LfmmAll). Missing data imputation, which was carried out with the Lfmm function ‘impute’ using the best ‘snmf’ run, had no effects on the results (not shown), possibly because they represented a small proportion of the data (< 0.05).

Associations between alleles with minimum allele frequency (MAF) larger than 5% and the Bioclim variables were sought with thirty MCMC replicates of 100,000 iterations, including 50,000 burn-in iterations, for several latent factors values close to the most likely *K* previously obtained with Tess3r (*K*=2-6). To increase the power of the Lfmm test statistic we combined the *z*-scores from the 30 replicates and used its median to obtain *p*-values adjusted with the ‘genomic control’ option (we did not observed measurable effects varying the inflation factors *λ* in the range 0.01-100, data not shown). Decisions on the appropriate number of latent factors for each Bioclim variable and the common FDR level(s) for all of them were taken as previously explained for the GS tests with Tess3r. For LfmmAll, the number of iterations was increased to 300,000, including 200,000 burn-in iterations.

### Adaptive Introgression

We are aware that detailed descriptions of adaptive introgression have not been possible without the aid of closely linked molecular markers (Kim *et al.*, 2008; Whitney *et al.*, 2015; Suarez-Gonzalez *et al.*, 2016; Bechsgaard *et al.*, 2017, Leroy *et al.*, 2019), and thus our sparsely located EST-SSR markers are not the best choice for detecting adaptive introgression in forest tree populations with large effective sizes and little LD. Further, robust validation of adaptive introgression requires the combination of several lines of evidence (Vekemans, 2010; Hedrick, 2013; Suarez-Gonzalez *et al.*, 2018), some of them without application in our study (for example, the positive effects of the allelic variants in the F1 hybrid). In spite of it, we selected local adaptation candidates shared between the two species and filtered them through several screens to gain support for hybridization and adaptive introgression by several lines of evidence.

First, we examined allele frequencies to ensure that shared outlier alleles belong to any of three interspecific population pairs (CABfg-CABpy, CJAfg-SNEpy, IZKfg-IZKpy). This does not evince adaptive introgression, but the opposite (i.e., outlier alleles shared between populations from different inter-specific pairs) should be taken as evidence of incomplete lineage sorting (ILS).

Second, we estimated the average squared distance (ASD; Goldstein *et al.*, 1995) between the three population pairs for the datasets of shared candidates (obtained in the previous introgression screen) and ‘neutral’ loci (as defined in the above Population Structure section) and obtained 10,000 bootstrapped replicates. Under the idealized strict Stepwise Mutation Model, ASD is linearly related to the ‘time to the most recent common ancestor’ (Tmrca) at least up to 2 million years ago (Sun *et al.*, 2009), and loss of linearity due to size changes is recovered when populations reach a new mutation-drift equilibrium (Goldstein *et al.*, 1996). Therefore, significantly larger ASD estimates for the mean genome (neutral dataset) than for the shared candidates dataset should be interpreted as evidence of introgression (Willis *et al.*, 2012). We are aware that Tmrca estimates are very sensitive to mutation rates, but we are just interested in comparing the observed ASD values between two groups of genes (see Gymrek *et al*., 2017, for sophisticated estimates of mutation rate and Tmrca). A serious concern in our approach is the small number of candidates within each population pair, which reduces the power of this tests (e.g., see the results for the CJA-SNE populations below; but see Bird, 2012). Thus, even if our tests don’t demonstrate introgression, we think they can still be useful to accumulate introgression evidences.

Third, we sought further evidence against ILS in markers adjacent to the EST-SSR outliers. For this screen, we considered ‘species-specific’ alleles those present in one of the species and in the introgressed population, although we allowed some residual frequencies in other populations at two candidates. Note that the possibility of introgression in the non-targeted population pair (e.g., FIR013, see Results), together with high mutation rates and homoplasy of the EST-SSR markers, make a good case for these allowances.

Fourth, we tested the adaptive nature of introgressed alleles at candidate loci by searching disruptive selection signatures in two-marker haplotypes within each species, using score tests for associations available in the R library ‘haplo.stats’ (Schaid *et al.*, 2002; Sinnwell and Schaid, 2009). Thus, two-marker haplotype frequencies of candidate outliers and introgressed markers from each population of the candidate pair were compared to haplotype frequencies in the remaining populations from the same species. Highly significant associations from common haplotypes in both within-species groups (candidate populations for adaptive introgression vs. other populations from the same species) would suggest different allelic combinations are selected in contrasting environments, i.e., disruptive selection. One final step in the characterization of hybrid introgression in the two Iberian oaks was the functional annotation of the candidate markers (next section).

### Gene Ontology and Gene References Into Functions

The Gene Ontolgy (GO) project was born at the end of the last century to address the need of consistent descriptions of gene products across databases. Three structured ontologies describe biological processes, cellular components and molecular functions associated to gene products, in a species independent style that allows uniform queries across databases (Ashburner *et al*., 2000; Gene Ontology Consortium, 2015). Gene References Into Functions (GeneRIFs) consist of short descriptions (less than 425 characters) of a gene function associated to a PubMed citation ID, which allows fast annotation of newly discovered functions (Mitchell *et al*., 2003). GeneRIFs annotations are available from the NCBI Gene databases.

GO terms for loci involved in GSs-GEAs were annotated with Blast2Go v2 (Götz *et al.*, 2008), which performs automated annotation augmentation, by searching the non-redundant *Arabidopsis* protein database with records from the NCBI ESTs databases (blastx). To retrieve those records, the EST database was searched with the sequence identifiers obtained from Durand *et al*. (2010) and the OakPortal SSRs database (http://ssrdatabase.pierroton.inra.fr/home). However, this approach failed to produce results for several EST-SSRs and could not be used with two gSSRs of unknown sequence. Blasting the NCBI records against the oak transcriptome OCV4 de-novo assembly (Lesur *et al*., 2015) and/or against the *Q. robur* v.1&2 whole-genome scaffolds assemblies (Plomion *et al*., 2016) identified larger contigs and/or the scaffold positions, which in turn allowed to obtain satisfactory ‘blastx’ searches. Several markers (POR030, POR038, ZQR87) were annotated by first searching the oak scaffold assemblies with the primer sequences (https://urgi.versailles.inra.fr/blast), which produced two alignments within unique scaffolds at positions that were compatible with the observed SSRs sizes. Blasting the surrounding scaffold sequences against the oak transcriptome assembly (Lesur *et al.*, 2015) and/or the *Arabidopsis* non-redundant protein database found the searched homology. The only exception was ZQR87, as PCR primers could be mapped to a unique scaffold but the upstream and downstream sequences hit different transcripts with identities lower than 90% and whose homology to *Arabidopsis* proteins was poor. Although this might be indicative of an intergenic sequence, we opted for showing the best blastx hit to the Viridiplantae group.

### Coalescent Estimates of Divergence with Gene Flow

We used ABCToolbox v.2 (Wegmann *et al.*, 2010) to analyze population dynamics and the approximate timing of inter-specific exchanges and adaptive introgression for each of the three population pairs. For this task, we used the ‘neutral’ dataset and compared first 5 simple alternative models: strict isolation (SI), isolation with migration (IM), ancient migration (AM), secondary contact (SC) and panmixia (PAN). Models SI and PAN were dropped from analysis soon, because at least one of the observed summary statistic fell out the simulated statistics envelope. Then, we considered variants of the three remaining models (AMc, IMc and SCc) that included two ghost ‘central’ populations of much larger effective population sizes, which sent genes into the sink ‘marginal’ populations at rates 10 times larger than inter-specific exchanges (Supplemental File 6, Figure S6-1). Mutation rate variation was only considered between the two groups of microsatellites (di- and tri/hexa-nucleotide repeat motifs), whereas different geometric parameters for the Generalized Stepwise Mutation (GSM) model imposed distinct rates of multistep mutations in the same two groups. Linkage relationships among the markers were discarded because run times increased prohibitively and all 70 simulated microsatellites were considered to be independent. Supplemental File 6 contains the full model descriptions and their complete parameterizations. Sequential model choice was carried out by first comparing the three models without ‘central’ populations (for each population pair), then comparing the three models with “central” populations and finally comparing the two best models for each population pair.

The observed data were 15 summary statistics of the three inter-specific population pairs, including the number of alleles from each population and their standard deviations (K1, K2, Ksd1, Ksd2), the mean number of alleles and its standard deviation (mean_K, sd_K), the total number of alleles (tot_K), the heterozygosity and its standard deviation from each population (H1, H2, Hsd1, Hsd2), the mean heterozygosity and its standard deviation (mean_H, sd_H), the total heterozygosity (tot_H) and population differentiation (Fst_2_1). Summary statistics were pruned to 7 (sd_K, tot_K, Hsd2, mean_H, sd_H, tot_H, Fst_2_1) for the ‘Estimation’ task of ABCtoolbox because of elevated correlations among variables in the simulated datasets.

Simulations (500,000 per run) were carried out with ‘fastsimcoal2’ (Excoffier *et al.*, 2013) and summary statistics were computed with ‘arlsumstat’ (Excoffier and Lischer, 2010). Validation techniques followed the steps detailed in the ABCToolbox wiki (https://bitbucket.org/phaentu/abctoolbox-public/wiki/Home). In brief, posterior densities of the estimated parameters were obtained by simple rejection sampling of the 1000 simulations closest to the observed data and by a post-sampling adjustment based on a general linear model (Leuenberger and Wegmann, 2010). Model choice was based on the Bayes factors from each compared model, which were obtained from their marginal densities according to Leuenberger and Wegmann (2010). Model choice validation was carried out by two statistical tests that measure the capacity of a model to reproduce the observed data. The marginal density *p*-value gives the fraction of retained simulations whose marginal density is smaller than or equal to the marginal density of the observed data. The Tukey depth *p*-value measures the fraction of retained simulations whose Tukey depth is lower than or equal to the Tukey depth of the observed data (in other words, the centrality of the data in a multidimensional space where each dimension represents a summary statistic). Small *p*-values of these tests would indicate poor fit of the model. Validation of parameters estimates was carried out by cross-validation techniques using pseudo-observed data (i.e., summary statistics of simulated datasets), and with non-parametric Kolmogorov-Smirnov tests on the posterior density distribution of parameter values.

## Results

Single locus statistics and LD values are described in Supplemental File 2.

### Population Structure

Among several Structure models, only the ‘Linkage model with independent allele frequencies’ was able to find a pattern in population structure, which grouped the 3 geographically related inter-specific population pairs into 3 common ancestors (even if support provided by the Δ*K* statistic is weak; Figures 1a-1c). Tess3r supported an ancestry model with *K*=5 groups that resulted from separating the southernmost inter-specific population pair (CJA-SNE) into two independent ancestors (Figures 1d-1f). Both methods agreed that geography was more important than phylogeny in shaping relationships among these Mediterranean white oaks, at least in the two closest inter-specific pairs of populations (CAB and IZK). Noteworthy, the Tess3r model with *K*=4 ancestral groups, joined populations as Structure did, but apportioned significant levels of admixture to 5 populations (Figure 2e). The geographic component of the global population structure showed that each population pair was derived from two different ancestors, whereas the phylogenetic component indicated that the 4 *Q. faginea* populations derived from two ancestors and that the 3 *Q. pyrenaica* populations had different ancestors (Supplemental File 3, Figure S3-1).

Serially subsampling the number of individuals per population (Figures S3-2) confirmed the stability of the Tess3r model, in contrast with the variability shown by the different Structure models. The ‘Linkage model with independent allele frequencies’ showed a relative stability with sub-samples of 36 and 24 individuals. Furthermore, the 3 repetitions of this model with samples of 8 individuals per population, were able to recover the phylogenetic signal in our data, grouping the populations according to ‘morphological species’. Inferences of two groups with other Structure models failed to recover the species signal and produced 2 ancestor groups each with populations from both species.

Truncating the dataset off the most and the least globally differentiated markers, which are the most likely to show footprints of positive and balancing selection respectively, affected the two models differently. The geographically explicit admixture model from Tess3r kept finding the same population structure with 5 ancestral groups, although admixture proportions increased significantly. Structure’s Admixture and Linkage models with independent allele frequencies recovered the phylogenetic signal in the sample, grouping the populations according to species (Figure S3-3). It follows we should find the loci responsible for creating the geographic population structure among the discarded EST-SSRs, which suggests some of them might be involved in local adaptation. Our results suggest that combining inferences from different models and datasets can help to find cryptic, not obvious relationships that might interest forest managers and evolutionary scientists.

Further study of the ‘neutral’ marker set revealed that effects on the geographic pairs were mostly restricted to the IZK populations, where residual levels of admixture were discovered in most trees from both species (Figure S3-4). An increase in shared ancestry levels was also observed when the two species were analyzed separately (Figure S3-4), but the 4 *Q. faginea* populations were assigned 4 different ancestors instead of the two ancestors obtained when all markers were used.

### Local Adaptation

The search for molecular signatures of environmental-related natural selection was carried out with a combination of population and landscape genetics approaches. We used two GS (BayeScan and Tess3r) and three GEA (BayeScEnv, Lfmm and LfmmAll) models to analyze candidate loci for local adaptations in the two biological replicates (species). We compared the results of all tests to search strong candidates into the outliers/associated loci detected by several models. Functional annotation of the candidate EST-SSRs provided, in several cases, additional evidence of their possible roles in local adaptation.

Bayescan and BayeScEnv are Bayesian models well suited to the use of FDR for providing significance levels, and they use a fixed FDR of 10% (Supplemental File 4; Table S4-1, Table S4-3). Tess3r and Lfmm-LfmmAll models need intervention to select a correct distribution of the adjusted *p*-values first, and the appropriate FDR levels afterwards (Figures S4-2, S4-4, S4-5). We selected appropriate *p*-value distributions based on the number of clusters (*K*), as the shapes of adjusted *p-*values histograms did not change with very different values of the genomic inflation factor (*λ*, from 0.01 to 100). However, most of the times we were able to select distributions of the appropriate shape for a correct FDR usage, and for the rest we favored the conservative options to reduce the false positives rate (Figures S4-2, S4-4). Another concern with our analyses could be the increased homozygosity caused by null alleles, which could be confounded with selection signatures. We tested this hypothesis with Bayescan results and found little evidence to support it (Fig. S4-1). First, the high rate of outliers common to both species confirms the similarities of the two biological replicates, in spite of general interspecific differences in null alleles frequencies. Second, 4 out of 10 common outliers by Bayescan (FIR013, PIE082, PIE137 and PIE219) did not show evidences of null alleles in any population (Table S2-1). Third, two other common outliers (POR008 and VIT023), displayed null alleles in two populations of one species but none in the other species. Fourth, another two outliers either in *Q. faginea* (FIR044) or in *Q. pyrenaica* (PIE204) did not show evidence of null alleles in any population of the respective species. Thus, it seems that influence of null alleles in the GS test, if any, is not large.

The complete results from the two GS and three GEA models are shown in Tables S4-1 to S4-6 and they are summarized in Table S4-7 (2 different columns are used for the LfmmAll modela with 27 and 10 Bioclim variables because of the large differences between them). A modified version of Table S4-7, obtained after dropping the loci detected as outliers only once within species and the LfmmAll model with 10 Bioclim variables, is shown in Table 1. Sixteen out of 30 outliers/associated loci were common to both species. Most of them were significant according to three or more models in each species, thus becoming strong candidates for local adaptation. Species-specific outliers/associated loci amounted to 6 in *Q. faginea* and 8 in *Q. pyrenaica.* Dropping Table S4-7 candidate loci according to a strict majority rule (significant in at least 3 tests per species) reduced their numbers to 21, 12 common and 2-7 species-specific in *Q. faginea* and *Q. pyrenaica* respectively. Under any of those stringencies, functional annotation of the candidate loci showed that both common and species-specific groups include metabolic, signal perception, regulatory and response functions (Tables 2a-2b).

**Table 1:**
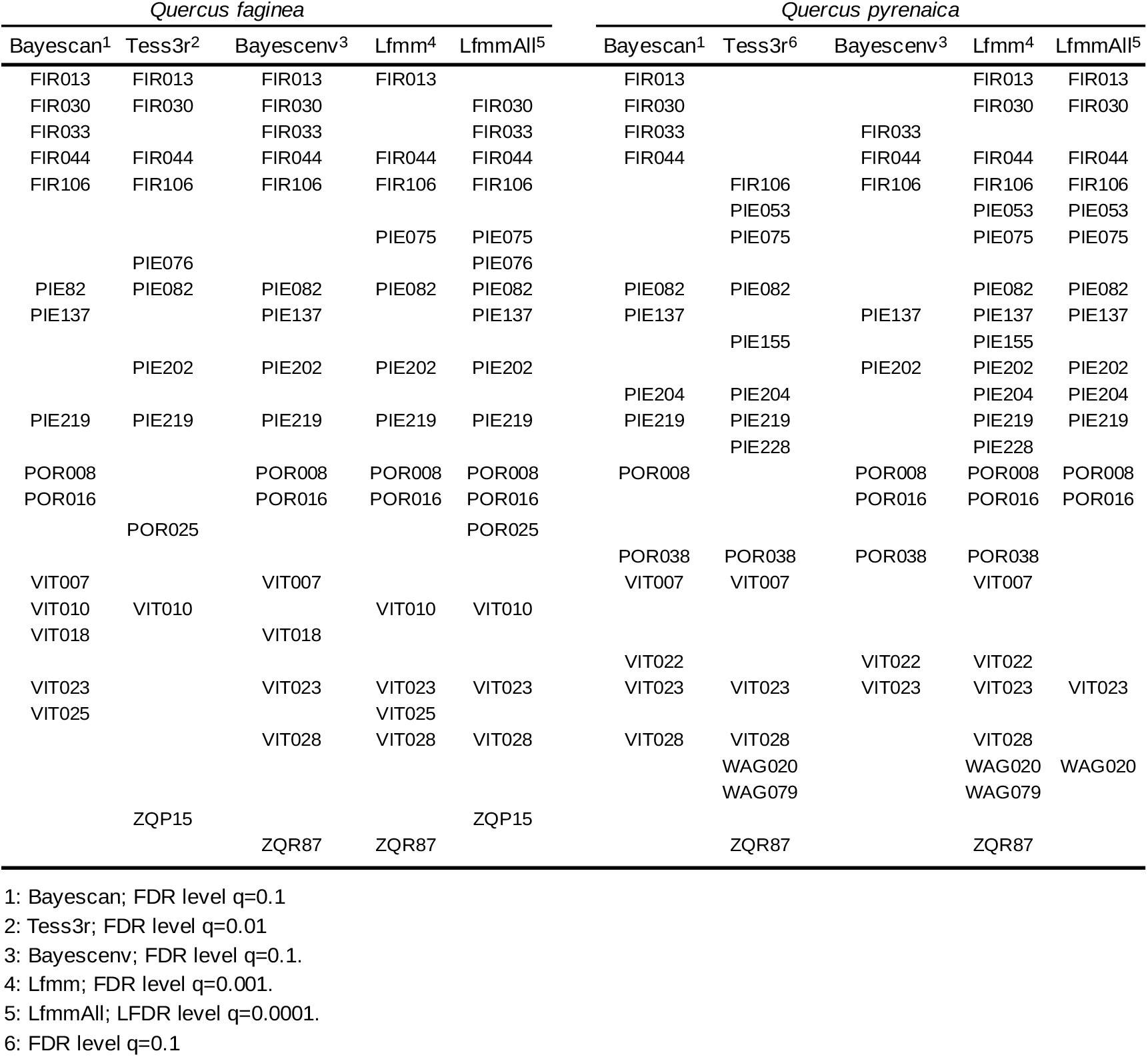
Comparison among GS outliers and loci involved in GEAs detected by the different models used in the study. Loci listed only once within species have been dropped.

**Table 2a:** Functional annotation of 16 local adaptation candidates shared by the two oak species.

| SeqName | Description | Length | #Hits<sup>†</sup> | e-Value | Gos and GeneRIFs |
| --- | --- | --- | --- | --- | --- |
| FIR013<br>FP026253.1 | B-box type zinc finger with CCT domain-containing (CO) | 711 | 6 | 1.18E-38 | P:far-red light signaling pathway;<br>P:circadian rhythm;<br>F: DNA-binding transcription factor activity |
| FIR030<br>FP056086.1 | NADH dehydrogenase ubiquinone 1 Alpha subcomplex subunit | 542 | 1 | 6.73E-26 | C:mitochondrial respiratory chain Complex 1 |
| FIR033<br>FP030475.1 | SAM domain containing | 824 | 4 | 9.80E-67 | F:phospholipase activity<br>Glycosyl groups |
| FIR044<br>FP066528.1 | zinc finger B-box (AT5G45410) | 783 | 2 | 1.83E-92 | P: defense response to bacterium;;<br>C:chloroplast; |
| FIR106<br>FP065330.1 | AAA-type ATPase family protein | 634 | 1 | 7.91E-55 | P:SNARE complex disassembly;<br>F: ATPase activity |
| PIE075<br>FP073223.1 | ralf-like 33 | 768 | 8 | 3.48E-37 | P:calcium-mediated signaling;<br>P: root development;<br>P: pollen tube growth;<br>F: hormone activity |
| PIE082<br>FP044493.1 | Ubiquitin-like superfamily protein | 679 | 1 | 1.00E-28 | P:response to UV-B |
| PIE137<br>FP050882.1 | PHYTOSULFOKINE 3 PRECURSOR | 599 | 1 | 1.97E-13 | P:cell differentiation;<br>P:cell proliferation;<br>F:growth factor activity; |
| PIE202<br>FP055682.1 | O-methyltransferase family protein | 643 | 11 | 2.89E-59 | GeneRIF: miRNAs degradation<br>P:melatonin biosynthetic process;<br>F: caffeate O-methyltransferase activity |
| PIE219<br>FP026114.1 | serine carboxypeptidase-like 22 | 830 | 20 | 8.44E-30 | P:brassinosteroid mediated<br>Signaling pathway;<br>P: cell wall proteolysis |
| POR008<br>FP038868.1 | prenylated RAB acceptor 1.H | 650 | 1 | 3.21E-54 | P:vesicle-mediated transport;<br>C: endoplasmic reticulum membrane |
| POR016<br>FP046659.1 | Tetratricopeptide repeat (TPR)-like Superfamily protein | 624 | 20 | 1.05E-83 | P:response to osmotic stress;<br>P: defense response;<br>F: phosphoprotein phosphatase activity |
| VIT007<br>FP067286 | BTB-POZ and MATH domain 2 | 328 | 8 | 2.22E-23 | P:response to water deprivation;<br>P:response to salt stress;<br>F: identical protein binding |
| VIT023<br>FP067992.1 | Integrase-type DNA-binding superfamily | 711 | 20 | 8.87E-26 | P:response to salt stress;<br>P:response to water deprivation;<br>F: DNA binding transcription factor activity |
| VIT028<br>FP072685 | transcription factor MYB44-like | 579 | 14<sup>a</sup> | 2.40E-19 | F: DNA binding |
| ZQR87<br>Q. robur V2-2N<br>Scaffold_0906 | rab9 effector protein with kelch motifs-like (Q. suber)<br><sup>8</sup>F-box/kelch-repeat protein At1g51550-like (Carica papaya) | 300 | 20<sup>a</sup> | 9.00E-16 | P: circadian rhythm<br>P: response to blue light;<br>F: ubiquitin-protein transferase |

†: All sequences blastx against *Arabidopsis thaliana* proteins, except: "a" blastx against seed plants proteins.

**Table 2b:**
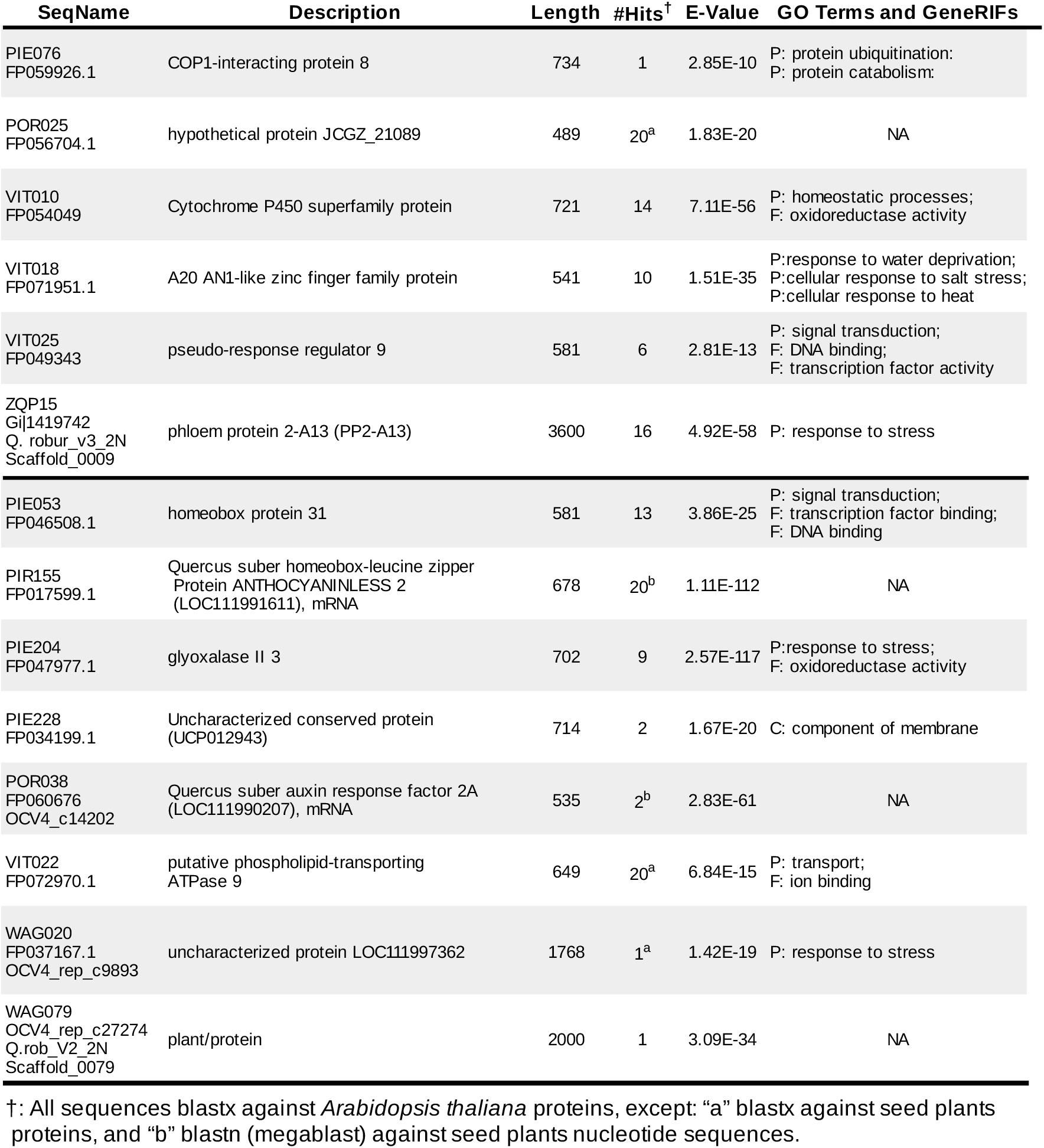
Functional annotation of the 14 species-specific local adaptation candidates (6 in *Quercus faginea*, top panel; and 8 in *Q. pyrenaica*, bottom panel).

### Adaptive Introgression

Allele frequencies from the 16 common candidate loci in Table 1 show that outlier/associated alleles are shared within the three inter-specific population pairs, 6 by the CAB populations, 4 by the CJA-SNE populations and 6 by the IZK populations (Supplemental File 5, Figure S5-1). Two introgression events were inferred for outliers FIR013 and FIR044, both in population pairs CAB and IZK, whereas POR016 showed private outlier alleles in populations CJA-SNE. Only POR016 and VIT028 showed allele frequencies that were not easily interpreted in terms of adaptive introgression. Note that all 4 common outliers affected by the strict majority rule (FIR033, PIE075, VIT007 and ZQR87) passed this first introgression screen.

Comparisons of the bootstrapped ASD values within pairs of populations are consistent with introgression of the candidates in populations CABfg-CABpy and IZKfg-IZKpy (Figure S5-2), as the smaller ASD values translate into shorter coalescence times. Populations CJAfg-SNEpy illustrate this test bias when one of the compared groups contains a small number of loci. One of the four candidate loci from this populations (FIR030) showed a large ASD, and bootstrapped values including several instances of this marker stood out with very large distances (Figure S5-2).

We sought further evidence against ILS in markers adjacent to the EST-SSR outliers. In spite of anticipated difficulties, we found putative introgressed alleles from adjacent loci within each of the three interspecific population pairs (Figure 3). The inferred double introgression of FIR013 in population pairs CAB and IZK contained adjacent introgessed alleles from a different flanking locus in each population pair (FIR048 and ZQR87 respectively), but both introgression instances ocurred into *Q. faginea.* Introgressed alleles from markers flanking FIR044 could be inferred only in populations CAB, although the direction could not be determined because the outliers and flanking alleles were geographically private. The inferred introgression of PIE082 in populations CJA-SNE was further confirmed by introgression of alleles from both flanking markers (VIT043 and PIE216) into *Q. pyrenaica.* Finally, the inferred introgression of VIT007 in the IZK populations was joined by introgression of alleles from adjacent locus WAG011 into *Q. faginea.* Note that some residual frequencies of putative introgressed alleles from adjacent loci were allowed based on the possibility of introgression in more than one population pair (e.g., FIR013), together with high mutation rates and homoplasy of the EST-SSR markers.

Fourth, we tested the adaptive nature of introgressed alleles at candidate loci by searching for disruptive selection signatures with two marker haplotypes within each species (Figure S5-3), i.e., two-marker haplotype frequencies from each population of the candidate pair were compared to haplotype frequencies in the remaining populations from the same species. Highly significant associations from common haplotypes in both within-species groups (candidate population for adaptive introgression vs. other populations from the same species) suggest different allelic combinations are selected in contrasting environments, i.e., disruptive selection among populations, within species.

**Figure 3:**
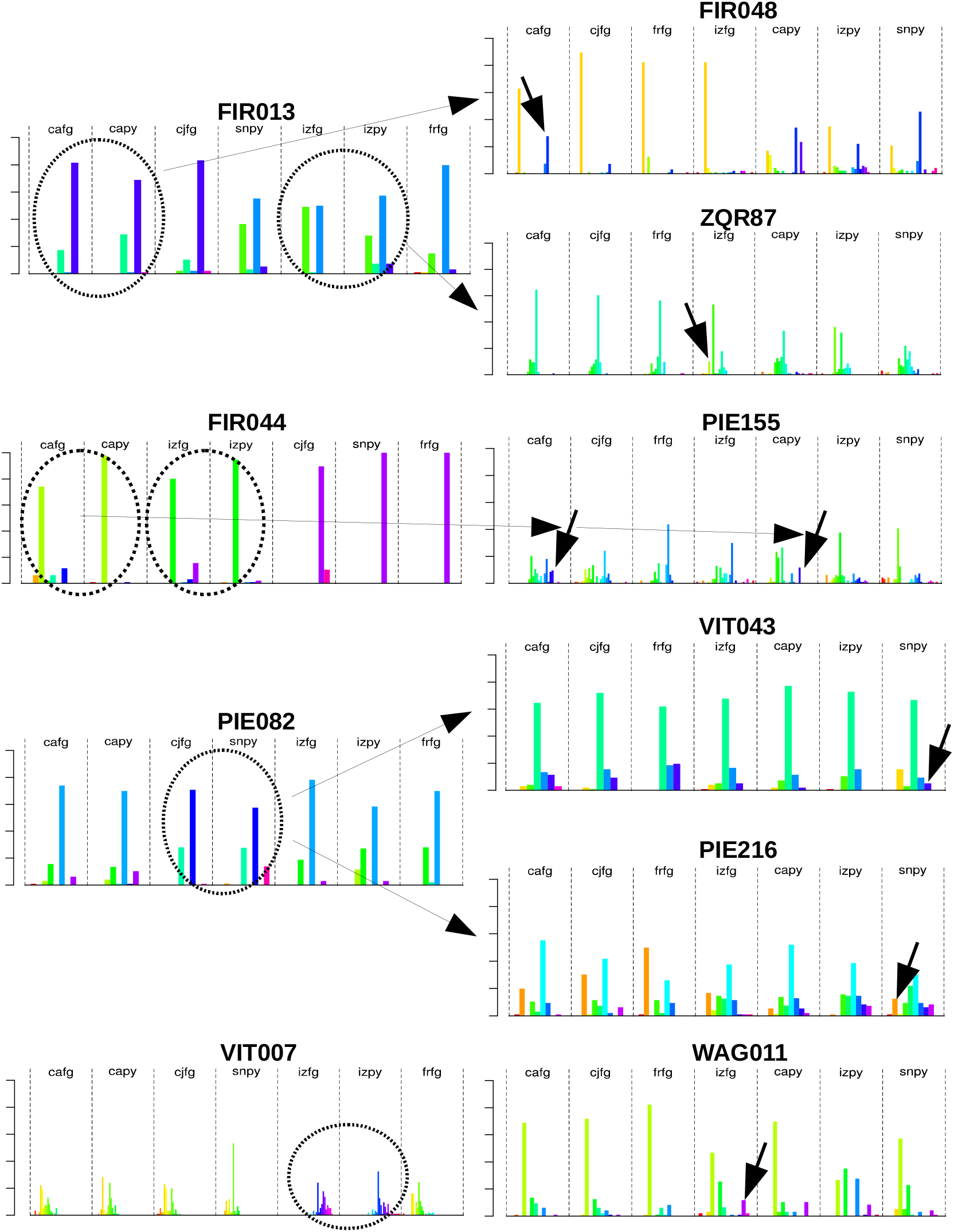
Introgressed alleles from candidate loci for local adaptation (left pane) and from flanking markers (right pane) in the three inter-specific population pairs. Dotted ovals signal the introgressed adaptive alleles. Arrows in the right pane signal towards ‘species-specific’ alleles introgressed in the focal population from the other species. Note that FIR044 introgressed alleles at candidate loci and flanking markers are geographically private, thus impeding to infer the introgression direction and not allowing to exclude incomplete lineage sorting.

One final step in the characterization of hybrid introgression in the two Iberian oaks was the functional annotation of the candidate markers. FIR013 was derived from several oak EST clones (NCBI sequence FP026253) with best blastx hits to the *Arabidopsis* CONSTANS-2 gene, thus eliciting the marker as one of the two known oak orthologs to the *Arabidopsis* CONSTANS gene (CO). CONSTANS is a regulatory RNA best-known for activating FLOWERING LOCUS T and promoting flowering in a large variety of plants (Griffiths *et al*., 2003; Valverde, 2011). In trees, CO participates not only in the photoperiodic flowering pathway, but it is also involved in Autumn growth-cessation and Spring bud burst (Böhlenius *et al.*, 2006; Maurya and Bhalerao, 2017). It would be tempting to associate this exonic EST-SSR to differences in flowering time between the two oak species. However, the two CAB populations contain the same COL2 alleles at similar frequencies but showed bud phenology differences amounting to nearly 1 month (our own observations at sampling time), which cast severe doubts on the function of the putative COL2 oak gene. Indeed, several molecular pathways collaborate to promote flowering in higher plants (Moon *et al*., 2005; He, 2012), which could provide a myriad explanations to the phenology differences in the absence of data from the complex genetic network that regulates flowering. Or else, the COL2 oak gene could be involved in other adaptive function(s) not related to phenology, such as regulation of abiotic stress tolerance through an abscisic acid dependent manner (At-COL4, Min *et al.*, 2015), or regulation of branching patterns and shade avoidance (At-COL7, Wang *et al.*, 2013). On the other hand, a similar COL2 polymorphism has been identified as species outlier in allopatric populations of the North American red oaks *Quercus rubra* and *Q. ellipsoidalis* (Lind-Riehl *et al.*, 2014). In sympatry, though, soil quality partially drives asymmetric introgression of the drought-tolerant *Q. ellipsoidalis* alleles into *Q. rubra* (Khodwekar and Gailing, 2017), which it is opposite to the soil requirements and introgression direction of this locus in the Iberian oaks. Indeed, functional characterization of the putative oak COL2 gene (e.g., complementation of *Arabidopsis* CO null mutants) might be needed to assess the gene function and putative adaptive nature of the introgressed alleles. The second candidate for adaptive introgression, PIE082, is ortholog to an *Arabidopsis* ubiquitin family protein putatively involved in the response to UV-B light (Wolf, 2011). Adaptive introgression of this gene has been inferred in the pair CJAfg-SNEpy, the two oak populations that grow at highest altitude and are exposed to highest solar radiation intensities, further supporting an adaptive role of the PIE082 alleles. Finally, VIT007 has been identified as ortholog to an *Arabidopsis* BTB-POZ and MATH domain 2 gene that is involved in the responses to salt and osmotic stresses and to water deprivation (Weber and Hellmann, 2009). Adaptive introgression of this gene has been identified in the IZK populations, which contrary to all other populations grow under nearly temperate climate conditions with likely winter snow and only sporadic drought periods

### ABC of Divergence with Gene Flow

Dating introgression events, or at least putting some time limits to them, was studied with coalescent simulations coupled to ABC model selection using Bayes factors (Supplemental File 6). Genealogies without gene flow or without divergence (SI and PAM models) were discarded soon and we concentrated on models of divergence with continuous gene flow (IM), divergence with gene flow that finished some time in the past (AM), and divergence with gene flow from secondary contact to present (SC). Model choice validation tests didn’t reject the best ABC models for the three inter-specific population pairs (Table S6-1). However, these p-values do not confirm hypothesis but only reject them, and they can be seen as filters that reject the models that do not fit the data (ABCtoolbox wiki). Further, best models poor fits of cross-validation and Kolmogorov-Smirnov tests (Figures S6-2, S6-3) indicated that parameter estimates were biased, i.e. low accuracy of the ABC-GLM model (ABCtoolbox wiki). Thus, we modified the models to incorporate source-sink gene flow from central to marginal populations (Figure S6-1 and Table S6-1 models IMc, AMc, SMc). We obtained better Bayes factors in two population pairs, but the accuracy of the ABC-GLM models kept low and parameter estimates biased. Reasons for such bias can be multiple, among others, a demographic history more complex than the simulated ones; e.g. including expansions and declines, extinctions and re-colonizations, or variable migration rates (Slatkin and Excoffier, 2012; Excoffier *et al.*, 2013; Peischl and Excoffier, 2015), differential interactions with other species from the same radiation (Fraïsse *et al.*, 2016), the local effects of RI barriers of variable strengths on the variance of gene flow rates across the genome (Roux *et al.*, 2016), or variable recombination rates across the genome (Burri *et al.*, 2015).

While low accuracy of parameter estimates recommends caution, best models Bayes factors strongly support ongoing inter-specific gene flow within the three population pairs. Further, a recent secondary contact is supported for the pairs CJA-SNE and IZKfg-IZKpy, while continuous inter-specific gene flow since species split is supported for the CAB populations; i.e., sympatric speciation (Table S6-1). Sequential model choice also supported intra-specific gene flow from ‘central’ to marginal populations in the population pairs CAB and IZK, at least with our parameterizations; whereas the comparison of best models in the CJA-SNE populations (SC vs. SCc) favored the model without intra-specific gene flow from ‘central’ to marginal populations (Table S6-1). As we discuss below, these inferences are supported by independent evidences, too.

## Discussion

Our study follows previous work on interspecific genetic differentiation among range-wide populations of the two Mediterranean white oaks, *Quercus faginea* Lam. and *Q. pyrenaica* (Matt.) Wild. In the present study, we shifted the focus towards genetic variation within species’ marginal populations, and to their associations with environmental variables (although both genetic structure and local adaptation revealed interspecific interactions). Starting from careful sampling considerations, we show that (1) the combination of EST-SSRs allele frequencies and *F*_ST_-based population and landscape genetics models can reliably inform the search of genes that are important for local adaptation, and (2) adaptive introgression of large-effect alleles lies at the base of local adaptations in marginal populations of the two Mediterranean white oaks, granting fast adaptation to extreme environments and favoring range expansions. This mutualistic biotic interaction provides new insights on population genetics at the edges, and supports the conservation value of marginal populations from sister-species. In addition, we show that, (3) coalescent simulations coupled to ABC model selection support the hypothesis of divergence with gene flow in the three interspecific population pairs, but a recent secondary contact was inferred in populations affected by contraction/expansion linked to Quaternary climatic oscillations whereas old continued introgression was inferred in populations from a refugia area with little climatic oscillationsand (4) population structure has a strong geographic component that is related to adaptive introgression, but the underlying neutral phylogenetic signal can be recovered by genotyping a few individuals per population, even if several markers are under heterogeneous natural selection.

### Mutualistic Introgression

Forest trees marginal-peripheral populations have attracted much attention during the last decade. The central-marginal hypothesis anticipates that peripheral populations exhibit lower genetic diversity and greater genetic differentiation than central populations due to smaller effective population size and increased geographical isolation, which has been confirmed to a certain degree (e.g., Eckert *et al.*, 2008), in spite of notable exceptions (e.g., Yakimoski and Eckert, 2008). Further, edge populations at the limits of the geographical/ecological ranges are expected to harbor adaptive alleles for heterogeneous natural selection at the range margins and beyond (Roschanski *et al.*, 2016, but see McLane and Aitken, 2012), although increased phenotypic plasticity might play a similar role (Chevin and Lande, 2011). Thus, there is little doubt that marginal populations are important not only to applied genetics, such as germplasm conservation and/or assisted strategies to counter climate change (Hampe and Petit, 2005; Petit *et al*., 2008), but also to theoretical developments in evolutionary and population genetics, such as range expansion and local adaptation (Bridle and Vines, 2007; Kremer *et al.*, 2012; Savolainen *et al.*, 2013).

In this work we show how ancient and recent introgression promotes local adaptation in sister-species marginal populations from the European white oaks syngameon. In spite of contributing only a qualitative vision of the process (quantifying introgression in oaks is now possible with available genomic assets and cheapening massive-genotyping technologies; Leroy *et al.*, 2019), we show it likely occurs in the three studied population pairs, suggesting it might extend along the contact areas between the two species. The main evolutionary benefit from this process is a fast acquisition of adaptive alleles already tested by natural selection (Abbott *et al.*, 2013), which can drive adaptation at a faster pace than standing variation when introgression is iterative (Hedrick, 2013). Further, we have shown that adaptive introgression in these oaks is a G×G×E interaction that occurs in both directions, the two species forming a rarely described geographic mosaic of mutualistic co-evolution (Thompson, 2005). This type of biotic interaction is likely to occur in other radiations too (e.g., white oaks of the Eastern and of the Southwestern United States and Mexico, Boecklen, 2017; fruitflies from America and Australia, Bergland *et al*., 2016), conferring an explicit evolutionary advantage to syngameons.

Adaptive hybridization/introgresion at marginal populations has been recognized as one of the main drivers of species’ range expansions (Rieseberg *et al*., 2003; Arnold, 2004; Abbott *et al.*, 2013) and it has been acknowledged, together with rare long-distance dispersal events, as two main drivers of oaks successful northward colonization of Europe after the last glaciation (Kremer, 2016). Adaptive introgression can contribute to solve the two main genetic risks leading marginal populations to face extinction, (i) the small effective population sizes, with small genetic and phenotypic variances, which might cause populations to collapse due to inbreeding; and (ii) the lack of adaptive alleles for local conditions, which might cause poor recruiting each new generation, populations becoming sinks rather than sources of dispersers (Dias, 1996). Rescuing edge populations by new mutations it’s unlikely because the lag time for favorable mutations is long, specially if adaptation depends on many genes of small effects (Barret and Schluter, 2008; Abbott *et al.*, 2013), whereas gene-flow from conspecific populations frequently consists of maladaptive alleles from central populations that often counter local adaptation (Bridle and Vines, 2007; Sexton *et al.*, 2009). Thus, adaptive introgression with resident relatives might become the best alternative to the supply of adaptive alleles to edge populations (Grant and Grant, 1994; Arnold and Martin, 2009), even for small-effect alleles from multiple loci if repeated introgression occurs during the transient LD phase (Abbott et al., 2013, Yeaman, 2015). Further, introgression might have an indirect adaptive effect by increasing effective population sizes and overall genetic diversity, which could increment survival chances until new favorable mutations arise or gene flow brings new adaptive alleles (Pfenning *et al*., 2016; Pierce *et al.*, 2017). Our study contains a wealth of data to support the hypothesis of direct introgression effects through the transfer of adaptive alleles that have been already tested by natural selection in the genetic context of a sister species. Although our experimental design and analysis methods are not totally appropriate to search indirect effects evidences (Pierce *et al.*, 2017), we would expect different adaptive alleles in the two species from each pair if indirect effects are predominant. However, our data show that adaptive loci shared between inter-specific population pairs also share the EST-SSR alleles, which suggests that local adaptation in marginal populations occurs without extended lag-times. Similarly, Europe’s fast colonization after the last glaciation, suggests adaptive alleles to northern conditions were already present in the front-end populations, and could extend into sister species by means of introgression.

An interesting follow-up to hybridization’s role in range expansions is the limitation imposed by genome swamping and the collapse of the focal marginal population into the resident species (Behm *et al.*, 2010; Kearns *et al.*, 2018). Three main solutions have been proposed to the problem of maintaining species boundaries in the face of introgression (Pfenning et al., 2016): i) hybridization might carry high costs but be a rare event, ii) hybridization could occur only during the first stages of contact, and iii) hybridization might generate fitness trade-offs, or might represent the “best of a bad situation”. The two oak species analyzed in this study suggest a mechanism to avoid genome swamping that could be made extensive to other biological systems: ecological speciation coupled to introgression at unrelated adaptive traits. Ecological discrimination between the two Iberian oaks is based on soil pH (Blanco-Castro *et al.*, 1997), which suggests an excellent candidate character to drive ecological speciation, and we have previously reported partial evidence that phenology might also contribute to ecological speciation between *Quercus faginea* and *Quercus pyrenaica* through assortative mating (Goicoechea *et al*., 2015). Adaptive introgressions described in this study involve loci governing environmental traits unrelated to soil pH, such as exposure to UV radiation or arid-temperate climate adaptations. Yet, to fully explain the maintenance of species barriers without the formation of hybrid swarms some kind of reinforcement is needed. This might be accomplished by phenology-driven assortative mating (Abadie *et al*., 2012; Lepais *et al*., 2013; Goicoechea *et al*., 2015), because oak’s asynchronous earlier masculine than feminine flower development allows hybrids to back-cross one parent or the other depending on their roles as pollen donors or recipients. In this case, a few admixture generations could replace the close linkage between genes governing soil adaptations and leaf/flower morphological characters, or else epistatic interactions, that is needed to explain association between ecological discrimination and taxonomic characters.

### Challenges To Local Adaptation Inferences

Using marginal populations to analyze the genetic footprints of local adaptation has some advantages but some drawbacks too. On the positive side, peripheral populations are prone to be under environmentally driven heterogeneous selection that is necessary for local adaptation to occur (Gavrilets, 2004). On the negative side, theory predicts that the genetic and demographic characteristics of marginal populations counteract local selection with varying intensities, causing ‘maladaptation’ sometimes, enhancing the evolutionary potential of small populations others (Kawecki, 2008; Roesti, 2018). Further, lasting patterns of genetic differentiation can arise and be maintained under a variety of scenarios in addition to differential effective gene flow between ‘permeable’ and ‘non-porous’ regions of the genome (Hoban *et al*., 2016; Wolf and Ellegren, 2017), although temperate oaks classical provenance trials have demonstrated local adaptations for phenology and growth traits in natural populations (Ducousso *et al*., 1996; Alberto *et al*., 2013) and molecular markers have identified several genes involved in adaptation to environmental conditions (Rellstab *et al.*, 2016; Leroy *et al*., 2019).

The geographic population structure (Figure 1) closely associated to the three main environments (arid-Mediterranean: CAB, high altitude-Mediterranean: CJA-SNE, temperate-like: IZK), together with recovery of the phylogenetic signal when the most and the least differentiated markers were excluded (Figure S3-2), were the first insights into the potential of GSs and GEAs to search loci that are important for local adaptation. The extraordinary concordance among 5 GS and GEA models, partly replicated between species (Table 1) and confirmed by the inferred introgression events (Figure S5-3), is persuasive evidence in favor of Table 1 candidates, even taking into account that the variances of our *F*_ST_ estimates might be raised as a consequence of high neutral differentiation (Figure S2-6).

Nevertheless, the combination of Bioclim variables and GEA models showed several results that challenge their usefulness in our data. First, the two GEA models that fit environmental variables each at a time (BayeScEnv and Lfmm) found extra candidate loci, in both species, among the highly correlated Bioclim variables (Table S4-7). On the contrary, the only model that fits all Bioclim variables at once (LfmmAll) found highly significant GEAs for the 10 ‘uncorrelated’ Bioclim variables that were not detected when all 27 Bioclim variables were fit (Table S4-7). Comparison among all GS-GEA models showed that LfmmAll was more related to the rest when all 27 Bioclim variables were fit than when only the 10 ‘uncorrelated’ variables were used. Whereas these results mean just a little inconvenience for BayeScEnv and Lfmm (as highly correlated Bioclim variables can’t be discarded), results from LfmmAll need further investigations that fall outside the scope of this study.

Second, inter-specific comparisons of candidates detected by BayeScEnv and Lfmm, using only the 10 ‘uncorrelated’ Bioclim variables for the sake of clarity, showed that the first GEA model found only 2 EST-SSRs associated to the same Bioclim variable in both species, whereas Lfmm showed 12 EST-SSRs associated to both species that involved 8 out of 10 Bioclim variables (Figure S4-6a). From an alternative point of view, the comparison of the two GEA models within each species showed common EST-SSRs associated to 6 out of 10 Bioclim variables in *Q. faginea* and to 7 out of 10 Bioclim variables in *Q. pyrenaica.* However, associations exclusive to each GEA model largely exceeded the common associations in both species (Figure S4-6b).

Third, we counted the number of Bioclim variables each of the 16 shared candidates was associated with, in each species (Figure S4-7). BayeScEnv found 7 candidates exclusive to *Q. faginea*, two of them associated to more than 10 Bioclim variables, whereas Lfmm found 4 candidates exclusive to *Q. pyrenaica*, one of them associated to 17 and two other associated to 8 Bioclim variables. One extreme case that exemplifies the interactions between GEA models and species was VIT007, which appeared as a BayeScEnv candidate associated to 8 Bioclim variables exclusively in *Q. faginea*, but as a Lfmm candidate associated to another 8 Bioclim variables exclusively in *Q. pyrenaica* (Figure S4-7).

In spite of all these examples, there is enough evidence to support that the combination of *F*_ST_-based GS and GEA models can reliably identify local adaptation candidates based on allele frequencies from EST-SSRs (Table 1, Figure S5-1). Indeed, the comparison of Table S4-7 and Table 1 suggests that false positives are appropriately controlled with our approach, whereas between model differences within the shared candidates, with similar allele frequencies patterns in the two species, question the power of the two studied GEA models to avoid false negatives. On the other hand, the different number of populations from the two species, and the environmental differences between IZKfg-IZKpy and specially between CJAfg-SNEpy are expected to cause inter-specific variation in candidates. Thus, the question becomes whether these environmental differences can explain all the within GEA models inter-specific variation or just a part of it. We acknowledge that our sampling design might not be the most appropriate to answer this question and that proving the usefulness of Bioclim variables, and hence environmental/niche modeling, as predictors of genetic differentiation linked to local adaptation might require the sampling of clines to take full advantage of the large environmental resolution provided by Bioclim variables. However, our results warn against the uncritical use of Bioclim variables for SDM/niche modeling and they suggest further research should be carried out.

We welcome the proliferation of good practices and road-maps for conducting and interpreting GSs and GEAs to assist the detection of molecular footprints from natural selection linked to local adaptation. In fact, our methods considered several of the most important suggestions in the literature, such as genotyping large sample sizes per population, using paired populations from two sister species to avoid idiosyncratic effects of introgression, and controlling the confounding signals of demography and population history, either with the population-specific parameter *β* in BayeScan and BayeScEnv, or with latent factors in Lfmm-LfmmAll (Lotterhos and Whitlock, 2015; Fraïsse *et al.*, 2016; Hoban *et al.*, 2016). Explicit IBD estimates (Nadeau *et al.*, 2016) were not necessary in our analysis because they contribute solely to global population differentiation. Further, we discarded the effects of correlations among genetic markers and environmental variables because environments are not clinal but patchy (see the negative auto-correlations in Figure S2-4), and the existence of geographic homogeneous selection was not a concern after we careful selected the sampling environments. Most important confounding effects not addressed in our study are linked selection, recombination variation and reduced genome representation (Tiffin and Ross-Ibarra, 2014; Burri *et al.*, 2015; Hoban *et al.*, 2016; Ravinet *et al.*, 2017), although the last might not be a concern if local adaptations to highly divergent environments affect many genes (Sork, 2017; Ahlstrand *et al.*, 2018; Tabas-Madrid *et al.*, 2018).

Polygenic traits with complex genetic architectures are predicted as another challenge for *F*_ST_-based GSs and for GEA models (Le Corre and Kremer, 2012; Yeaman, 2015), specially when neutral differentiation is high (Pérez-Figueroa *et al.*, 2010) and when selection is weak in comparison to gene flow (Beaumont and Balding, 2004). However, we have shown that our populations show high neutral differentiation (Figures S2-6) and that migration from central to marginal populations is relevant at least in populations CAB and IZK (Table S6-1), yet we were able to find strong candidates for various cellular functions underlying the successful adaptation to extreme environments (such as a cellular machinery highly efficient at elevated temperatures, an efficient redox mechanism to overcome hyper-osmotic stress, a sensitive glucose metabolism, a complex regulatory system, and so on). The reason for this disagreement might dwell at the definition of ‘polygenic trait with complex genetic architecture’, because adaptation to an extreme environment probably becomes a binary trait with a selection coefficient *s*=1 (survival:death), within the collection of phenotypes and genotypes from our ecologically distant marginal populations. Thus, sampling from very different environments could become a strategy to overcome the predicted difficulties to detect local adaptation candidates linked to polygenic traits with complex genetic architecture in tree populations with *F*_ST_-based GS an GEAs.

### Independent Evidences for ABC-Supported Coalescent Models

Among several divergence with gene flow tests, coalescent simulations of models with and without gene flow, coupled to ABC model selection with Bayes factors, have increased popularity during the last years (Nosil, 2012). As a consequence, simulation models have advanced to take into account several confounding factors, including among-locus variation in mutation and recombination rates, genetic drift and introgression, and even linked background selection (Excoffier *et al*., 2013; Elyashiv *et al.*, 2016; Roux *et al.*, 2016; Leroy *et al.*, 2018). These advancements rely on DNA sequence characteristics and their application to SSR data has not been sufficiently tested. We selected ABCtoolbox among available frameworks for microsatellite analysis (Wegmann *et al*., 2010; Excoffier *et al*., 2013, Cornuet *et al*., 2014;) because it allows to take into account among-locus variation in mutation and recombination rates (using ‘fastsimcoal2’), to select best models with Bayes factors, and because it provides validation tests for model choice and model fit (Wegmann *et al*., 2010). Best models and parameterizations passed model choice validation tests but failed model fit validation tests, which cast some doubts about their capabilities to describe the demographic and gene flow histories of the three interspecific population pairs. This isn’t unexpected since, in addition to some confounding factors named above, we have not considered demographic events related to Quaternary climate oscillations and associated changes in gene flow frequencies, possible admixture/hybridization events with other species, or genetic linkage among markers. However, selected models agree well with other independent sources of knowledge, as we discuss next.

Populations CABfg-CABpy grow in an area usually considered as refugia during Quaternary glaciations (Hewitt, 2000; Brewer *et al*., 2002), subject to minor climate oscillations that should allow a constant (co)-evolutionary history since populations from both species first colonized this area. This agrees with an almost pure IM model, with populations from both species exchanging genes since populations within species differentiated, shortly after speciation took place, till present; such as it was inferred from our simulations. Furthermore, these populations survive at the boundaries of their respective distribution areas, which supports the source-sink model of intra-specific gene flow with the respective central populations that was also inferred in our analyses.

Marginality of the IZK populations also supports the intra-specific source-sink gene flow model for both populations, whereas their northern latitude supports expansion/retreat or colonization/extinction dynamics that agree with a recent secondary contact. Further support to current gene flow between these populations comes from the oaks morphological and taxonomic diversity in the area between the two ‘pure-species’ populations, which suggests it is a hybrid zone maintained by ongoing gene flow and divergent selection for soil quality (Uribe-Echebarría, 2001).

Finally, the location of population SNEpy within the main distribution area of *Q. faginea* and far away from the distribution area of *Q. pyrenaica*, could be explained by two alternative hypothesis, either (i) a rare, very long distance dispersal event, or (ii) a gradual, step-by-step ‘colonization by hybridization’ (Petit *et al.*, 2004), of small mountain areas with acid soils bassets within the predominant calcareous landscape in southern Iberia. We can’t discard the first hypothesis because any footprints of such old strong bottleneck on diversity and allelic richness would be wiped out by time and because pollen records in the area are very recent (Brewer *et al*., 2002; Anderson *et al.*, 2011). However, the distribution of oaks cpDNA haplotypes within the Iberian peninsula (Olalde *et al.*, 2002) and the introgression direction detected in this study suggest the step-by-step ‘colonization by hybridization’ is a more parsimonious hypothesis, and it could explain the population remnants and scattered *Q. pyrenaica* individuals along the way from the *Q. pyrenaica* main distribution area to Sierra Nevada mountains. On the other hand, population expansion/contraction dynamics in the rear end mountain tops are contrary to northern populations (i.e., expansion during glacial periods and contraction during the interglacials), which could facilitate the blend of the CJA-SNE gene pools during the Würm (110,000-10,000 years ago) and thus explain the inference of a recent secondary contact in these populations (SC model, Table S6-1). Furthermore, long-distance isolation of SNEpy from the central *Q. pyrenaica* distribution area supports a model without source-sink intra-specific gene flow in this population. ABC inferences favored the model without any central population (Table S6-1), although a model with just one central population (corresponding to *Q. faginea*) seems more appropriate.

### Population Structure

Genetic population structure measures differentiation within and among populations, a variable quantity not only because it depends on the number and identity of populations, but because populations themselves are dynamic entities that change along time due to stochastic and deterministic processes. The enormous success of this parameter during the last 20 years of Population and Evolutionary Genetics must be explained by theoretical developments initiated at the end of the last century that allowed to estimate the proportions of individual genomes originating from *K* ancestral populations and the corresponding ancestral genotype frequencies, making thus possible to calculate an *F*_ST_ analogue measuring differentiation of each population from their ancestral ones (Pritchard *et al*., 2000). Structure’s model-based Bayesian clustering method soon became the standard, and it has maintained a predominant position since then (Novembre, 2016). However, the model has assumed Hardy-Weimberg equilibrium over successive improvements, a condition that is not always met in real data. Further, the subtle footprints of recent adaptive introgression at ecologically relevant genes might be difficult to discern within the phylogenetic background.

Throughout this study we have considered phylogenetic and geographic signals in population structure, although it is clear by now that the phylogenetic signal is an old genome-wide reminder of a common ancestor, whereas the geographic signal is the result of adaptive introgression at environment-related genes between geographically structured populations from sister species. Previous range-wide studies of inter-specific differences within the European white oaks had shown the persistence of a phylogenetic signal in paired interspecific samples (Goicoechea *et al.*, 2012; Goicoechea *et al.*, 2015), in spite of null-alleles and H-W disequilibrium in an important proportion of markers, and evidences of natural selection acting at several genes. Those studies prioritized the number of populations in ample distribution areas over the number of samples per population, whereas present study focus on environmental adaptations and adaptive introgression at species range limits requires a larger intra-population sampling that we have quantified by serial sub-sampling. Structure’s models with uncorrelated alleles frequencies needed all 48 individuals per population to converge into the solution revealing geographically structured introgression, sample sizes of 8 individuals per population converged to the solution revealing inter-specific differences, and intermediate sample sizes lead to non-convergent solutions, often indicating no relationships among the 7 populations. We are aware that it is possible to average different Structure runs into a unique solution (e.g, Jakobsson and Rosenberg, 2007; Verity and Nichols, 2016), but these methods should be applied when the different runs for a given *K* converge into the same solution, not to hide different local minima that could indicate complex population structures. Indeed, the ‘neutral’ markers failed to reveal local adaptation and introgression, converging to the inter-specific differences solution. Therefore, both types of sub-sampling (individuals and markers) can be used to search for the neutral genetic structure, although deciphering whether markers are under natural selection might be difficult, or even impossible, in many studies carried out with just a few markers.

An interesting result concerning the possibility to detect introgression of a few genes by means of population structure analysis is the grouping of *Q. faginea* populations (Fig. S3-1a). The common ancestor for CABfg and CJAfg finds a logic on the basis of close geographic location and partially similar environmental adaptations. But the relationship between populations FROfg and IZKfg is not easily conciliated with geographic, phylogenetic or climate data. The two populations are too far apart to assume any direct gene flow between them, they belong to distinct maternal lineages extending into different phylogeographic areas (Olalde *et al*., 2002), and the current study shows that climate at both locations is rather different. An hypothesis to conciliate the inferred population structure with these observations is that both *Q. faginea* populations share common pervasive alleles from another, non-targeted species of the same evolutionary radiation (Fraïsse *et al*., 2014). Such unattended species would be *Quercus pubescens*, which arriving to Iberia from eastern Europe colonized the river Ebro valley and could admix with both *Q. faginea* populations. Partial support to this hypothesis is found on the existence of *Quercus cerrioides* (= *Q. faginea* x *Q. pubescens)* hyrbrids in the flora catalog of the Font Roja Park (and own observations), and in the proximity of the Izki forest to the westernmost *Q.-pubescens* populations (Blanco-Castro *et al.*, 1997)..

## Supporting information

ABCToolbox files and scripts

All Supplemental materials

## Supplemental Files

**1- SuppFilesGoikoEtal2019.pdf** – All cited supplemental mateial, with exception of

**2- ABCTboxGoikoEtal2019** - Compressed folder with all necessary files to fully parameterizate the coalescent models (simulation), example files to perform model-choice and model-fit validation tests (estimation), and rudimentary R scripts to draw the graphs.

## Acknowledgments

This work was partly supported by grant CGL2009-07670 from the Spanish Ministry of Science and Innovation (MICINN). PG Goicoechea is indebted to Parks managers and crews, specially to Angel Gómez Manzaneque, who enormously facilitated sampling oak forests within the protected areas. Thanks are also due to Drs. de Villemereuil, François and Wegmann who kindly provided the ultimate software updates and offered very useful comments/recommendations to implement the analyses and to interpret the results.

