## Supplementary material for "Adaptive Introgression Promotes Fast Adaptation In Oaks Marginal Populations": ABCToolbox files and scripts: paramValidation.R

```
random <- read.delim("ABC_smodelFit_model0_RandomValidation.txt")
retained <- read.delim("ABC_modelFit_model0_RetainedValidation_Obs0.txt")
labels <- names(random)


##remember to change the main title for different models

## random validation
pdf("randomValidation_mode.pdf",width=9,height=11)
par(mfrow=c(3,3), oma=c(0,0,2,0))

for (n in c(3,8,13,18,23,28,33,38,43)) {
  xrange <- as.vector(range(random[,n]))
  plot(random[,n],random[,(n+1)], xlim=xrange, ylim=xrange, xlab=labels[n], ylab=labels[n+1])
  title("CAB", outer=T)
}
dev.off()

pdf("randomValidation_mean.pdf",width=9,height=12)
par(mfrow=c(3,3), oma=c(0,0,2,0))

for (n in c(3,8,13,18,23,28,33,38,43)) {
  xrange <- as.vector(range(random[,n]))
  plot(random[,n],random[,(n+2)], xlim=xrange, ylim=xrange, xlab=labels[n], ylab=labels[n+2])
  title("CAB", outer=T)
}
dev.off()


## retained validation
pdf("retainedValidation_mode.pdf",width=9,height=10)
par(mfrow=c(3,3), oma=c(0,0,2,0))

for (n in c(3,8,13,18,23,28,33,38,43)) {
  xrange <- as.vector(range(retained[,n]))
  yrange <- as.vector(range(retained[,(n+2)]))
  plot(retained[,n],retained[,(n+1)], xlim=xrange, ylim=yrange, xlab=labels[n], ylab=labels[n+1])
  title("CAB Retained Validation (mode)", outer=T)
}
dev.off()

pdf("retainedValidation_mean.pdf",width=9,height=10)
par(mfrow=c(3,3), oma=c(0,0,2,0))

for (n in c(3,8,13,18,23,28,33,38,43)) {
  xrange <- as.vector(range(retained[,n]))
  yrange <- as.vector(range(retained[,(n+2)]))
  plot(retained[,n],retained[,(n+2)], xlim=xrange, ylim=yrange, xlab=labels[n], ylab=labels[n+2])
  title("CAB Retained Validation (mean)", outer=T)
}
dev.off()
```
