## Supplementary material for "Adaptive Introgression Promotes Fast Adaptation In Oaks Marginal Populations": ABCToolbox files and scripts: plotPosteriors.R

```
ABC_GLM<-read.delim("ABC_modelFit_model0_MarginalPosteriorDensities_Obs0.txt")
ABC_rej<-read.delim("ABC_modelFit_model0_BestSimsParamStats_Obs0.txt")

params <- names(ABC_rej)

pdf("posteriorEstimation.pdf",width=9, height=11)
par(mfrow=c(3,3), oma=c(1,1,2,1))


for (i in 3:11) {
  j=2*(i-2)
  k=(j+1)
  dr <- density(ABC_rej[,i])
  xrange<-as.vector(range(c(dr$x, ABC_GLM[,j])))
  yrange<-as.vector(range(c(dr$y,ABC_GLM[,k])))
  plot(dr, xlab=params[i], xlim=xrange, ylim=yrange, main="", type="l", col="blue")
  lines(ABC_GLM[,j],ABC_GLM[,k], type="l", col="red")
  mtext("Posteriors Estimation (random blue vs. retained red)", font=1, side=3, line=-2, outer=T)
}
dev.off()
```
