## Supplementary material for "Adaptive Introgression Promotes Fast Adaptation In Oaks Marginal Populations": ABCToolbox files and scripts: posteriorBias.R

```
random.sim <- read.delim("ABC_modelFit_model0_RandomValidation.txt")
retained.sim <- read.delim("ABC_modelFit_model0_RetainedValidation_Obs0.txt")
labels <- names(random.sim)

# Little function to paste the K-S test label
#' @param xfrac The fraction over from the left side.
#' @param yfrac The fraction down from the top.
#' @param label The text to label with.
#' @param pos Position to pass to text()
#' @param ... Anything extra to pass to text(), e.g. cex, col.
add_label <- function(xfrac, yfrac, label, pos = 4, ...) {
  u <- par("usr")
  x <- u[1] + xfrac * (u[2] - u[1])
  y <- u[4] - yfrac * (u[4] - u[3])
  text(x, y, label, pos = pos, ...)
}

## Remember to change the main titles for the diferent models

##Random Posteriors bias
pdf("posterior_bias_random.pdf",width=9,height=13)
par(mfrow=c(3,3), oma=c(0.5, 0.75, 2, 0.25), mar=c(4, 4, 4, 4))
for (n in c(6,11,16,21,26,31,36,41,46)) {
  ks<-ks.test(random.sim[,n], "qunif")
  hist(random.sim[,n], main="", xlab=labels[n], ylab="Frequency")
  add_label(0.3, 0.07, paste("K-S test = ", round(ks$statistic, digits=3), sep=""))
  add_label(0.3, 0.15, paste("p-value = ", round(ks$p.value, digits=3), sep=""))
}
  title("IZK posterior bias", outer=T)
dev.off()


##Retained Posteriors bias
pdf("posterior_bias_retained.pdf",width=9,height=10)
par(mfrow=c(3,3), oma=c(0.5, 0.75, 2, 0.25), mar=c(4, 4, 4, 4))

for (n in c(6,11,16,21,26,31,36,41,46)) {
  ks<-ks.test(retained.sim[,n], "qunif")
  hist(retained.sim[,n], main="", xlab=labels[n], ylab="Frequency")
  add_label(0.3, 0.07, paste("K-S test = ", round(ks$statistic, digits=3), sep=""))
  add_label(0.3, 0.15, paste("p-value = ", round(ks$p.value, digits=3), sep=""))
}
  title("CAB Posterior Bias (retained simulations)", outer=T)
dev.off()
```
