## Supplementary material for "Adaptive Introgression Promotes Fast Adaptation In Oaks Marginal Populations": All Supplemental materials

[4]: BIOGECO, INRA- Univ. Bordeaux, 33610 Cestas, France

[5]: INIA Forest Res Ctr, Dept Forest Ecol & Genet, E-28040 Madrid, Spain

**Keywords:** marginal populations, range expansion/limits, adaptive introgression, local adaptation, landscape genetics, genome scans, oaks.

**Running Title:** Adaptive introgression in oaks marginal populations

**Supplemental FILE 1:** ENVIRONMENTAL VARIABLES

**Table S1-1:** Mean monthly values of the Accumulated Precipitation (Table S1-1a), the daily Minimum Temperatures (Table S1-1b), the daily Maximum Temperatures (Table S1-1c) and the daily Solar Radiation (Table S1-1d), for the seven populations from this study.

**Table S1-2:** Centered-standardized values of the 27 BIOCLIM environmental variables used in this study for each oak population. The descriptions in the second column indicate how to compute each variable. Variables in dark background were highly correlated with at least one of the 10 'uncorrelated' variables.

**Table S1-1a**

| Populations | January | February | March | April | May | June |
|---|---|---|---|---|---|---|
| CAB-fg | 75.3 | 75.5 | 47.9 | 69.4 | 58.5 | 37.1 |
| CJA-fg | 65.6 | 62.2 | 48.9 | 57.6 | 50.5 | 31.0 |
| FRO-fg | 40.6 | 33.6 | 58.9 | 63.0 | 91.7 | 57.5 |
| IZK-fg | 78.0 | 62.2 | 59.5 | 88.3 | 67.4 | 52.9 |
| CAB-py | 75.3 | 75.5 | 47.9 | 69.4 | 58.5 | 37.1 |
| IZK-py | 80.9 | 67.9 | 71.9 | 92.2 | 76.6 | 59.9 |
| SNE-py | 76.4 | 73.9 | 72.6 | 71.7 | 65.6 | 36.8 |

| Populations | July | August | September | October | November | December |
|---|---|---|---|---|---|---|
| CAB-fg | 14.3 | 11.8 | 32.3 | 60.5 | 79.3 | 95.7 |
| CJA-fg | 9.1 | 8.2 | 25.7 | 52.8 | 70.6 | 90.6 |
| FRO-fg | 30.2 | 28.1 | 41.9 | 38.5 | 54.5 | 39.8 |
| IZK-fg | 35.8 | 37.5 | 43.7 | 71.3 | 89.0 | 91.4 |
| CAB-py | 14.3 | 11.8 | 32.3 | 60.5 | 79.3 | 95.7 |
| IZK-py | 36.5 | 32.6 | 55.4 | 79.9 | 100.8 | 101.5 |
| SNE-py | 12.2 | 11.3 | 22.7 | 49.1 | 83.9 | 93.0 |

**Table S1-1b**

| Populations | January | February | March | April | May | June |
|---|---|---|---|---|---|---|
| CAB-fg | 0.7 | 1.9 | 3.5 | 5.5 | 8.9 | 13.3 |
| CJA-fg | 0.0 | 0.9 | 2.6 | 4.5 | 8.0 | 12.2 |
| FRO-fg | -1.8 | -0.3 | 1.4 | 2.9 | 6.5 | 9.9 |
| IZK-fg | 0.7 | 0.9 | 2.6 | 3.8 | 7.2 | 10.2 |
| CAB-py | 0.7 | 1.9 | 3.5 | 5.5 | 8.9 | 13.3 |
| IZK-py | -0.1 | 0.5 | 2.1 | 3.5 | 6.9 | 9.8 |
| SNE-py | -3.7 | -2.1 | -0.3 | 0.9 | 4.2 | 7.9 |

| Populations | July | August | September | October | November | December |
|---|---|---|---|---|---|---|
| CAB-fg | 16.5 | 16.2 | 13.3 | 8.5 | 4.0 | 1.6 |
| CJA-fg | 15.6 | 15.3 | 12.3 | 7.7 | 3.6 | 1.0 |
| FRO-fg | 14.1 | 13.4 | 10.7 | 5.2 | 1.4 | -1.0 |
| IZK-fg | 12.6 | 12.7 | 10.4 | 7.4 | 3.7 | 1.9 |
| CAB-py | 16.5 | 16.2 | 13.3 | 8.5 | 4.0 | 1.6 |
| IZK-py | 12.3 | 12.3 | 10.1 | 6.7 | 3.0 | 1.0 |
| SNE-py | 11.6 | 11.1 | 8.4 | 3.0 | -0.2 | -2.7 |

**Table S1-1c**

| Populations | January | February | March | April | May | June |
|---|---|---|---|---|---|---|
| CAB-fg | 10.4 | 12.2 | 15.8 | 17.9 | 22.4 | 28.3 |
| CJA-fg | 9.9 | 11.4 | 14.9 | 16.9 | 21.6 | 27.7 |
| FRO-fg | 9.5 | 10.4 | 13.1 | 15.0 | 19.2 | 23.8 |
| IZK-fg | 8.0 | 9.6 | 12.6 | 13.9 | 18.5 | 22.7 |
| CAB-py | 10.4 | 12.2 | 15.8 | 17.9 | 22.4 | 28.3 |
| IZK-py | 7.6 | 9.2 | 12.3 | 13.7 | 18.2 | 22.6 |
| SNE-py | 7.4 | 8.2 | 11.2 | 12.6 | 17.6 | 23.5 |

| Populations | July | August | September | October | November | December |
|---|---|---|---|---|---|---|
| CAB-fg | 33.5 | 33.1 | 28.2 | 20.9 | 14.7 | 10.9 |
| CJA-fg | 33.1 | 32.6 | 22.6 | 20.1 | 14.1 | 10.4 |
| FRO-fg | 29.1 | 28.9 | 24.6 | 18.1 | 13.1 | 10.2 |
| IZK-fg | 26.7 | 26.7 | 23.3 | 17.5 | 11.6 | 8.8 |
| CAB-py | 33.5 | 33.1 | 28.2 | 20.9 | 14.7 | 10.9 |
| IZK-py | 26.5 | 26.4 | 23.0 | 17.2 | 11.4 | 8.3 |
| SNE-py | 28.9 | 28.6 | 23.8 | 15.9 | 11.1 | 7.5 |

**Table S1-1d:**

| Populations | January | February | March | April | May | June |
|---|---|---|---|---|---|---|
| CAB-fg | 696 | 1120 | 1780 | 2474 | 2989 | 3214 |
| CJA-fg | 869 | 1311 | 1965 | 2616 | 3080 | 3270 |
| FRO-fg | 276 | 615 | 1267 | 2021 | 2647 | 2938 |
| IZK-fg | 968 | 1423 | 2092 | 2712 | 3126 | 3288 |
| CAB-py | 696 | 1120 | 1780 | 2474 | 2989 | 3214 |
| IZK-py | 819 | 1254 | 1909 | 2583 | 3061 | 3256 |
| SNE-py | 482 | 869 | 1533 | 2269 | 2839 | 3103 |

| Populations | July | August | September | October | November | December |
|---|---|---|---|---|---|---|
| CAB-fg | 3084 | 2676 | 2043 | 1353 | 801 | 553 |
| CJA-fg | 3159 | 2803 | 2216 | 1545 | 977 | 715 |
| FRO-fg | 2777 | 2271 | 1558 | 847 | 358 | 182 |
| IZK-fg | 3187 | 2864 | 2329 | 1679 | 1111 | 826 |
| CAB-py | 3084 | 2676 | 2043 | 1353 | 801 | 553 |
| IZK-py | 3139 | 2766 | 2170 | 1486 | 930 | 668 |
| SNE-py | 2951 | 2495 | 1812 | 1111 | 570 | 344 |

**Table S1-2:**

| Variable | Meaning | CAB-fg | CJA-fg | FRO-fg | IZK-fg | CAB-py | IZK-py | SNE-py |
|---|---|---|---|---|---|---|---|---|
| BIO1 | Annual Mean Temperature | 1.174 | 0.616 | -0.385 | -0.466 | 1.174 | -0.695 | -1.419 |
| BIO2 | Mean Diurnal Range (Mean of monthly (max temp - min temp)) | 0.590 | 0.381 | 0.456 | -1.537 | 0.590 | -1.352 | 0.872 |
| BIO3 | Isothermality (BIO2/BIO7) (* 100) | -0.592 | -1.653 | 1.337 | 0.502 | -0.592 | 0.376 | 0.622 |
| BIO4 | Temperature Seasonality (standard deviation *100) | 0.954 | 0.641 | -0.178 | -1.427 | 0.954 | -1.264 | 0.320 |
| BIO5 | Max Temperature of Warmest Month | 1.057 | 0.929 | -0.346 | -1.111 | 1.057 | -1.175 | -0.410 |
| BIO6 | Min Temperature of Coldest Month | 0.720 | 0.300 | -0.780 | 0.720 | 0.720 | 0.240 | -1.920 |
| BIO7 | Temperature Annual Range (BIO5-BIO6) | 0.685 | 0.782 | 0.069 | -1.519 | 0.685 | -1.324 | 0.620 |
| BIO8 | Mean Temperature of Wettest Quarter | 0.350 | 0.072 | 1.682 | -0.290 | 0.350 | -0.585 | -1.579 |
| BIO9 | Mean Temperature of Driest Quarter | 1.239 | 0.545 | -0.252 | -0.879 | 1.239 | -1.013 | -0.879 |
| BIO10 | Mean Temperature of Warmest Quarter | 1.139 | 0.825 | -0.294 | -0.893 | 1.139 | -1.022 | -0.893 |
| BIO11 | Mean Temperature of Coldest Quarter | 1.010 | 0.501 | -0.319 | 0.041 | 1.010 | -0.382 | -1.860 |
| BIO12 | Annual Precipitation | -0.230 | -1.055 | -1.002 | 0.932 | -0.230 | 1.702 | -0.117 |
| BIO13 | Precipitation of Wettest Month | 0.388 | -0.957 | -0.667 | -0.746 | 0.388 | 1.918 | -0.324 |
| BIO14 | Precipitation of Driest Month | -0.695 | -1.002 | 0.696 | 1.354 | -0.695 | 1.080 | -0.738 |
| BIO15 | Precipitation Seasonality (standard deviation *100) | 0.601 | 0.997 | -0.591 | -1.303 | 0.601 | -1.198 | 0.893 |
| BIO16 | Precipitation of Wettest Quarter | 0.103 | -0.945 | -1.533 | 0.464 | 0.103 | 1.570 | 0.237 |
| BIO17 | Precipitation of Driest Quarter | -0.574 | -1.019 | 0.635 | 1.120 | -0.574 | 1.337 | -0.926 |
| BIO18 | Precipitation of Warmest Quarter | -0.523 | -0.977 | 0.604 | 1.116 | -0.523 | 1.344 | -1.041 |
| BIO19 | Precipitation of Coldest Quarter | 0.513 | -0.064 | -2.208 | 0.207 | 0.513 | 0.591 | 0.447 |
| BIO20 | Annual Mean Radiation (W m-2) | 0.087 | 0.735 | -1.786 | 1.137 | 0.087 | 0.552 | -0.811 |
| BIO21 | Highest monthly radiation (W m-2) | 0.248 | 0.699 | -1.978 | 0.844 | 0.248 | 0.586 | -0.647 |
| BIO22 | Lowest monthly radiation (W m-2) | 0.019 | 0.750 | -1.653 | 1.250 | 0.019 | 0.538 | -0.923 |
| BIO23 | Radiation seasonality (standard deviation * 100) | -0.114 | -0.723 | 1.821 | -1.102 | -0.114 | -0.552 | 0.785 |
| BIO24 | Radiation of wettest quarter (W m-2) | -0.392 | -0.009 | 2.174 | -0.071 | -0.392 | -0.427 | -0.883 |
| BIO25 | Radiation of driest quarter (W m-2) | 0.277 | 0.574 | -2.172 | 0.725 | 0.277 | 0.493 | -0.174 |
| BIO26 | Radiation of warmest quarter (W m-2) | 0.143 | 0.730 | -1.864 | 1.030 | 0.143 | 0.571 | -0.751 |
| BIO27 | Radiation of coldest quarter (W m-2) | 0.043 | 0.757 | -1.715 | 1.194 | 0.043 | 0.548 | -0.871 |

**Supplemental FILE 2:** DATA AND BASIC GENETIC PARAMETERS.

**File S2-1:** RESULTS section accounting for several descriptive statistcs: null alleles, diversity estimates, population differentiation, autocorrelation of genetic and geographic distances and linkage disequilibrium).

**Table S2-1:** List of markers used in this study, their linkage groups, repeat motifs, PCR primers and Annealing Temperatures. Basic population genetic parameters (Allelic Richness AR, Observed Heterozygosity Ho; Expected Heterozygosity He, and Chi-square Tests of Hardy-Weinberg Equilibrium) are shown for each marker in each population. Values in red show significant deviations from HWE, after Bonferroni corrections.

**Table S2-2:** Null-allele frequencies for each marker in each population. Grayed cells indicate frequencies non-significantly different from zero. The number of markers with significant null-alleles frequencies in each population is shown in the bottom lane.

**Table S2-3:** Pairwise differentiation, Jost's D and Nei's Gst. among all populations. Confidence intervals of the estimates are between brackets.

**Figure S2-1:** Observed heterozygosities (Ho) among populations within species, for *Q. faginea* (Fig. S2-1a) and *Q. pyrenaica* (S2-1b). The FROfg population has been dropped for simplicity.

**Figure S2-2:** Autocorrelation plots between geographic and genetic distances (IBD) for each population and for the whole dataset with and without markers with extreme $F_{ST}$ values.

**Figure S2-3:** Linkage disequilibrium estimates for consecutive markers from each population (a-g) and for the two species (h). Points above the horizontal dashed lines are highly significant. The scale of the y-axis is very different in (h), which leads to hidden significance limit dashed lines at the bottom of the graph.

**File S2-1:** DESCRIPTIVE STATISTICS

Null alleles tests indicated that populations are probably at H-W equilibrium, although they contain an ample number of loci showing evidences of null alleles (ranging from 28-29 loci in the two CAB populations to 42-45 in the IZKfg and the CJAfg populations, Table S2-2). Diversity estimates did not show large differences with respect to mean allelic richness or expected heterozygosities (Table S2-1), which can not be easily translated into population sizes as admixture proportions probably play a large role in the estimates (Fraïsse *et al.*, 2016). Indeed, 2-bp EST-SSRs were more diverse than 3/6-bp microsatellites in all populations, which can be attributed to different mutation rates for both types of markers. The comparison of observed heterozygosities among populations within-species showed a large variance among markers, both within *Q. faginea* and *Q. pyrenaica* (Figures S2-1a, S2-1b). A large number of markers showed very large differences among different populations that can not be explained solely on the basis of the null alleles estimates. Thus, CJAfg showed several consecutive markers from LGs#5 and 6 with very low heterozygosities, which could be a footprint of ancient admixture. Unfortunately, the lack of central populations that could help to isolate the effects of ancient admixture on different marginal populations, and the variable null alleles frequencies across markers and populations, prevented us to further pursue this issue.

Pairwise differentiation coefficients (Table S2-3) were coherent with trends anticipated by ancestry results. Thus, the two CAB, the two IZK and the pair formed by CJA and SNE populations were less differentiated between them than with any other population from the respective species. Two other intra-specific differentiation values contributed to the observed geographic pattern. The CABfg and FROfg populations were most closely related than any other *Q. faginea* pair, in spite of the shortest distance between CABfg and CJAfg. Furthermore, the IZKpy and SNEpy populations were less differentiated between them than with the geographically intermediate CABpy population.

Spatial autocorrelation plots between genetic and physical distances were obtained for the whole dataset and for each population (Figure S2-2). At the local within populations scale, IBD patterns resembled those usually found in natural forest-tree populations (Neale and Savolainen, 2004;

Savolainen *et al.*, 2007), slightly positive autocorrelation at the shortest distance classes and non-significant subsequently. The regional scale showed a double structure of isolation components. The two closest inter-specific population pairs (CAB and IZK, distance class 168 km) showed highly significant positive autocorrelation, which could be attributed to either isolation-by-distance or to isolation-by-adaptation (IBD, IBA). All other distance classes were negatively correlated, a pattern that probably requires complex interactions among landscape characteristics and maximum dispersal distances (van Strien *et al.*, 2015).

Gametic linkage disequilibrium among consecutive pairs of loci was scarce in 5 out of the 7 studied populations (Figure S2-3). These populations, with 3 to 8 significant LD values, conform to the null hypothesis for tree populations under equilibrium, as large effective population sizes quickly erode LD (Neal and Savolainen, 2004; Savolainen *et al.*, 2007). Populations CABpy and FROfg contained a few more marker intervals at significant LD (17 and 16, respectively), but the number of highly significant intervals kept low. This increment can be explained by the spatial distribution of population FROfg, which it is spread into an altitudinal gradient of several small forests that increase the Wahlund effect. But other reasons should be sought for the CABpy population, which spreads continuously in both banks of a summer-dry creek. A very large number of marker intervals at highly significant LD were detected when populations within-species were merged, as expected due to isolate breaking.

**Table S2-1**

| Marker | LG | Repeat | Forward Primer | Reverse Primer | TA | Cabañeros *Q. faginea* | | | | Collado Jardines *Q. faginea* | | | | Font Roja *Q. faginea* | | | | Izki *Q. faginea* | | | | Cabañero *Q. pyrenaica* | | | | Izki *Q. pyrenaica* | | | | Sierra Nevada *Q. pyrenaica* | | | |
|---|---|---|---|---|---|---|---|---|---|---|---|---|---|---|---|---|---|---|---|---|---|---|---|---|---|---|---|---|---|---|---|---|---|
| | | | | | | AR | Ho | He | Chi-square | AR | Ho | He | Chi-square | AR | Ho | He | Chi-square | AR | Ho | He | Chi-square | AR | Ho | He | Chi-square | AR | Ho | He | Chi-square | AR | Ho | He | Chi-square |
| PIE076 | 1 | GAGGAA | CGATTTCCTCAGGATTGAGC | TTGCCGATCTCGATTCTCTT | 60.5 | 1.98 | 0.10 | 0.10 | 1.000 | 2.89 | 0.83 | 0.51 | 0.000 | 2.00 | 0.10 | 0.10 | 1.000 | 2.89 | 0.35 | 0.30 | 0.732 | 2.00 | 0.38 | 0.31 | 0.300 | 1.95 | 0.06 | 0.06 | 1.000 | 2.72 | 0.13 | 0.12 | 1.000 |
| FIR073 | 1 | TA | TCGCTGAGAAAAGAAACAAAAA | GAATTTCTGGGAAAGCGTGA | 59.5 | 21.70 | 0.75 | 0.95 | 0.000 | 22.02 | 0.70 | 0.94 | 0.000 | 20.36 | 0.77 | 0.92 | 0.004 | 22.67 | 0.65 | 0.94 | 0.000 | 12.76 | 0.44 | 0.79 | 0.000 | 17.09 | 0.64 | 0.91 | 0.000 | 17.09 | 0.65 | 0.92 | 0.000 |
| VIT026 | 1 | CCA | CCACCTAAACCAAACCCTCA | TGTTGCCAATACCGATTCCT | 60.5 | 7.40 | 0.35 | 0.71 | 0.000 | 8.06 | 0.46 | 0.76 | 0.000 | 9.32 | 0.46 | 0.81 | 0.000 | 9.26 | 0.38 | 0.84 | 0.000 | 6.54 | 0.48 | 0.67 | 0.002 | 7.48 | 0.40 | 0.74 | 0.000 | 6.94 | 0.47 | 0.75 | 0.001 |
| PIE238 | 1 | AT | CTTTGGTAGCATTCCTGTGC | TTCCCCCATTTGTGCTAAAC | 61.5 | 14.23 | 0.85 | 0.86 | 0.826 | 11.95 | 0.60 | 0.63 | 0.579 | 15.69 | 0.71 | 0.89 | 0.000 | 17.34 | 0.79 | 0.91 | 0.013 | 17.17 | 0.64 | 0.90 | 0.000 | 23.42 | 0.58 | 0.94 | 0.000 | 20.70 | 0.65 | 0.91 | 0.000 |
| POR040 | 1 | GA | CATCATACGACCCGAAAACC | TCTTCTGCTCCTCTCCCCTA | 59.5 | 16.31 | 0.83 | 0.86 | 0.568 | 16.04 | 0.75 | 0.90 | 0.000 | 15.67 | 0.52 | 0.82 | 0.000 | 13.48 | 0.81 | 0.86 | 0.282 | 12.41 | 0.69 | 0.74 | 0.044 | 16.04 | 0.88 | 0.89 | 0.004 | 14.73 | 0.81 | 0.89 | 0.149 |
| VIT007 | 1 | TC | ACCCTTTCCTCTTCCCAAAG | GGAAATCGGAAATCGGAAAT | 60.5 | 12.00 | 0.85 | 0.87 | 0.080 | 12.54 | 0.35 | 0.85 | 0.000 | 12.36 | 0.53 | 0.86 | 0.000 | 13.08 | 0.88 | 0.87 | 0.178 | 11.50 | 0.58 | 0.83 | 0.000 | 16.74 | 0.69 | 0.85 | 0.003 | 10.51 | 0.49 | 0.68 | 0.002 |
| WAG011 | 1 | CT | AAAAACCCCATTTCACCCTC | TTTGAGAAACTTTGGCCACC | 59.5 | 5.11 | 0.48 | 0.50 | 0.101 | 6.18 | 0.48 | 0.46 | 0.338 | 4.94 | 0.35 | 0.31 | 1.000 | 8.14 | 0.60 | 0.70 | 0.011 | 5.75 | 0.40 | 0.49 | 0.011 | 5.27 | 0.77 | 0.72 | 0.569 | 7.41 | 0.56 | 0.61 | 0.166 |
| FIR040 | 1 | GGA | GAAACGGAAACGCAAACG | GAGCAGAGGGCTAAACTCCA | 59.5 | 4.92 | 0.69 | 0.73 | 0.137 | 3.94 | 0.65 | 0.65 | 0.649 | 4.00 | 0.71 | 0.69 | 0.964 | 3.67 | 0.50 | 0.57 | 0.726 | 6.49 | 0.72 | 0.75 | 0.508 | 3.95 | 0.58 | 0.64 | 0.156 | 3.98 | 0.65 | 0.61 | 0.682 |
| PIE223 | 2 | GGT | AGAAGCCCAACACGGCTAC | AGCAAAACACAAACGCACAA | 63.5 | 7.90 | 0.74 | 0.77 | 0.617 | 9.48 | 0.71 | 0.81 | 0.088 | 9.60 | 0.60 | 0.84 | 0.000 | 13.24 | 0.78 | 0.89 | 0.000 | 7.07 | 0.51 | 0.70 | 0.003 | 9.52 | 0.56 | 0.67 | 0.003 | 14.26 | 0.67 | 0.82 | 0.073 |
| FIR065 | 2 | CTT | ATTCCCATGCATCAAAATCC | TCCTTCAGTTTGAGAGCTCCTT | 59.5 | 4.03 | 0.44 | 0.51 | 0.105 | 3.49 | 0.23 | 0.44 | 0.000 | 2.00 | 0.29 | 0.33 | 0.389 | 4.35 | 0.63 | 0.68 | 0.022 | 3.56 | 0.60 | 0.63 | 0.314 | 4.25 | 0.50 | 0.58 | 0.499 | 5.64 | 0.63 | 0.65 | 0.780 |
| ZQR87 | 2 | AG | TCCCACCACTTTGGTCTCTCA | GTTGTCAGCAGTGGGATGGGTA | 56.0 | 6.17 | 0.56 | 0.55 | 0.482 | 6.36 | 0.54 | 0.60 | 0.007 | 9.17 | 0.68 | 0.64 | 0.892 | 9.49 | 0.64 | 0.67 | 0.368 | 12.06 | 0.88 | 0.83 | 0.331 | 9.09 | 0.81 | 0.75 | 0.000 | 13.75 | 0.85 | 0.87 | 0.152 |
| FIR048 | 2 | CT | TGCACCAAAATTGGAGGATG | TTGATGCAAGGTGCAGTTTC | 59.5 | 4.13 | 0.68 | 0.52 | 0.237 | 3.49 | 0.17 | 0.19 | 0.410 | 4.26 | 0.33 | 0.31 | 0.160 | 9.39 | 0.33 | 0.32 | 0.626 | 7.97 | 0.72 | 0.78 | 0.660 | 13.62 | 0.70 | 0.81 | 0.001 | 11.36 | 0.81 | 0.73 | 0.566 |
| FIR013 | 2 | CAG | CGGGGAGGTTGATGAGTATT | AACACTGTCACCCCCATAGC | 58.5 | 2.58 | 0.30 | 0.31 | 0.182 | 4.25 | 0.31 | 0.29 | 0.659 | 4.28 | 0.38 | 0.34 | 0.398 | 2.67 | 0.50 | 0.51 | 1.000 | 3.17 | 0.42 | 0.44 | 0.471 | 4.00 | 0.63 | 0.58 | 0.128 | 3.98 | 0.60 | 0.56 | 0.870 |
| ZQP119 | 2 | AG | GATCAGTGATAGTGCCTCTC | GATCAACAAGCCCAAGGCAC | 56.0 | 18.06 | 0.78 | 0.90 | 0.037 | 15.15 | 0.54 | 0.83 | 0.000 | 18.80 | 0.49 | 0.90 | 0.000 | 20.52 | 0.60 | 0.92 | 0.000 | 17.35 | 0.60 | 0.91 | 0.000 | 18.05 | 0.98 | 0.89 | 0.092 | 17.04 | 0.63 | 0.89 | 0.000 |
| POR038 | 2 | CT | GCCTGCTTTTTGTTCCAGAT | CGAGCTGTTCTTGAAGGTTG | 59.5 | 14.24 | 0.94 | 0.89 | 0.078 | 16.63 | 0.90 | 0.92 | 0.385 | 12.53 | 0.63 | 0.86 | 0.000 | 12.99 | 0.75 | 0.86 | 0.025 | 9.57 | 0.85 | 0.83 | 0.234 | 15.13 | 0.79 | 0.79 | 0.784 | 7.56 | 0.38 | 0.72 | 0.000 |
| FIR032 | 2 | TA | TGAAACTGGTCTCAGATTTTGTG | ACACTACACACACTTGCTCTTTTT | 58.5 | 9.38 | 0.60 | 0.73 | 0.322 | 13.19 | 0.77 | 0.87 | 0.000 | 12.97 | 0.53 | 0.79 | 0.000 | 14.48 | 0.71 | 0.83 | 0.000 | 14.83 | 0.90 | 0.90 | 0.075 | 15.64 | 0.81 | 0.90 | 0.000 | 14.04 | 0.59 | 0.87 | 0.000 |
| PIE197 | 2 | CTA | CAAGTACCACACCCACAACC | GAAAGAGCACTGCCTCCAAC | 58.0 | 2.92 | 0.50 | 0.48 | 1.000 | 2.50 | 0.44 | 0.45 | 0.555 | 5.64 | 0.58 | 0.62 | 0.000 | 3.35 | 0.48 | 0.46 | 1.000 | 3.62 | 0.43 | 0.35 | 0.768 | 3.95 | 0.50 | 0.46 | 0.423 | 3.71 | 0.46 | 0.44 | 0.839 |
| PIE075 | 3 | TATTTT | GTGCGCTTCAGAGGAACACT | CCAATCCCAATCTCTTTCGT | 59.5 | 2.97 | 0.27 | 0.35 | 0.005 | 3.00 | 0.22 | 0.58 | 0.000 | 3.86 | 0.13 | 0.28 | 0.000 | 4.93 | 0.04 | 0.42 | 0.000 | 2.56 | 0.40 | 0.42 | 0.321 | 5.25 | 0.15 | 0.66 | 0.000 | 4.90 | 0.15 | 0.40 | 0.000 |
| PIE135 | 3 | CT | TTGTCTACCTTTATCAAACATCATT | CAAATACGAGAATACCACAGCAAC | 59.5 | 13.04 | 0.50 | 0.88 | 0.000 | 14.45 | 0.67 | 0.90 | 0.000 | 16.38 | 0.60 | 0.89 | 0.000 | 14.05 | 0.60 | 0.89 | 0.000 | 8.44 | 0.85 | 0.82 | 0.219 | 12.67 | 0.80 | 0.86 | 0.152 | 12.98 | 0.71 | 0.89 | 0.014 |
| FIR047 | 3 | TCG | AACCCATTTGAGGTTTTTGC | CCGAAAAAGAAAACCCACCT | 58.0 | 1.00 | 0.00 | 0.00 | 1.000 | 1.54 | 0.02 | 0.02 | 1.000 | 2.86 | 0.08 | 0.12 | 0.014 | 1.90 | 0.04 | 0.04 | 1.000 | 2.49 | 0.07 | 0.07 | 1.000 | 1.98 | 0.08 | 0.08 | 1.000 | 2.00 | 0.09 | 0.08 | 1.000 |
| GOT021 | 3 | AT | AGAAAGTTCCAGGGAAAGCA | CTTCGTCCCCAGTTGAATGT | 59.5 | 2.36 | 0.06 | 0.06 | 1.000 | 1.97 | 0.10 | 0.10 | 1.000 | 1.00 | 0.00 | 0.00 | 1.000 | 3.58 | 0.34 | 0.29 | 0.761 | 3.36 | 0.52 | 0.53 | 0.583 | 3.62 | 0.60 | 0.56 | 0.444 | 3.64 | 0.50 | 0.44 | 0.338 |
| PIE021 | 3 | TC | CACCTCTCTTTCCTTCTTTAGCA | CCCAGATAAGAGAACCCCATA | 59.5 | 19.49 | 0.83 | 0.90 | 0.009 | 21.07 | 0.72 | 0.92 | 0.000 | 15.42 | 0.72 | 0.84 | 0.001 | 23.16 | 0.54 | 0.94 | 0.000 | 15.13 | 0.56 | 0.89 | 0.000 | 24.27 | 0.63 | 0.94 | 0.000 | 20.46 | 0.69 | 0.88 | 0.000 |
| VIT107 | 3 | TA | TGATCACAGATTGGAGCTTAACA | CCCCCACTTAGGAAAGAAGC | 60.5 | 12.69 | 0.84 | 0.87 | 0.816 | 9.24 | 0.83 | 0.86 | 0.083 | 7.18 | 0.79 | 0.79 | 0.281 | 15.14 | 0.88 | 0.88 | 0.409 | 15.45 | 0.92 | 0.92 | 0.209 | 16.41 | 0.85 | 0.86 | 0.353 | 12.88 | 0.92 | 0.73 | 0.767 |
| POR011 | 3 | CTT | ATCCTTCAACGCCCTTTAGC | GGCAAGCTTAGAAACGGAGA | 59.5 | 8.61 | 0.81 | 0.80 | 0.705 | 8.94 | 0.78 | 0.78 | 0.788 | 7.51 | 0.79 | 0.78 | 0.412 | 8.79 | 0.83 | 0.83 | 0.523 | 7.92 | 0.86 | 0.83 | 0.205 | 7.62 | 0.90 | 0.83 | 0.034 | 9.31 | 0.87 | 0.81 | 0.373 |
| VIT022 | 3 | AG | TTCTGGGAAAGTAGCTTTGGA | TTCAACGCAGACTTGGTTTG | 60.5 | 9.62 | 0.57 | 0.75 | 0.000 | 9.22 | 0.69 | 0.65 | 0.705 | 9.71 | 0.87 | 0.72 | 0.224 | 11.56 | 0.60 | 0.73 | 0.000 | 7.35 | 0.48 | 0.52 | 0.170 | 6.37 | 0.40 | 0.45 | 0.351 | 13.75 | 0.77 | 0.85 | 0.433 |
| PIE242 | 3 | TA | GGAGGGAAAAGAACAATGC | TTGCAATCCTCCAAATTTAATG | 60.5 | 8.51 | 0.88 | 0.79 | 0.000 | 9.10 | 0.83 | 0.85 | 0.131 | 10.17 | 0.79 | 0.77 | 0.725 | 16.44 | 0.73 | 0.89 | 0.000 | 9.99 | 0.69 | 0.78 | 0.027 | 14.47 | 0.65 | 0.88 | 0.000 | 9.21 | 0.65 | 0.75 | 0.317 |
| PIE051 | 3 | ACCAAC | TCAAAACCAAAGCCAAAACC | GCAGCTTCCAAAAGATCACC | 59.5 | 5.86 | 0.54 | 0.52 | 0.895 | 5.15 | 0.36 | 0.46 | 0.137 | 5.29 | 0.50 | 0.43 | 0.548 | 6.33 | 0.75 | 0.61 | 0.562 | 6.36 | 0.58 | 0.67 | 0.002 | 5.13 | 0.51 | 0.45 | 0.410 | 4.70 | 0.54 | 0.58 | 0.047 |
| PIE088 | 4 | TCTTTC | AGAATGGCCTGTACTTTCCTG | CATACTGGAGTCCTGGCAGAG | 58.0 | 8.07 | 0.17 | 0.73 | 0.000 | 6.32 | 0.33 | 0.39 | 0.016 | 5.52 | 0.33 | 0.52 | 0.006 | 8.26 | 0.20 | 0.71 | 0.000 | 5.80 | 0.19 | 0.72 | 0.000 | 6.23 | 0.13 | 0.65 | 0.000 | 8.69 | 0.23 | 0.80 | 0.000 |
| VIT043 | 4 | CA | GATGGGTTGAGCGGACTATC | CAACCAAAATGGAAACATCG | 59.5 | 5.80 | 0.58 | 0.55 | 0.506 | 4.25 | 0.33 | 0.45 | 0.014 | 3.00 | 0.54 | 0.54 | 0.357 | 5.66 | 0.29 | 0.51 | 0.000 | 4.61 | 0.33 | 0.39 | 0.172 | 3.62 | 0.44 | 0.43 | 0.792 | 4.98 | 0.48 | 0.52 | 0.157 |
| PIE082 | 4 | CAGCAA | GAATTCGATTTTTGATCGAAGC | GAGCAGAGTCCGGTTGTGAC | 59.5 | 4.47 | 0.42 | 0.42 | 0.482 | 2.50 | 0.48 | 0.42 | 0.172 | 2.88 | 0.23 | 0.43 | 0.000 | 2.97 | 0.31 | 0.35 | 0.426 | 5.08 | 0.48 | 0.48 | 0.027 | 3.95 | 0.67 | 0.57 | 0.024 | 3.73 | 0.70 | 0.57 | 0.041 |
| PIE216 | 4 | GAT | ACCATTACCAAAACCCTTCAAC | TCTTTTTGGAAGGATTGATGAGA | 59.5 | 6.03 | 0.81 | 0.64 | 0.150 | 6.23 | 0.58 | 0.71 | 0.168 | 5.53 | 0.63 | 0.66 | 0.229 | 8.01 | 0.94 | 0.78 | 0.016 | 7.35 | 0.74 | 0.68 | 0.441 | 6.62 | 0.92 | 0.77 | 0.108 | 8.44 | 0.71 | 0.82 | 0.005 |
| PIE228 | 4 | AGA | TGGAGGAGCACTGCATATTG | CACTGTGGCTGGAGCATCTA | 60.5 | 8.93 | 0.67 | 0.74 | 0.067 | 7.35 | 0.67 | 0.69 | 0.302 | 11.10 | 0.79 | 0.84 | 0.059 | 9.24 | 0.77 | 0.82 | 0.015 | 12.08 | 0.84 | 0.85 | 0.048 | 10.36 | 0.83 | 0.84 | 0.252 | 11.41 | 0.73 | 0.75 | 0.037 |
| POR029 | 4 | ACC | TGGGTTGTGTCTCTCAATGG | AAGCCTGCAATGAAGCTGAT | 63.0 | 9.68 | 0.92 | 0.82 | 0.397 | 7.43 | 0.83 | 0.80 | 0.276 | 6.60 | 0.73 | 0.77 | 0.070 | 10.57 | 0.88 | 0.81 | 0.000 | 6.97 | 0.98 | 0.82 | 0.000 | 9.56 | 0.81 | 0.85 | 0.001 | 5.96 | 0.74 | 0.76 | 0.333 |
| VIT003 | 4 | TA | GCAGTATCCAAACAGACCCTAA | GCTTTGCTCTAAACAGCAACTT | 59.5 | 11.08 | 0.66 | 0.83 | 0.002 | 11.99 | 0.42 | 0.88 | 0.000 | 11.93 | 0.72 | 0.87 | 0.015 | 12.26 | 0.52 | 0.88 | 0.000 | 12.43 | 0.63 | 0.89 | 0.000 | 12.71 | 0.63 | 0.86 | 0.000 | 12.59 | 0.49 | 0.86 | 0.000 |
| VIT018 | 4 | (CCA)(AAC) | TGATGCGTTTGTGTTTGTTG | CCAACAACATGTGCCAAAAC | 59.5 | 1.56 | 0.02 | 0.02 | 1.000 | 2.38 | 0.08 | 0.08 | 1.000 | 4.20 | 0.09 | 0.32 | 0.000 | 2.79 | 0.04 | 0.08 | 0.024 | 1.81 | 0.04 | 0.04 | 1.000 | 2.59 | 0.09 | 0.08 | 1.000 | 1.73 | 0.02 | 0.02 | 1.000 |
| VIT017 | 4 | CGG | GATCCGAATCCGAAACTGAA | GTTACGAGCGAGGACGTTGT | 59.5 | 4.89 | 0.44 | 0.39 | 0.824 | 4.07 | 0.19 | 0.19 | 0.170 | 3.54 | 0.17 | 0.16 | 1.000 | 5.21 | 0.31 | 0.33 | 0.054 | 2.95 | 0.19 | 0.17 | 1.000 | 4.43 | 0.27 | 0.26 | 0.224 | 5.69 | 0.46 | 0.42 | 0.215 |
| PIE002 | 4 | AG | CTCCTCCATTTCCCAATTCA | TCGCTTGTGGTACATCTTGG | 60.5 | 11.47 | 0.52 | 0.81 | 0.000 | 8.97 | 0.60 | 0.71 | 0.139 | 10.13 | 0.60 | 0.75 | 0.004 | 7.90 | 0.71 | 0.71 | 0.387 | 5.11 | 0.69 | 0.76 | 0.347 | 9.06 | 0.56 | 0.74 | 0.000 | 9.12 | 0.77 | 0.81 | 0.014 |
| PIE029 | 5 | TTC | TCACGTGGTTTGTCGTCTTT | CAATACTGTTTTTCGGCAATGA | 60.5 | 9.66 | 0.46 | 0.74 | 0.000 | 6.51 | 0.29 | 0.55 | 0.000 | 5.13 | 0.31 | 0.44 | 0.005 | 7.75 | 0.54 | 0.72 | 0.003 | 5.00 | 0.22 | 0.29 | 0.001 | 7.65 | 0.52 | 0.62 | 0.001 | 7.36 | 0.48 | 0.61 | 0.130 |
| WAG043 | 5 | AG | GCCCACAGGCAAATAACAAC | AACTCAGGCATGGGCTACAG | 59.5 | 16.57 | 0.81 | 0.90 | 0.004 | 18.92 | 0.80 | 0.92 | 0.008 | 14.68 | 0.77 | 0.88 | 0.019 | 22.11 | 0.75 | 0.92 | 0.000 | 15.16 | 0.85 | 0.89 | 0.004 | 20.26 | 0.85 | 0.93 | 0.007 | 18.56 | 0.66 | 0.85 | 0.000 |
| POR014 | 5 | CT | TTCCCACACATCAATTTTTCC | TGAGACGTAAGAGCCAATCCT | 59.5 | 11.03 | 0.81 | 0.85 | 0.063 | 12.19 | 0.75 | 0.88 | 0.075 | 11.02 | 0.69 | 0.80 | 0.041 | 9.57 | 0.48 | 0.80 | 0.000 | 10.73 | 0.39 | 0.78 | 0.000 | 15.14 | 0.63 | 0.87 | 0.000 | 12.47 | 0.70 | 0.86 | 0.042 |
| GOT065 | 5 | CTT | AATGCAGGCATGACCTCTCT | TCAACCCGACTACGGAGAAC | 59.5 | 3.98 | 0.67 | 0.60 | 0.035 | 4.47 | 0.56 | 0.57 | 0.112 | 4.53 | 0.40 | 0.48 | 0.149 | 7.01 | 0.65 | 0.71 | 0.108 | 4.79 | 0.71 | 0.69 | 0.009 | 5.25 | 0.77 | 0.71 | 0.558 | 5.44 | 0.77 | 0.70 | 0.001 |
| VIT023 | 5 | ATA | AATGCGAACGACATGAACAA | CTCTCGTCGGAGACTCAACC | 58.0 | 1.99 | 0.11 | 0.10 | 1.000 | 4.75 | 0.35 | 0.37 | 0.329 | 3.30 | 0.15 | 0.17 | 0.328 | 4.67 | 0.50 | 0.52 | 0.588 | 4.00 | 0.60 | 0.56 | 0.030 | 3.63 | 0.60 | 0.51 | 0.744 | 3.00 | 0.26 | 0.32 | 0.068 |
| FIR106 | 5 | TAT | GGGTGAACATGGTGGAGATT | GAGAAAACCACACCTCACAAGA | 60.5 | 4.13 | 0.89 | 0.67 | 0.000 | 3.75 | 0.08 | 0.46 | 0.000 | 1.00 | 0.00 | 0.00 | 1.000 | 4.24 | 0.08 | 0.18 | 0.000 | 2.56 | 0.96 | 0.51 | 0.000 | 1.98 | 0.08 | 0.08 | 1.000 | 3.84 | 0.17 | 0.16 | 1.000 |
| POR030 | 5 | CCG | CTCACTCCAAACTCCGAAGC | CTAACCCTCCCACCTCCATT | 60.5 | 5.36 | 0.71 | 0.70 | 0.682 | 4.00 | 0.50 | 0.71 | 0.011 | 4.65 | 0.88 | 0.70 | 0.000 | 5.56 | 0.83 | 0.68 | 0.185 | 4.54 | 0.65 | 0.63 | 0.999 | 4.79 | 0.54 | 0.46 | 0.444 | 4.00 | 0.73 | 0.70 | 0.003 |
| POR039 | 5 | CCT | CGAAGGAAAAGCCAAAGAA | GCATTCAGTGCAACAAGTCC | 62.5 | 7.52 | 0.63 | 0.63 | 0.147 | 6.16 | 0.65 | 0.67 | 0.870 | 7.05 | 0.66 | 0.59 | 0.035 | 4.97 | 0.92 | 0.67 | 0.000 | 4.92 | 0.73 | 0.70 | 0.152 | 6.62 | 0.96 | 0.71 | 0.000 | 8.36 | 0.79 | 0.70 | 0.237 |
| VIT021 | 5 | GA | GCCGATCCTCGATCTGATAC | CGCTGAGACCTTCTTCAACC | 59.5 | 13.68 | 0.91 | 0.89 | 0.826 | 15.41 | 0.88 | 0.91 | 0.322 | 12.53 | 0.96 | 0.87 | 0.021 | 14.78 | 0.94 | 0.91 | 0.190 | 11.30 | 0.83 | 0.84 | 0.305 | 13.82 | 0.94 | 0.86 | 0.618 | 13.11 | 0.88 | 0.83 | 0.770 |
| POR016 | 6 | GGT | AGCAACAGCAGAGCCAAAAT | CAGCGGCTTTGAGGTAATTC | 60.5 | 4.65 | 0.25 | 0.26 | 0.447 | 4.00 | 0.23 | 0.31 | 0.162 | 3.53 | 0.02 | 0.17 | 0.000 | 5.43 | 0.17 | 0.28 | 0.002 | 2.00 | 0.17 | 0.15 | 1.000 | 3.48 | 0.17 | 0.29 | 0.002 | 7.84 | 0.29 | 0.79 | 0.000 |
| FIR021 | 6 | CAC | CTCATCATCGGAGGTGGAGT | TTTGAGAAGAGGCGAGTCGT | 60.5 | 7.33 | 0.40 | 0.73 | 0.000 | 5.91 | 0.29 | 0.74 | 0.000 | 4.95 | 0.31 | 0.68 | 0.000 | 8.24 | 0.46 | 0.79 | 0.000 | 4.72 | 0.54 | 0.68 | 0.034 | 6.23 | 0.69 | 0.71 | 0.015 | 7.64 | 0.67 | 0.79 | 0.041 |
| POR024 | 6 | AG | GCCCCGGATTTATTTCTCTG | TGAGCCTCAACCAAAACAAA | 60.5 | 11.14 | 0.81 | 0.84 | 0.042 | 4.66 | 0.17 | 0.25 | 0.000 | 15.56 | 0.77 | 0.78 | 0.565 | 15.03 | 0.81 | 0.84 | 0.109 | 9.27 | 0.91 | 0.66 | 0.000 | 10.32 | 0.77 | 0.81 | 0.080 | 11.28 | 0.90 | 0.86 | 0.079 |
| PIE125 | 6 | GGAAGC | AATACAAATCGCAGGAGGTG | CTAACCCATCGTTCATGGAG | 59.5 | 3.56 | 0.63 | 0.54 | 0.393 | 2.99 | 0.29 | 0.43 | 0.003 | 3.61 | 0.27 | 0.33 | 0.294 | 4.35 | 0.54 | 0.63 | 0.257 | 6.73 | 0.81 | 0.71 | 0.015 | 3.62 | 0.52 | 0.63 | 0.133 | 4.00 | 0.74 | 0.66 | 0.562 |
| POR034 | 6 | TTC | TGCGTCTCAATCACTCTCTCTT | TCGATCCGTGAAAATCTGAA | 59.5 | 5.47 | 0.69 | 0.70 | 0.706 | 6.00 | 0.79 | 0.71 | 0.971 | 5.88 | 0.83 | 0.75 | 0.863 | 6.66 | 0.75 | 0.73 | 0.107 | 7.43 | 0.75 | 0.74 | 0.000 | 6.18 | 0.71 | 0.66 | 0.401 | 4.72 | 0.48 | 0.65 | 0.011 |
| VIT028 | 6 | AG | CGGAGTTGGTAGGCTCGAT | CTCTCCCTCACTCTCCCTCA | 59.5 | 12.45 | 0.48 | 0.84 | 0.000 | 6.37 | 0.23 | 0.51 | 0.000 | 9.10 | 0.29 | 0.75 | 0.000 | 12.47 | 0.49 | 0.88 | 0.000 | 6.03 | 0.60 | 0.77 | 0.005 | 8.85 | 0.62 | 0.79 | 0.006 | 5.72 | 0.49 | 0.67 | 0.022 |
| POR027 | 6 | GA | AGGCCCAGTTCATCCTTACA | TGGTAATCCAACTCCAAGCTG | 59.5 | 6.05 | 0.67 | 0.58 | 0.689 | 2.91 | 0.19 | 0.17 | 1.000 | 5.28 | 0.42 | 0.44 | 0.821 | 4.96 | 0.58 | 0.52 | 0.438 | 3.94 | 0.15 | 0.16 | 0.028 | 5.55 | 0.60 | 0.57 | 0.767 | 5.68 | 0.44 | 0.53 | 0.024 |
| PIE071 | 6 | TATTTT | GTCGGCGATGAGAGGAAAG | GGAGACATAGTTCCAACTCAAGC | 59.5 | 10.70 | 0.65 | 0.84 | 0.000 | 10.40 | 0.72 | 0.86 | 0.005 | 11.59 | 0.85 | 0.88 | 0.569 | 14.72 | 0.83 | 0.89 | 0.339 | 9.40 | 0.75 | 0.80 | 0.025 | 16.21 | 0.81 | 0.92 | 0.003 | 11.76 | 0.67 | 0.87 | 0.001 |
| PIE219 | 7 | ACC | CAACAGAGTCTGGTAGCGAAA | CCAGTCATGCATGTTCTTCG | 60.5 | 5.58 | 0.46 | 0.43 | 0.617 | 5.75 | 0.42 | 0.49 | 0.050 | 5.66 | 0.21 | 0.20 | 1.000 | 4.66 | 0.77 | 0.62 | 0.042 | 8.37 | 0.85 | 0.78 | 0.105 | 4.48 | 0.58 | 0.49 | 0.091 | 4.90 | 0.63 | 0.55 | 0.567 |
| PIE137 | 7 | TC | TCTCCTACCACCAAAACAAGG | CACTTCTCTAACCCCATCAGC | 59.5 | 4.94 | 0.40 | 0.36 | 1.000 | 4.93 | 0.47 | 0.49 | 0.300 | 4.54 | 0.36 | 0.35 | 0.098 | 9.41 | 0.67 | 0.70 | 0.198 | 4.97 | 0.62 | 0.68 | 0.050 | 4.62 | 0.48 | 0.48 | 0.324 | 3.98 | 0.31 | 0.33 | 0.236 |
| VIT046 | 7 | CT | ACATCACATGCTGGGGAAAT | GCCAAGTGCTATCAGTGCAA | 58.0 | 6.22 | 0.69 | 0.70 | 0.080 | 4.94 | 0.75 | 0.71 | 0.464 | 4.32 | 0.83 | 0.66 | 0.052 | 7.46 | 0.63 | 0.75 | 0.101 | 4.72 | 0.54 | 0.52 | 0.880 | 5.77 | 0.47 | 0.48 | 0.602 | 5.69 | 0.50 | 0.55 | 0.400 |
| FIR043 | 7 | TC | TTCTCCATTTCACACGCTTC | ACGACATCGTTTTGGAGCTT | 60.5 | 9.52 | 0.65 | 0.80 | 0.001 | 15.45 | 0.63 | 0.88 | 0.000 | 8.79 | 0.74 | 0.73 | 0.276 | 10.79 | 0.69 | 0.80 | 0.001 | 9.25 | 0.51 | 0.76 | 0.000 | 10.29 | 0.89 | 0.82 | 0.000 | 11.24 | 0.81 | 0.83 | 0.007 |
| FIR030 | 7 | AG | GGACATATTATCTAGGAGACGAG | ATGTCCCATAGCACAGAGCA | 59.5 | 10.81 | 0.81 | 0.83 | 0.120 | 10.07 | 0.42 | 0.78 | 0.000 | 9.21 | 0.46 | 0.80 | 0.000 | 13.24 | 0.55 | 0.69 | 0.024 | 7.00 | 0.25 | 0.48 | 0.000 | 9.71 | 0.38 | 0.43 | 0.037 | 10.18 | 0.38 | 0.56 | 0.000 |
| FIR095 | 7 | CT | TCCCACTTTCCCTCTCACAC | TGGAATATGGAAGTGGGTTCA | 59.5 | 10.13 | 0.83 | 0.86 | 0.260 | 14.43 | 0.77 | 0.90 | 0.000 | 6.65 | 0.83 | 0.81 | 0.931 | 13.31 | 0.81 | 0.87 | 0.181 | 10.06 | 0.90 | 0.87 | 0.777 | 10.99 | 0.83 | 0.86 | 0.959 | 10.61 | 0.88 | 0.83 | 0.209 |
| PIE127 | 7 | ACACCA | CTTTTGGGGATGATCTGTTG | AGGATCCTTTGAATGGGAAG | 60.5 | 2.00 | 0.28 | 0.27 | 1.000 | 3.50 | 0.48 | 0.43 | 1.000 | 2.54 | 0.06 | 0.06 | 1.000 | 3.67 | 0.46 | 0.53 | 0.401 | 3.00 | 0.56 | 0.51 | 0.666 | 2.00 | 0.23 | 0.26 | 0.306 | 3.00 | 0.35 | 0.41 | 0.206 |
| VIT009 | 7 | AG | CTGCTGCTGCTTCTTCCTCT | GCGATCCGTAGGGTTAGGTC | 59.5 | 12.15 | 0.64 | 0.86 | 0.000 | 8.34 | 0.88 | 0.84 | 0.325 | 10.47 | 0.58 | 0.86 | 0.000 | 8.63 | 0.73 | 0.81 | 0.197 | 12.51 | 0.71 | 0.85 | 0.001 | 13.10 | 0.74 | 0.89 | 0.000 | 11.59 | 0.88 | 0.87 | 0.049 |
| POR025 | 7 | TC | CACACAAACCCATATGATCTGAA | TCTCTTTCGATCCCTTCTGC | 59.5 | 13.43 | 0.77 | 0.86 | 0.048 | 6.60 | 0.94 | 0.64 | 0.000 | 15.08 | 0.94 | 0.90 | 0.022 | 14.55 | 0.77 | 0.89 | 0.006 | 13.51 | 0.98 | 0.81 | 0.000 | 16.83 | 0.79 | 0.89 | 0.010 | 13.10 | 0.77 | 0.84 | 0.017 |
| WAG021 | 8 | TTG | ACCATCCTCTGATGCTGCTC | GTAAGGAAGAGACCGTGGGG | 59.5 | 4.61 | 0.71 | 0.65 | 0.384 | 5.51 | 0.60 | 0.56 | 0.462 | 4.99 | 0.67 | 0.69 | 0.285 | 6.64 | 0.73 | 0.76 | 0.178 | 4.97 | 0.56 | 0.62 | 0.317 | 5.85 | 0.77 | 0.74 | 0.018 | 6.56 | 0.77 | 0.73 | 0.937 |
| POR008 | 8 | GTG | AGGGGTCATCAGAGTTGGTG | ATCCCTCAGTCTTCCCGAAC | 60.5 | 2.98 | 0.02 | 0.24 | 0.000 | 3.32 | 0.11 | 0.10 | 1.000 | 1.99 | 0.08 | 0.08 | 1.000 | 3.96 | 0.19 | 0.35 | 0.000 | 1.00 | 0.00 | 0.00 | 1.000 | 2.00 | 0.13 | 0.15 | 0.299 | 1.98 | 0.06 | 0.06 | 1.000 |
| PIE023 | 8 | AG | GGTCTCTCTTATTTTCTCTTTCTT | TGTGAAGAAGAACAAGACCCTTT | 59.5 | 2.36 | 0.06 | 0.06 | 1.000 | 2.47 | 0.04 | 0.12 | 0.005 | 3.95 | 0.27 | 0.26 | 0.046 | 4.31 | 0.23 | 0.21 | 1.000 | 2.00 | 0.23 | 0.20 | 1.000 | 3.11 | 0.08 | 0.08 | 1.000 | 2.99 | 0.10 | 0.36 | 0.000 |
| PIE101 | 8 | AT | GCGACAGTCACAATTAAGCTAC | CACCCAATTTTCAATCTGTG | 59.5 | 9.36 | 0.54 | 0.68 | 0.011 | 8.86 | 0.48 | 0.82 | 0.000 | 8.17 | 0.54 | 0.80 | 0.000 | 12.59 | 0.65 | 0.86 | 0.004 | 9.12 | 0.51 | 0.78 | 0.000 | 9.95 | 0.46 | 0.84 | 0.000 | 8.41 | 0.48 | 0.80 | 0.000 |
| PIE175 | 8 | TCT | ATTTGTCCACCGAGGTTCAG | CTTGGAAATGTGGGTTTTGG | 59.5 | 9.20 | 0.60 | 0.63 | 0.017 | 9.16 | 0.65 | 0.73 | 0.313 | 7.55 | 0.69 | 0.76 | 0.131 | 8.14 | 0.73 | 0.78 | 0.615 | 12.63 | 0.92 | 0.88 | 0.000 | 8.46 | 0.56 | 0.79 | 0.000 | 9.24 | 0.60 | 0.73 | 0.080 |
| VIT013 | 8 | AAC | GCCATGGCTTACACCACTTC | ATCAACAGAGCGGGTTTGTC | 59.5 | 5.38 | 0.32 | 0.69 | 0.000 | 5.75 | 0.46 | 0.72 | 0.000 | 5.96 | 0.56 | 0.68 | 0.035 | 6.65 | 0.48 | 0.64 | 0.017 | 5.80 | 0.40 | 0.78 | 0.000 | 4.95 | 0.45 | 0.71 | 0.000 | 4.00 | 0.48 | 0.71 | 0.000 |
| PIE204 | 8 | CAA | GCCATGGCTTACACCACTTC | ACACGTCGGCAAGCAAGTAG | 59.5 | 8.57 | 0.63 | 0.76 | 0.000 | 5.73 | 0.69 | 0.73 | 0.584 | 5.99 | 0.77 | 0.73 | 0.915 | 9.86 | 0.73 | 0.73 | 0.609 | 6.47 | 0.60 | 0.63 | 0.027 | 9.96 | 0.75 | 0.81 | 0.034 | 4.00 | 0.81 | 0.73 | 0.733 |
| VIT031.1 | 8 | CA | GCCACCCACCACTTTACAA | TTGAAGCTTGGAGGAGATGAA | 59.5 | 3.00 | 0.63 | 0.62 | 0.698 | 3.00 | 0.47 | 0.60 | 0.146 | 3.67 | 0.57 | 0.66 | 0.011 | 4.57 | 0.40 | 0.60 | 0.002 | 2.47 | 0.08 | 0.08 | 1.000 | 2.87 | 0.11 | 0.18 | 0.002 | 3.00 | 0.20 | 0.25 | 0.085 |
| VIT031.2 | 8 | CT | GCCACCCACCACTTTACAA | TTGAAGCTTGGAGGAGATGAA | 59.5 | 3.11 | 0.44 | 0.51 | 0.342 | 4.75 | 0.65 | 0.55 | 0.345 | 3.66 | 0.70 | 0.63 | 0.494 | 4.61 | 0.46 | 0.57 | 0.212 | 3.99 | 0.63 | 0.64 | 0.091 | 4.94 | 0.72 | 0.64 | 0.593 | 4.92 | 0.52 | 0.61 | 0.116 |
| PIE155 | 8 | GA | CCAGAAAATCAAGGCTGCTC | ACAACTCACTCATCACCACTGA | 59.5 | 11.83 | 0.77 | 0.88 | 0.008 | 14.07 | 0.71 | 0.88 | 0.000 | 10.16 | 0.60 | 0.75 | 0.000 | 15.34 | 0.77 | 0.86 | 0.185 | 9.61 | 0.85 | 0.82 | 0.039 | 16.54 | 0.69 | 0.83 | 0.001 | 16.14 | 0.75 | 0.80 | 0.073 |
| FIR044 | 8 | TGTT | TGTCCTTGTTTCCTACAAGGTG | TGCCGGAAACTAGTCCTCAA | 60.5 | 4.79 | 0.31 | 0.43 | 0.000 | 2.00 | 0.21 | 0.19 | 1.000 | 1.00 | 0.00 | 0.00 | 1.000 | 3.64 | 0.31 | 0.33 | 0.799 | 2.11 | 0.04 | 0.04 | 1.000 | 3.73 | 0.10 | 0.10 | 1.000 | 1.00 | 0.00 | 0.00 | 1.000 |
| VIT020 | 9 | CT | CCAAACACGGCTTACAAAGAA | CCGCTCGAGGCTTTTCTAGT | 58.5 | 12.74 | 0.85 | 0.90 | 0.440 | 12.23 | 0.48 | 0.89 | 0.000 | 11.65 | 0.84 | 0.87 | 0.001 | 12.90 | 0.85 | 0.89 | 0.095 | 7.75 | 0.73 | 0.78 | 0.170 | 9.59 | 0.71 | 0.78 | 0.032 | 10.54 | 0.48 | 0.86 | 0.000 |
| FIR033 | 9 | CT | AAGCTTTCCACCGACTTTCA | TGAAGTTGCGGAGAAGAGAGA | 59.5 | 8.78 | 0.22 | 0.62 | 0.000 | 3.99 | 0.19 | 0.59 | 0.000 | 4.53 | 0.31 | 0.59 | 0.000 | 4.91 | 0.33 | 0.63 | 0.000 | 4.72 | 0.17 | 0.60 | 0.000 | 6.29 | 0.43 | 0.76 | 0.000 | 3.72 | 0.46 | 0.65 | 0.000 |
| PIE259 | 9 | AC | TGGCTTTATGACACCTTTCG | CCTTTGGCATTTAGCATCAAA | 59.5 | 9.80 | 0.71 | 0.86 | 0.073 | 13.47 | 0.63 | 0.88 | 0.000 | 8.32 | 0.76 | 0.80 | 0.611 | 11.65 | 0.81 | 0.86 | 0.046 | 6.36 | 0.29 | 0.76 | 0.000 | 11.76 | 0.63 | 0.86 | 0.000 | 10.33 | 0.40 | 0.82 | 0.000 |
| WAG064 | 9 | CAG | ACTCAGCAGCAGCATCAAAG | CTGATTGGGAGGAGCATAGC | 59.5 | 7.55 | 0.73 | 0.84 | 0.023 | 6.87 | 0.85 | 0.80 | 0.774 | 7.84 | 0.85 | 0.79 | 0.109 | 7.34 | 0.79 | 0.79 | 0.113 | 6.13 | 0.60 | 0.72 | 0.346 | 7.20 | 0.88 | 0.79 | 0.838 | 6.69 | 0.77 | 0.75 | 0.889 |
| PIE059 | 9 | GAGCAC | GAGAGATGGAGCACGAGCA | GGGTCAGGACCATCTTCTTGT | 59.5 | 3.92 | 0.63 | 0.61 | 0.590 | 4.00 | 0.79 | 0.69 | 0.108 | 5.47 | 0.48 | 0.48 | 0.873 | 3.99 | 0.58 | 0.62 | 0.746 | 4.00 | 0.77 | 0.72 | 0.196 | 4.00 | 0.85 | 0.70 | 0.134 | 4.00 | 0.58 | 0.61 | 0.301 |
| PIE264 | 9 | AT | GCGCAGAGTCAGAGTTCCTT | ACGAGAATCAAGCGCAAAAT | 59.5 | 5.36 | 0.81 | 0.65 | 0.175 | 5.29 | 0.73 | 0.65 | 0.136 | 6.27 | 0.71 | 0.66 | 0.409 | 7.46 | 0.75 | 0.70 | 0.880 | 4.92 | 0.71 | 0.68 | 0.688 | 5.98 | 0.75 | 0.79 | 0.193 | 5.64 | 0.75 | 0.67 | 0.408 |
| ZQP15 | 9 | AG | CGATTTGATAATGACACTATGG | CATCGACTCATTGTTAAGCAC | 59.5 | 6.33 | 0.35 | 0.76 | 0.000 | 6.44 | 0.31 | 0.69 | 0.000 | 6.96 | 0.53 | 0.79 | 0.000 | 11.13 | 0.40 | 0.86 | 0.000 | 3.00 | 0.42 | 0.58 | 0.000 | 4.60 | 0.52 | 0.59 | 0.000 | 4.48 | 0.50 | 0.58 | 0.000 |
| FIR005 | 9 | CAA | GGCAACAAGGCCTTAAATCA | GGCAAGGGTTAAGGTGAACA | 59.5 | 5.56 | 0.77 | 0.74 | 0.765 | 6.57 | 0.69 | 0.70 | 0.225 | 6.61 | 0.77 | 0.73 | 0.000 | 6.00 | 0.69 | 0.76 | 0.006 | 4.98 | 0.71 | 0.64 | 0.539 | 5.62 | 0.60 | 0.73 | 0.085 | 5.98 | 0.73 | 0.73 | 0.128 |
| PIE081 | 9 | GACAGG | GCAGAAGAACGGAAGAAGCA | ATCCCGACTACCTCGAGCAT | 59.5 | 4.53 | 0.31 | 0.32 | 0.022 | 3.75 | 0.40 | 0.42 | 0.021 | 1.96 | 0.06 | 0.06 | 1.000 | 3.57 | 0.27 | 0.26 | 0.385 | 4.54 | 0.60 | 0.59 | 0.001 | 3.94 | 0.56 | 0.50 | 0.139 | 4.72 | 0.56 | 0.54 | 0.001 |
| VIT025 | 9 | AG | AGGCAGCCTCTCCTACCAAG | ACTGCAAGGACCAAGATGCT | 59.5 | 8.75 | 0.49 | 0.76 | 0.000 | 11.14 | 0.41 | 0.81 | 0.000 | 7.74 | 0.48 | 0.64 | 0.022 | 13.26 | 0.52 | 0.85 | 0.000 | 6.57 | 0.50 | 0.70 | 0.031 | 7.14 | 0.31 | 0.65 | 0.000 | 8.07 | 0.42 | 0.62 | 0.006 |
| FIR023 | 10 | AAC | CACACCCATCATCAAACACC | CGCTGGTAGGACCAGAACAT | 59.5 | 6.84 | 0.71 | 0.79 | 0.511 | 6.88 | 0.85 | 0.80 | 0.227 | 6.53 | 0.83 | 0.77 | 0.809 | 6.58 | 0.74 | 0.76 | 0.827 | 4.91 | 0.67 | 0.64 | 0.476 | 6.24 | 0.77 | 0.78 | 0.788 | 4.00 | 0.71 | 0.70 | 0.765 |
| POR028 | 10 | AAT | TCTTCAAAACCCCTTCAACG | TGAGGATGCCATAACCATCA | 59.5 | 7.66 | 0.71 | 0.82 | 0.023 | 8.07 | 0.56 | 0.76 | 0.001 | 10.77 | 0.38 | 0.84 | 0.000 | 9.46 | 0.56 | 0.82 | 0.000 | 7.94 | 0.38 | 0.81 | 0.000 | 6.96 | 0.54 | 0.78 | 0.000 | 12.42 | 0.60 | 0.81 | 0.002 |
| ZQR96 | 10 | AG | CCCAGTCACATCCACTACTGTCC | GGTTGGGAAAAGGAGATCAGA | 59.5 | 15.49 | 0.85 | 0.91 | 0.058 | 12.23 | 0.83 | 0.85 | 0.988 | 11.88 | 0.88 | 0.82 | 0.809 | 14.21 | 0.92 | 0.89 | 0.407 | 10.70 | 0.77 | 0.80 | 0.076 | 13.50 | 0.79 | 0.84 | 0.051 | 14.46 | 0.90 | 0.88 | 0.538 |
| PIE148 | 10 | GA | GGTGTTGAAGTTTTTGTTTCC | ATCCTTTGGTGGGTGTGG | 55.5 | 15.39 | 0.91 | 0.91 | 0.547 | 15.99 | 0.85 | 0.92 | 0.204 | 12.39 | 0.85 | 0.81 | 0.099 | 16.48 | 0.90 | 0.88 | 0.301 | 13.51 | 0.98 | 0.89 | 0.022 | 17.05 | 0.89 | 0.89 | 0.488 | 15.61 | 0.90 | 0.90 | 0.551 |
| ZQR11 | 10 | AG | CCTTGAACTCGAAGGTGTCCTT | GTAGGTCAAAACCATTGGTTGACT | 58.0 | 9.94 | 0.39 | 0.70 | 0.000 | 6.87 | 0.07 | 0.73 | 0.000 | 8.80 | 0.49 | 0.80 | 0.000 | 12.89 | 0.60 | 0.83 | 0.000 | 11.63 | 0.82 | 0.85 | 0.509 | 13.05 | 0.51 | 0.80 | 0.000 | 11.89 | 0.80 | 0.87 | 0.002 |
| WAG020 | 10 | CAA | AGATCAGGATGTGGAGGTGC | AAATCCAAACCCCAGTTTC | 59.5 | 2.00 | 1.00 | 0.50 | 0.000 | 6.11 | 0.55 | 0.70 | 0.025 | 4.64 | 0.42 | 0.59 | 0.000 | 8.24 | 0.58 | 0.77 | 0.004 | 2.92 | 0.96 | 0.53 | 0.000 | 5.94 | 0.63 | 0.64 | 0.488 | 5.62 | 0.60 | 0.61 | 0.828 |
| PIE186 | 10 | TAA | CCATCACAAAACTGCGCTATC | CACGCTCATATACACCCAAATG | 58.0 | 6.34 | 0.70 | 0.69 | 0.781 | 6.44 | 0.75 | 0.77 | 0.267 | 6.28 | 0.69 | 0.66 | 0.138 | 6.66 | 0.88 | 0.74 | 0.088 | 7.80 | 0.81 | 0.82 | 0.488 | 9.23 | 0.88 | 0.84 | 0.377 | 8.93 | 0.79 | 0.83 | 0.036 |
| PIE053 | 10 | AAAGAA | GAGCTGCATATTTTCCTTAGTCA | GAACTCTTCCTCTTCATCTTCTGC | 59.5 | 7.28 | 0.75 | 0.75 | 0.908 | 5.86 | 0.83 | 0.78 | 0.612 | 5.96 | 0.67 | 0.73 | 0.256 | 8.91 | 0.71 | 0.81 | 0.356 | 5.61 | 0.65 | 0.58 | 0.302 | 6.80 | 0.52 | 0.59 | 0.031 | 6.43 | 0.67 | 0.66 | 0.691 |
| PIE187 | 10 | ACA | TTCTCAGCCCTTCGATCAAC | TTTGTGGTGGTCTTTGTTGG | 58.0 | 5.11 | 0.81 | 0.68 | 0.057 | 5.25 | 0.77 | 0.73 | 0.974 | 4.00 | 0.68 | 0.69 | 0.958 | 5.35 | 0.73 | 0.72 | 0.620 | 4.95 | 0.69 | 0.67 | 0.106 | 4.86 | 0.63 | 0.60 | 0.420 | 4.66 | 0.62 | 0.63 | 0.656 |
| PIE144 | 11 | CT | TGCAAGGTGGGAACTTTGAT | CGGCATTAAAAGAGCCTTCTC | 59.5 | 12.13 | 0.85 | 0.84 | 0.404 | 14.85 | 0.94 | 0.89 | 0.349 | 13.32 | 0.91 | 0.87 | 0.171 | 15.35 | 0.94 | 0.86 | 0.909 | 13.61 | 0.88 | 0.84 | 0.313 | 15.99 | 0.90 | 0.91 | 0.004 | 13.67 | 0.90 | 0.91 | 0.208 |
| PIE257 | 11 | CT | GCCAACGAGAGCTAAGTTGC | GCAAGGTCAAGGTCAGGGTA | 59.5 | 9.66 | 0.88 | 0.81 | 0.634 | 8.12 | 0.69 | 0.82 | 0.252 | 8.16 | 0.70 | 0.80 | 0.049 | 9.78 | 0.77 | 0.85 | 0.158 | 9.32 | 0.85 | 0.85 | 0.424 | 9.54 | 0.73 | 0.80 | 0.173 | 9.91 | 0.77 | 0.86 | 0.269 |
| WAG079 | 11 | CT | CAAAACGACAAGGAAGGAGC | CTAACACTGGCTGCTGTTGC | 58.0 | 13.88 | 0.96 | 0.88 | 0.815 | 14.04 | 0.87 | 0.88 | 0.088 | 12.06 | 0.75 | 0.81 | 0.070 | 15.62 | 0.91 | 0.91 | 0.249 | 17.36 | 0.91 | 0.88 | 0.291 | 11.79 | 0.58 | 0.60 | 0.239 | 11.15 | 0.80 | 0.74 | 0.523 |
| PIE036 | 11 | CTT | TGACCTCTGAATTCCAAGAACC | TATGAAGGTGTTGATTCCCTTTT | 59.5 | 5.90 | 0.38 | 0.45 | 0.047 | 7.86 | 0.35 | 0.70 | 0.000 | 5.53 | 0.28 | 0.53 | 0.000 | 6.21 | 0.38 | 0.67 | 0.000 | 6.52 | 0.24 | 0.62 | 0.000 | 7.90 | 0.31 | 0.74 | 0.000 | 7.80 | 0.58 | 0.68 | 0.009 |
| FIR104 | 11 | GGT | TTAACTCGGTTTGCGACTCA | AGCACGTGACTCGACCTGTA | 60.5 | 7.85 | 0.88 | 0.79 | 0.212 | 12.74 | 0.79 | 0.89 | 0.177 | 11.28 | 0.66 | 0.76 | 0.079 | 10.46 | 0.81 | 0.86 | 0.509 | 8.65 | 0.67 | 0.72 | 0.070 | 5.98 | 0.69 | 0.73 | 0.635 | 9.54 | 0.69 | 0.77 | 0.196 |
| PIE202 | 11 | AG | GGGAAGAGAGAAAGAGATGGAAG | GCTGGTGATACAAGTGTTTGG | 59.5 | 2.65 | 0.13 | 0.28 | 0.000 | 2.00 | 0.98 | 0.50 | 0.000 | 2.88 | 1.00 | 0.52 | 0.000 | 2.00 | 0.73 | 0.46 | 0.000 | 2.00 | 0.15 | 0.14 | 1.000 | 2.00 | 0.06 | 0.26 | 0.000 | 1.00 | 0.00 | 0.00 | 1.000 |
| PIE033 | 11 | TTC | AAGCGACTTCTCCTGTCTGG | CCATGCAACAACCAAATGAT | 59.5 | 22.64 | 0.90 | 0.94 | 0.004 | 20.07 | 0.92 | 0.94 | 0.107 | 20.59 | 0.92 | 0.93 | 0.152 | 21.57 | 0.81 | 0.92 | 0.013 | 17.90 | 0.91 | 0.92 | 0.007 | 25.59 | 0.94 | 0.95 | 0.405 | 23.85 | 0.91 | 0.94 | 0.045 |
| PIE102 | 11 | GAA | ACCTTCCATGCTCAAAGATG | CATTTCTGAAAAGCCCAATATG | 59.5 | 7.97 | 0.53 | 0.58 | 0.118 | 9.89 | 0.81 | 0.80 | 0.379 | 9.86 | 0.85 | 0.82 | 0.234 | 12.32 | 0.79 | 0.84 | 0.281 | 8.62 | 0.77 | 0.75 | 0.093 | 5.73 | 0.58 | 0.57 | 0.220 | 8.67 | 0.96 | 0.79 | 0.261 |
| PIE089 | 12 | TTAGGG | CGCCCTTAGACCAGGATTTT | TTGAGTCTTGCTTGCCATTG | 59.0 | 5.30 | 0.51 | 0.61 | 0.000 | 7.22 | 0.52 | 0.78 | 0.000 | 7.30 | 0.52 | 0.80 | 0.000 | 7.21 | 0.56 | 0.74 | 0.118 | 8.39 | 0.60 | 0.76 | 0.001 | 7.24 | 0.48 | 0.80 | 0.000 | 7.74 | 0.53 | 0.78 | 0.001 |
| VIT010 | 12 | TC | CACGTGCATGTAAAGGTGCT | CTGTGTCTGTGATGCGTGTTT | 58.0 | 5.68 | 0.57 | 0.61 | 0.171 | 3.88 | 0.65 | 0.58 | 0.474 | 5.82 | 0.58 | 0.71 | 0.026 | 7.58 | 0.52 | 0.48 | 0.419 | 7.65 | 0.77 | 0.68 | 0.558 | 5.92 | 0.53 | 0.63 | 0.292 | 7.42 | 0.55 | 0.70 | 0.026 |
| VIT050 | 12 | ATTTTG | TGGTTCCCTCAAGGTTGGT | TGGTTTCAAACACAAACCAG | 58.5 | 4.51 | 0.52 | 0.55 | 0.874 | 3.88 | 0.63 | 0.62 | 0.933 | 3.53 | 0.25 | 0.53 | 0.000 | 5.39 | 0.48 | 0.67 | 0.001 | 3.99 | 0.33 | 0.43 | 0.001 | 3.61 | 0.29 | 0.31 | 0.228 | 3.92 | 0.33 | 0.38 | 0.039 |
| PIE196 | 12 | GTG | GTCTGCGTGTCTCAACTTCG | CACAGCACTCTCACCACAGC | 59.5 | 7.07 | 0.13 | 0.62 | 0.000 | 6.38 | 0.21 | 0.52 | 0.000 | 2.88 | 0.04 | 0.25 | 0.000 | 8.84 | 0.35 | 0.52 | 0.000 | 9.02 | 0.61 | 0.65 | 0.167 | 10.51 | 0.59 | 0.70 | 0.054 | 6.99 | 0.81 | 0.80 | 0.000 |
| VIT037 | 12 | TC | GAAGCCACCAAAAGCTCTGT | GAGTGGAAATGAAAAACCCAGA | 59.5 | 9.79 | 0.79 | 0.84 | 0.390 | 11.64 | 0.69 | 0.84 | 0.043 | 12.72 | 0.77 | 0.87 | 0.133 | 10.42 | 0.52 | 0.85 | 0.000 | 8.95 | 0.81 | 0.81 | 0.023 | 9.83 | 0.73 | 0.81 | 0.319 | 10.98 | 0.77 | 0.81 | 0.678 |
| PIE236 | 12 | GAA | TCAATCCCATCTTCCTGCTC | TCTATGCGATTCCGCTTACC | 59.5 | 5.47 | 0.63 | 0.66 | 0.144 | 6.11 | 0.46 | 0.57 | 0.002 | 5.85 | 0.72 | 0.68 | 0.550 | 6.78 | 0.69 | 0.69 | 0.434 | 7.03 | 0.79 | 0.69 | 0.041 | 5.59 | 0.57 | 0.67 | 0.022 | 4.70 | 0.58 | 0.63 | 0.559 |
| ZQR112 | 12 | AG | TTCTTGCTTTGGTGCGCG | GTGGTCAGAGACTCGGTAAGTATTC | 58.5 | 6.28 | 0.79 | 0.71 | 0.625 | 4.97 | 0.73 | 0.74 | 0.842 | 5.41 | 0.48 | 0.56 | 0.018 | 9.33 | 0.75 | 0.77 | 0.707 | 3.52 | 0.56 | 0.46 | 0.398 | 3.57 | 0.44 | 0.51 | 0.217 | 4.43 | 0.50 | 0.55 | 0.118 |
| POR020 | 12 | GAT | CAAATCCGCTCAACTTGTCA | ACCATCGTCTCCAACATCAA | 59.5 | 8.23 | 0.75 | 0.71 | 0.620 | 7.43 | 0.77 | 0.74 | 0.463 | 6.63 | 0.73 | 0.77 | 0.116 | 7.77 | 0.89 | 0.81 | 0.283 | 8.28 | 0.58 | 0.69 | 0.026 | 6.88 | 0.58 | 0.75 | 0.005 | 6.72 | 0.58 | 0.78 | 0.058 |
| ZQR30 | 12 | AG | TGCTCCGTCATAATCTTGCTCTGA | GCAATCCTATCATGCACATGCACAT | 59.5 | 19.74 | 0.88 | 0.92 | 0.321 | 17.97 | 0.85 | 0.92 | 0.031 | 11.03 | 0.53 | 0.83 | 0.000 | 22.79 | 0.94 | 0.94 | 0.135 | 11.85 | 0.88 | 0.86 | 0.310 | 23.24 | 0.94 | 0.93 | 0.666 | 17.77 | 0.75 | 0.91 | 0.004 |
| | | | | Average 2bp | | 10.29 | 0.67 | 0.74 | | 9.95 | 0.59 | 0.71 | | 9.60 | 0.63 | 0.72 | | 11.98 | 0.66 | 0.77 | | 9.12 | 0.63 | 0.71 | | 11.24 | 0.64 | 0.73 | | 10.44 | 0.62 | 0.73 | |
| | | | | Average 3-6 bp | | 5.95 | 0.53 | 0.58 | | 5.90 | 0.52 | 0.59 | | 5.72 | 0.47 | 0.53 | | 6.82 | 0.57 | 0.63 | | 5.83 | 0.57 | 0.59 | | 6.17 | 0.54 | 0.59 | | 6.32 | 0.56 | 0.61 | |
| | | | | Average-All | | 8.08 | 0.60 | 0.66 | | 7.89 | 0.56 | 0.65 | | 7.62 | 0.55 | 0.63 | | 9.35 | 0.61 | 0.70 | | 7.44 | 0.60 | 0.65 | | 8.66 | 0.59 | 0.66 | | 8.34 | 0.59 | 0.67 | |

**Table S2-2:**

| Locus | CAB-fg | CJA-fg | FRO-fg | IZK-fg | CAB-py | IZK-py | SNE-py |
|---|---|---|---|---|---|---|---|
| PIE076 | -0.05 | -0.54 | -0.05 | -0.19 | -0.21 | -0.03 | -0.06 |
| FIR073 | 0.10 | 0.13 | 0.08 | 0.16 | 0.23 | 0.15 | 0.15 |
| VIT026 | 0.24 | 0.19 | 0.21 | 0.27 | 0.14 | 0.22 | 0.18 |
| PIE238 | 0.01 | 0.00 | 0.10 | 0.07 | 0.14 | 0.19 | 0.14 |
| POR040 | 0.02 | 0.08 | 0.18 | 0.02 | 0.04 | 0.01 | 0.04 |
| VIT007 | 0.01 | 0.29 | 0.19 | -0.01 | 0.15 | 0.09 | 0.13 |
| WAG011 | 0.03 | 0.00 | -0.18 | 0.07 | 0.11 | -0.03 | 0.05 |
| FIR040 | 0.03 | -0.01 | -0.01 | 0.05 | 0.02 | 0.05 | -0.03 |
| PIE223 | 0.01 | 0.06 | 0.14 | 0.06 | 0.14 | 0.10 | 0.09 |
| FIR065 | 0.06 | 0.21 | 0.05 | 0.04 | 0.02 | 0.07 | 0.03 |
| ZQR87 | -0.03 | 0.08 | -0.03 | 0.00 | -0.03 | -0.08 | 0.01 |
| FIR048 | -0.21 | 0.05 | -0.04 | -0.03 | 0.03 | 0.07 | -0.08 |
| FIR013 | 0.02 | -0.03 | -0.07 | 0.01 | 0.03 | -0.05 | -0.05 |
| ZQP119 | 0.06 | 0.18 | 0.23 | 0.17 | 0.17 | -0.06 | 0.14 |
| POR038 | -0.03 | 0.01 | 0.14 | 0.06 | -0.01 | 0.00 | 0.22 |
| FIR032 | 0.09 | 0.05 | 0.16 | 0.07 | 0.00 | 0.05 | 0.17 |
| PIE197 | -0.02 | 0.02 | 0.02 | -0.02 | -0.23 | -0.02 | -0.04 |
| PIE075 | 0.13 | 0.29 | 0.21 | 0.34 | 0.03 | 0.36 | 0.26 |
| PIE135 | 0.21 | 0.13 | 0.16 | 0.16 | -0.02 | -0.03 | 0.10 |
| FIR047 | 0.00 | -0.01 | 0.09 | -0.02 | -0.04 | -0.04 | -0.04 |
| GOT021 | -0.03 | -0.05 | 0.00 | -0.18 | 0.01 | -0.05 | -0.07 |
| PIE021 | 0.04 | 0.11 | 0.07 | 0.21 | 0.18 | 0.17 | 0.11 |
| VIT107 | 0.02 | 0.01 | 0.00 | 0.00 | 0.00 | 0.00 | -0.23 |
| POR011 | -0.01 | 0.00 | -0.06 | -0.01 | -0.02 | -0.04 | -0.05 |
| VIT022 | 0.13 | -0.05 | -0.15 | 0.09 | 0.04 | 0.07 | 0.05 |
| PIE242 | -0.07 | 0.01 | -0.03 | 0.09 | 0.06 | 0.13 | 0.08 |
| PIE051 | -0.04 | 0.08 | -0.13 | -0.18 | 0.07 | -0.06 | 0.04 |
| PIE088 | 0.37 | 0.07 | 0.16 | 0.34 | 0.34 | 0.37 | 0.34 |
| VIT043 | -0.04 | 0.11 | -0.01 | 0.20 | 0.07 | 0.01 | 0.07 |
| PIE082 | 0.02 | -0.07 | 0.20 | 0.05 | 0.03 | -0.10 | -0.11 |
| PIE216 | -0.20 | 0.10 | 0.03 | -0.12 | -0.08 | -0.12 | 0.07 |
| PIE228 | 0.05 | 0.02 | 0.03 | 0.04 | 0.00 | 0.00 | 0.01 |
| POR029 | -0.06 | -0.03 | 0.03 | -0.05 | -0.10 | 0.03 | 0.03 |
| VIT003 | 0.11 | 0.26 | 0.08 | 0.20 | 0.15 | 0.14 | 0.21 |
| VIT018 | -0.01 | -0.04 | 0.28 | 0.11 | -0.02 | -0.04 | -0.01 |
| VIT017 | -0.09 | 0.04 | -0.09 | -0.03 | -0.10 | -0.01 | -0.06 |
| PIE002 | 0.18 | 0.08 | 0.10 | 0.00 | 0.05 | 0.12 | 0.02 |
| PIE029 | 0.20 | 0.21 | 0.13 | 0.13 | 0.11 | 0.09 | 0.10 |
| WAG043 | 0.04 | 0.06 | 0.06 | 0.09 | 0.02 | 0.04 | 0.12 |
| POR014 | -0.02 | 0.07 | 0.07 | 0.20 | 0.25 | 0.15 | 0.09 |
| GOT065 | -0.06 | 0.00 | 0.09 | 0.05 | -0.01 | -0.04 | -0.05 |
| VIT023 | -0.05 | 0.00 | 0.06 | 0.04 | -0.07 | -0.11 | 0.11 |
| FIR106 | -0.22 | 0.34 | 0.00 | 0.18 | -0.76 | -0.04 | -0.08 |
| POR030 | -0.01 | 0.15 | -0.17 | -0.13 | -0.01 | -0.01 | -0.03 |
| POR039 | 0.01 | 0.02 | -0.06 | -0.23 | -0.03 | -0.23 | -0.06 |
| VIT021 | -0.02 | 0.02 | -0.06 | -0.01 | 0.01 | -0.04 | -0.03 |
| POR016 | 0.01 | 0.10 | 0.24 | 0.16 | -0.09 | 0.16 | 0.31 |
| FIR021 | 0.21 | 0.29 | 0.26 | 0.20 | 0.10 | 0.02 | 0.08 |
| POR024 | 0.02 | 0.16 | 0.00 | 0.02 | -0.31 | 0.02 | -0.02 |
| PIE125 | -0.08 | 0.16 | 0.07 | 0.07 | -0.08 | 0.08 | -0.07 |
| POR034 | 0.02 | -0.06 | -0.06 | -0.01 | -0.03 | -0.02 | 0.12 |
| VIT028 | 0.22 | 0.24 | 0.30 | 0.22 | 0.11 | 0.11 | 0.13 |
| POR027 | -0.10 | -0.10 | 0.03 | -0.10 | 0.06 | -0.02 | 0.10 |
| PIE071 | 0.11 | 0.08 | 0.01 | 0.03 | 0.04 | 0.06 | 0.12 |
| PIE219 | -0.06 | 0.09 | -0.11 | -0.15 | -0.06 | -0.13 | -0.09 |
| PIE137 | -0.08 | 0.03 | -0.04 | 0.04 | 0.05 | 0.01 | 0.02 |
| VIT046 | 0.00 | -0.03 | -0.13 | 0.09 | -0.05 | 0.02 | 0.05 |
| FIR043 | 0.09 | 0.14 | -0.01 | 0.07 | 0.17 | -0.05 | 0.02 |
| FIR030 | 0.02 | 0.22 | 0.21 | 0.09 | 0.22 | 0.08 | 0.18 |
| FIR095 | 0.02 | 0.07 | -0.02 | 0.04 | -0.02 | 0.02 | -0.03 |
| PIE127 | -0.02 | -0.08 | -0.03 | 0.08 | -0.07 | 0.06 | 0.08 |
| VIT009 | 0.13 | -0.02 | 0.16 | 0.05 | 0.08 | 0.08 | 0.00 |
| POR025 | 0.06 | -0.34 | -0.02 | 0.07 | -0.13 | 0.05 | 0.04 |
| WAG021 | -0.05 | -0.04 | 0.01 | 0.02 | 0.04 | -0.02 | -0.03 |
| POR008 | 0.28 | -0.05 | -0.04 | 0.19 | 0.00 | 0.07 | -0.03 |
| PIE023 | -0.03 | 0.16 | -0.01 | -0.12 | -0.01 | -0.04 | 0.27 |
| PIE101 | 0.11 | 0.20 | 0.16 | 0.12 | 0.17 | 0.23 | 0.20 |
| PIE175 | 0.04 | 0.06 | 0.05 | 0.03 | -0.03 | 0.15 | 0.08 |
| VIT013 | 0.25 | 0.17 | 0.09 | 0.13 | 0.24 | 0.18 | 0.15 |
| PIE204 | 0.10 | 0.03 | -0.03 | 0.02 | 0.04 | 0.04 | -0.06 |
| VIT031.1 | -0.01 | 0.10 | 0.06 | 0.16 | -0.03 | 0.13 | 0.08 |
| VIT031.2 | 0.07 | -0.10 | -0.06 | 0.10 | 0.01 | -0.07 | 0.07 |
| PIE155 | 0.06 | 0.10 | 0.12 | 0.05 | -0.03 | 0.10 | 0.03 |
| FIR044 | 0.11 | -0.11 | 0.00 | 0.02 | -0.02 | -0.05 | 0.00 |
| VIT020 | 0.02 | 0.23 | 0.02 | 0.02 | 0.03 | 0.05 | 0.22 |
| FIR033 | 0.30 | 0.30 | 0.22 | 0.22 | 0.33 | 0.20 | 0.12 |
| PIE259 | 0.09 | 0.14 | 0.03 | 0.03 | 0.29 | 0.13 | 0.25 |
| WAG064 | 0.06 | -0.03 | -0.05 | -0.01 | 0.08 | -0.05 | -0.01 |
| PIE059 | -0.01 | -0.08 | -0.01 | 0.04 | -0.03 | -0.12 | 0.00 |
| PIE264 | -0.01 | -0.07 | -0.04 | -0.03 | -0.02 | 0.02 | -0.07 |
| ZQP15 | 0.25 | 0.27 | 0.15 | 0.26 | 0.11 | 0.05 | 0.06 |
| FIR005 | -0.01 | 0.01 | -0.02 | 0.04 | -0.07 | 0.09 | 0.01 |
| PIE081 | 0.02 | 0.01 | -0.03 | -0.01 | -0.03 | -0.08 | -0.04 |
| VIT025 | 0.18 | 0.24 | 0.12 | 0.19 | 0.14 | 0.23 | 0.15 |
| FIR023 | 0.05 | -0.03 | -0.04 | 0.01 | -0.03 | 0.01 | -0.01 |
| POR028 | 0.07 | 0.14 | 0.27 | 0.15 | 0.26 | 0.15 | 0.13 |
| ZQR96 | 0.03 | 0.01 | -0.06 | -0.02 | 0.02 | 0.03 | -0.01 |
| PIE148 | 0.00 | 0.04 | -0.02 | -0.01 | -0.05 | 0.00 | 0.00 |
| ZQR11 | 0.22 | 0.42 | 0.18 | 0.13 | 0.02 | 0.18 | 0.04 |
| WAG020 | -0.01 | 0.10 | 0.16 | 0.12 | -0.63 | -0.01 | 0.00 |
| PIE186 | -0.05 | 0.02 | -0.03 | -0.11 | 0.00 | -0.03 | 0.02 |
| PIE053 | 0.00 | -0.04 | 0.04 | 0.07 | -0.08 | 0.09 | 0.00 |
| PIE187 | -0.13 | -0.04 | 0.01 | 0.00 | -0.03 | -0.02 | 0.02 |
| PIE144 | 0.00 | -0.03 | -0.02 | -0.05 | -0.04 | 0.01 | 0.01 |
| PIE257 | -0.05 | 0.08 | 0.06 | 0.04 | 0.00 | 0.05 | 0.05 |
| WAG079 | -0.04 | 0.00 | 0.03 | 0.00 | -0.02 | 0.01 | -0.04 |
| PIE036 | 0.08 | 0.24 | 0.22 | 0.21 | 0.27 | 0.27 | 0.07 |
| FIR104 | -0.06 | 0.05 | 0.07 | 0.03 | 0.04 | 0.03 | 0.05 |
| PIE202 | 0.20 | -0.86 | -0.82 | -0.48 | -0.08 | 0.25 | 0.00 |
| PIE033 | 0.02 | 0.01 | 0.01 | 0.06 | 0.02 | 0.01 | 0.01 |
| PIE102 | 0.05 | -0.02 | -0.03 | 0.03 | -0.02 | -0.01 | -0.11 |
| PIE089 | 0.10 | 0.16 | 0.17 | 0.12 | 0.11 | 0.19 | 0.16 |
| VIT010 | 0.05 | -0.07 | 0.08 | -0.08 | -0.09 | 0.09 | 0.10 |
| VIT050 | 0.02 | -0.01 | 0.24 | 0.14 | 0.11 | 0.03 | 0.04 |
| PIE196 | 0.36 | 0.27 | 0.27 | 0.18 | 0.04 | 0.09 | -0.03 |
| VIT037 | 0.03 | 0.09 | 0.06 | 0.19 | -0.01 | 0.05 | 0.03 |
| PIE236 | 0.03 | 0.12 | -0.04 | 0.00 | -0.09 | 0.06 | 0.04 |
| ZQR112 | -0.06 | 0.01 | 0.09 | 0.01 | -0.14 | 0.07 | 0.09 |
| POR020 | -0.04 | -0.03 | 0.01 | -0.05 | 0.67 | 0.11 | 0.12 |
| ZQR30 | 0.03 | 0.04 | 0.20 | 0.00 | -0.02 | -0.01 | 0.09 |
| Nb significant | 29 | 45 | 39 | 42 | 28 | 31 | 38 |

**Table S2-3**

| | G$_{ST}$_Nei | | | | | | |
|---|---|---|---|---|---|---|---|
| D_Jost | **CAB-fg** | **CJA-fg** | **FRO-fg** | **IZK-fg** | **CAB-py** | **IZK-py** | **SNE-py** |
| **CAB-fg** | | 0.069 (0.066-0.072) | 0.071 (0.068-0.074) | 0.056 (0.053-0.058) | 0.049 (0.046-0.052) | 0.073 (0.070-0.077) | 0.081 (0.078-0.084) |
| **CJA-fg** | 0.288 (0.277-0.293) | | 0.064 (0.060-0.067) | 0.057 (0.054-0.060) | 0.082 (0.079-0.085) | 0.083 (0.079-0.086) | 0.050 (0.047-0.054) |
| **FRO-fg** | 0.280 (0.270-0.291) | 0.247 (0.237-0.258) | | 0.046 (0.043-0.048) | 0.088 (0.085-0.092) | 0.074 (0.070-0.077) | 0.068 (0.065-0.072) |
| **IZK-fg** | 0.258 (0.247-0.268) | 0.260 (0.250-0.271) | 0.193 (0.182-0.204) | | 0.069 (0.066-0.072) | 0.035 (0.032-0.037) | 0.056 (0.053-0.059) |
| **CAB-py** | 0.201 (0.190-0.212) | 0.343 (0.332-0.355) | 0.349 (0.337-0.361) | 0.319 (0.308-0.330) | | 0.064 (0.061-0.067) | 0.075 (0.072-0.078) |
| **IZK-py** | 0.318 (0.306-0.330) | 0.357 (0.344-0.370) | 0.296 (0.284-0.308) | 0.159 (0.148-0.171) | 0.268 (0.259-0.278) | | 0.055 (0.053-0.058) |
| **SNE-py** | 0.360 (0.349-0.371) | 0.213 (0.200-0.225) | 0.277 (0.266-0.289) | 0.265 (0.253-0.278) | 0.326 (0.316-0.337) | 0.242 (0.232-0.251) | |

**Figure S2-1a**

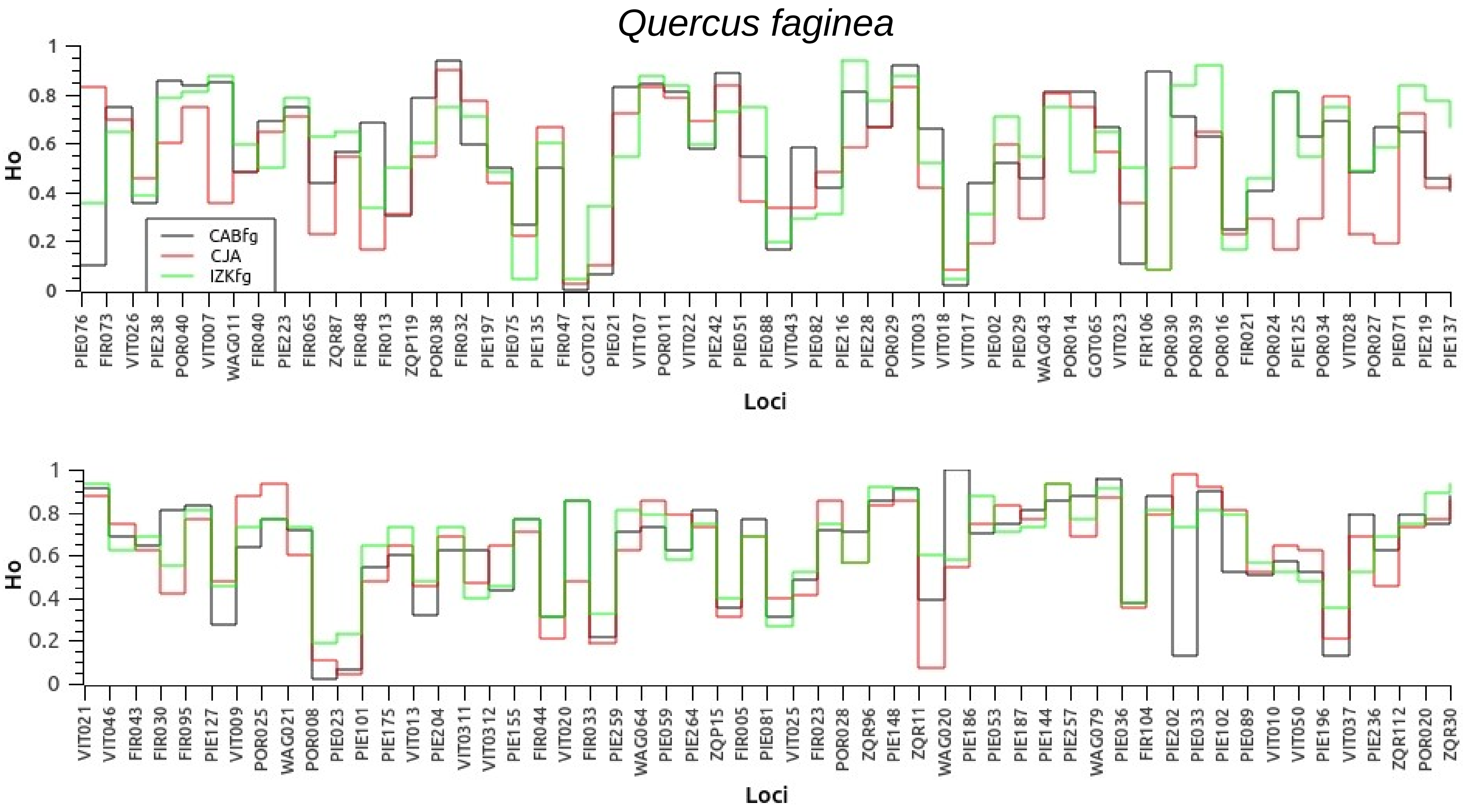

Quercus faginea
Ho
1
0.8
0.6
0.4
0.2
0
CABFg
CJA
IZKfg
PIE076 FIR073 VIT026 PIE238 POR040 VIT007 WAG011 FIR040 PIE223 FIR065 ZQR87 FIR048 FIR013 ZQP119 POR038 FIR032 PIE197 PIE075 PIE135 FIR047 GOT021 PIE021 VIT107 POR011 VIT022 PIE242 PIE051 PIE088 VIT043 PIE082 PIE216 PIE228 POR029 VIT003 VIT018 VIT017 PIE002 PIE029 WAG043 POR014 GOT065 VIT023 FIR106 POR030 POR039 POR016 FIR021 POR024 PIE125 POR034 VIT028 POR027 PIE071 PIE219 PIE137
Loci
VIT021 VIT046 FIR043 FIR030 FIR095 PIE127 VIT009 POR025 WAG021 POR008 PIE023 PIE101 PIE175 VIT013 PIE204 VIT0311 VIT0312 PIE155 FIR044 VIT020 FIR033 PIE259 WAG064 PIE059 PIE264 ZQP15 FIR005 PIE081 VIT025 FIR023 POR028 ZQR96 PIE148 ZQR11 WAG020 PIE186 PIE053 PIE187 PIE144 PIE257 WAG079 PIE036 FIR104 PIE202 PIE033 PIE102 PIE089 VIT010 VIT050 PIE196 VIT037 PIE236 ZQR112 POR020 ZQR30
Loci

**Figure S2-1b**

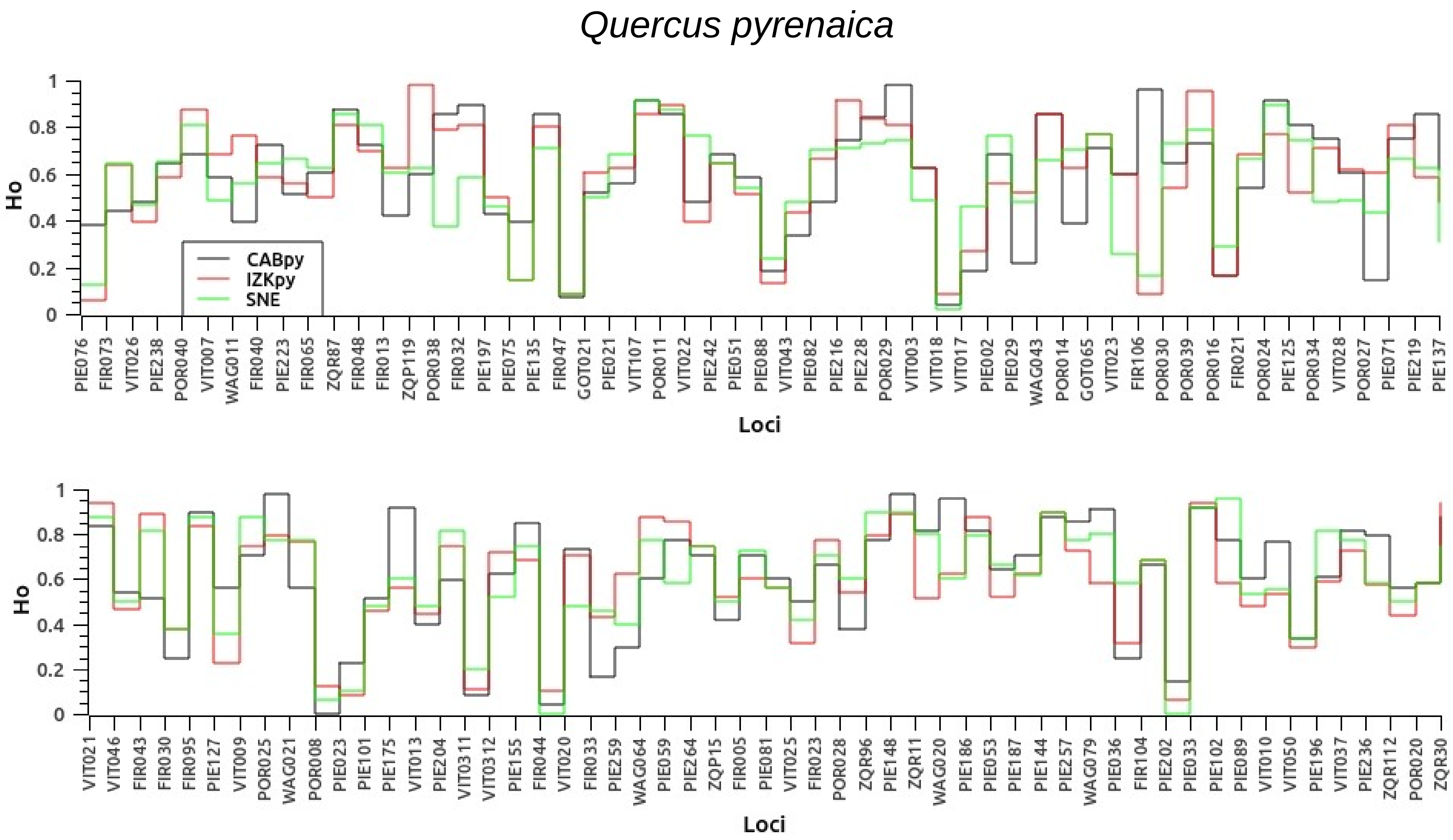
Quercus pyrenaica
Ho
Loci
CABpy
IZKpy
SNE
1
0.8
0.6
0.4
0.2
0
PIE076 FIR073 VIT026 PIE238 POR040 VIT007 WAG011 FIR040 PIE223 FIR065 ZQR87 FIR048 FIR013 ZQP119 POR038 FIR032 PIE197 PIE075 PIE135 FIR047 GOT021 PIE021 VIT107 POR011 VIT022 PIE242 PIE051 PIE088 VIT043 PIE082 PIE216 PIE228 POR029 VIT003 VIT018 VIT017 PIE002 PIE029 WAG043 POR014 GOT065 VIT023 FIR106 POR030 POR039 POR016 FIR021 POR024 PIE125 POR034 VIT028 POR027 PIE071 PIE219 PIE137
VIT021 VIT046 FIR043 FIR030 FIR095 PIE127 VIT009 POR025 WAG021 POR008 PIE023 PIE101 PIE175 VIT013 PIE204 VIT0311 VIT0312 PIE155 FIR044 VIT020 FIR033 PIE259 WAG064 PIE059 PIE264 ZQP15 FIR005 PIE081 VIT025 FIR023 POR028 ZQR96 PIE148 ZQR11 WAG020 PIE186 PIE053 PIE187 PIE144 PIE257 WAG079 PIE036 FIR104 PIE202 PIE033 PIE102 PIE089 VIT010 VIT050 PIE196 VIT037 PIE236 ZQR112 POR020 ZQR30

**Figure S2-2:**

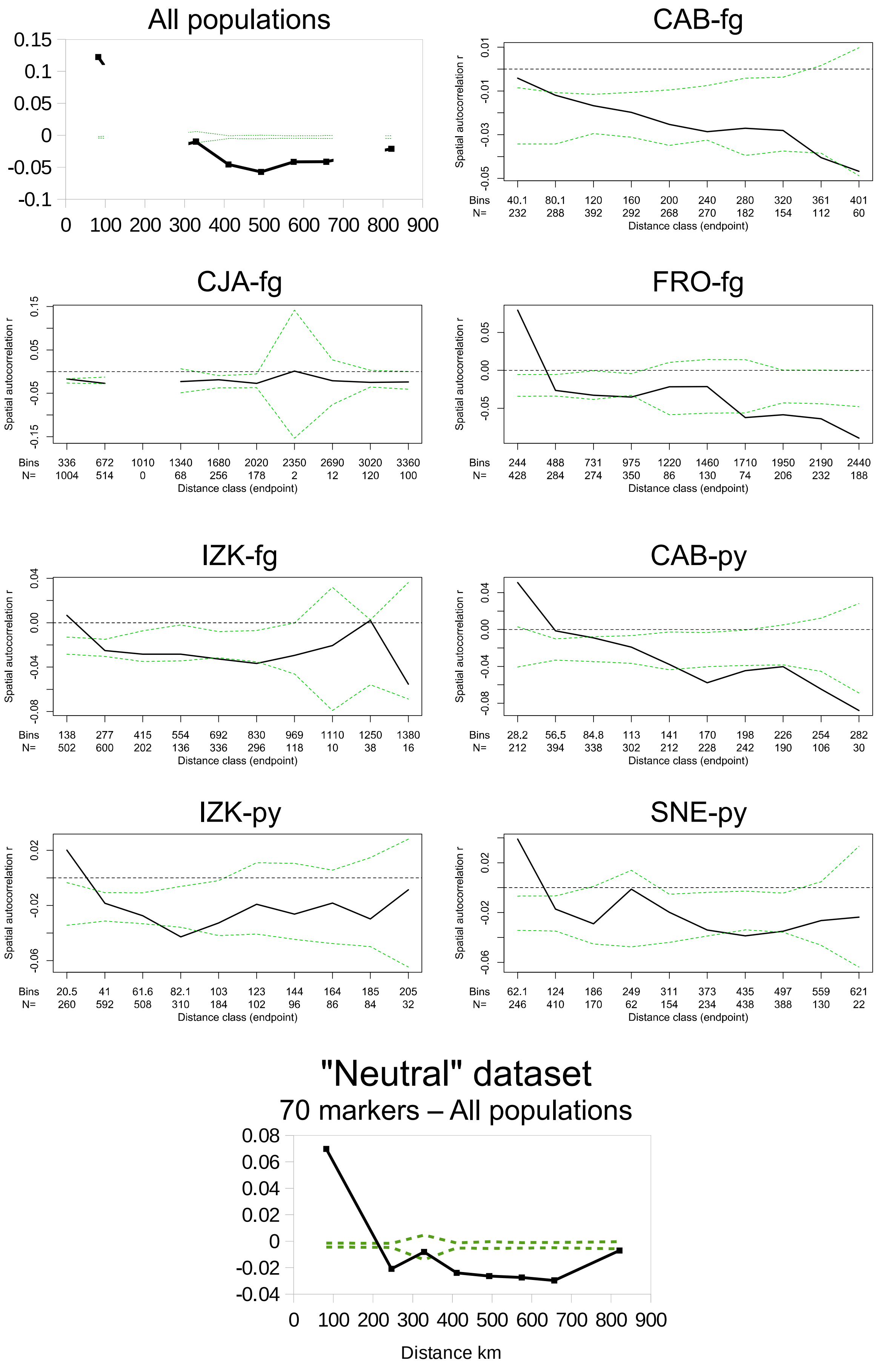

All populations
CAB-fg
CJA-fg
FRO-fg
IZK-fg
CAB-py
IZK-py
SNE-py
Spatial autocorrelation r
Distance class (endpoint)
Bins
N=
"Neutral" dataset
70 markers – All populations
Distance km

**Figure S2-3:**

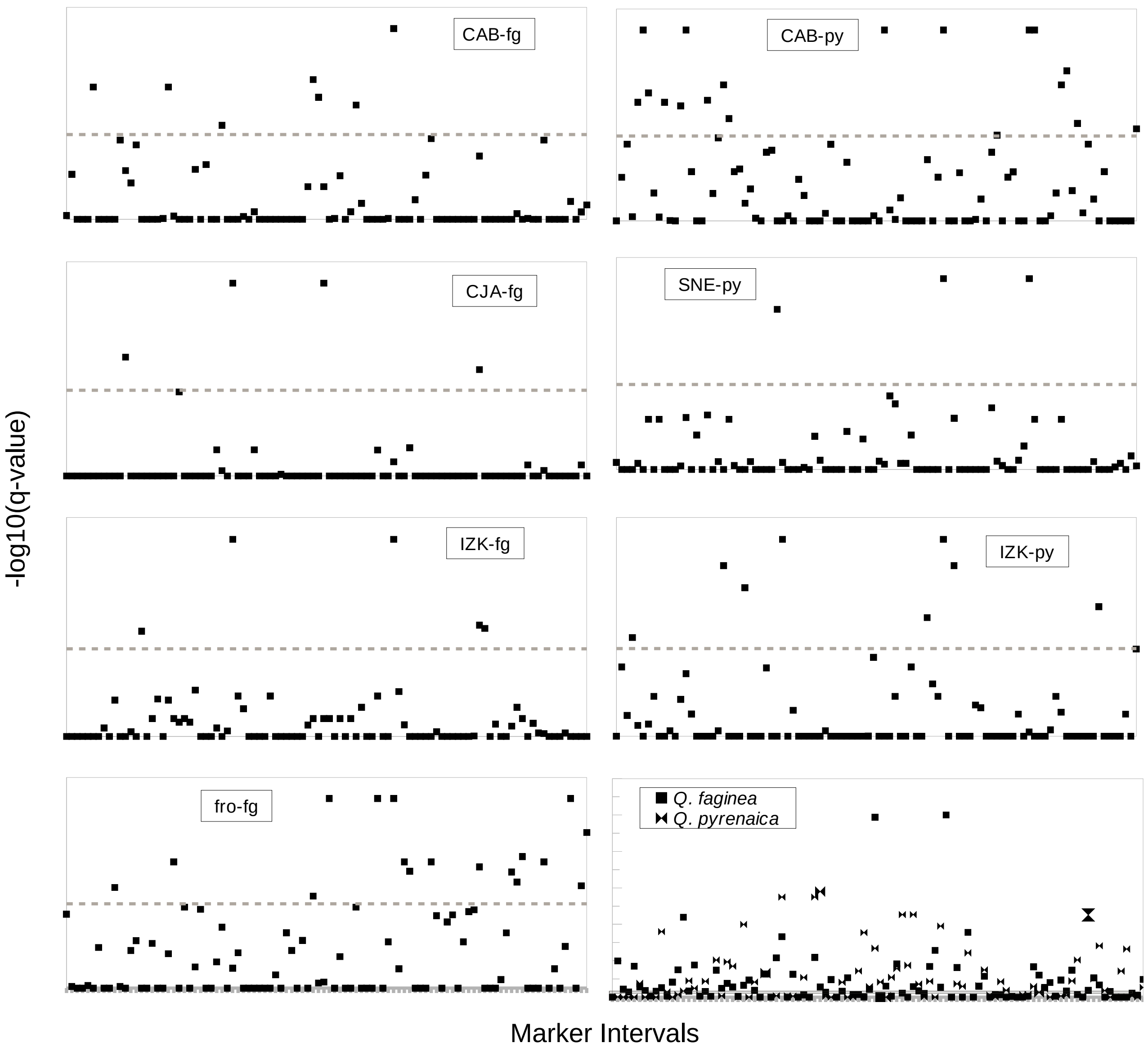
CAB-fg
CAB-py
CJA-fg
SNE-py
IZK-fg
IZK-py
fro-fg
Q. faginea
Q. pyrenaica
-log10(q-value)
Marker Intervals

**Supplemental FILE 3:** POPULATION STRUCTURE

**Figure S3-1:** Geographic (between inter-specific pairs) and phylogenetic (within species) population structure signals obtained with Structure (Fig. S3-1a) and Tess3r (Fig. S3-1b).

**Figure S3-2:** Model comparisons for the sub-sampled datasets: Structure's Linkage model with correlated allele frequencies and equal rates of population differentiation (Fig. S3-2a), the same model with different rates of population differentiation (Fig. S3-2b) and the linkage model with independent allele frequencies (Fig. S3-2c). Tess3r was resilient to subsampling (Fig. S3-2d).

**Figure S3-3:** Effects of the 'neutral' dataset on population structure inferences with Structure' Admixture and Linkage models, and with Tess3r (Fig. S3-3a). Results of the independent allele frequencies model on the geographic pairs and populations within-species are also shown (Fig. S3-3b).

**Figure S3-1a**

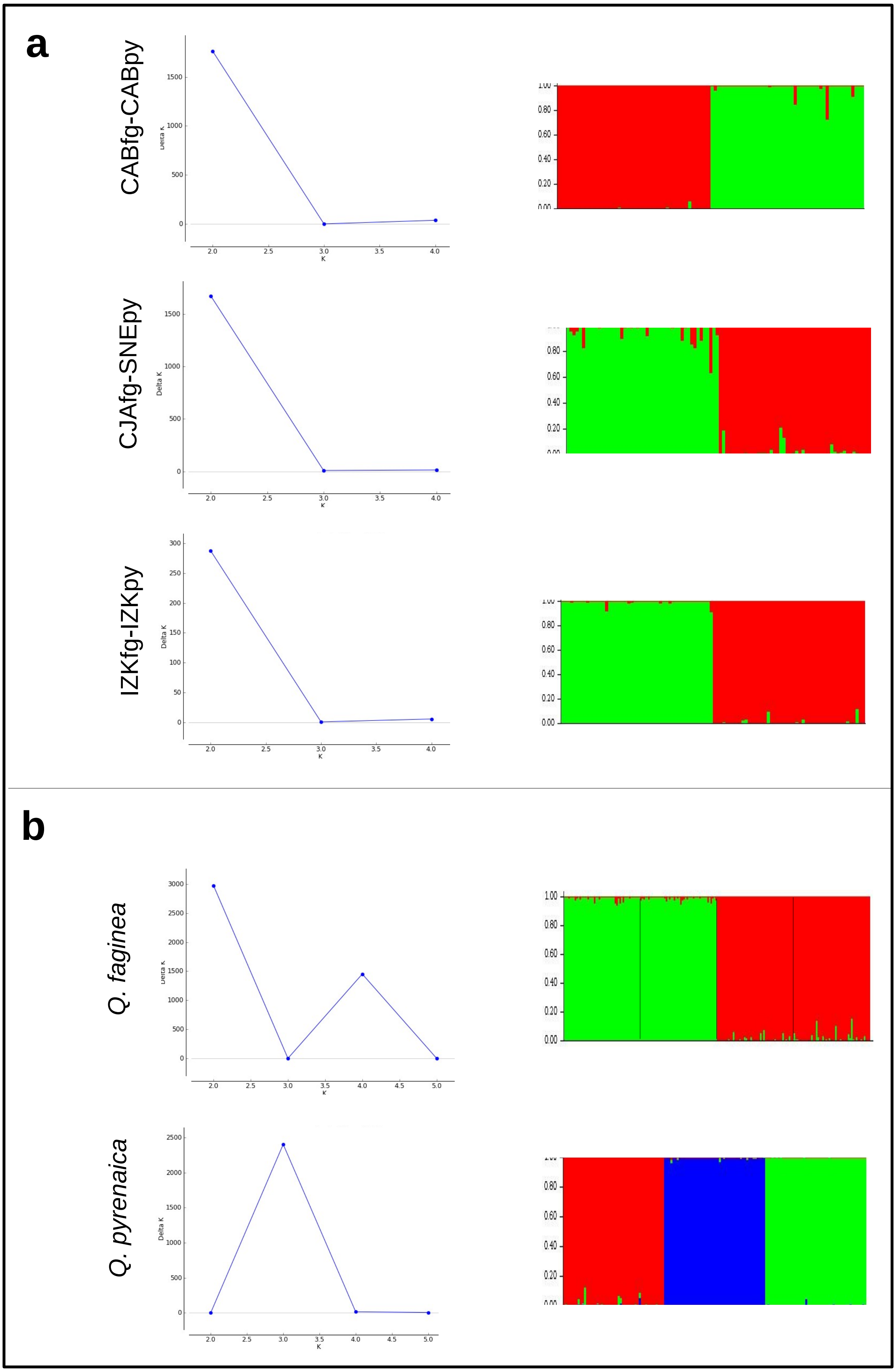

a
CABfg-CABpy
CJAfg-SNEpy
IZKfg-IZKpy
b
Q. faginea
Q. pyrenaica
Delta K
K

**Figure S3-1b**

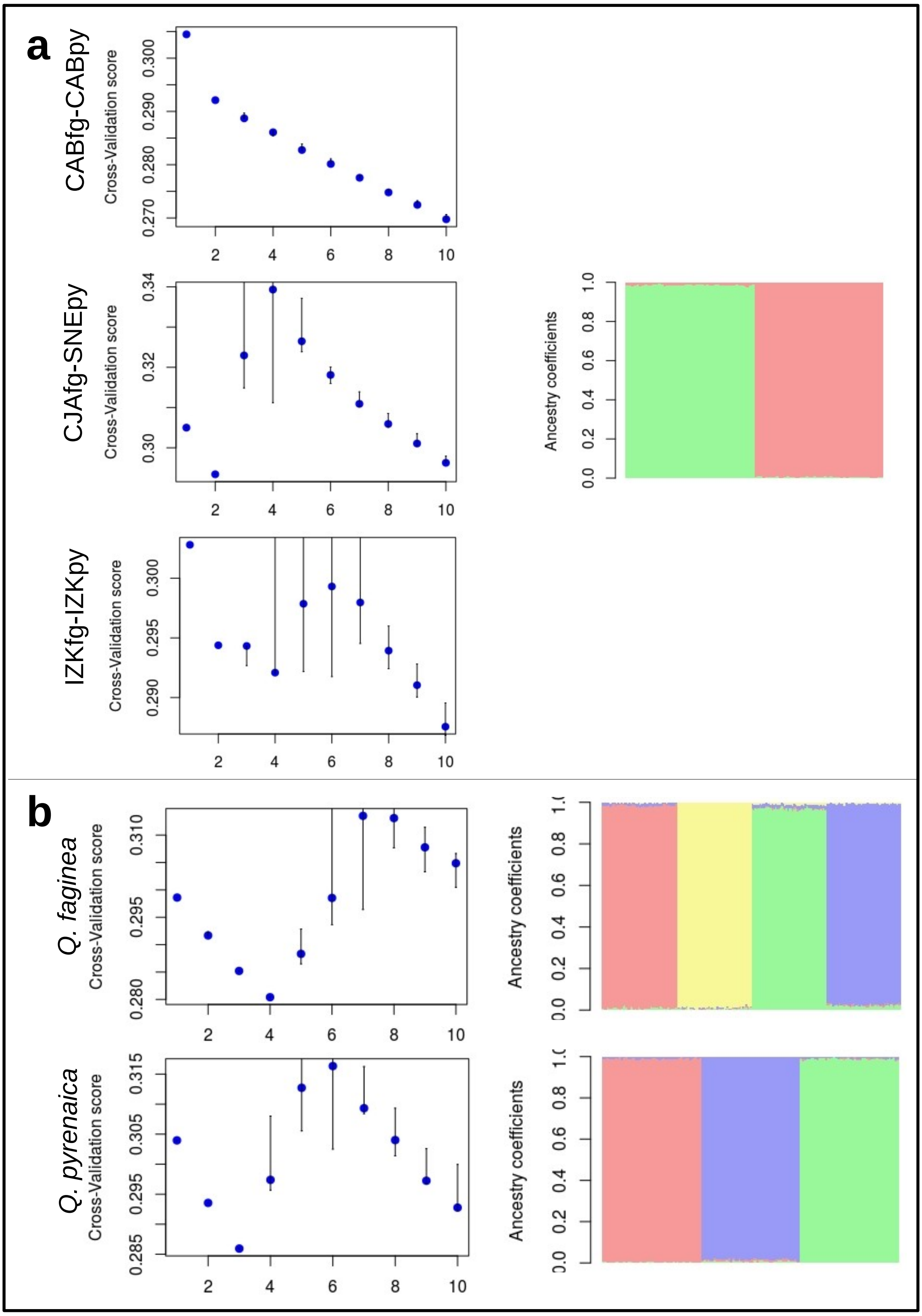
a
CABfg-CABpy
CJAfg-SNEpy
IZKfg-IZKpy
b
Q. faginea
Q. pyrenaica
Cross-Validation score
Ancestry coefficients

### Figure S3-2a

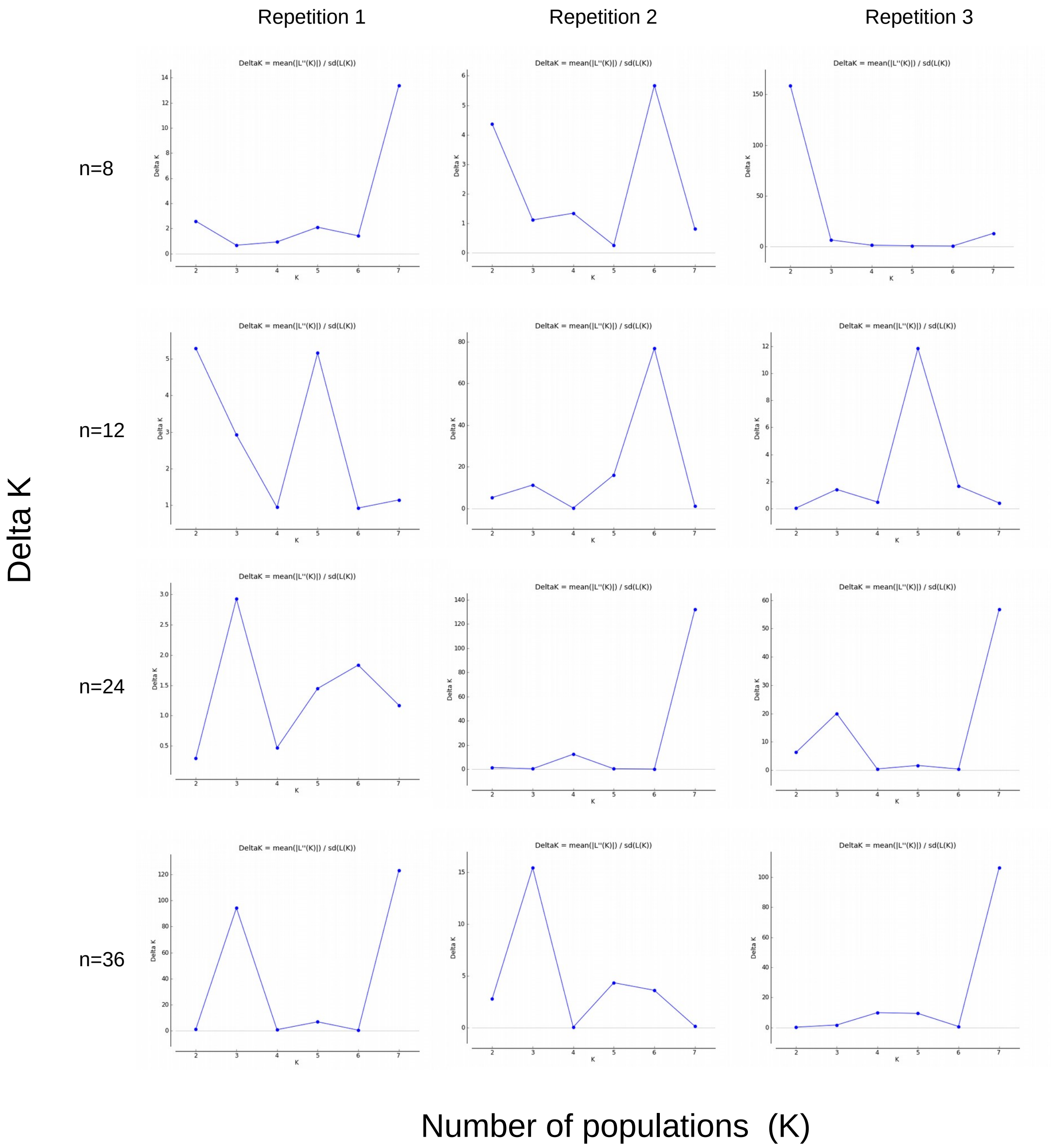
Repetition 1
Repetition 2
Repetition 3
n=8
n=12
n=24
n=36
Delta K
DeltaK = mean(|L''(K)|) / sd(L(K))
Number of populations (K)

**Figure S3-2b**

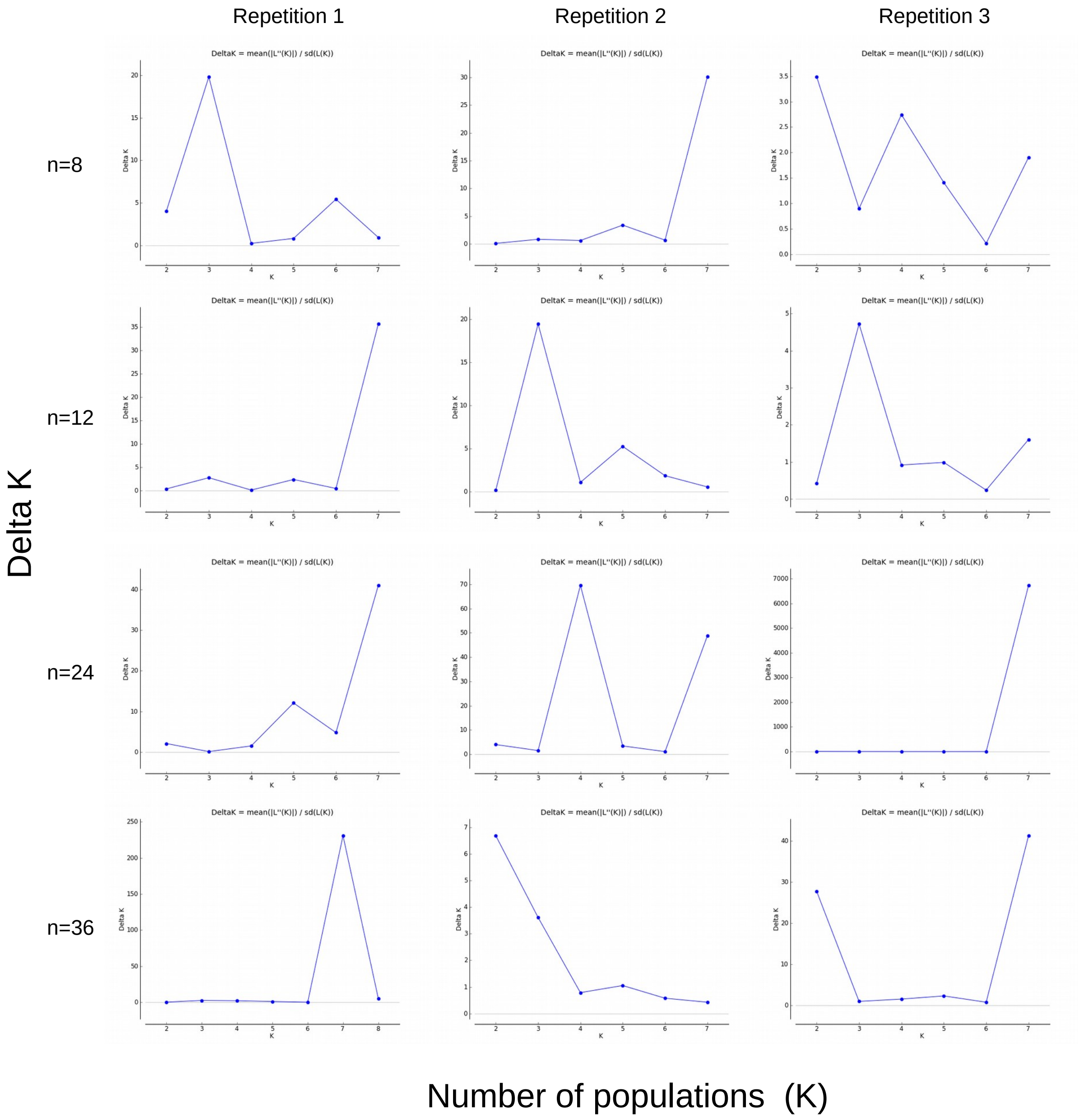
Repetition 1
Repetition 2
Repetition 3
n=8
n=12
n=24
n=36
Delta K
Number of populations (K)
DeltaK = mean(|L''(K)|) / sd(L(K))

**Figure S3-2c**

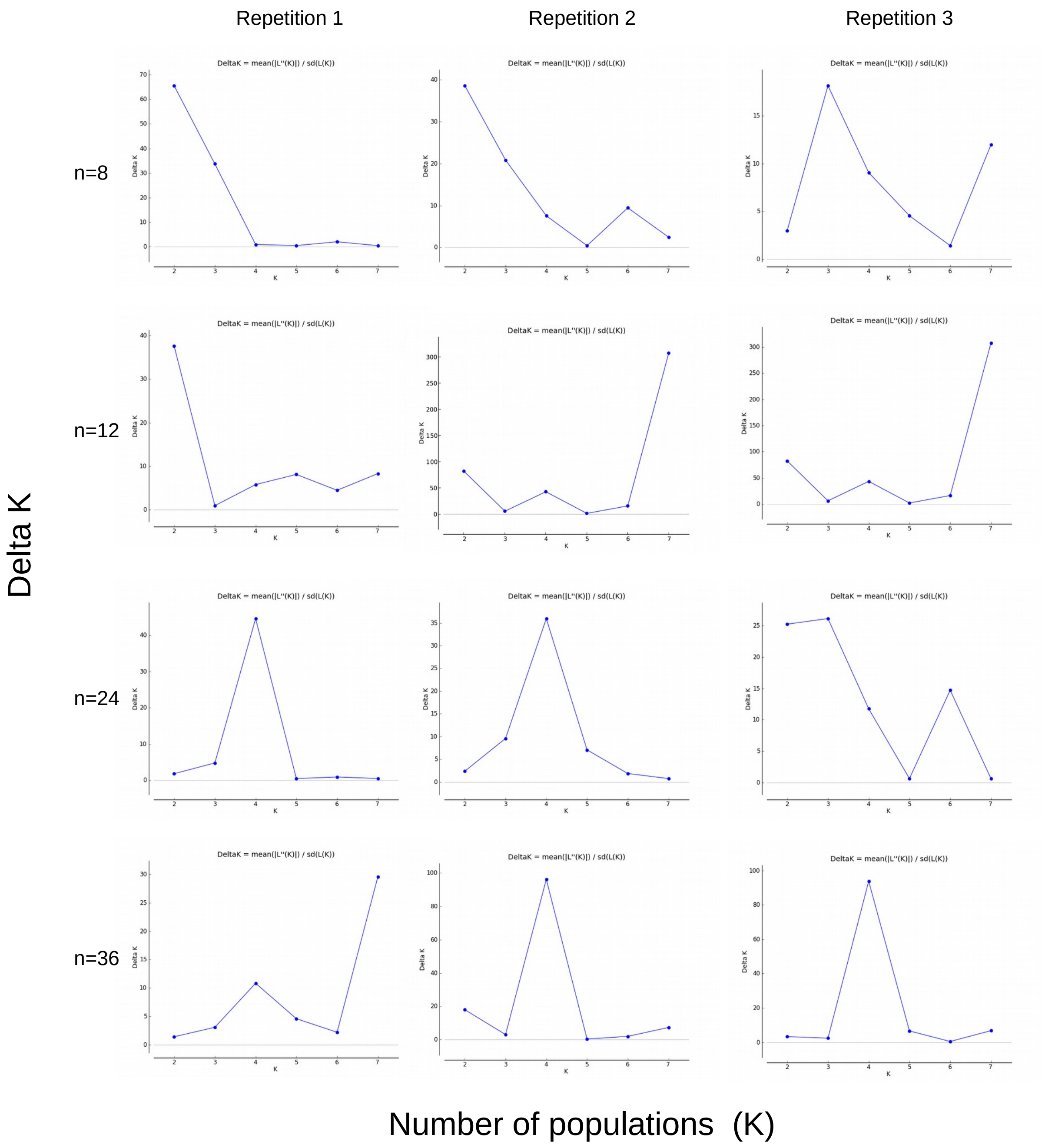
Repetition 1
Repetition 2
Repetition 3
n=8
n=12
n=24
n=36
Delta K
Number of populations (K)
DeltaK = mean(|L''(K)|) / sd(L(K))
Delta K
K

**Figure S3-2d:**

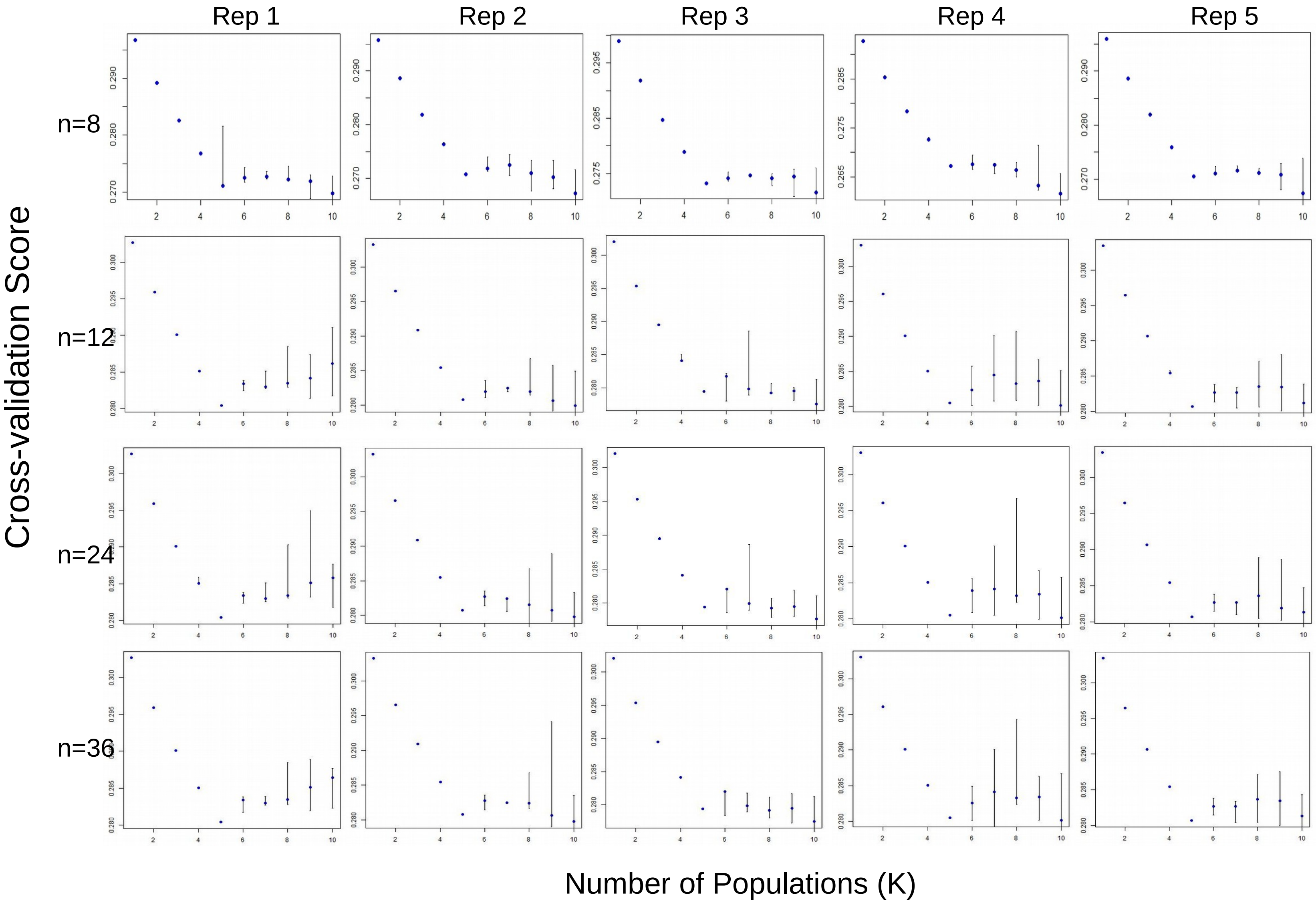
Rep 1
Rep 2
Rep 3
Rep 4
Rep 5
n=8
n=12
n=24
n=36
Cross-validation Score
Number of Populations (K)

**Figure S3-3a:**

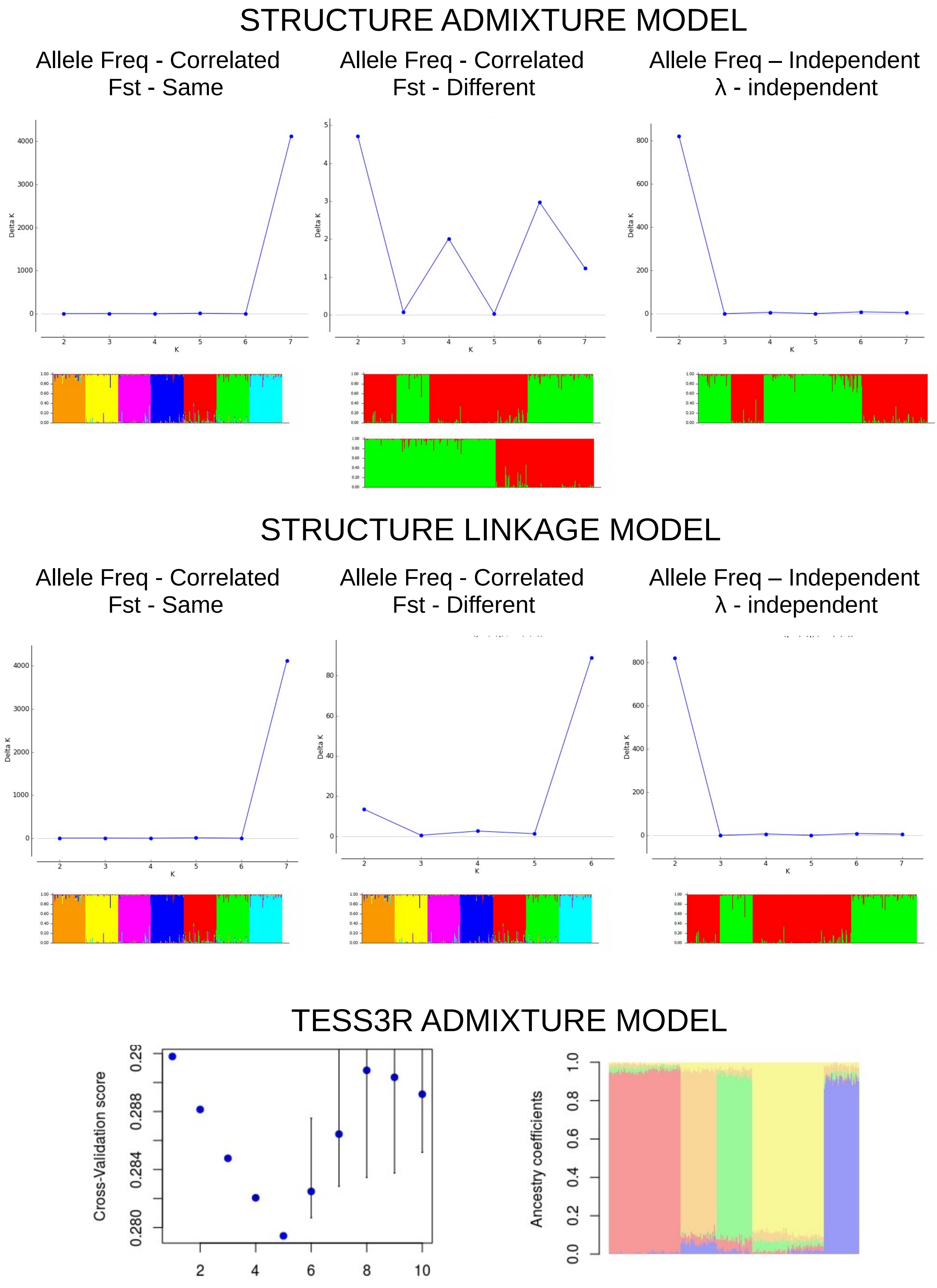
STRUCTURE ADMIXTURE MODEL
Allele Freq - Correlated
Fst - Same
Allele Freq - Correlated
Fst - Different
Allele Freq – Independent
λ - independent
Delta K
K
STRUCTURE LINKAGE MODEL
Allele Freq - Correlated
Fst - Same
Allele Freq - Correlated
Fst - Different
Allele Freq – Independent
λ - independent
Delta K
K
TESS3R ADMIXTURE MODEL
Cross-Validation score
Ancestry coefficients

**Figure S3-3b:**

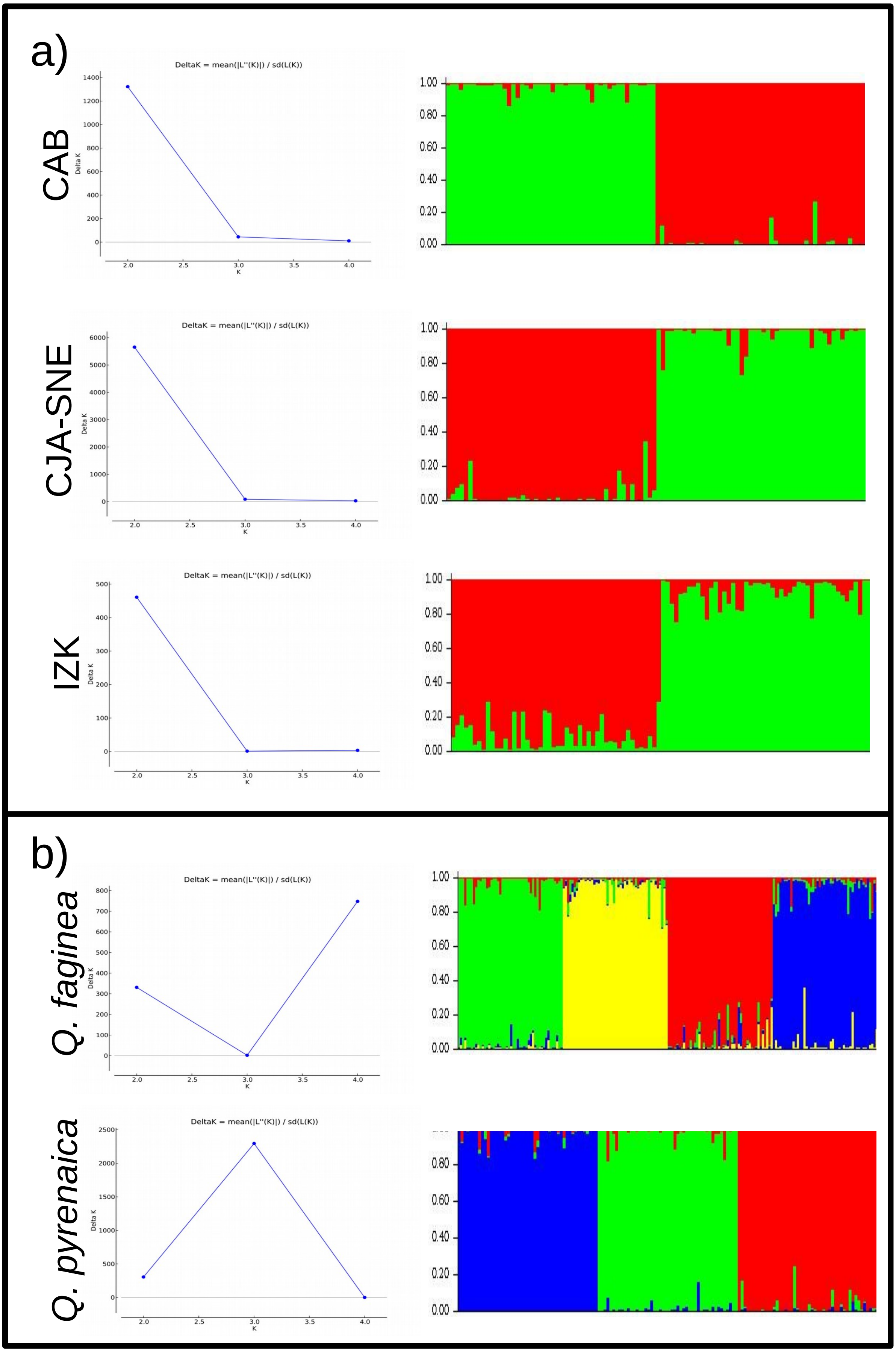
a)
CAB
CJA-SNE
IZK
b)
Q. faginea
Q. pyrenaica
DeltaK = mean(|L''(K)|) / sd(L(K))
Delta K
K

**Supplemental FILE 4:** LOCAL ADAPTATION

**Table S4-1:** List of outlier loci detected by Bayescan in *Quercus faginea* and *Q. pyrenaica*.

**Table S4-2:** List of outlier detected by Tess3r in *Quercus faginea* and *Q. pyrenaica*.

**Table S4-3:** List of genetic-environmental associations detected by Bayescenv in *Quercus faginea* (S4-3a) and *Q. pyrenaica* (S4-3b) for each of the 27 Bioclim variables.

**Table S4-4:** List of genetic-environmental associations detected by Lfmm in *Quercus faginea* (S4-4a) and *Q. pyrenaica* (S4-4b) for each of the 27 Bioclim variables.

**Table S4-5:** List of genetic-environmental associations detected by LfmmAll in *Quercus faginea* (S4-5a) and *Q. pyrenaica* (S4-5b) for each of the 27 Bioclim variables.

**Table S4-6:**List of genetic-environmental associations detected by LfmmAll in *Quercus faginea* (S4-6a) and *Q. pyrenaica* (S4-6b) for each of the 10 'uncorrelated' Bioclim variables.

**Table S4-7:** Comparison of outliers/associated loci among the genome scan (Bayescan, Tess3r) and the genetic-environmental association models (Bayescenv, Lfmm. LfmmAll) for *Quercus faginea* and *Q. pyrenaica*. Results of the LfmmAll model are shown independently for all 27 and for the 10 'uncorrelated' Bioclim variables only to highlight the 'private' loci in the model with 10 variables.

**Figure S4-1:** Graphical output from Bayescan tests among populations from the two oak species. Dot color indicate the absence/presence of null alleles (black: absence of null alleles, dark-gray: one population with null alleles, light-gray: two or more populations with null alleles).

**Figure S4-2:** Visual selection of *K* values (number of ancestral populations) and false discovery rates (FDR) to be used in genomes scans performed with Tess3r in *Q. faginea* (S4-2a) and *Q. pyrenaica* (S4-2b). Histograms show adjusted *p*-values frequencies and Manhattan plots show outliers (red dots) at different FDRs.

**Figure S4-3:** Visual selection of K and FDR values in genomic-environmental association tests with Lfmm for *Q. faginea* (S4-3a) and *Q. pyrenaica* (S4-3b).

**Figure S4-4:** Visual selection of K in GEA tests performed fitting all 27 environmental variables at once (LfmmAll) for *Q. faginea* (S4-4a) and *Q. pyrenaica* (S4-4b).

**Figure S4-5:** Visual selection of K in GEA tests performed fitting the 10 ‘uncorrelated’ Bioclim variables at once (LfmmAll) for *Q. faginea* (S4-4a) and *Q. pyrenaica* (S4-4b)

**Figure S4-6:** Between-species comparison of genetic-environmental associations (loci) detected by Bayescan and Lfmm for each ‘uncorrelated’ Bioclim variable (S4-6a), and between-models comparison of GEAs (loci) detected in *Quercus faginea* and *Q. pyrenaica* for each uncorrelated Bioclim variable (S4-6b).

### Table S4-1:

| *Quercus faginea* | | | | | | *Quercus pyrenaica* | | | | | |
|---|---|---|---|---|---|---|---|---|---|---|---|
| Loci† | prob | log10(PO) | qval | alpha | fst | Loci† | prob | log10(PO) | qval | alpha | fst |
| 1 | 0.949 | 1.3 | 0.007 | -1.241 | 0.028 | 2 | 1.000 | 3.4 | 0.000 | 1.396 | 0.272 |
| 2 | 1.000 | 1000.0 | 0.000 | 1.999 | 0.389 | 5 | 1.000 | 1000.0 | 0.000 | 1.526 | 0.194 |
| 5 | 1.000 | 4.4 | 0.000 | 1.041 | 0.142 | 7 | 1.000 | 1000.0 | 0.000 | 2.231 | 0.327 |
| 7 | 1.000 | 1000.0 | 0.000 | 1.703 | 0.243 | 10 | 1.000 | 1000.0 | 0.000 | 2.660 | 0.558 |
| 24 | 1.000 | 3.7 | 0.000 | -1.240 | 0.026 | 33 | 0.979 | 1.7 | 0.006 | 1.289 | 0.255 |
| 33 | 0.999 | 3.1 | 0.000 | 1.488 | 0.284 | 37 | 0.887 | 0.9 | 0.024 | 0.920 | 0.122 |
| 41 | 1.000 | 4.4 | 0.000 | 1.404 | 0.193 | 41 | 1.000 | 1000.0 | 0.000 | 1.757 | 0.236 |
| 43 | 0.980 | 1.7 | 0.003 | -0.892 | 0.024 | 46 | 0.955 | 1.3 | 0.016 | -1.120 | 0.032 |
| 50 | 1.000 | 1000.0 | 0.000 | 2.059 | 0.406 | 51 | 0.998 | 2.8 | 0.000 | 1.247 | 0.243 |
| 53 | 1.000 | 1000.0 | 0.000 | 1.432 | 0.268 | 52 | 0.966 | 1.5 | 0.010 | -1.347 | 0.026 |
| 54 | 0.724 | 0.4 | 0.042 | -0.587 | 0.050 | 53 | 0.979 | 1.7 | 0.004 | 1.023 | 0.205 |
| 62 | 1.000 | 1000.0 | 0.000 | 2.090 | 0.411 | 62 | 1.000 | 1000.0 | 0.000 | 2.723 | 0.571 |
| 63 | 1.000 | 1000.0 | 0.000 | -2.014 | 0.013 | 63 | 0.817 | 0.6 | 0.032 | -0.895 | 0.041 |
| 65 | 1.000 | 1000.0 | 0.000 | 2.069 | 0.405 | 74 | 1.000 | 1000.0 | 0.000 | 1.223 | 0.151 |
| 73 | 0.929 | 1.1 | 0.012 | -1.211 | 0.029 | 78 | 1.000 | 1000.0 | 0.000 | 1.005 | 0.125 |
| 77 | 0.899 | 0.9 | 0.013 | -0.769 | 0.028 | 86 | 0.999 | 3.2 | 0.000 | 1.164 | 0.146 |
| 78 | 0.999 | 3.3 | 0.000 | 0.827 | 0.118 | 87 | 1.000 | 1000.0 | 0.000 | 1.686 | 0.331 |
| 80 | 1.000 | 1000.0 | 0.000 | 1.662 | 0.234 | 90 | 0.814 | 0.6 | 0.042 | 0.903 | 0.124 |
| 83 | 1.000 | 1000.0 | 0.000 | 1.731 | 0.333 | | | | | | |
| 85 | 1.000 | 1000.0 | 0.000 | -1.408 | 0.015 | | | | | | |
| 87 | 1.000 | 1000.0 | 0.000 | 1.696 | 0.323 | | | | | | |
| 90 | 1.000 | 1000.0 | 0.000 | 1.269 | 0.172 | | | | | | |
| 102 | 0.970 | 1.5 | 0.003 | -1.166 | 0.029 | | | | | | |
| 104 | 0.996 | 2.4 | 0.001 | -0.834 | 0.025 | | | | | | |

†: loci are alphabetically sorted

**Table S4-2:**

| *Q. faginea* | | | *Q. pyrenaica* | | |
| --- | --- | --- | --- | --- | --- |
| Loci | Alleles | adj. p-values | Loci | Alleles | adj. p-values |
| FIR013 | FIR013_157 | 1.8E-04 | FIR106 | FIR106_236 | 6.5E-04 |
| | FIR013_163 | 9.3E-05 | | FIR106_242 | 1.0E-02 |
| FIR030 | FIR030_137 | 4.2E-05 | PIE053 | PIE053_184 | 4.7E-02 |
| FIR044 | FIR044_122 | 2.8E-05 | PIE075 | PIE075_251 | 1.7E-01 |
| FIR048 | FIR048_212 | 1.0E-05 | PIE082 | PIE082_234 | 5.4E-03 |
| FIR104 | FIR104_236 | 1.4E-04 | | PIE082_240 | 3.4E-02 |
| | FIR104_243 | 1.4E-05 | PIE155 | PIE155_256 | 2.0E-01 |
| FIR106 | FIR106_237 | 4.1E-16 | PIE196 | PIE196_223 | 2.0E-02 |
| | FIR106_243 | 1.7E-07 | PIE204 | PIE204_111 | 7.1E-03 |
| | FIR106_242 | 7.3E-07 | | PIE204_119 | 3.3E-02 |
| PIE076 | PIE076_173 | 3.3E-14 | PIE219 | PIE219_124 | 1.1E-02 |
| | PIE076_188 | 4.8E-06 | | PIE219_129 | 2.4E-01 |
| PIE082 | PIE082_234 | 1.5E-08 | | PIE219_130 | 4.9E-02 |
| | PIE082_240 | 3.9E-08 | | PIE219_132 | 1.5E-02 |
| PIE089 | PIE089_261 | 2.0E-04 | PIE228 | PIE228_205 | 3.9E-02 |
| PIE202 | PIE202_124 | 4.5E-19 | POR024 | POR024_198 | 4.3E-02 |
| | PIE202_122 | 3.5E-13 | POR025 | POR025_114 | 6.9E-02 |
| | PIE202_126 | 4.9E-05 | POR030 | POR030_121 | 3.0E-02 |
| PIE204 | PIE204_111 | 1.3E-05 | POR038 | POR038_137 | 5.3E-03 |
| | PIE204_119 | 7.4E-05 | POR039 | POR039_174 | 1.1E-02 |
| PIE219 | PIE219_129 | 2.0E-07 | VIT007 | VIT007_160 | 5.7E-02 |
| PIE242 | PIE242_124 | 2.7E-07 | VIT023 | VIT023_132 | 1.1E-02 |
| POR014 | POR014_186 | 4.3E-05 | | VIT023_138 | 4.8E-02 |
| POR025 | POR025_114 | 3.9E-10 | VIT028 | VIT028_167 | 3.8E-02 |
| | POR025_118 | 3.5E-07 | VIT107 | VIT107_169 | 5.0E-04 |
| POR030 | POR030_118 | 1.9E-04 | WAG020 | WAG020_292 | 6.5E-04 |
| POR039 | POR039_222 | 3.0E-05 | WAG043 | WAG043_307 | 4.0E-03 |
| POR040 | POR040_189 | 7.4E-05 | WAG079 | WAG079_323 | 4.8E-02 |
| VIT010 | VIT010_152 | 5.8E-05 | ZQR30 | ZQR30_220 | 2.7E-03 |
| | VIT010_145 | 2.0E-05 | ZQR87 | ZQR87_126 | 1.8E-02 |
| WAG020 | WAG020_292 | 3.5E-12 | | | |
| | WAG020_298 | 1.3E-04 | | | |
| ZQP15 | ZQP15_126 | 6.2E-05 | | | |
| | ZQP15_128 | 6.2E-05 | | | |

**Table S4-3a:**

| Variable | Marker | PEP | q-value | g |
|---|---|---|---|---|
| BIO1 | FIR013 | 0.011 | 0.009 | 2.762 |
| | FIR044 | 0.026 | 0.015 | 3.693 |
| | PIE219 | 0.176 | 0.046 | 1.541 |
| | POR008 | 0.008 | 0.008 | 2.829 |
| | VIT023 | 0.016 | 0.012 | 2.548 |
| BIO2 | FIR013 | 0.010 | 0.010 | 2.617 |
| | PIE219 | 0.011 | 0.010 | 1.624 |
| BIO3 | FIR044 | 0.001 | 0.001 | 4.919 |
| | PIE082 | 0.054 | 0.028 | 2.059 |
| | GOT021 | 0.026 | 0.021 | 4.456 |
| | PIE137 | 0.017 | 0.017 | 1.817 |
| | VIT028 | 0.057 | 0.033 | 1.432 |
| BIO04 | VIT007 | 0.004 | 0.004 | 1.184 |
| | ZQR87 | 0.050 | 0.027 | 1.287 |
| | FIR065 | 0.007 | 0.007 | 2.548 |
| | PIE219 | 0.038 | 0.022 | 1.514 |
| | VIT018 | 0.073 | 0.039 | 2.568 |
| BIO05 | FIR033 | 0.037 | 0.024 | 2.182 |
| | VIT007 | 0.012 | 0.012 | 1.195 |
| | ZQR87 | 0.053 | 0.034 | 1.380 |
| | FIR013 | 0.003 | 0.003 | 2.687 |
| | FIR044 | 0.074 | 0.035 | 3.323 |
| | PIE202 | 0.021 | 0.015 | 3.272 |
| | PIE219 | 0.020 | 0.011 | 1.792 |
| | POR016 | 0.067 | 0.027 | 2.443 |
| | VIT018 | 0.023 | 0.017 | 2.800 |
| BIO6 | NA | NA | NA | NA |
| BIO07 | VIT007 | 0.003 | 0.003 | 1.163 |
| | ZQR87 | 0.083 | 0.043 | 1.163 |
| | FIR013 | 0.027 | 0.027 | 2.402 |
| | PIE219 | 0.032 | 0.029 | 1.467 |
| BIO8 | FIR044 | 0.053 | 0.028 | 5.123 |
| | FIR106 | 0.004 | 0.004 | 3.765 |
| BIO09 | VIT007 | 0.026 | 0.026 | 1.134 |
| | FIR013 | 0.004 | 0.004 | 2.672 |
| | PIE219 | 0.074 | 0.043 | 1.561 |
| | VIT018 | 0.050 | 0.027 | 2.565 |
| BIO10 | VIT007 | 0.018 | 0.018 | 1.179 |
| | ZQR87 | 0.062 | 0.040 | 1.358 |
| | FIR013 | 0.003 | 0.003 | 2.721 |
| | FIR044 | 0.066 | 0.028 | 3.349 |
| | PIE202 | 0.024 | 0.015 | 3.300 |
| | PIE219 | 0.030 | 0.018 | 1.781 |
| | POR008 | 0.074 | 0.036 | 2.457 |
| | POR016 | 0.099 | 0.045 | 2.408 |
| | VIT018 | 0.017 | 0.010 | 2.809 |
| BIO11 | POR008 | 0.010 | 0.010 | 2.675 |
| | VIT023 | 0.039 | 0.025 | 2.367 |
| BIO12 | PIE219 | 0.020 | 0.020 | 1.578 |
| BIO13 | VIT023 | 0.034 | 0.034 | 2.335 |
| BIO14 | FIR033 | 0.010 | 0.010 | 2.515 |
| | VIT031.1 | 0.083 | 0.040 | 1.555 |
| | FIR013 | 0.009 | 0.009 | 2.621 |
| | FIR044 | 0.016 | 0.013 | 3.755 |
| | PIE082 | 0.090 | 0.030 | 2.081 |
| | PIE202 | 0.015 | 0.012 | 3.290 |
| | PIE219 | 0.022 | 0.017 | 1.829 |
| | POR016 | 0.021 | 0.015 | 2.578 |
| | VIT018 | 0.037 | 0.020 | 2.678 |
| BIO15 | FIR033 | 0.009 | 0.009 | 2.575 |
| | VIT007 | 0.022 | 0.015 | 1.192 |
| | VIT031.1 | 0.070 | 0.033 | 1.576 |
| | ZQR87 | 0.089 | 0.047 | 1.338 |
| | FIR013 | 0.012 | 0.012 | 2.574 |
| | PIE082 | 0.098 | 0.037 | 2.004 |
| | PIE202 | 0.018 | 0.015 | 3.229 |
| | PIE219 | 0.022 | 0.017 | 1.792 |
| | POR016 | 0.027 | 0.020 | 2.498 |
| | VIT018 | 0.052 | 0.027 | 2.606 |
| BIO16 | FIR013 | 0.114 | 0.048 | 2.283 |
| | FIR106 | 0.011 | 0.007 | 2.974 |
| | POR016 | 0.028 | 0.017 | 2.826 |
| | PIE219 | 0.100 | 0.037 | 1.574 |
| | PIE202 | 0.054 | 0.024 | 3.184 |
| | FIR044 | 0.003 | 0.003 | 4.220 |
| BIO17 | FIR030 | 0.092 | 0.049 | 1.280 |
| | FIR033 | 0.008 | 0.008 | 2.620 |
| | VIT007 | 0.054 | 0.035 | 1.137 |
| | VIT031.1 | 0.042 | 0.025 | 1.640 |
| | FIR013 | 0.024 | 0.016 | 2.547 |
| | FIR044 | 0.009 | 0.009 | 3.885 |
| | FIR106 | 0.157 | 0.045 | 2.120 |
| | PIE082 | 0.053 | 0.029 | 2.193 |
| | PIE202 | 0.015 | 0.012 | 3.275 |
| | PIE219 | 0.031 | 0.019 | 1.811 |
| | POR016 | 0.016 | 0.013 | 2.584 |
| | VIT018 | 0.053 | 0.024 | 2.555 |
| BIO18 | FIR033 | 0.007 | 0.007 | 2.632 |
| | VIT007 | 0.046 | 0.032 | 1.146 |
| | VIT031.1 | 0.042 | 0.025 | 1.637 |
| | FIR013 | 0.023 | 0.017 | 2.541 |
| | FIR044 | 0.012 | 0.012 | 3.870 |
| | PIE082 | 0.059 | 0.026 | 2.156 |
| | PIE202 | 0.017 | 0.014 | 3.247 |
| | VIT018 | 0.066 | 0.032 | 2.514 |
| BIO19 | FIR106 | 0.010 | 0.010 | 5.287 |
| | POR008 | 0.029 | 0.019 | 3.143 |
| BIO20 | FIR106 | 0.007 | 0.007 | 3.608 |
| BIO21 | FIR044 | 0.001 | 0.001 | 5.830 |
| | PIE082 | 0.085 | 0.030 | 2.443 |
| BIO22 | FIR106 | 0.007 | 0.007 | 3.499 |
| BIO23 | FIR044 | 0.061 | 0.033 | 4.843 |
| | FIR106 | 0.005 | 0.005 | 3.682 |
| BIO24 | GOT021 | 0.017 | 0.017 | 5.079 |
| | FIR044 | 0.005 | 0.005 | 5.550 |
| | POR008 | 0.035 | 0.020 | 3.033 |
| BIO25 | FIR044 | 0.041 | 0.022 | 5.018 |
| | FIR106 | 0.004 | 0.004 | 3.714 |
| BIO26 | GOT021 | 0.033 | 0.025 | 4.140 |
| | FIR044 | 0.015 | 0.015 | 4.807 |
| BIO27 | FIR106 | 0.006 | 0.006 | 3.533 |

**Table S4-3b:**

| Variable | Marker | PEP | q-value | g | Variable | Marker | PEP | q-value | g |
|---|---|---|---|---|---|---|---|---|---|
| BIO1 | FIR044 | 0.029 | 0.015 | 3.917 | BIO13 | FIR106 | 0.007 | 0.007 | 5.240 |
| | PIE202 | 0.044 | 0.025 | 6.602 | BIO14 | VIT022 | 0.032 | 0.032 | 1.669 |
| | PIE264 | 0.001 | 0.001 | 5.777 | | FIR106 | 0.001 | 0.001 | 5.500 |
| BIO2 | FIR044 | 0.122 | 0.048 | 3.050 | | POR008 | 0.008 | 0.005 | 5.089 |
| | FIR106 | 0.002 | 0.002 | 5.468 | BIO15 | FIR047 | 0.883 | 0.422 | 0.362 |
| | POR008 | 0.020 | 0.011 | 4.942 | | FIR106 | 0.002 | 0.002 | 5.463 |
| BIO3 | FIR044 | 0.023 | 0.011 | 3.974 | | POR008 | 0.023 | 0.012 | 4.935 |
| | FIR106 | 0.132 | 0.049 | 3.696 | BIO16 | VIT022 | 0.015 | 0.015 | 1.759 |
| | PIE202 | 0.042 | 0.022 | 6.463 | | FIR106 | 0.002 | 0.002 | 5.536 |
| | POR008 | 0.000 | 0.000 | 5.903 | | POR008 | 0.005 | 0.004 | 5.183 |
| | FIR033 | 0.049 | 0.049 | 2.695 | BIO17 | FIR044 | 0.114 | 0.047 | 3.087 |
| BIO04 | VIT022 | 0.007 | 0.007 | 1.817 | | FIR106 | 0.002 | 0.002 | 5.464 |
| | FIR106 | 0.010 | 0.009 | 5.387 | | POR008 | 0.026 | 0.014 | 4.900 |
| | POR008 | 0.009 | 0.009 | 5.270 | BIO18 | PIE137 | 0.047 | 0.047 | 2.343 |
| BIO05 | VIT022 | 0.025 | 0.025 | 1.724 | | FIR106 | 0.003 | 0.003 | 5.437 |
| | FIR106 | 0.052 | 0.028 | 4.459 | | POR008 | 0.067 | 0.035 | 4.622 |
| | POR008 | 0.004 | 0.004 | 5.592 | BIO19 | PIE137 | 0.017 | 0.017 | 2.381 |
| BIO6 | FIR044 | 0.001 | 0.001 | 4.524 | BIO20 | FIR044 | 0.023 | 0.023 | 3.678 |
| | FIR106 | 0.119 | 0.044 | 3.755 | | VIT023 | 0.070 | 0.039 | 1.960 |
| | PIE202 | 0.051 | 0.019 | 6.265 | BIO21 | PIE137 | 0.005 | 0.005 | 2.536 |
| | POR008 | 0.005 | 0.003 | 5.290 | | POR038 | 0.062 | 0.034 | 1.351 |
| | FIR033 | 0.088 | 0.038 | 2.807 | | FIR106 | 0.007 | 0.007 | 3.968 |
| | POR038 | 0.008 | 0.008 | 1.571 | | VIT023 | 0.059 | 0.029 | 2.049 |
| BIO07 | VIT022 | 0.021 | 0.021 | 1.725 | BIO22 | PIE137 | 0.008 | 0.008 | 2.482 |
| BIO07 | FIR106 | 0.001 | 0.001 | 5.498 | | FIR044 | 0.032 | 0.031 | 3.596 |
| BIO07 | POR008 | 0.007 | 0.004 | 5.126 | | FIR106 | 0.029 | 0.029 | 4.599 |
| BIO8 | NA | NA | NA | NA | | VIT023 | 0.074 | 0.045 | 1.933 |
| BIO09 | FIR033 | 0.041 | 0.041 | 2.761 | BIO23 | PIE137 | 0.006 | 0.006 | 2.506 |
| | PIE023 | 0.050 | 0.045 | 3.775 | | FIR044 | 0.018 | 0.018 | 3.753 |
| | FIR044 | 0.093 | 0.047 | 3.440 | | FIR106 | 0.023 | 0.020 | 4.668 |
| | FIR106 | 0.018 | 0.009 | 4.817 | | VIT023 | 0.067 | 0.036 | 1.976 |
| | POR008 | 0.000 | 0.000 | 5.920 | BIO24 | VIT022 | 0.007 | 0.007 | 1.816 |
| | POR016 | 0.077 | 0.032 | 2.511 | | FIR106 | 0.011 | 0.011 | 5.320 |
| BIO10 | FIR033 | 0.040 | 0.040 | 2.764 | | POR008 | 0.011 | 0.011 | 5.262 |
| | PIE023 | 0.047 | 0.043 | 3.784 | BIO25 | PIE137 | 0.007 | 0.007 | 2.497 |
| | FIR106 | 0.018 | 0.009 | 4.785 | | FIR106 | 0.025 | 0.023 | 4.647 |
| | FIR044 | 0.096 | 0.049 | 3.418 | | VIT023 | 0.070 | 0.039 | 1.955 |
| | POR008 | 0.000 | 0.000 | 5.911 | BIO26 | NA | NA | NA | NA |
| | POR016 | 0.082 | 0.034 | 2.481 | BIO27 | PIE137 | 0.007 | 0.007 | 2.485 |
| BIO11 | NA | NA | NA | NA | | FIR044 | 0.032 | 0.032 | 3.598 |
| | | | | | | FIR106 | 0.031 | 0.031 | 4.580 |
| | | | | | | VIT023 | 0.074 | 0.046 | 1.938 |

**Table S4-4a:**

| Variable | Loci | Alleles | adj. p-values | Variable | Loci | Alleles | adj. p-values |
|---|---|---|---|---|---|---|---|
| BIO01 | PIE075 | PIE075_251 | 1.7E-12 | BIO11 | PIE075 | PIE075_251 | 1.6E-12 |
| BIO01 | FIR013 | FIR013_166 | 1.8E-11 | BIO11 | FIR013 | FIR013_166 | 3.3E-10 |
| BIO01 | POR008 | POR008_139 | 1.2E-07 | BIO11 | POR008 | POR008_139 | 1.2E-08 |
| BIO02 | FIR044 | FIR044_124 | 6.1E-10 | BIO12 | PIE219 | PIE219_130 | 3.5E-09 |
| BIO02 | ZQR87_ | ZQR87_128 | 3.8E-08 | BIO12 | FIR044 | FIR044_124 | 7.3E-09 |
| BIO02 | FIR013 | FIR013_166 | 5.2E-07 | BIO12 | FIR044 | FIR044_134 | 1.3E-08 |
| BIO03 | FIR013 | FIR013_166 | 2.4E-10 | BIO13 | POR008 | POR008_148 | 1.4E-15 |
| BIO03 | PIE075 | PIE075_251 | 2.7E-09 | BIO13 | FIR044 | FIR044_122 | 1.2E-09 |
| BIO03 | FIR106 | FIR106_242 | 6.1E-09 | BIO13 | PIE219 | PIE219_129 | 1.5E-09 |
| BIO03 | FIR013 | FIR013_163 | 9.4E-09 | BIO13 | POR008 | POR008_139 | 4.0E-09 |
| BIO03 | VIT025 | VIT025_162 | 2.7E-08 | BIO13 | FIR044 | FIR044_134 | 6.2E-08 |
| BIO03 | POR016 | POR016_134 | 2.1E-07 | BIO13 | VIT010 | VIT010_152 | 2.9E-07 |
| BIO03 | POR016 | POR016_144 | 4.0E-07 | BIO13 | PIE202 | PIE202_128 | 3.0E-07 |
| BIO04 | FIR013 | FIR013_166 | 1.1E-11 | BIO14 | FIR013 | FIR013_166 | 1.1E-20 |
| BIO04 | FIR044 | FIR044_124 | 1.5E-10 | BIO14 | FIR106 | FIR106_242 | 1.7E-07 |
| BIO04 | PIE075 | PIE075_249 | 1.6E-08 | BIO15 | FIR013 | FIR013_166 | 1.5E-12 |
| BIO04 | PIE219 | PIE219_124 | 3.6E-08 | BIO16 | PIE219 | PIE219_130 | 1.4E-15 |
| BIO04 | ZQR087 | ZQR87_128 | 4.0E-08 | BIO16 | PIE075 | PIE075_250 | 1.3E-09 |
| BIO04 | PIE075 | PIE075_251 | 2.6E-07 | BIO16 | FIR044 | FIR044_134 | 8.1E-08 |
| BIO05 | FIR013 | FIR013_166 | 5.6E-15 | BIO17 | FIR013 | FIR013_166 | 1.7E-20 |
| BIO06 | FIR044 | FIR044_134 | 3.1E-07 | BIO17 | FIR106 | FIR106_242 | 4.2E-08 |
| BIO07 | FIR044 | FIR044_124 | 2.9E-20 | BIO18 | FIR013 | FIR013_166 | 3.7E-19 |
| BIO07 | PIE075 | PIE075_249 | 1.8E-12 | BIO18 | FIR106 | FIR106_242 | 1.5E-07 |
| BIO07 | PIE219 | PIE219_124 | 8.4E-12 | BIO19 | FIR013 | FIR013_166 | 5.3E-16 |
| BIO07 | FIR013 | FIR013_166 | 1.2E-09 | BIO19 | PIE075 | PIE075_251 | 1.5E-10 |
| BIO07 | ZQR087 | ZQR87_128 | 3.4E-09 | BIO19 | FIR106 | FIR106_242 | 1.3E-09 |
| BIO08 | NA | NA | NA | BIO20 | FIR013 | FIR013_166 | 1.3E-15 |
| BIO09 | PIE075 | PIE075_251 | 7.3E-09 | BIO20 | PIE075 | PIE075_251 | 4.4E-10 |
| BIO09 | FIR013 | FIR013_166 | 8.3E-08 | BIO20 | FIR106 | FIR106_242 | 2.6E-09 |
| BIO10 | FIR013 | FIR013_166 | 4.5E-17 | BIO21 | NA | NA | NA |
| BIO10 | PIE075 | PIE075_251 | 1.2E-08 | BIO22 | NA | NA | NA |
| BIO10 | FIR106 | FIR106_242 | 3.5E-07 | BIO23 | NA | NA | NA |
| | | | | BIO24 | NA | NA | NA |
| | | | | BIO25 | NA | NA | NA |
| | | | | BIO26 | NA | NA | NA |
| | | | | BIO27 | NA | NA | NA |

**Table S4-4b:**

| Variable | Loci | Alleles | adj. p-values |
|---|---|---|---|
| BIO01 | PIE202 | PIE202_126 | 5.8E-16 |
| BIO01 | POR008 | POR008_139 | 7.4E-14 |
| BIO01 | POR008 | POR008_148 | 9.6E-14 |
| BIO01 | FIR044 | FIR044_122 | 2.4E-13 |
| BIO01 | PIE202 | PIE202_128 | 1.0E-10 |
| BIO01 | PIE075 | PIE075_251 | 5.3E-10 |
| BIO01 | PIE053 | PIE053_184 | 1.1E-07 |
| BIO02 | FIR044 | FIR044_124 | 8.4E-30 |
| BIO02 | PIE219 | PIE219_124 | 3.3E-14 |
| BIO02 | VIT022 | VIT022_174 | 1.3E-09 |
| BIO02 | PIE137 | PIE137_350 | 4.9E-08 |
| BIO03 | POR008 | POR008_139 | 2.9E-21 |
| BIO03 | FIR044 | FIR044_122 | 1.4E-20 |
| BIO03 | POR008 | POR008_148 | 5.1E-20 |
| BIO03 | PIE202 | PIE202_126 | 1.7E-18 |
| BIO03 | PIE202 | PIE202_128 | 1.4E-15 |
| BIO03 | PIE075 | PIE075_251 | 9.4E-12 |
| BIO03 | PIE053 | PIE053_184 | 1.6E-09 |
| BIO03 | FIR106 | FIR106_236 | 5.1E-07 |
| BIO03 | PIE204 | PIE204_111 | 8.6E-07 |
| BIO04 | FIR044 | FIR044_124 | 3.3E-18 |
| BIO04 | PIE219 | PIE219_124 | 8.5E-09 |
| BIO04 | VIT022 | VIT022_174 | 2.9E-07 |
| BIO05 | FIR044 | FIR044_122 | 3.4E-08 |
| BIO05 | POR008 | POR008_139 | 4.3E-08 |
| BIO05 | PIE202 | PIE202_128 | 4.7E-07 |
| BIO06 | FIR044 | FIR044_134 | 1.7E-28 |
| BIO06 | FIR030 | FIR030_137 | 5.9E-10 |
| BIO06 | PIE219 | PIE219_130 | 1.1E-09 |
| BIO06 | VIT023 | VIT023_138 | 1.9E-08 |
| BIO06 | VIT023 | VIT023_141 | 2.0E-08 |
| BIO06 | POR016 | POR016_134 | 5.6E-08 |
| BIO06 | PIE137 | PIE137_352 | 3.2E-07 |
| BIO07 | FIR044 | FIR044_124 | 8.9E-30 |
| BIO07 | PIE219 | PIE219_124 | 1.7E-12 |
| BIO07 | VIT022 | VIT022_174 | 4.0E-09 |
| BIO08 | PIE202 | PIE202_126 | 5.1E-08 |
| BIO09 | POR008 | POR008_139 | 1.8E-23 |
| BIO09 | FIR044 | FIR044_122 | 4.9E-23 |
| BIO09 | POR008 | POR008_148 | 1.8E-17 |
| BIO09 | PIE202 | PIE202_128 | 1.9E-17 |
| BIO09 | PIE075 | PIE075_251 | 9.9E-12 |
| BIO09 | PIE053 | PIE053_184 | 4.0E-09 |
| BIO09 | PIE202 | PIE202_126 | 6.5E-09 |
| BIO09 | FIR106 | FIR106_236 | 1.7E-06 |
| BIO10 | POR008 | POR008_139 | 3.0E-22 |
| BIO10 | FIR044 | FIR044_122 | 4.3E-22 |
| BIO10 | PIE202 | PIE202_128 | 2.2E-16 |
| BIO10 | POR008 | POR008_148 | 2.3E-16 |
| BIO10 | PIE075 | PIE075_251 | 3.8E-11 |
| BIO10 | PIE053 | PIE053_184 | 1.9E-08 |
| BIO10 | PIE202 | PIE202_126 | 1.9E-08 |
| BIO11 | NA | NA | NA |
| BIO12 | FIR044 | FIR044_124 | 1.4E-24 |
| BIO12 | PIE219 | PIE219_124 | 1.9E-11 |
| BIO12 | VIT022 | VIT022_174 | 6.9E-11 |
| BIO13 | FIR044 | FIR044_124 | 1.0E-19 |
| BIO13 | PIE219 | PIE219_124 | 4.2E-15 |
| BIO13 | PIE137 | PIE137_350 | 4.8E-11 |
| BIO13 | WAG079 | WAG079_321 | 5.1E-08 |
| BIO13 | PIE075 | PIE075_250 | 8.8E-08 |
| BIO13 | VIT007 | VIT007_134 | 2.6E-07 |
| BIO14 | FIR044 | FIR044_124 | 1.5E-30 |
| BIO14 | PIE219 | PIE219_124 | 1.1E-13 |
| BIO14 | PIE137 | PIE137_350 | 7.7E-08 |
| BIO15 | FIR044 | FIR044_124 | 4.5E-22 |
| BIO15 | PIE219 | PIE219_124 | 9.7E-14 |
| BIO15 | VIT022 | VIT022_174 | 8.7E-10 |
| BIO15 | PIE137 | PIE137_350 | 3.2E-08 |
| BIO16 | FIR044 | FIR044_124 | 1.0E-24 |
| BIO16 | PIE219 | PIE219_124 | 1.1E-10 |
| BIO17 | FIR044 | FIR044_124 | 2.5E-22 |
| BIO17 | PIE219 | PIE219_124 | 1.3E-14 |
| BIO17 | VIT022 | VIT022_174 | 6.4E-10 |
| BIO17 | PIE137 | PIE137_350 | 6.7E-09 |
| BIO18 | FIR044 | FIR044_124 | 1.1E-24 |
| BIO18 | PIE219 | PIE219_124 | 1.6E-15 |
| BIO18 | PIE137 | PIE137_350 | 3.5E-10 |
| BIO19 | FIR044 | FIR044_134 | 1.5E-16 |
| BIO19 | PIE219 | PIE219_124 | 6.3E-15 |
| BIO19 | PIE137 | PIE137_350 | 2.3E-11 |
| BIO19 | VIT007 | VIT007_134 | 1.7E-10 |
| BIO19 | PIE219 | PIE219_130 | 4.1E-10 |
| BIO19 | VIT023 | VIT023_141 | 1.6E-09 |
| BIO19 | FIR030 | FIR030_137 | 2.8E-09 |
| BIO19 | VIT022 | VIT022_174 | 4.3E-09 |
| BIO19 | PIE137 | PIE137_352 | 4.5E-09 |
| BIO19 | VIT023 | VIT023_138 | 1.4E-08 |
| BIO19 | PIE075 | PIE075_250 | 3.1E-08 |
| BIO19 | PIE082 | PIE082_240 | 1.4E-07 |
| BIO19 | WAG079 | WAG079_323 | 2.0E-07 |
| BIO19 | WAG079 | WAG079_321 | 2.3E-07 |
| BIO20 | FIR044 | FIR044_134 | 1.6E-27 |
| BIO20 | PIE137 | PIE137_352 | 2.6E-12 |
| BIO20 | PIE219 | PIE219_130 | 1.1E-11 |
| BIO20 | FIR044 | FIR044_124 | 1.3E-11 |
| BIO20 | VIT023 | VIT023_138 | 3.5E-09 |
| BIO20 | FIR030 | FIR030_137 | 3.7E-09 |
| BIO20 | PIE075 | PIE075_250 | 3.7E-09 |
| BIO20 | PIE219 | PIE219_124 | 7.1E-09 |
| BIO20 | VIT023 | VIT023_141 | 1.0E-08 |
| BIO20 | PIE137 | PIE137_350 | 6.0E-08 |
| BIO20 | VIT007 | VIT007_134 | 6.6E-08 |
| BIO20 | FIR030 | FIR030_134 | 4.1E-07 |
| BIO20 | PIE082 | PIE082_237 | 1.8E-06 |
| BIO21 | FIR044 | FIR044_134 | 2.3E-15 |
| BIO21 | PIE075 | PIE075_250 | 3.3E-13 |
| BIO21 | PIE137 | PIE137_352 | 2.4E-12 |
| BIO21 | VIT023 | VIT023_141 | 5.4E-12 |
| BIO21 | PIE219 | PIE219_130 | 1.0E-10 |
| BIO21 | FIR030 | FIR030_137 | 1.9E-10 |
| BIO21 | VIT023 | VIT023_138 | 1.2E-09 |
| BIO21 | PIE082 | PIE082_240 | 2.6E-09 |
| BIO21 | POR016 | POR016_134 | 1.8E-08 |
| BIO21 | VIT007 | VIT007_134 | 2.2E-08 |
| BIO21 | WAG079 | WAG079_323 | 1.9E-07 |
| BIO21 | FIR030 | FIR030_134 | 2.3E-07 |
| BIO21 | POR038 | POR038_189 | 2.5E-06 |
| BIO22 | PIE075 | PIE075_250 | 1.6E-14 |
| BIO22 | FIR044 | FIR044_134 | 2.5E-10 |
| BIO22 | VIT007 | VIT007_134 | 6.2E-10 |
| BIO22 | PIE137 | PIE137_352 | 6.5E-10 |
| BIO22 | VIT023 | VIT023_141 | 3.4E-09 |
| BIO22 | PIE219 | PIE219_130 | 2.2E-08 |
| BIO22 | VIT023 | VIT023_138 | 2.9E-08 |
| BIO22 | FIR030 | FIR030_137 | 3.0E-08 |
| BIO22 | PIE082 | PIE082_240 | 7.2E-08 |
| BIO22 | WAG079 | WAG079_323 | 1.4E-07 |
| BIO22 | FIR030 | FIR030_134 | 5.7E-07 |
| BIO22 | POR016 | POR016_134 | 7.1E-07 |
| BIO23 | PIE075 | PIE075_250 | 1.6E-14 |
| BIO23 | FIR044 | FIR044_134 | 4.6E-12 |
| BIO23 | PIE137 | PIE137_352 | 2.4E-11 |
| BIO23 | VIT023 | VIT023_141 | 3.4E-10 |
| BIO23 | VIT007 | VIT007_134 | 1.0E-09 |
| BIO23 | PIE219 | PIE219_130 | 2.8E-09 |
| BIO23 | FIR030 | FIR030_137 | 4.4E-09 |
| BIO23 | VIT023 | VIT023_138 | 1.5E-08 |
| BIO23 | PIE082 | PIE082_240 | 2.0E-08 |
| BIO23 | WAG079 | WAG079_323 | 6.8E-08 |
| BIO23 | POR016 | POR016_134 | 2.2E-07 |
| BIO23 | FIR030 | FIR030_134 | 7.3E-07 |
| BIO24 | FIR044 | FIR044_124 | 7.6E-13 |
| BIO25 | PIE075 | PIE075_250 | 2.2E-14 |
| BIO25 | FIR044 | FIR044_134 | 8.2E-12 |
| BIO25 | PIE137 | PIE137_352 | 1.3E-10 |
| BIO25 | VIT023 | VIT023_141 | 5.0E-10 |
| BIO25 | VIT007 | VIT007_134 | 8.8E-10 |
| BIO25 | PIE219 | PIE219_130 | 5.2E-09 |
| BIO25 | FIR030 | FIR030_137 | 5.6E-09 |
| BIO25 | VIT023 | VIT023_138 | 1.3E-08 |
| BIO25 | PIE082 | PIE082_240 | 4.1E-08 |
| BIO25 | WAG079 | WAG079_323 | 2.1E-07 |
| BIO25 | POR016 | POR016_134 | 3.3E-07 |
| BIO25 | FIR030 | FIR030_134 | 4.8E-07 |
| BIO26 | PIE202 | PIE202_126 | 5.0E-09 |
| BIO27 | PIE075 | PIE075_250 | 9.1E-14 |
| BIO27 | VIT007 | VIT007_134 | 6.2E-10 |
| BIO27 | FIR044 | FIR044_134 | 6.9E-10 |
| BIO27 | PIE137 | PIE137_352 | 9.9E-10 |
| BIO27 | VIT023 | VIT023_141 | 2.8E-09 |
| BIO27 | PIE219 | PIE219_130 | 3.1E-08 |
| BIO27 | FIR030 | FIR030_137 | 5.1E-08 |
| BIO27 | VIT023 | VIT023_138 | 8.7E-08 |
| BIO27 | PIE082 | PIE082_240 | 1.9E-07 |
| BIO27 | WAG079 | WAG079_323 | 2.4E-07 |
| BIO27 | FIR030 | FIR030_134 | 9.8E-07 |
| BIO27 | POR016 | POR016_134 | 1.8E-06 |

**Table S4-5a:**

| Variable | Loci | Alleles | adj. p-val |
|---|---|---|---|
| BIO01 | VIT026 | VIT026_261 | 3.4E-10 |
| BIO01 | PIE002 | PIE002_179 | 2.0E-07 |
| BIO01 | PIE036 | PIE036_184 | 4.9E-07 |
| BIO01 | PIE076 | PIE076_188 | 7.1E-07 |
| BIO02 | PIE219 | PIE219_130 | 3.7E-17 |
| BIO02 | FIR044 | FIR044_134 | 7.1E-16 |
| BIO02 | PIE075 | PIE075_250 | 3.7E-10 |
| BIO02 | FIR044 | FIR044_124 | 1.1E-07 |
| BIO03 | POR016 | POR016_134 | 1.5E-17 |
| BIO03 | VIT023 | VIT023_138 | 1.1E-15 |
| BIO03 | POR016 | POR016_144 | 5.7E-14 |
| BIO03 | PIE137 | PIE137_350 | 1.0E-11 |
| BIO03 | PIE082 | PIE082_237 | 3.5E-11 |
| BIO03 | VIT028 | VIT028_163 | 1.2E-10 |
| BIO03 | VIT023 | VIT023_141 | 2.0E-10 |
| BIO03 | FIR044 | FIR044_134 | 2.7E-10 |
| BIO03 | PIE082 | PIE082_240 | 3.3E-10 |
| BIO03 | PIE137 | PIE137_352 | 9.1E-10 |
| BIO03 | FIR106 | FIR106_241 | 1.3E-09 |
| BIO03 | POR008 | POR008_148 | 4.5E-09 |
| BIO04 | PIE219 | PIE219_130 | 3.3E-17 |
| BIO04 | FIR044 | FIR044_134 | 5.2E-16 |
| BIO04 | POR016 | POR016_134 | 4.9E-09 |
| BIO04 | POR008 | POR008_148 | 5.3E-08 |
| BIO04 | VIT023 | VIT023_138 | 3.0E-07 |
| BIO04 | PIE075 | PIE075_250 | 3.1E-07 |
| BIO04 | POR016 | POR016_144 | 3.3E-07 |
| BIO04 | POR024 | POR024_181 | 1.0E-06 |
| BIO05 | POR016 | POR016_144 | 6.0E-16 |
| BIO05 | POR016 | POR016_134 | 2.4E-15 |
| BIO05 | FIR044 | FIR044_134 | 1.3E-14 |
| BIO05 | PIE082 | PIE082_237 | 2.7E-14 |
| BIO05 | VIT023 | VIT023_138 | 3.4E-13 |
| BIO05 | PIE219 | PIE219_130 | 1.0E-12 |
| BIO05 | VIT023 | VIT023_141 | 5.9E-11 |
| BIO05 | POR008 | POR008_148 | 1.2E-10 |
| BIO05 | PIE082 | PIE082_240 | 1.1E-09 |
| BIO05 | FIR106 | FIR106_241 | 2.0E-09 |
| BIO05 | PIE137 | PIE137_350 | 1.2E-08 |
| BIO05 | PIE137 | PIE137_352 | 5.5E-08 |
| BIO05 | VIT028 | VIT028_163 | 8.1E-08 |
| BIO06 | FIR044 | FIR044_134 | 6.1E-23 |
| BIO06 | PIE219 | PIE219_130 | 2.8E-16 |
| BIO06 | PIE075 | PIE075_250 | 2.5E-09 |
| BIO06 | FIR044 | FIR044_124 | 5.2E-09 |
| BIO06 | PIE075 | PIE075_249 | 3.6E-07 |
| BIO07 | FIR044 | FIR044_134 | 1.2E-18 |
| BIO07 | PIE219 | PIE219_130 | 1.8E-16 |
| BIO07 | POR016 | POR016_144 | 8.9E-15 |
| BIO07 | POR016 | POR016_134 | 4.6E-13 |
| BIO07 | VIT023 | VIT023_138 | 1.9E-11 |
| BIO07 | POR008 | POR008_148 | 4.8E-11 |
| BIO07 | FIR106 | FIR106_241 | 1.3E-10 |
| BIO07 | PIE082 | PIE082_237 | 1.6E-10 |
| BIO07 | VIT023 | VIT023_141 | 4.3E-10 |
| BIO07 | PIE137 | PIE137_352 | 9.1E-10 |
| BIO07 | VIT028 | VIT028_163 | 6.3E-09 |
| BIO07 | PIE219 | PIE219_129 | 3.4E-07 |
| BIO07 | PIE082 | PIE082_240 | 3.6E-07 |
| BIO08 | POR016 | POR016_134 | 3.3E-10 |
| BIO08 | VIT023 | VIT023_138 | 1.2E-07 |
| BIO09 | POR016 | POR016_134 | 2.6E-09 |
| BIO10 | FIR044 | FIR044_134 | 6.2E-10 |
| BIO10 | POR016 | POR016_144 | 1.2E-09 |
| BIO10 | PIE082 | PIE082_237 | 1.7E-09 |
| BIO10 | POR016 | POR016_134 | 1.6E-08 |
| BIO11 | FIR044 | FIR044_134 | 2.4E-10 |
| BIO11 | PIE219 | PIE219_130 | 4.0E-08 |
| BIO12 | FIR044 | FIR044_134 | 2.4E-12 |
| BIO12 | PIE082 | PIE082_240 | 5.9E-11 |
| BIO12 | POR016 | POR016_144 | 9.4E-11 |
| BIO12 | VIT023 | VIT023_138 | 9.4E-11 |
| BIO12 | VIT023 | VIT023_141 | 1.7E-10 |
| BIO12 | POR016 | POR016_134 | 5.1E-10 |
| BIO12 | POR008 | POR008_148 | 2.1E-09 |
| BIO12 | FIR106 | FIR106_241 | 4.2E-09 |
| BIO12 | PIE137 | PIE137_352 | 4.3E-09 |
| BIO12 | PIE137 | PIE137_350 | 4.8E-08 |
| BIO12 | PIE082 | PIE082_237 | 1.3E-07 |
| BIO12 | PIE219 | PIE219_130 | 4.0E-07 |
| BIO13 | VIT023 | VIT023_138 | 4.6E-21 |
| BIO13 | POR016 | POR016_134 | 4.6E-20 |
| BIO13 | POR016 | POR016_144 | 2.7E-17 |
| BIO13 | PIE082 | PIE082_237 | 1.8E-16 |
| BIO13 | FIR044 | FIR044_134 | 1.7E-14 |
| BIO13 | PIE082 | PIE082_240 | 2.4E-14 |
| BIO13 | VIT023 | VIT023_141 | 2.7E-14 |
| BIO13 | FIR106 | FIR106_241 | 6.3E-14 |
| BIO13 | PIE137 | PIE137_350 | 1.1E-13 |
| BIO13 | PIE137 | PIE137_352 | 7.2E-13 |
| BIO13 | VIT028 | VIT028_163 | 1.3E-12 |
| BIO13 | POR008 | POR008_148 | 5.7E-12 |
| BIO13 | POR008 | POR008_139 | 5.3E-09 |
| BIO13 | PIE202 | PIE202_128 | 1.7E-08 |
| BIO13 | POR024 | POR024_181 | 9.1E-08 |
| BIO13 | PIE202 | PIE202_124 | 1.3E-07 |
| BIO13 | FIR044 | FIR044_122 | 2.7E-07 |
| BIO13 | PIE219 | PIE219_129 | 3.0E-07 |
| BIO13 | PIE219 | PIE219_130 | 1.5E-06 |
| BIO13 | POR025 | POR025_114 | 2.5E-06 |
| BIO13 | PIE088 | PIE088_199 | 4.0E-06 |
| BIO14 | POR016 | POR016_134 | 1.0E-21 |
| BIO14 | POR016 | POR016_144 | 2.2E-20 |
| BIO14 | VIT023 | VIT023_138 | 1.6E-19 |
| BIO14 | FIR044 | FIR044_134 | 5.8E-19 |
| BIO14 | PIE082 | PIE082_237 | 3.6E-18 |
| BIO14 | PIE082 | PIE082_240 | 9.3E-16 |
| BIO14 | VIT023 | VIT023_141 | 3.6E-15 |
| BIO14 | FIR106 | FIR106_241 | 5.5E-14 |
| BIO14 | VIT028 | VIT028_163 | 6.6E-14 |
| BIO14 | PIE137 | PIE137_350 | 2.0E-13 |
| BIO14 | PIE137 | PIE137_352 | 1.5E-12 |
| BIO14 | POR008 | POR008_148 | 2.0E-12 |
| BIO14 | PIE219 | PIE219_130 | 1.9E-10 |
| BIO14 | POR024 | POR024_181 | 1.4E-07 |
| BIO15 | VIT023 | VIT023_138 | 6.2E-16 |
| BIO15 | FIR044 | FIR044_134 | 5.5E-15 |
| BIO15 | POR016 | POR016_134 | 1.3E-14 |
| BIO15 | POR016 | POR016_144 | 1.1E-13 |
| BIO15 | VIT023 | VIT023_141 | 1.9E-12 |
| BIO15 | PIE137 | PIE137_350 | 2.2E-12 |
| BIO15 | PIE082 | PIE082_240 | 6.0E-12 |
| BIO15 | POR008 | POR008_148 | 1.7E-11 |
| BIO15 | PIE082 | PIE082_237 | 2.1E-10 |
| BIO15 | FIR106 | FIR106_241 | 7.8E-10 |
| BIO15 | PIE137 | PIE137_352 | 1.2E-09 |
| BIO15 | VIT028 | VIT028_163 | 1.3E-08 |
| BIO15 | PIE202 | PIE202_124 | 4.7E-08 |
| BIO15 | PIE219 | PIE219_130 | 5.1E-08 |
| BIO15 | POR008 | POR008_139 | 9.8E-08 |
| BIO15 | PIE219 | PIE219_129 | 9.3E-07 |
| BIO15 | POR024 | POR024_181 | 2.1E-06 |
| BIO15 | FIR044 | FIR044_122 | 2.2E-06 |
| BIO16 | PIE082 | PIE082_237 | 8.5E-14 |
| BIO16 | VIT023 | VIT023_138 | 3.1E-13 |
| BIO16 | PIE082 | PIE082_240 | 1.3E-12 |
| BIO16 | POR008 | POR008_148 | 2.2E-12 |
| BIO16 | VIT028 | VIT028_163 | 5.5E-11 |
| BIO16 | PIE137 | PIE137_352 | 8.1E-10 |
| BIO16 | VIT023 | VIT023_141 | 1.0E-09 |
| BIO16 | FIR106 | FIR106_241 | 4.6E-09 |
| BIO16 | POR024 | POR024_181 | 2.0E-07 |
| BIO16 | PIE137 | PIE137_350 | 2.2E-07 |
| BIO16 | POR008 | POR008_139 | 4.9E-07 |
| BIO17 | POR016 | POR016_134 | 1.2E-18 |
| BIO17 | VIT023 | VIT023_138 | 8.6E-18 |
| BIO17 | POR016 | POR016_144 | 3.5E-16 |
| BIO17 | PIE082 | PIE082_237 | 5.2E-16 |
| BIO17 | VIT023 | VIT023_141 | 3.5E-15 |
| BIO17 | PIE082 | PIE082_240 | 3.8E-14 |
| BIO17 | FIR044 | FIR044_134 | 7.8E-12 |
| BIO17 | PIE137 | PIE137_352 | 2.8E-11 |
| BIO17 | PIE137 | PIE137_350 | 1.6E-10 |
| BIO17 | VIT028 | VIT028_163 | 1.9E-09 |
| BIO17 | POR008 | POR008_148 | 3.1E-09 |
| BIO17 | FIR106 | FIR106_241 | 9.9E-08 |
| BIO17 | PIE202 | PIE202_124 | 1.2E-07 |
| BIO17 | PIE219 | PIE219_130 | 1.9E-07 |
| BIO17 | POR008 | POR008_139 | 8.1E-07 |
| BIO17 | POR024 | POR024_181 | 1.9E-06 |
| BIO17 | PIE219 | PIE219_129 | 2.0E-06 |
| BIO18 | POR016 | POR016_134 | 1.7E-12 |
| BIO18 | PIE082 | PIE082_240 | 1.7E-11 |
| BIO18 | VIT023 | VIT023_138 | 2.7E-11 |
| BIO18 | FIR044 | FIR044_134 | 2.8E-11 |
| BIO18 | VIT023 | VIT023_141 | 5.1E-11 |
| BIO18 | PIE082 | PIE082_237 | 1.1E-10 |
| BIO18 | FIR106 | FIR106_241 | 5.1E-10 |
| BIO18 | POR016 | POR016_144 | 1.3E-09 |
| BIO18 | POR008 | POR008_148 | 1.4E-09 |
| BIO18 | PIE137 | PIE137_350 | 4.2E-09 |
| BIO18 | VIT028 | VIT028_163 | 2.1E-08 |
| BIO18 | PIE137 | PIE137_352 | 3.5E-08 |
| BIO18 | FIR044 | FIR044_122 | 1.0E-06 |
| BIO19 | PIE075 | PIE075_250 | 2.9E-11 |
| BIO19 | FIR044 | FIR044_134 | 6.5E-08 |
| BIO20 | VIT023 | VIT023_138 | 5.0E-12 |
| BIO20 | POR016 | POR016_134 | 1.7E-10 |
| BIO20 | PIE137 | PIE137_350 | 5.6E-08 |
| BIO20 | VIT028 | VIT028_163 | 7.5E-08 |
| BIO20 | VIT023 | VIT023_141 | 3.5E-07 |
| BIO20 | POR016 | POR016_144 | 3.9E-07 |
| BIO20 | PIE082 | PIE082_237 | 1.1E-06 |
| BIO20 | PIE082 | PIE082_240 | 1.4E-06 |
| BIO21 | VIT023 | VIT023_138 | 8.7E-19 |
| BIO21 | POR016 | POR016_134 | 4.2E-16 |
| BIO21 | VIT023 | VIT023_141 | 1.8E-12 |
| BIO21 | POR016 | POR016_144 | 6.6E-12 |
| BIO21 | PIE137 | PIE137_350 | 1.6E-11 |
| BIO21 | PIE082 | PIE082_240 | 6.5E-10 |
| BIO21 | PIE137 | PIE137_352 | 9.6E-08 |
| BIO21 | VIT028 | VIT028_163 | 8.7E-07 |
| BIO22 | POR016 | POR016_134 | 6.5E-11 |
| BIO22 | VIT023 | VIT023_138 | 1.0E-09 |
| BIO22 | PIE137 | PIE137_350 | 1.3E-07 |
| BIO22 | POR016 | POR016_144 | 1.5E-07 |
| BIO22 | VIT023 | VIT023_141 | 5.2E-07 |
| BIO23 | POR016 | POR016_134 | 2.1E-11 |
| BIO23 | VIT023 | VIT023_138 | 4.2E-11 |
| BIO23 | POR016 | POR016_144 | 1.1E-08 |
| BIO23 | PIE137 | PIE137_350 | 3.6E-08 |
| BIO23 | VIT028 | VIT028_163 | 8.2E-08 |
| BIO23 | PIE082 | PIE082_237 | 1.0E-07 |
| BIO24 | NA | NA | NA |
| BIO25 | VIT023 | VIT023_138 | 2.9E-12 |
| BIO25 | PIE137 | PIE137_350 | 3.2E-11 |
| BIO25 | VIT023 | VIT023_141 | 5.2E-11 |
| BIO25 | POR016 | POR016_134 | 1.1E-10 |
| BIO25 | FIR106 | FIR106_241 | 5.5E-09 |
| BIO25 | POR016 | POR016_144 | 5.8E-09 |
| BIO25 | PIE082 | PIE082_240 | 2.0E-08 |
| BIO25 | PIE082 | PIE082_237 | 3.0E-07 |
| BIO25 | VIT028 | VIT028_163 | 6.5E-07 |
| BIO26 | POR016 | POR016_134 | 3.3E-15 |
| BIO26 | VIT023 | VIT023_138 | 5.6E-14 |
| BIO26 | POR016 | POR016_144 | 2.5E-12 |
| BIO26 | VIT028 | VIT028_163 | 1.6E-11 |
| BIO26 | PIE137 | PIE137_350 | 2.0E-09 |
| BIO26 | VIT023 | VIT023_141 | 5.0E-09 |
| BIO26 | PIE088 | PIE088_199 | 5.7E-08 |
| BIO26 | FIR106 | FIR106_241 | 1.2E-07 |
| BIO26 | PIE082 | PIE082_240 | 6.3E-07 |
| BIO27 | VIT023 | VIT023_138 | 1.4E-15 |
| BIO27 | PIE137 | PIE137_350 | 1.3E-12 |
| BIO27 | POR016 | POR016_134 | 6.4E-12 |
| BIO27 | POR016 | POR016_144 | 7.4E-11 |
| BIO27 | VIT023 | VIT023_141 | 3.1E-08 |
| BIO27 | PIE137 | PIE137_352 | 8.2E-08 |
| BIO27 | FIR106 | FIR106_241 | 2.9E-07 |

**Table S4-5b:**

| Variable | Loci | Alleles | adj. p-val | Variable | Loci | Alleles | adj. p-val |
|---|---|---|---|---|---|---|---|
| BIO01 | FIR044 | FIR044_122 | 1.4E-12 | BIO10 | FIR044 | FIR044_122 | 1.4E-16 |
| BIO01 | POR008 | POR008_148 | 8.8E-12 | BIO10 | POR008 | POR008_139 | 3.6E-11 |
| BIO01 | FIR044 | FIR044_134 | 4.8E-11 | BIO10 | PIE202 | PIE202_126 | 1.1E-10 |
| BIO01 | POR008 | POR008_139 | 5.3E-11 | BIO10 | PIE202 | PIE202_128 | 1.3E-09 |
| BIO01 | PIE202 | PIE202_128 | 8.8E-11 | BIO10 | PIE204 | PIE204_111 | 8.2E-09 |
| BIO01 | PIE202 | PIE202_126 | 1.4E-10 | BIO10 | POR008 | POR008_148 | 2.0E-08 |
| BIO01 | PIE075 | PIE075_251 | 4.7E-08 | BIO10 | FIR044 | FIR044_134 | 7.0E-08 |
| BIO01 | POR016 | POR016_134 | 2.6E-07 | BIO10 | FIR030 | FIR030_134 | 2.0E-07 |
| BIO02 | PIE071 | PIE071_246 | 1.1E-08 | BIO10 | FIR013 | FIR013_166 | 7.2E-07 |
| BIO02 | VIT020 | VIT020_144 | 4.4E-08 | BIO10 | VIT023 | VIT023_141 | 7.5E-07 |
| BIO02 | POR020 | POR020_130 | 4.5E-08 | BIO11 | PIE202 | PIE202_128 | 4.6E-12 |
| BIO03 | POR008 | POR008_139 | 2.1E-09 | BIO11 | FIR044 | FIR044_122 | 3.3E-11 |
| BIO03 | PIE202 | PIE202_126 | 4.1E-08 | BIO11 | POR008 | POR008_139 | 4.4E-11 |
| BIO03 | PIE202 | PIE202_128 | 1.0E-07 | BIO11 | POR008 | POR008_148 | 1.1E-10 |
| BIO03 | FIR044 | FIR044_122 | 5.6E-07 | BIO11 | PIE202 | PIE202_126 | 3.9E-09 |
| BIO03 | POR008 | POR008_148 | 8.1E-07 | BIO11 | PIE075 | PIE075_251 | 1.5E-08 |
| BIO04 | POR008 | POR008_139 | 2.1E-09 | BIO11 | FIR044 | FIR044_134 | 2.0E-08 |
| BIO04 | PIE202 | PIE202_126 | 4.1E-08 | BIO11 | POR016 | POR016_134 | 2.1E-07 |
| BIO04 | PIE202 | PIE202_128 | 1.0E-07 | BIO11 | PIE082 | PIE082_237 | 1.3E-06 |
| BIO04 | FIR044 | FIR044_122 | 5.5E-07 | BIO12 | POR008 | POR008_139 | 1.4E-12 |
| BIO04 | POR008 | POR008_148 | 8.1E-07 | BIO12 | FIR044 | FIR044_134 | 7.0E-12 |
| BIO05 | POR008 | POR008_139 | 1.0E-08 | BIO12 | PIE202 | PIE202_128 | 1.1E-11 |
| BIO05 | FIR044 | FIR044_122 | 1.4E-07 | BIO12 | FIR044 | FIR044_122 | 5.4E-11 |
| BIO05 | PIE075 | PIE075_251 | 1.7E-07 | BIO12 | POR008 | POR008_148 | 6.2E-11 |
| BIO06 | FIR044 | FIR044_122 | 3.5E-11 | BIO12 | PIE202 | PIE202_126 | 2.5E-10 |
| BIO06 | POR008 | POR008_139 | 2.5E-10 | BIO12 | POR016 | POR016_134 | 2.0E-07 |
| BIO06 | POR008 | POR008_148 | 2.4E-09 | BIO12 | FIR013 | FIR013_166 | 8.5E-07 |
| BIO06 | PIE202 | PIE202_128 | 3.6E-09 | BIO13 | FIR044 | FIR044_124 | 1.2E-07 |
| BIO06 | PIE202 | PIE202_126 | 7.7E-09 | BIO14 | NA | NA | NA |
| BIO06 | FIR044 | FIR044_134 | 2.1E-07 | BIO15 | NA | NA | NA |
| BIO06 | PIE082 | PIE082_237 | 1.1E-06 | BIO16 | NA | NA | NA |
| BIO07 | FIR044 | FIR044_122 | 7.9E-16 | BIO17 | NA | NA | NA |
| BIO07 | POR008 | POR008_139 | 1.0E-15 | BIO18 | NA | NA | NA |
| BIO07 | PIE202 | PIE202_126 | 2.6E-15 | BIO19 | NA | NA | NA |
| BIO07 | FIR044 | FIR044_134 | 1.4E-13 | BIO20 | PIE202 | PIE202_126 | 1.6E-07 |
| BIO07 | POR008 | POR008_148 | 2.0E-12 | BIO21 | POR008 | POR008_139 | 1.4E-10 |
| BIO07 | PIE202 | PIE202_128 | 6.2E-12 | BIO21 | FIR044 | FIR044_134 | 2.9E-07 |
| BIO07 | PIE053 | PIE053_184 | 8.5E-08 | BIO21 | PIE202 | PIE202_128 | 4.1E-07 |
| BIO07 | PIE075 | PIE075_251 | 1.9E-07 | BIO21 | FIR044 | FIR044_122 | 4.2E-07 |
| BIO07 | VIT023 | VIT023_141 | 6.4E-07 | BIO21 | POR008 | POR008_148 | 5.1E-07 |
| BIO08 | FIR044 | FIR044_124 | 4.2E-09 | BIO22 | POR008 | POR008_139 | 6.5E-11 |
| BIO09 | POR008 | POR008_139 | 8.0E-16 | BIO22 | FIR044 | FIR044_134 | 7.0E-08 |
| BIO09 | PIE202 | PIE202_128 | 2.0E-15 | BIO22 | PIE202 | PIE202_126 | 2.4E-07 |
| BIO09 | FIR044 | FIR044_134 | 1.8E-14 | BIO23 | NA | NA | NA |
| BIO09 | PIE202 | PIE202_126 | 8.0E-14 | BIO24 | NA | NA | NA |
| BIO09 | FIR044 | FIR044_122 | 3.7E-13 | BIO25 | NA | NA | NA |
| BIO09 | POR008 | POR008_148 | 6.9E-12 | BIO26 | POR008 | POR008_139 | 5.1E-10 |
| BIO09 | PIE053 | PIE053_184 | 2.2E-09 | BIO26 | PIE202 | PIE202_126 | 2.9E-07 |
| BIO09 | VIT023 | VIT023_141 | 8.3E-08 | BIO26 | FIR044 | FIR044_134 | 5.2E-07 |
| BIO09 | POR016 | POR016_134 | 5.7E-07 | BIO27 | FIR044 | FIR044_134 | 5.0E-09 |

**Table S4-6a:**

| Variable | Loci | Alleles | adj. p-val | Variable | Loci | Alleles | adj. p-val |
|---|---|---|---|---|---|---|---|
| BIO01 | ZQR11 | ZQR11_276 | 2.6E-12 | BIO11 | FIR044 | FIR044_134 | 1.2E-23 |
| BIO01 | WAG020 | WAG020_292 | 2.6E-11 | BIO11 | PIE219 | PIE219_130 | 2.3E-16 |
| BIO01 | PIE148 | PIE148_206 | 4.6E-10 | BIO11 | PIE082 | PIE082_237 | 1.1E-12 |
| BIO01 | VIT031 | VIT0312_110 | 1.5E-08 | BIO11 | POR016 | POR016_144 | 9.5E-10 |
| BIO01 | FIR032 | FIR032_177 | 3.5E-08 | BIO11 | POR016 | POR016_134 | 2.1E-08 |
| BIO01 | POR028 | POR028_132 | 3.3E-07 | BIO11 | VIT023 | VIT023_138 | 2.5E-08 |
| BIO01 | POR034 | POR034_186 | 3.7E-07 | BIO11 | PIE137 | PIE137_352 | 6.5E-07 |
| BIO01 | WAG011 | WAG011_240 | 5.4E-07 | BIO11 | VIT028 | VIT028_163 | 8.9E-07 |
| BIO01 | PIE029 | PIE029_199 | 1.6E-06 | BIO11 | PIE075 | PIE075_250 | 1.4E-06 |
| BIO01 | PIE204 | PIE204_119 | 1.7E-06 | BIO11 | FIR106 | FIR106_241 | 1.6E-06 |
| BIO02 | PIE219 | PIE219_130 | 6.7E-32 | BIO12 | FIR044 | FIR044_134 | 6.4E-23 |
| BIO02 | FIR044 | FIR044_134 | 3.9E-31 | BIO12 | VIT023 | VIT023_138 | 7.5E-19 |
| BIO02 | PIE075 | PIE075_250 | 1.7E-20 | BIO12 | POR016 | POR016_134 | 6.6E-16 |
| BIO02 | FIR044 | FIR044_124 | 4.7E-18 | BIO12 | POR016 | POR016_144 | 2.1E-15 |
| BIO02 | PIE075 | PIE075_249 | 2.4E-14 | BIO12 | PIE219 | PIE219_130 | 4.2E-15 |
| BIO02 | PIE219 | PIE219_124 | 9.1E-09 | BIO12 | VIT023 | VIT023_141 | 1.2E-14 |
| BIO02 | VIT022 | VIT022_174 | 1.3E-06 | BIO12 | PIE082 | PIE082_237 | 3.9E-14 |
| BIO03 | POR016 | POR016_134 | 3.4E-20 | BIO12 | VIT028 | VIT028_163 | 2.2E-12 |
| BIO03 | VIT023 | VIT023_138 | 7.4E-20 | BIO12 | POR008 | POR008_148 | 1.2E-10 |
| BIO03 | POR016 | POR016_144 | 9.7E-17 | BIO12 | PIE137 | PIE137_352 | 1.5E-10 |
| BIO03 | VIT023 | VIT023_141 | 9.8E-16 | BIO12 | PIE082 | PIE082_240 | 3.4E-10 |
| BIO03 | PIE082 | PIE082_237 | 1.6E-14 | BIO12 | FIR106 | FIR106_241 | 2.4E-09 |
| BIO03 | PIE137 | PIE137_350 | 7.3E-14 | BIO12 | POR008 | POR008_139 | 3.0E-07 |
| BIO03 | PIE082 | PIE082_240 | 7.3E-13 | BIO12 | POR024 | POR024_181 | 6.6E-07 |
| BIO03 | PIE155 | PIE155_256 | 8.0E-13 | BIO12 | PIE137 | PIE137_350 | 1.1E-06 |
| BIO03 | FIR106 | FIR106_241 | 3.3E-12 | BIO12 | PIE202 | PIE202_128 | 2.1E-06 |
| BIO03 | POR024 | POR024_181 | 4.4E-12 | BIO13 | VIT023 | VIT023_138 | 4.4E-26 |
| BIO03 | PIE137 | PIE137_352 | 5.1E-12 | BIO13 | POR016 | POR016_144 | 3.9E-24 |
| BIO03 | VIT028 | VIT028_163 | 6.0E-12 | BIO13 | VIT023 | VIT023_141 | 5.3E-22 |
| BIO03 | POR008 | POR008_148 | 1.9E-10 | BIO13 | POR016 | POR016_134 | 7.0E-22 |
| BIO03 | FIR044 | FIR044_134 | 1.2E-08 | BIO13 | PIE082 | PIE082_240 | 3.8E-17 |
| BIO03 | POR008 | POR008_139 | 1.6E-07 | BIO13 | PIE082 | PIE082_237 | 2.2E-16 |
| BIO03 | VIT003 | VIT003_160 | 2.4E-07 | BIO13 | FIR106 | FIR106_241 | 9.4E-16 |
| BIO03 | PIE219 | PIE219_129 | 3.7E-07 | BIO13 | VIT028 | VIT028_163 | 1.3E-14 |
| BIO03 | POR025 | POR025_114 | 1.2E-06 | BIO13 | PIE137 | PIE137_352 | 1.3E-14 |
| BIO03 | PIE202 | PIE202_124 | 1.7E-06 | BIO13 | PIE137 | PIE137_350 | 4.3E-14 |
| BIO03 | FIR044 | FIR044_122 | 2.2E-06 | BIO13 | FIR044 | FIR044_134 | 1.6E-11 |
| BIO03 | PIE202 | PIE202_128 | 3.5E-06 | BIO13 | POR008 | POR008_148 | 2.3E-11 |
| BIO03 | PIE088 | PIE088_199 | 4.0E-06 | BIO13 | POR008 | POR008_139 | 5.2E-10 |
| BIO06 | FIR044 | FIR044_134 | 5.0E-19 | BIO13 | POR024 | POR024_181 | 1.6E-08 |
| BIO06 | PIE219 | PIE219_130 | 9.8E-18 | BIO13 | PIE202 | PIE202_128 | 3.5E-08 |
| BIO06 | PIE075 | PIE075_250 | 5.6E-12 | BIO13 | PIE219 | PIE219_130 | 5.8E-08 |
| BIO06 | FIR044 | FIR044_124 | 2.3E-11 | BIO13 | FIR044 | FIR044_122 | 1.8E-07 |
| BIO06 | PIE075 | PIE075_249 | 3.2E-08 | BIO13 | PIE219 | PIE219_129 | 2.3E-07 |
| BIO06 | PIE219 | PIE219_124 | 6.7E-07 | BIO13 | PIE202 | PIE202_124 | 2.6E-07 |
| BIO08 | VIT023 | VIT023_138 | 8.2E-14 | BIO13 | PIE238 | PIE238_222 | 2.3E-06 |
| BIO08 | POR016 | POR016_134 | 1.3E-11 | BIO19 | VIT023 | VIT023_138 | 7.9E-18 |
| BIO08 | PIE137 | PIE137_350 | 1.1E-10 | BIO19 | POR016 | POR016_134 | 1.7E-17 |
| BIO08 | POR016 | POR016_144 | 7.6E-10 | BIO19 | POR016 | POR016_144 | 2.6E-16 |
| BIO08 | VIT023 | VIT023_141 | 1.2E-09 | BIO19 | VIT023 | VIT023_141 | 4.2E-15 |
| BIO08 | FIR106 | FIR106_241 | 3.6E-08 | BIO19 | PIE082 | PIE082_237 | 2.2E-13 |
| BIO08 | PIE082 | PIE082_240 | 4.3E-08 | BIO19 | FIR044 | FIR044_134 | 4.8E-13 |
| BIO08 | VIT010 | VIT010_150 | 1.1E-06 | BIO19 | PIE082 | PIE082_240 | 6.7E-13 |
| BIO08 | PIE137 | PIE137_352 | 1.2E-06 | BIO19 | FIR106 | FIR106_241 | 1.4E-11 |
| BIO08 | VIT028 | VIT028_163 | 1.6E-06 | BIO19 | PIE137 | PIE137_352 | 6.9E-11 |
| BIO08 | PIE088 | PIE088_199 | 2.0E-06 | BIO19 | VIT028 | VIT028_163 | 7.7E-11 |
| | | | | BIO19 | POR008 | POR008_148 | 1.0E-10 |
| | | | | BIO19 | PIE137 | PIE137_350 | 8.7E-10 |
| | | | | BIO19 | POR008 | POR008_139 | 7.1E-08 |
| | | | | BIO19 | POR024 | POR024_181 | 5.9E-07 |
| | | | | BIO19 | PIE219 | PIE219_129 | 1.2E-06 |
| | | | | BIO19 | PIE202 | PIE202_128 | 1.3E-06 |
| | | | | BIO19 | PIE219 | PIE219_130 | 1.4E-06 |
| | | | | BIO20 | NA | NA | NA |

**Table S4-6b:**

| Variable | Loci | Alleles | adj. p-values |
|---|---|---|---|
| BIO01 | NA | NA | NA |
| BIO02 | FIR044 | FIR044_124 | 4.6E-16 |
| BIO02 | FIR044 | FIR044_134 | 6.3E-11 |
| BIO02 | PIE219 | PIE219_124 | 8.7E-09 |
| BIO02 | PIE137 | PIE137_350 | 1.2E-08 |
| BIO02 | VIT022 | VIT022_174 | 3.2E-08 |
| BIO02 | FIR030 | FIR030_134 | 1.1E-07 |
| BIO02 | VIT023 | VIT023_141 | 2.1E-07 |
| BIO02 | PIE137 | PIE137_352 | 3.6E-07 |
| BIO02 | PIE075 | PIE075_250 | 4.9E-07 |
| BIO03 | NA | NA | NA |
| BIO06 | FIR044 | FIR044_124 | 2.2E-15 |
| BIO06 | PIE137 | PIE137_350 | 1.7E-14 |
| BIO06 | FIR044 | FIR044_134 | 9.4E-13 |
| BIO06 | FIR030 | FIR030_134 | 7.0E-08 |
| BIO06 | POR016 | POR016_134 | 1.8E-07 |
| BIO06 | VIT023 | VIT023_141 | 3.0E-07 |
| BIO06 | PIE075 | PIE075_250 | 4.0E-07 |
| BIO06 | PIE219 | PIE219_124 | 8.7E-07 |
| BIO08 | FIR044 | FIR044_124 | 4.0E-12 |
| BIO08 | FIR044 | FIR044_134 | 1.8E-08 |
| BIO08 | FIR030 | FIR030_134 | 1.5E-07 |
| BIO11 | FIR044 | FIR044_124 | 3.6E-14 |
| BIO11 | FIR044 | FIR044_134 | 1.7E-08 |
| BIO11 | PIE137 | PIE137_350 | 4.7E-07 |
| BIO12 | FIR044 | FIR044_124 | 6.4E-22 |
| BIO12 | PIE137 | PIE137_350 | 6.2E-15 |
| BIO12 | FIR044 | FIR044_134 | 1.5E-11 |
| BIO12 | PIE075 | PIE075_250 | 6.7E-09 |
| BIO12 | VIT022 | VIT022_174 | 3.0E-08 |
| BIO12 | FIR030 | FIR030_134 | 7.3E-08 |
| BIO12 | PIE219 | PIE219_124 | 2.1E-07 |
| BIO12 | FIR030 | FIR030_137 | 2.3E-07 |
| BIO12 | VIT007 | VIT007_134 | 4.4E-07 |
| BIO12 | PIE223 | PIE223_227 | 1.7E-06 |
| BIO13 | FIR044 | FIR044_124 | 1.3E-29 |
| BIO13 | FIR044 | FIR044_134 | 3.0E-16 |
| BIO13 | PIE219 | PIE219_124 | 2.5E-15 |
| BIO13 | PIE137 | PIE137_350 | 2.1E-14 |
| BIO13 | VIT022 | VIT022_174 | 1.8E-12 |
| BIO13 | FIR030 | FIR030_134 | 4.3E-12 |
| BIO13 | PIE137 | PIE137_352 | 7.4E-11 |
| BIO13 | PIE075 | PIE075_250 | 1.1E-10 |
| BIO13 | VIT023 | VIT023_141 | 8.6E-10 |
| BIO13 | POR016 | POR016_134 | 1.5E-09 |
| BIO13 | FIR030 | FIR030_137 | 5.3E-09 |
| BIO13 | PIE219 | PIE219_130 | 8.6E-08 |
| BIO13 | VIT007 | VIT007_134 | 5.5E-07 |
| BIO13 | WAG079 | WAG079_321 | 9.0E-07 |
| BIO13 | POR038 | POR038_137 | 1.1E-06 |
| BIO19 | FIR044 | FIR044_124 | 1.1E-15 |
| BIO19 | PIE137 | PIE137_350 | 9.3E-13 |
| BIO19 | FIR030 | FIR030_134 | 4.8E-11 |
| BIO19 | FIR044 | FIR044_134 | 9.5E-09 |
| BIO19 | PIE219 | PIE219_124 | 3.8E-07 |
| BIO19 | FIR030 | FIR030_137 | 4.0E-07 |
| BIO19 | PIE219 | PIE219_130 | 8.5E-07 |
| BIO19 | VIT007 | VIT007_134 | 1.3E-06 |
| BIO20 | PIE137 | PIE137_350 | 3.1E-12 |
| BIO20 | FIR044 | FIR044_124 | 2.1E-11 |
| BIO20 | FIR044 | FIR044_134 | 2.1E-10 |
| BIO20 | FIR030 | FIR030_134 | 4.7E-07 |

**Table S4-7:**

| *Quercus faginea* | | | | | | *Quercus pyrenaica* | | | | | |
|---|---|---|---|---|---|---|---|---|---|---|---|
| Bayescan[1] | Tess3r[2] | Bayescenv[3] | Lfmm[4] | LfmmAll[5] | 10-LfmmAll[6] | Bayescan[1] | Tess3r[7] | Bayescenv[3] | Lfmm[4] | LfmmAll[5] | 10-LfmmAll[6] |
| FIR013 | FIR013 | FIR013 | FIR013 | | | FIR013 | | | FIR013 | FIR013 | |
| | | | FIR023 | | | | | | | | |
| FIR030 | FIR030 | FIR030 | | FIR030 | | FIR030 | | | FIR030 | FIR030 | FIR030 |
| | | | | | FIR032 | | | | | | |
| FIR033 | | FIR033 | | FIR033 | | FIR033 | | FIR033 | FIR033 | | |
| FIR044 | FIR044 | FIR044 | FIR044 | FIR044 | FIR044 | FIR044 | | FIR044 | FIR044 | FIR044 | FIR044 |
| | | | | | | | | FIR047 | | | |
| | FIR048 | | | | | | | | | | |
| | | FIR065 | | | | | | | | | |
| | FIR104 | | | | | | | | | | |
| FIR106 | FIR106 | FIR106 | FIR106 | FIR106 | FIR106 | | FIR106 | FIR106 | FIR106 | FIR106 | |
| | | GOT021 | | | | | | | | | |
| | | | | PIE002 | | | | | | | |
| | | | | | | | | PIE023 | | | |
| | | | PIE029 | | | | | | | | |
| | | | | PIE036 | | | | | | | |
| | | | | PIE053 | | | PIE053 | | PIE053 | PIE053 | |
| | | | | | | | | | | PIE071 | |
| | | | PIE075 | PIE075 | PIE075 | | PIE075 | | PIE075 | PIE075 | PIE075 |
| | PIE076 | | | PIE076 | | | | | | | |
| PIE82 | PIE082 | PIE082 | PIE082 | PIE082 | PIE082 | PIE082 | PIE082 | | PIE082 | PIE082 | |
| | | | | PIE088 | PIE088 | | | | | | |
| | PIE089 | | | | | | | | | | |
| | | | PIE102 | | | PIE102 | | | | | |
| PIE137 | | PIE137 | | PIE137 | PIE137 | PIE137 | | PIE137 | PIE137 | PIE137 | PIE137 |
| | | | | | PIE148 | | | | | | |
| | | | | | PIE155 | | PIE155 | | PIE155 | | |
| | | | | | | | | | | PIE175 | |
| | | | PIE196 | | | | PIE196 | | | | |
| | PIE202 | PIE202 | PIE202 | PIE202 | PIE202 | | | PIE202 | PIE202 | PIE202 | |
| | PIE204 | | | | PIE204 | PIE204 | PIE204 | | PIE204 | PIE204 | |
| PIE219 | PIE219 | PIE219 | PIE219 | PIE219 | PIE219 | PIE219 | PIE219 | | PIE219 | PIE219 | PIE219 |
| PIE223 | | | | | | | | | | | PIE223 |
| | | | | | | | PIE228 | | PIE228 | | |
| | | | | | PIE238 | | | | | | |
| | PIE242 | | | | | | | | | | |
| | | | | | | | | | | PIE259 | |
| | | | | | | | | PIE264 | | | |
| POR008 | | POR008 | POR008 | POR008 | POR008 | POR008 | | POR008 | POR008 | POR008 | |
| | POR014 | | | | | | | | | | |
| POR016 | | POR016 | POR016 | POR016 | POR016 | | | POR016 | POR016 | POR016 | POR016 |
| | | | | | | | | | | POR020 | |
| | | | | POR024 | POR024 | | POR024 | | | | |
| | POR025 | | | POR025 | POR025 | | POR025 | | | | |
| | | | | | POR028 | | | | | | |
| | POR030 | | | | | | POR030 | | | | |
| | | | | | POR034 | | | | | | |
| | | | | | | POR038 | POR038 | POR038 | POR038 | | POR038 |
| | POR039 | | | | | | POR039 | | | | |
| | POR040 | | | | | | | | | | |
| | | | | | VIT003 | | | | | | |
| VIT007 | | VIT007 | | | | VIT007 | VIT007 | | VIT007 | | VIT007 |
| VIT010 | VIT010 | | VIT010 | VIT010 | VIT10 | | | | | | |
| VIT018 | | VIT018 | | | | | | | | | |
| | | | | | | | | | | VIT020 | |
| | | | | VIT022 | VIT022 | VIT022 | | VIT022 | VIT022 | | VIT022 |
| VIT023 | | VIT023 | VIT023 | VIT023 | VIT023 | VIT023 | VIT023 | VIT023 | VIT023 | VIT023 | VIT023 |
| VIT025 | | | VIT025 | | | | | | | | |
| | | | | VIT026 | | | | | | | |
| | | VIT028 | VIT028 | VIT028 | VIT028 | VIT028 | VIT028 | | VIT028 | | |
| | | VIT031.1 | | | VIT031.1 | | | | | | |
| | | | | | | | | | | VIT037 | |
| | | | VIT107 | | | | VIT107 | | | | |
| | | | | | WAG011 | | | | | | |
| | WAG020 | | | | WAG020 | | WAG020 | | WAG020 | WAG020 | |
| | | | | | | | WAG043 | | | | |
| | | | | | | | WAG079 | | WAG079 | | WAG079 |
| | ZQP15 | | | ZQP15 | | | | | | | |
| | | | | | ZQR11 | | | | | | |
| | | | | | | | ZQR30 | | | | |
| | | ZQR87 | ZQR87 | | | | ZQR87 | | ZQR87 | | |

1: Bayescan; FDR level q=0.1
2: FDR level q=0.01
3: Bayescenv; FDR level q=0.1. Gray cells, additional loci involved in the 17 'correlated' Bioclim variables
4: Lfmm; Bayescenv; FDR level q=0.001. Gray cells, additional loci in the 17 'correlated' Bioclim variables
5: FDR level q=0.0001; 27 Bioclim variables fit at once
6: FDR level q=0.0001: 10 uncorrelated Biocllim variables fit at once
7: FDR level q=0.1

**Figure S4-1**

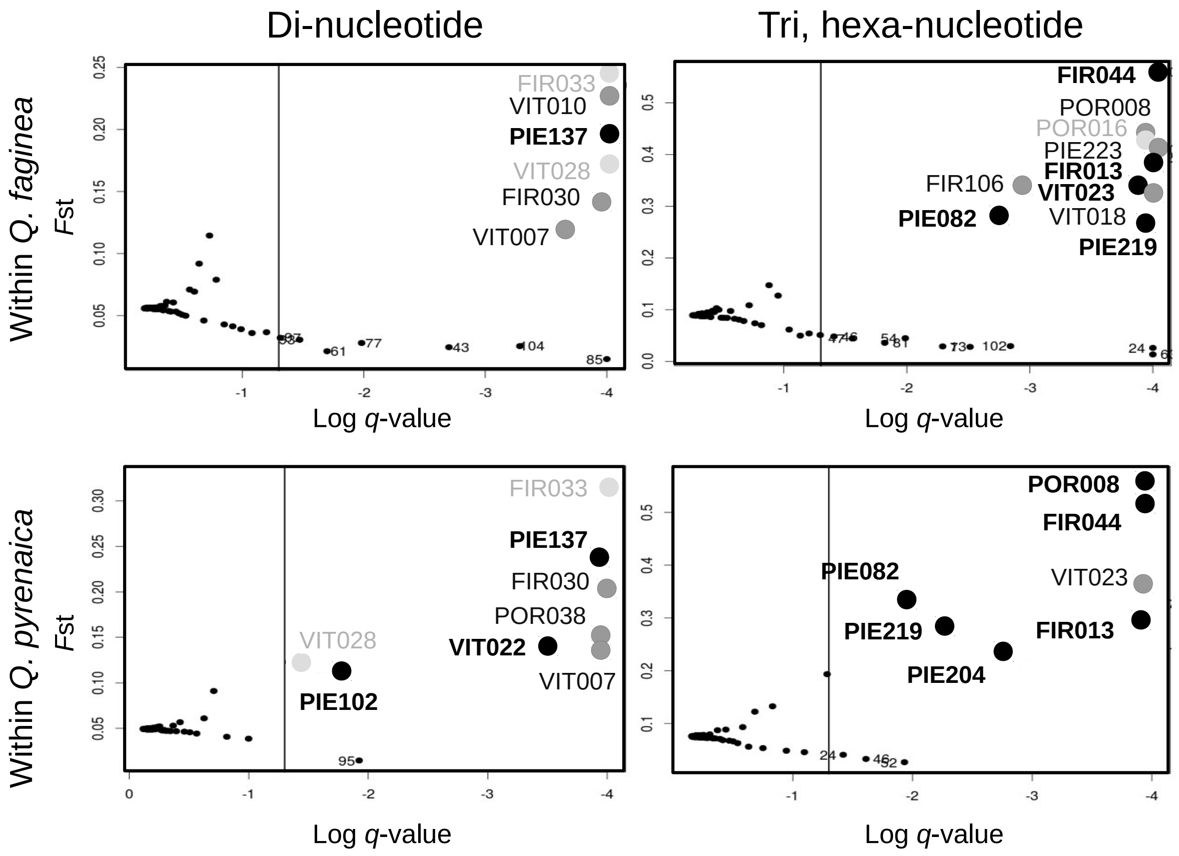
Di-nucleotide
Tri, hexa-nucleotide
Within Q. faginea
Within Q. pyrenaica
Fst
Log q-value
FIR033
VIT010
PIE137
VIT028
FIR030
VIT007
FIR044
POR008
POR016
PIE223
FIR013
VIT023
FIR106
VIT018
PIE082
PIE219
POR038
VIT022
PIE102
PIE204

**Figure S4-2a**

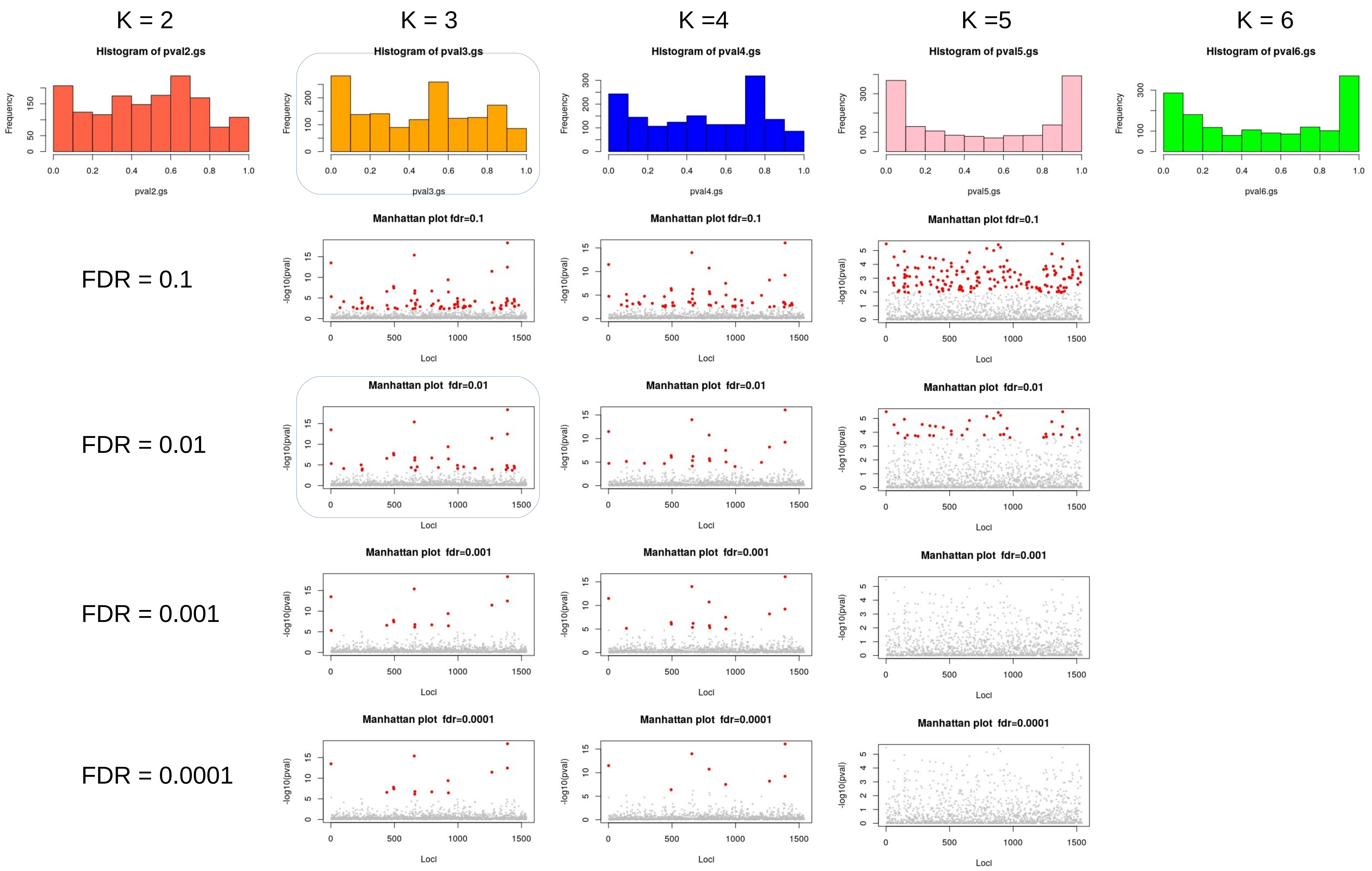

K = 2
K = 3
K =4
K =5
K = 6
Histogram of pval2.gs
Histogram of pval3.gs
Histogram of pval4.gs
Histogram of pval5.gs
Histogram of pval6.gs
Frequency
pval2.gs
pval3.gs
pval4.gs
pval5.gs
pval6.gs
FDR = 0.1
FDR = 0.01
FDR = 0.001
FDR = 0.0001
Manhattan plot fdr=0.1
Manhattan plot fdr=0.01
Manhattan plot fdr=0.001
Manhattan plot fdr=0.0001
-log10(pval)
Loci

**Figure S4-2b**

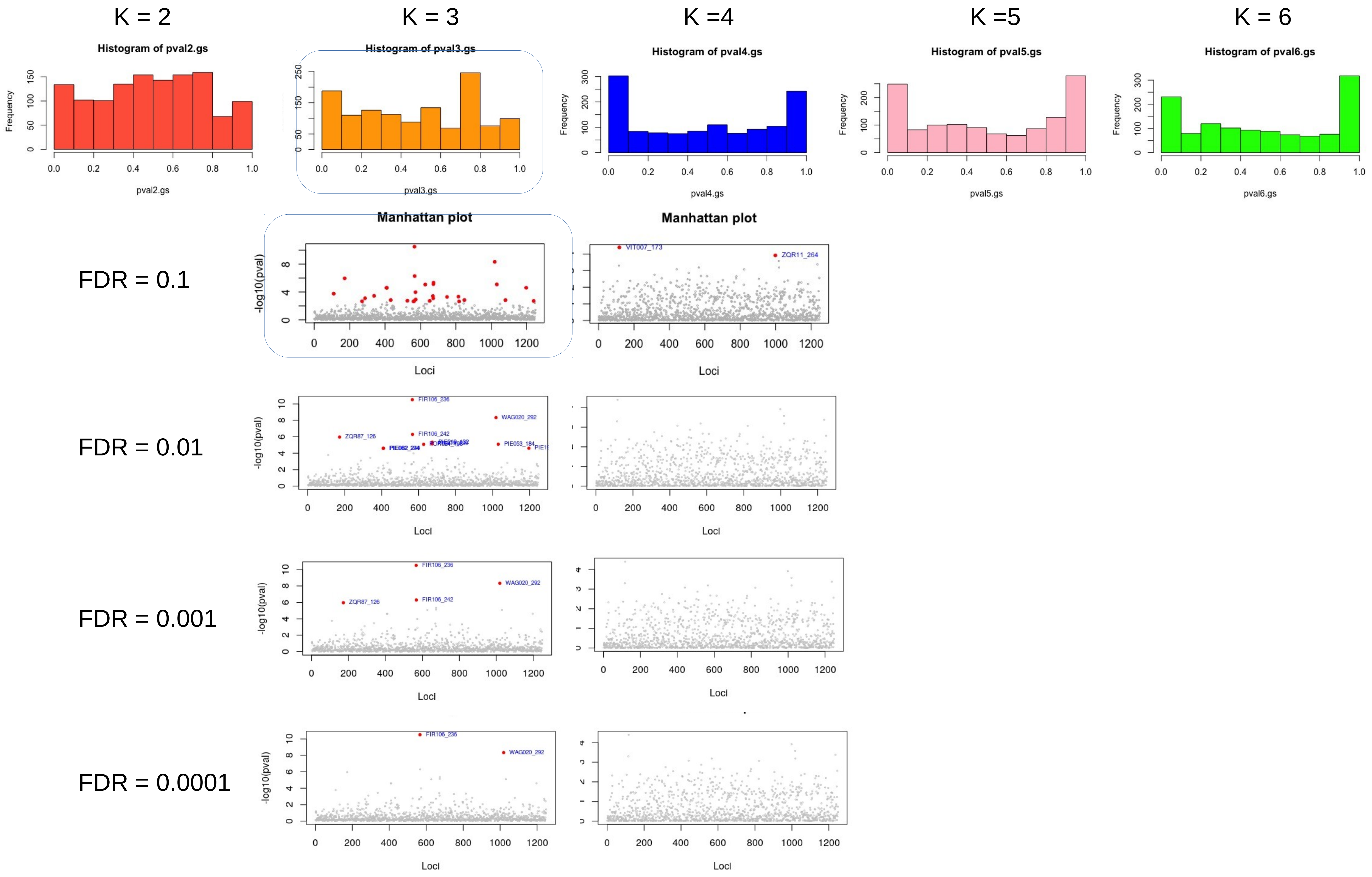

K = 2
K = 3
K =4
K =5
K = 6
Histogram of pval2.gs
Histogram of pval3.gs
Histogram of pval4.gs
Histogram of pval5.gs
Histogram of pval6.gs
Frequency
pval2.gs
pval3.gs
pval4.gs
pval5.gs
pval6.gs
FDR = 0.1
FDR = 0.01
FDR = 0.001
FDR = 0.0001
Manhattan plot
-log10(pval)
Loci
VIT007_173
ZQR11_264
FIR106_236
WAG020_292
ZQR87_126
FIR106_242
PIE053_184

**Figure S4-3a.**

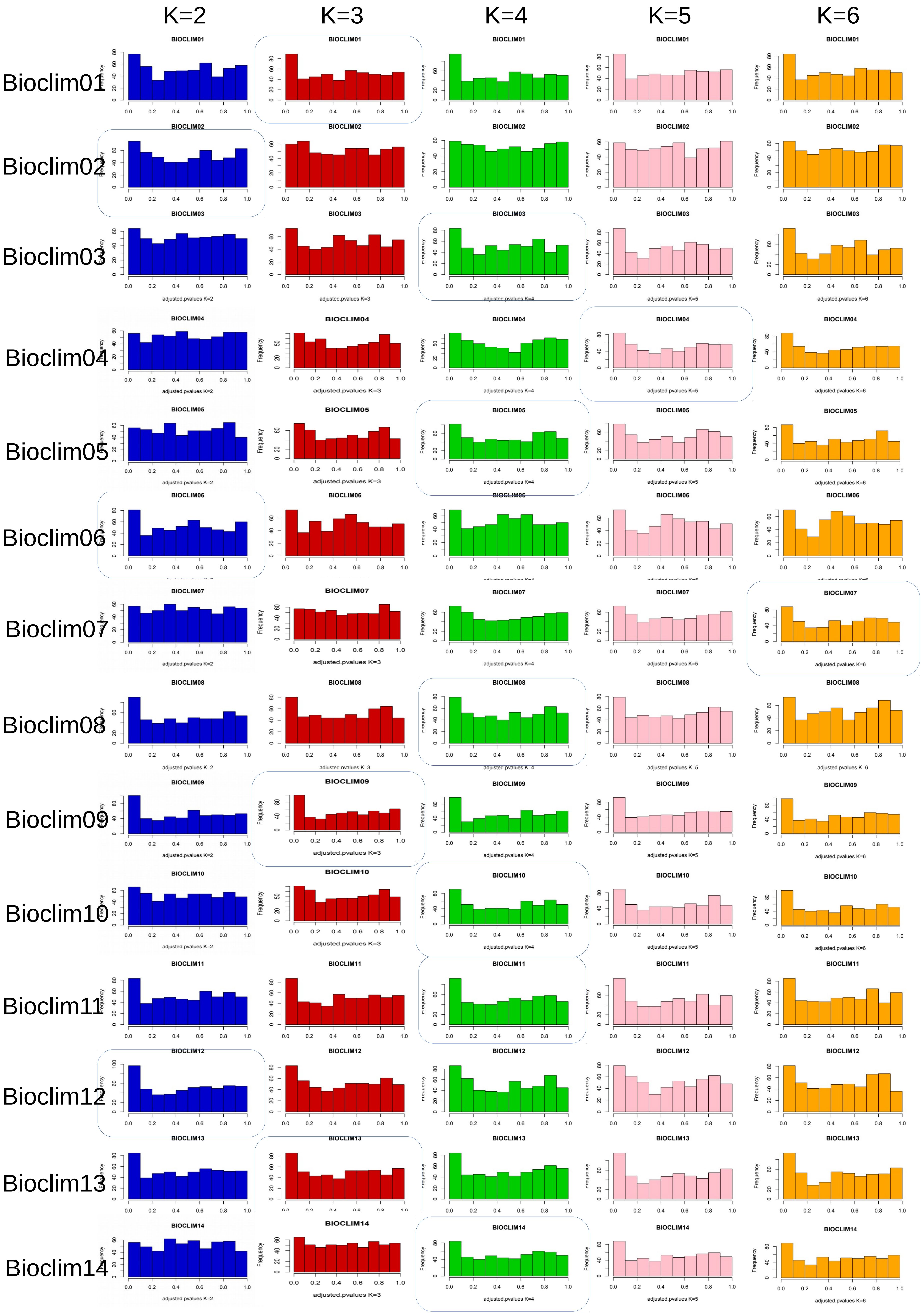
K=2
K=3
K=4
K=5
K=6
Bioclim01
Bioclim02
Bioclim03
Bioclim04
Bioclim05
Bioclim06
Bioclim07
Bioclim08
Bioclim09
Bioclim10
Bioclim11
Bioclim12
Bioclim13
Bioclim14

**Figure S4-3a (continue):**

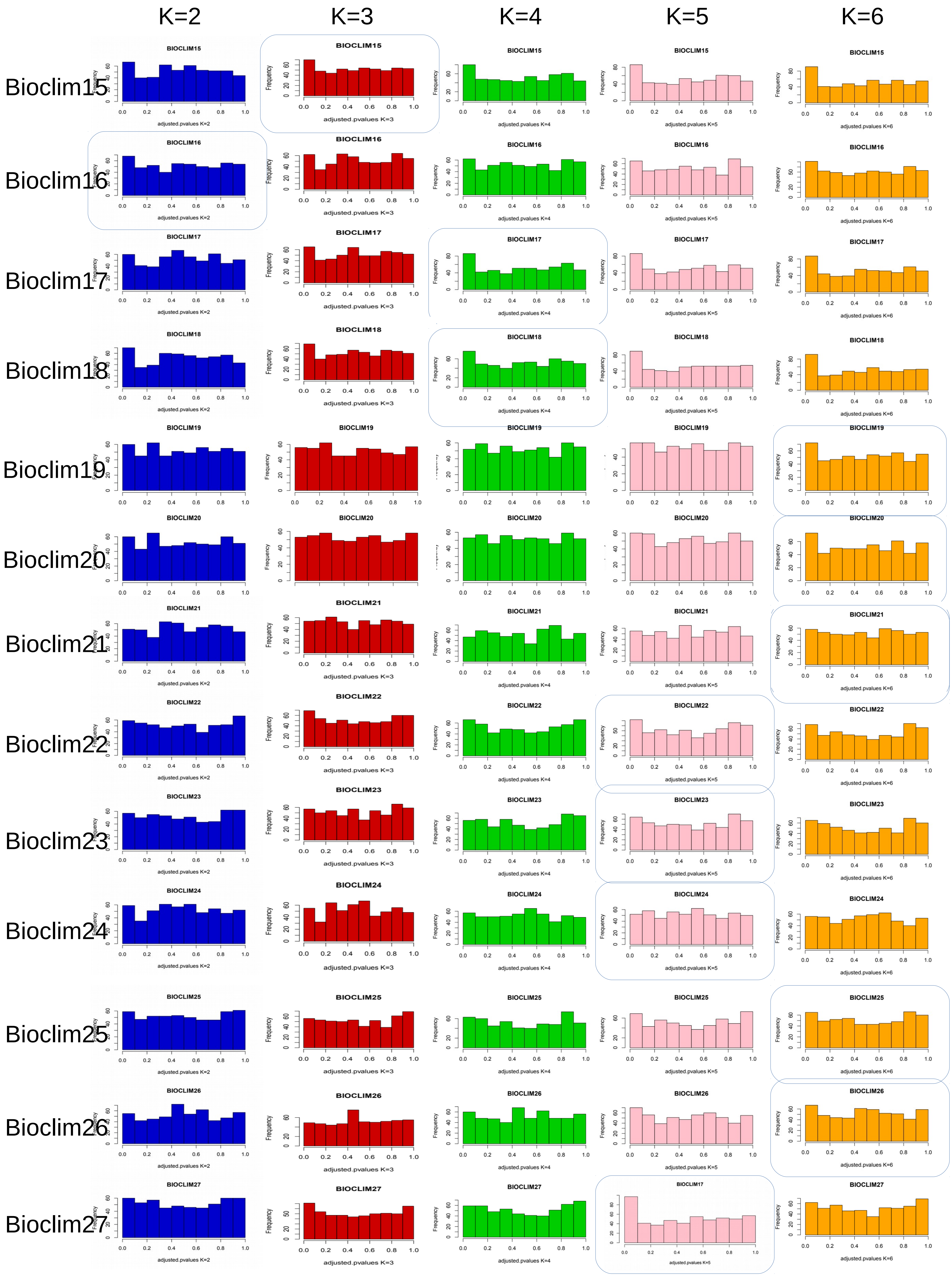

**Figure S4-3a (continue):**

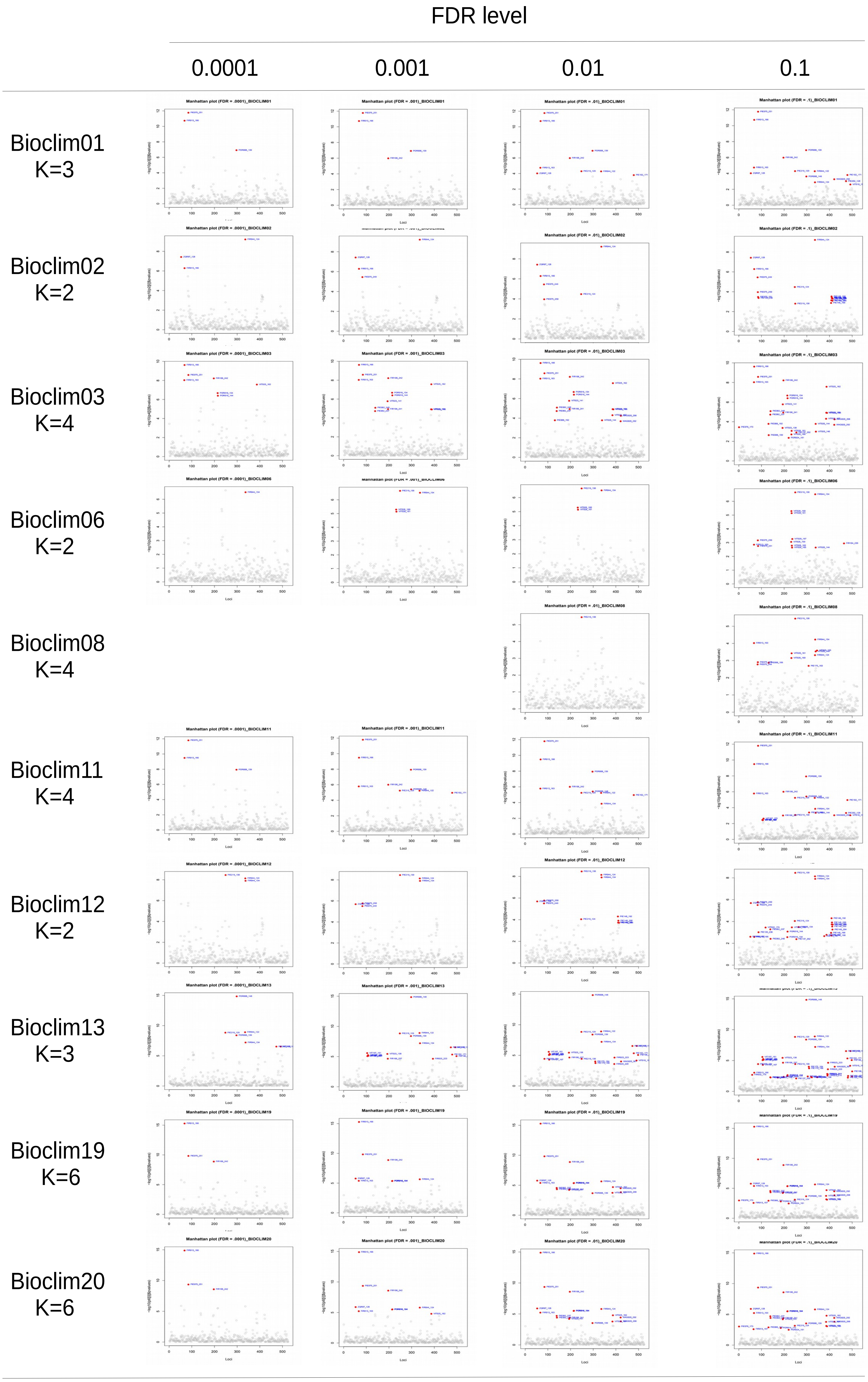

**Figure S4-3b:**

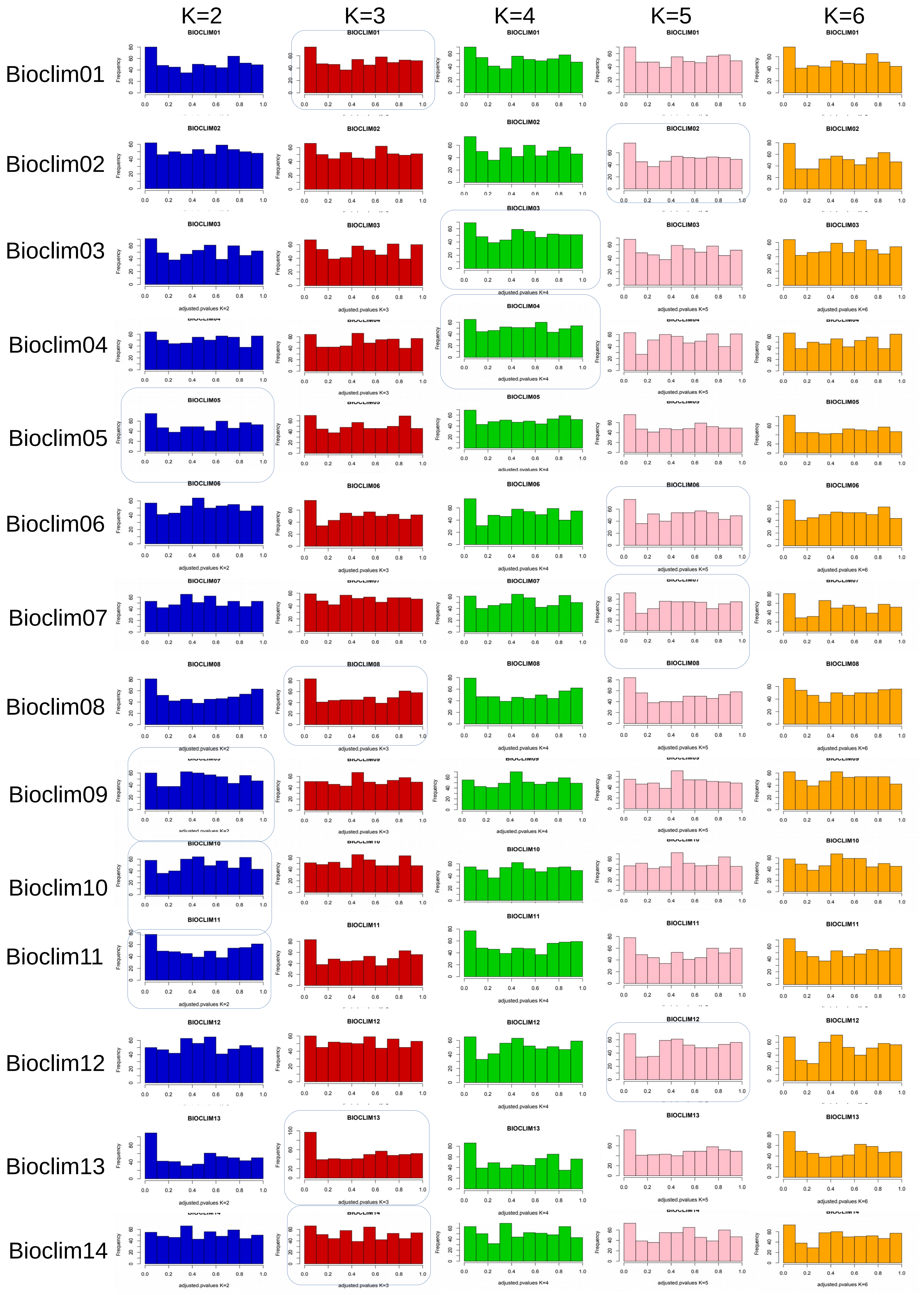
K=2
K=3
K=4
K=5
K=6
Bioclim01
Bioclim02
Bioclim03
Bioclim04
Bioclim05
Bioclim06
Bioclim07
Bioclim08
Bioclim09
Bioclim10
Bioclim11
Bioclim12
Bioclim13
Bioclim14

**Figure S4-3b (continue):**

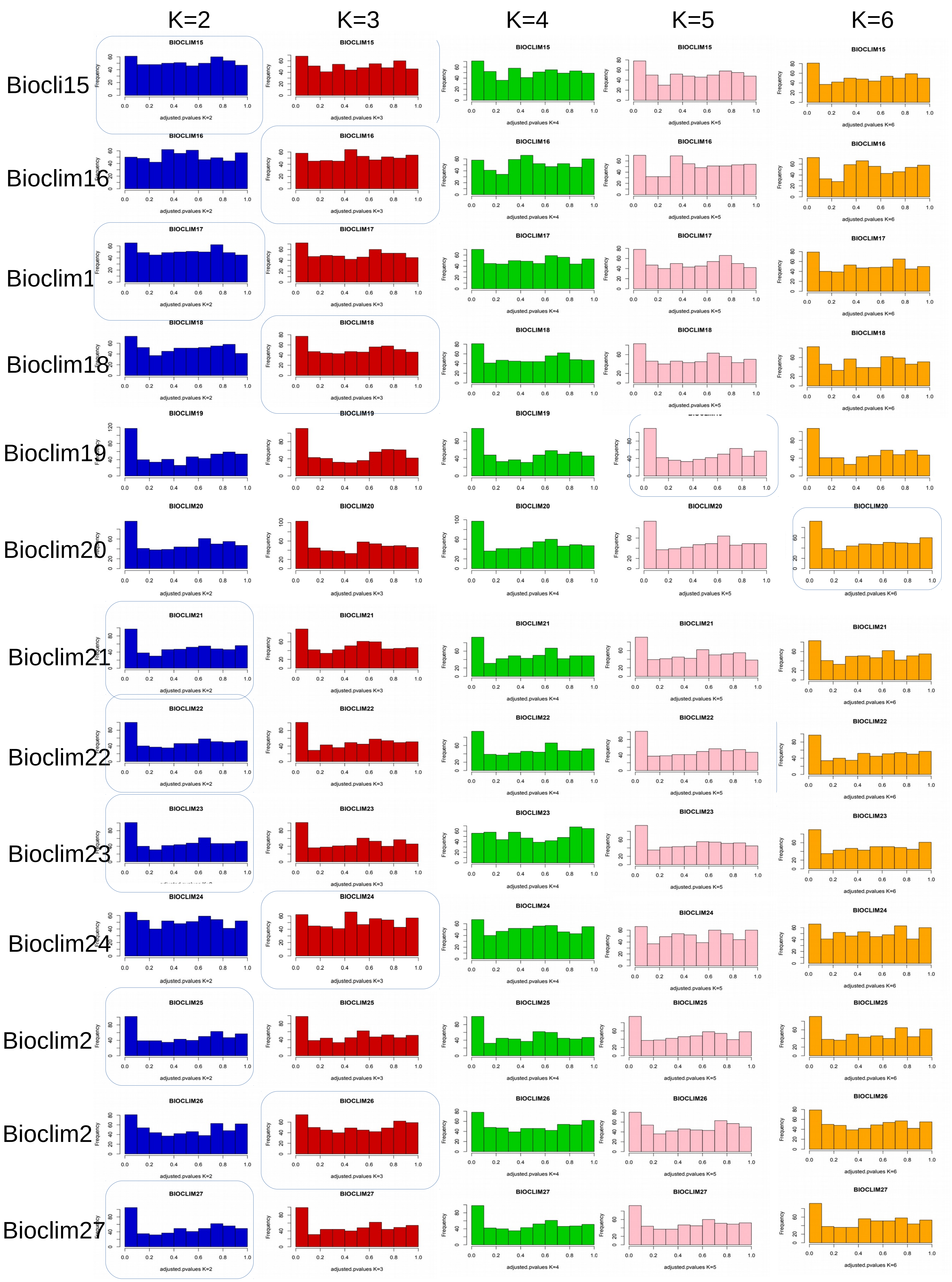

**Figure S4-4a:**

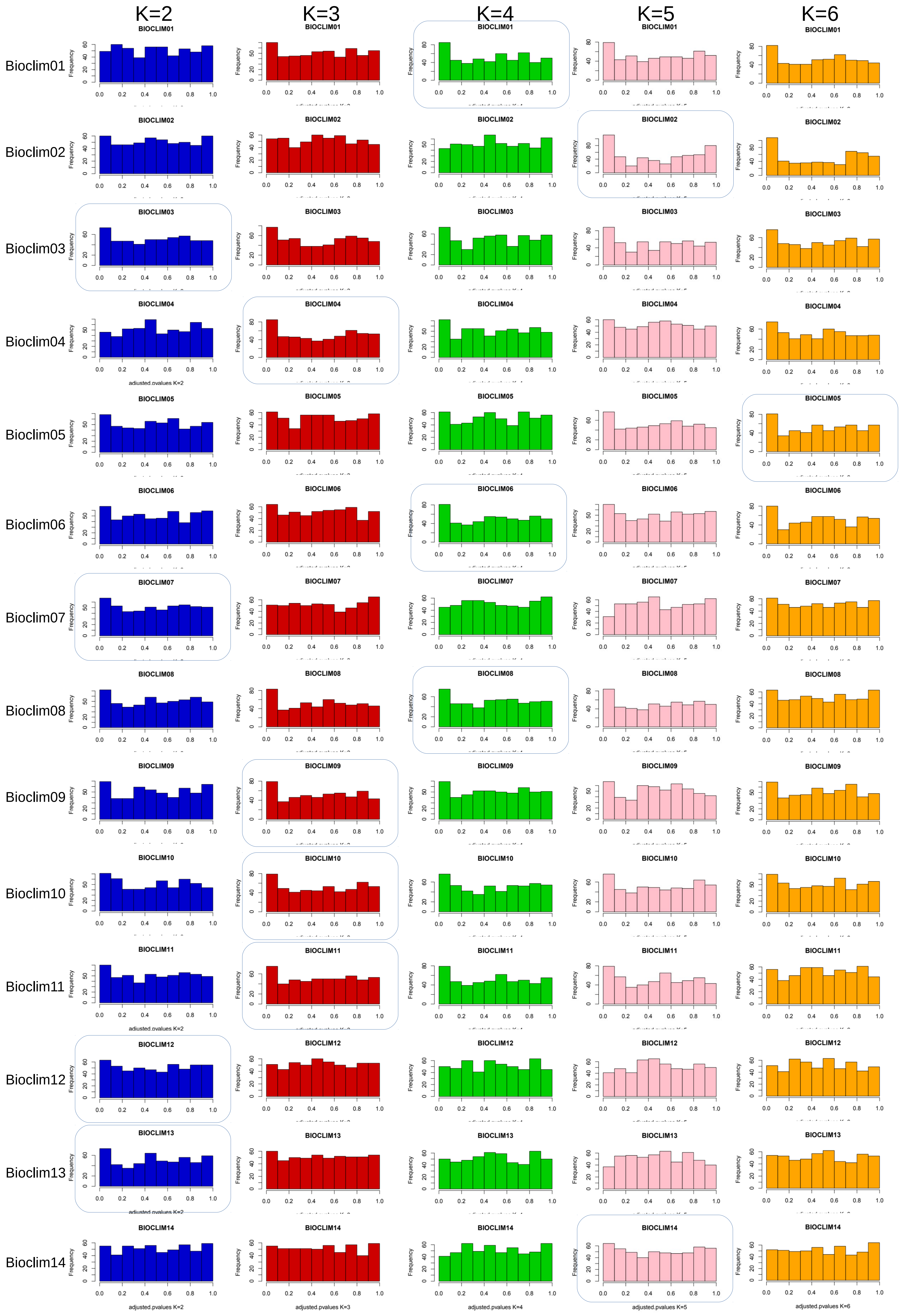
K=2
K=3
K=4
K=5
K=6
Bioclim01
Bioclim02
Bioclim03
Bioclim04
Bioclim05
Bioclim06
Bioclim07
Bioclim08
Bioclim09
Bioclim10
Bioclim11
Bioclim12
Bioclim13
Bioclim14
adjusted.pvalues K=2
adjusted.pvalues K=3
adjusted.pvalues K=4
adjusted.pvalues K=5
adjusted.pvalues K=6

**Figure S4-4a (continue):**

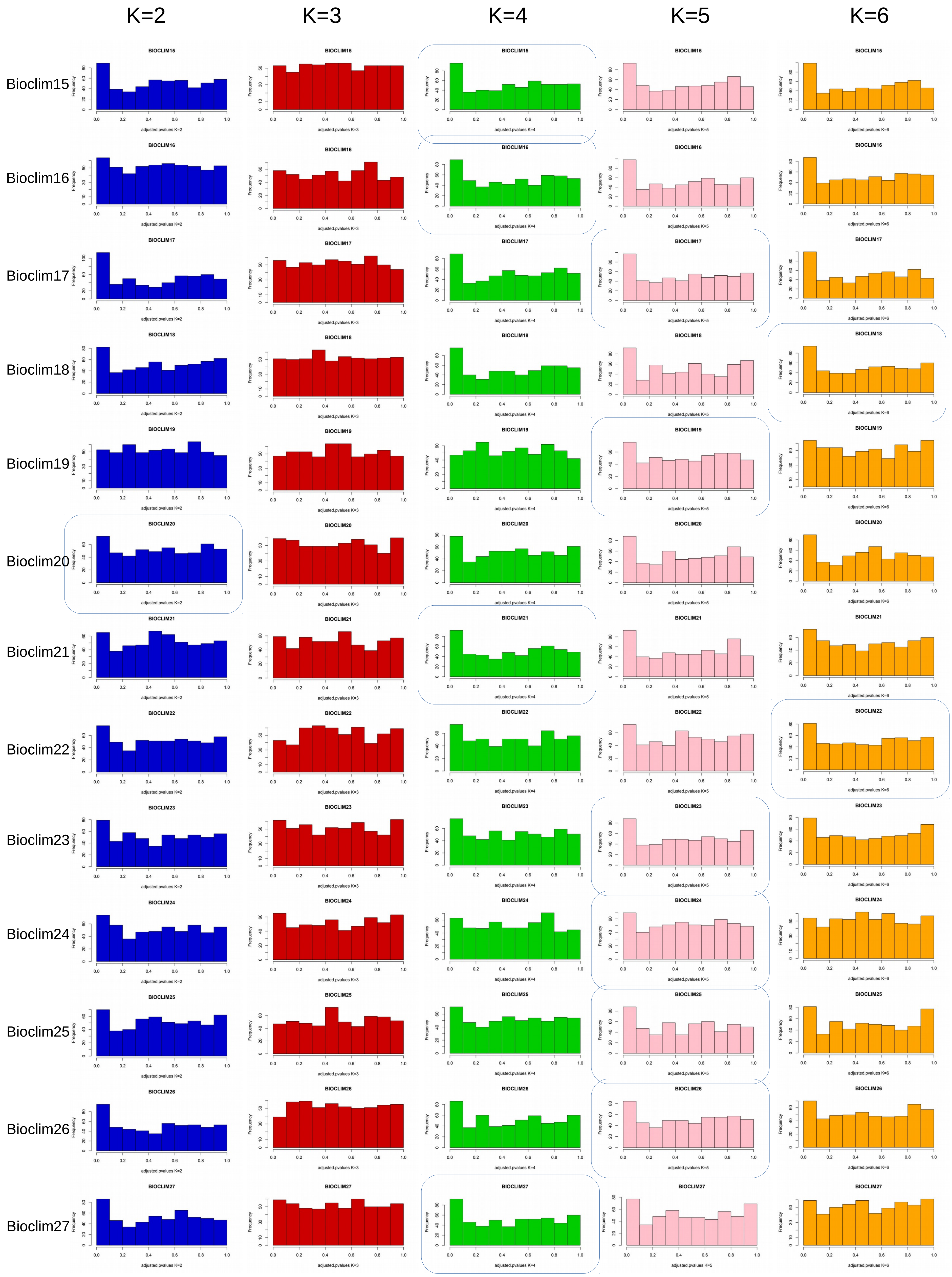

**Figure S4-4b:**

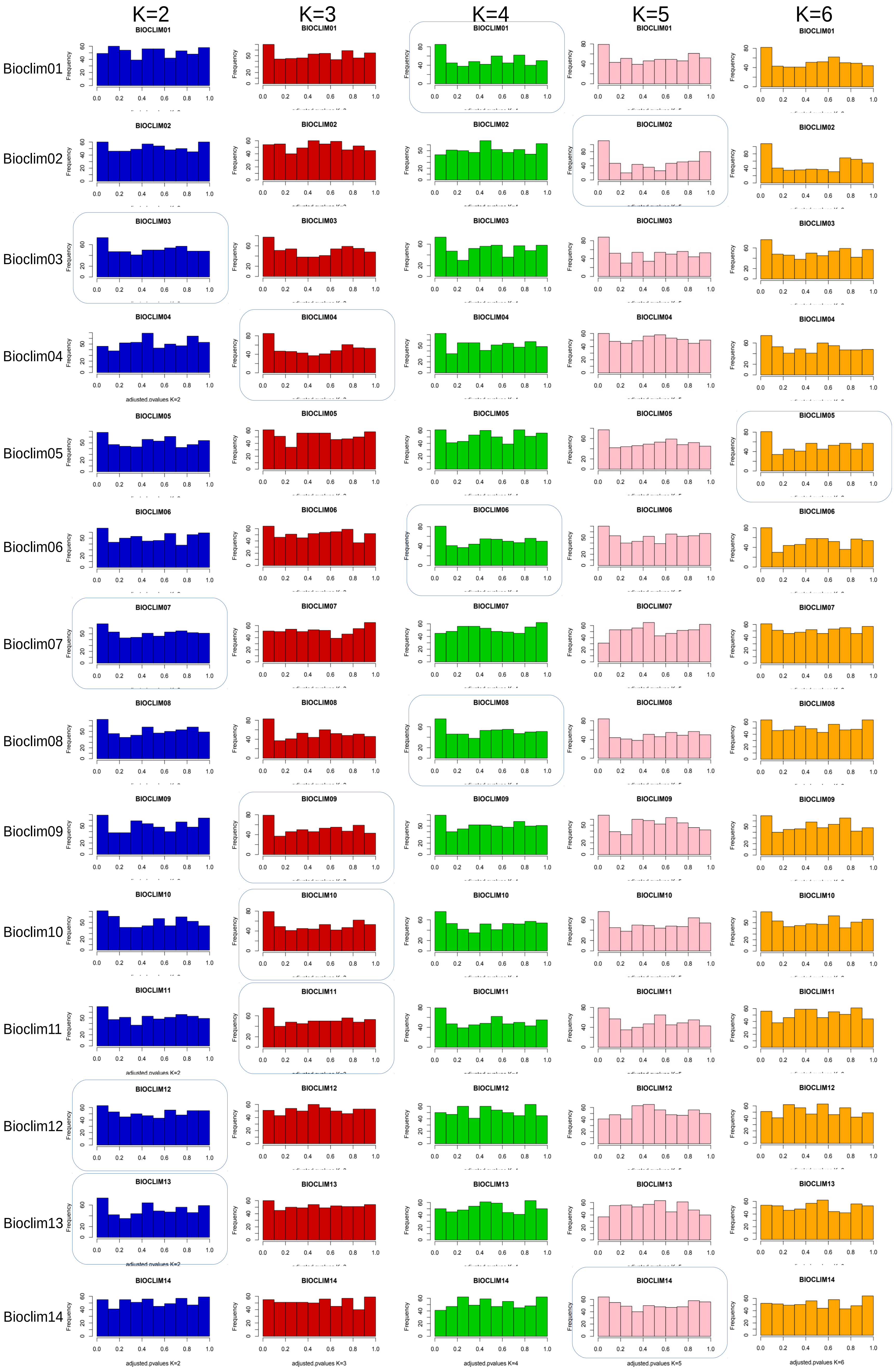
K=2
K=3
K=4
K=5
K=6
Bioclim01
Bioclim02
Bioclim03
Bioclim04
Bioclim05
Bioclim06
Bioclim07
Bioclim08
Bioclim09
Bioclim10
Bioclim11
Bioclim12
Bioclim13
Bioclim14
Frequency
adjusted.pvalues K=2
adjusted.pvalues K=3
adjusted.pvalues K=4
adjusted.pvalues K=5
adjusted.pvalues K=6

**Figure S4-4b (continue)**

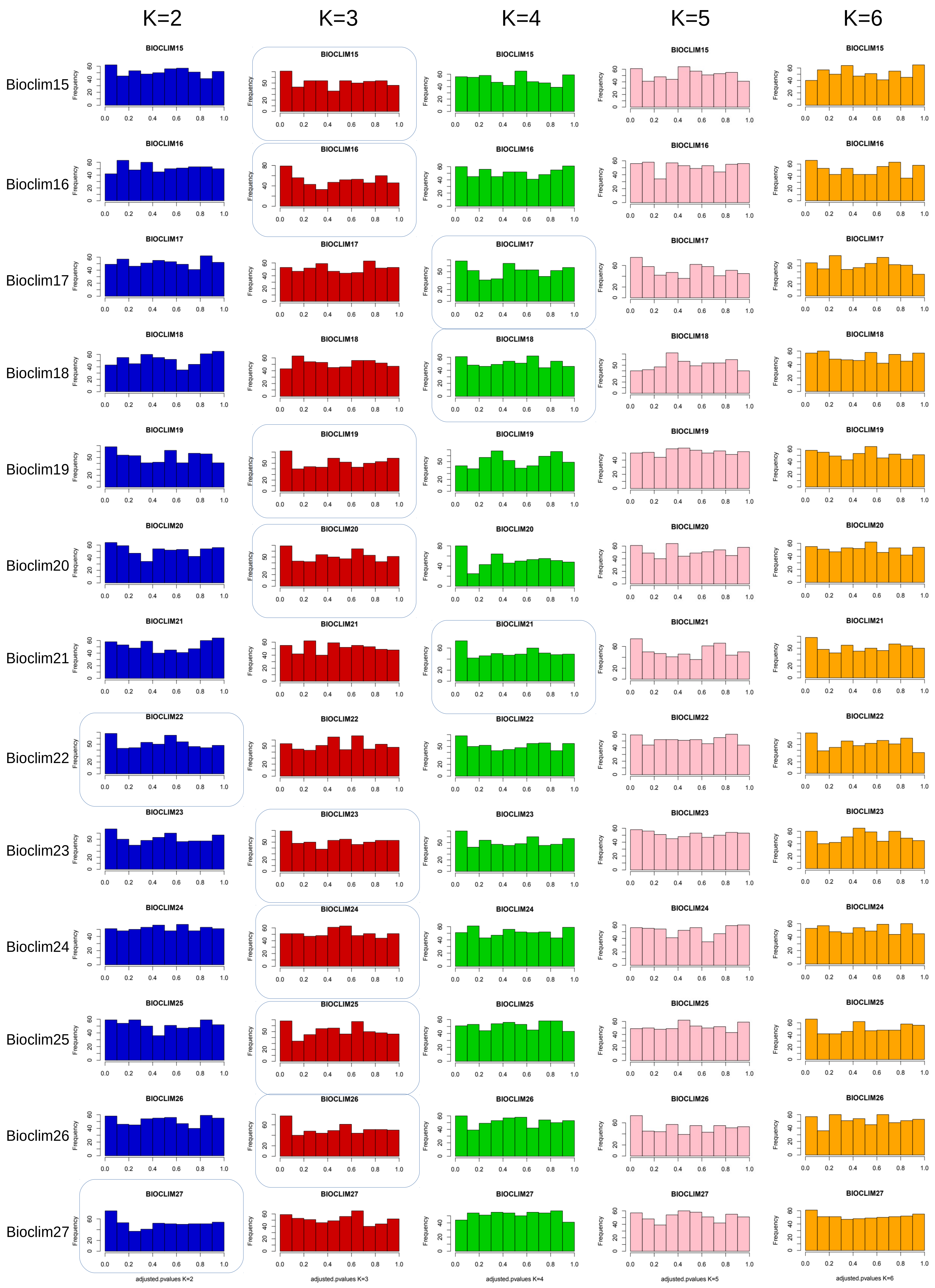

**Fgure S4-5a**

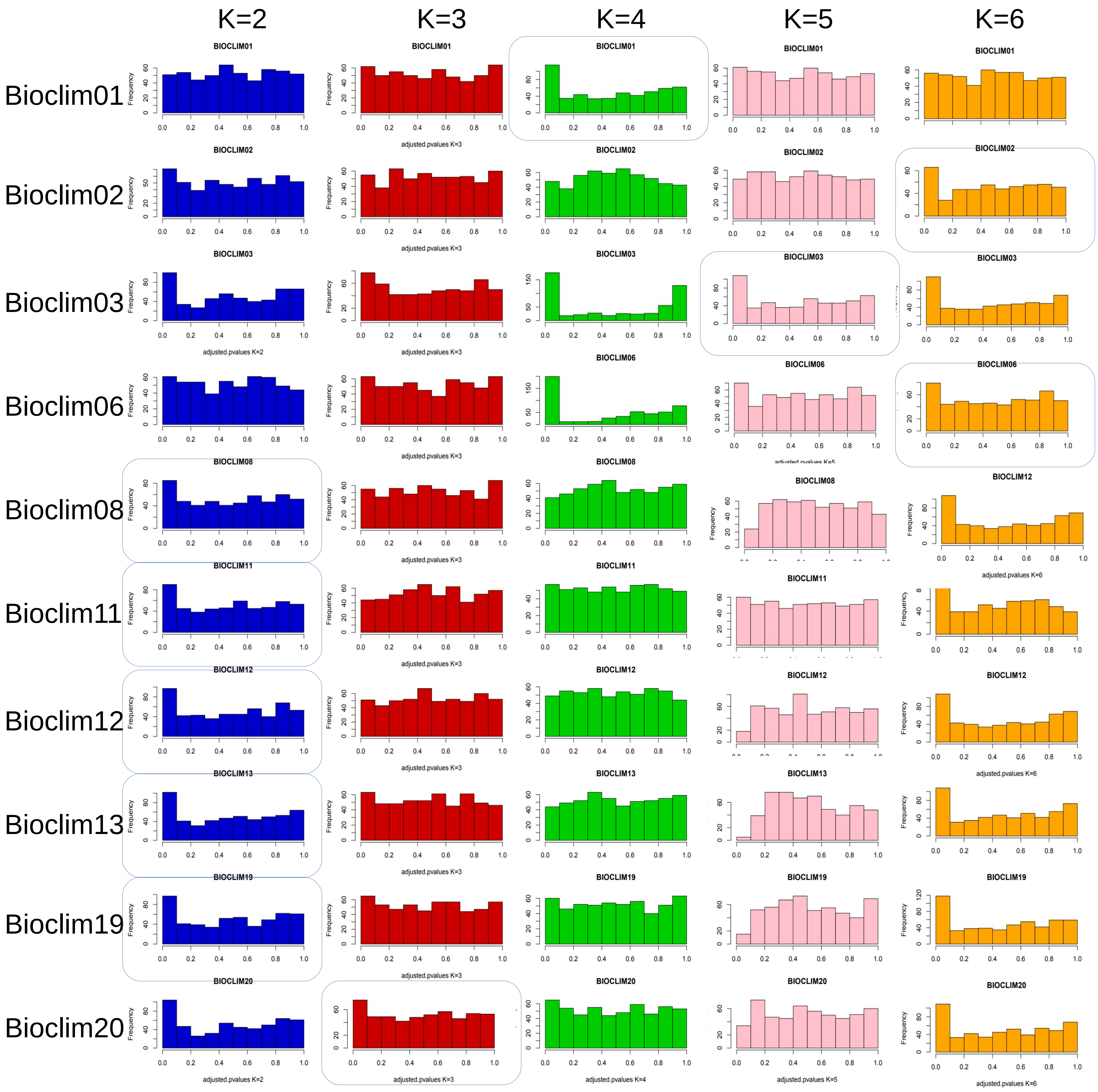
K=2
K=3
K=4
K=5
K=6
Bioclim01
Bioclim02
Bioclim03
Bioclim06
Bioclim08
Bioclim11
Bioclim12
Bioclim13
Bioclim19
Bioclim20
adjusted.pvalues K=2
adjusted.pvalues K=3
adjusted.pvalues K=4
adjusted.pvalues K=5
adjusted.pvalues K=6

**Figure S4-5b**

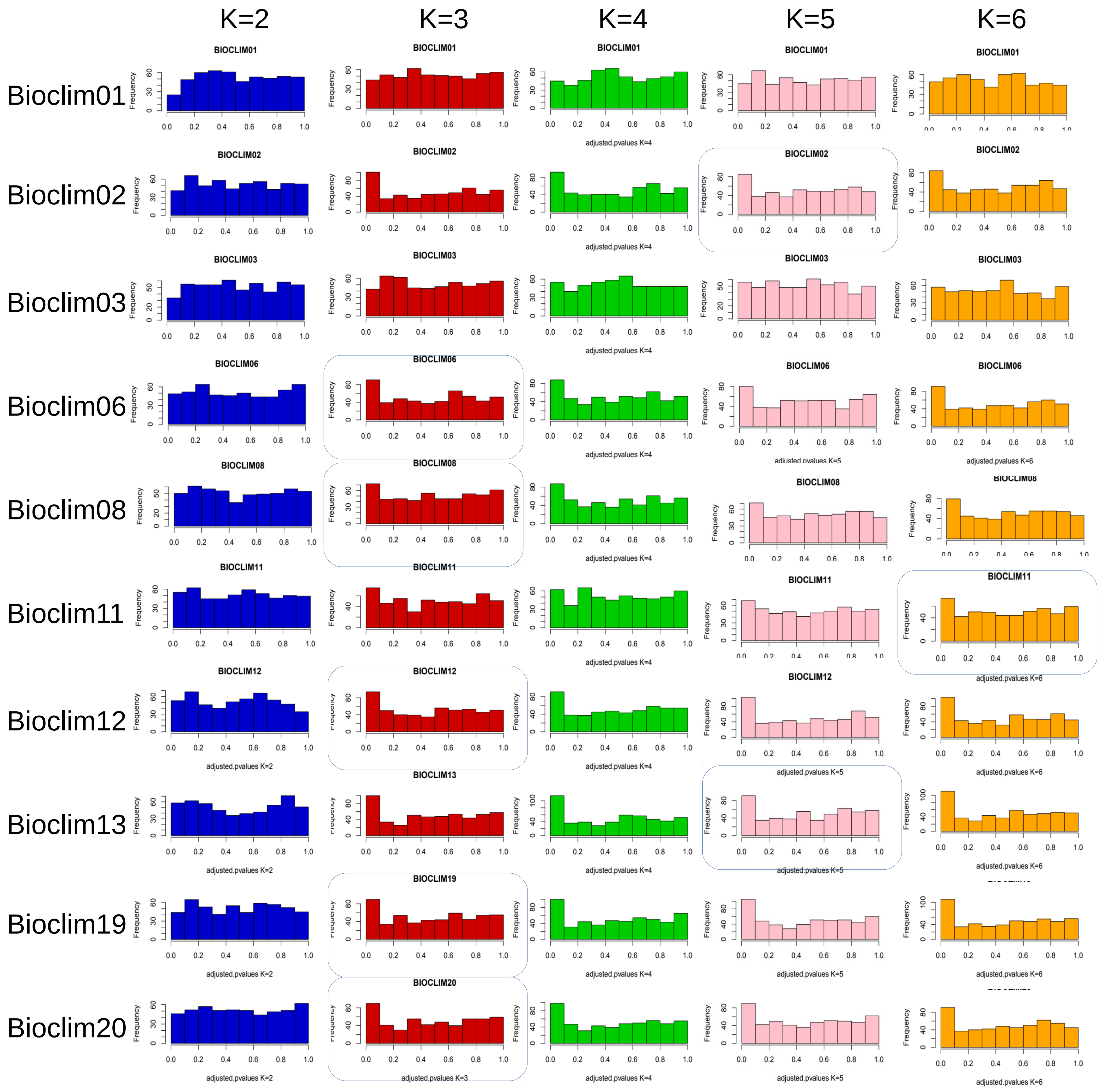

K=2
K=3
K=4
K=5
K=6
Bioclim01
Bioclim02
Bioclim03
Bioclim06
Bioclim08
Bioclim11
Bioclim12
Bioclim13
Bioclim19
Bioclim20
BIOCLIM01
BIOCLIM02
BIOCLIM03
BIOCLIM06
BIOCLIM08
BIOCLIM11
BIOCLIM12
BIOCLIM13
BIOCLIM19
BIOCLIM20
Frequency
adjusted.pvalues K=2
adjusted.pvalues K=3
adjusted.pvalues K=4
adjusted.pvalues K=5
adjusted.pvalues K=6

**Figure S4-6:**

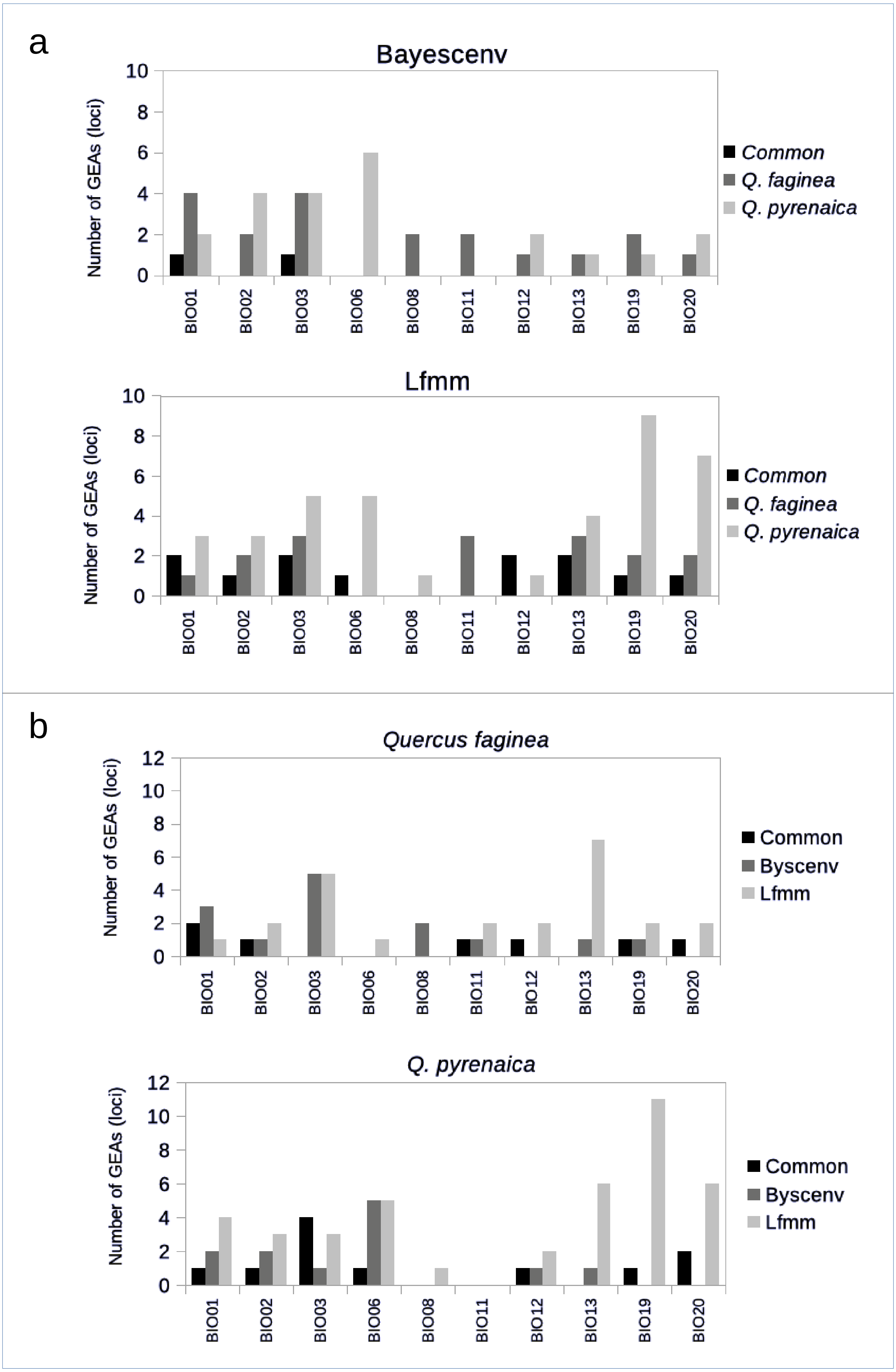
a
Bayescenv
Number of GEAs (loci)
0
2
4
6
8
10
BIO01
BIO02
BIO03
BIO06
BIO08
BIO11
BIO12
BIO13
BIO19
BIO20
*Common*
*Q. faginea*
*Q. pyrenaica*
Lfmm
Number of GEAs (loci)
*Common*
*Q. faginea*
*Q. pyrenaica*
b
*Quercus faginea*
Number of GEAs (loci)
12
Common
Byscenv
Lfmm
*Q. pyrenaica*
Number of GEAs (loci)
Common
Byscenv
Lfmm

**Figure S4-7:**

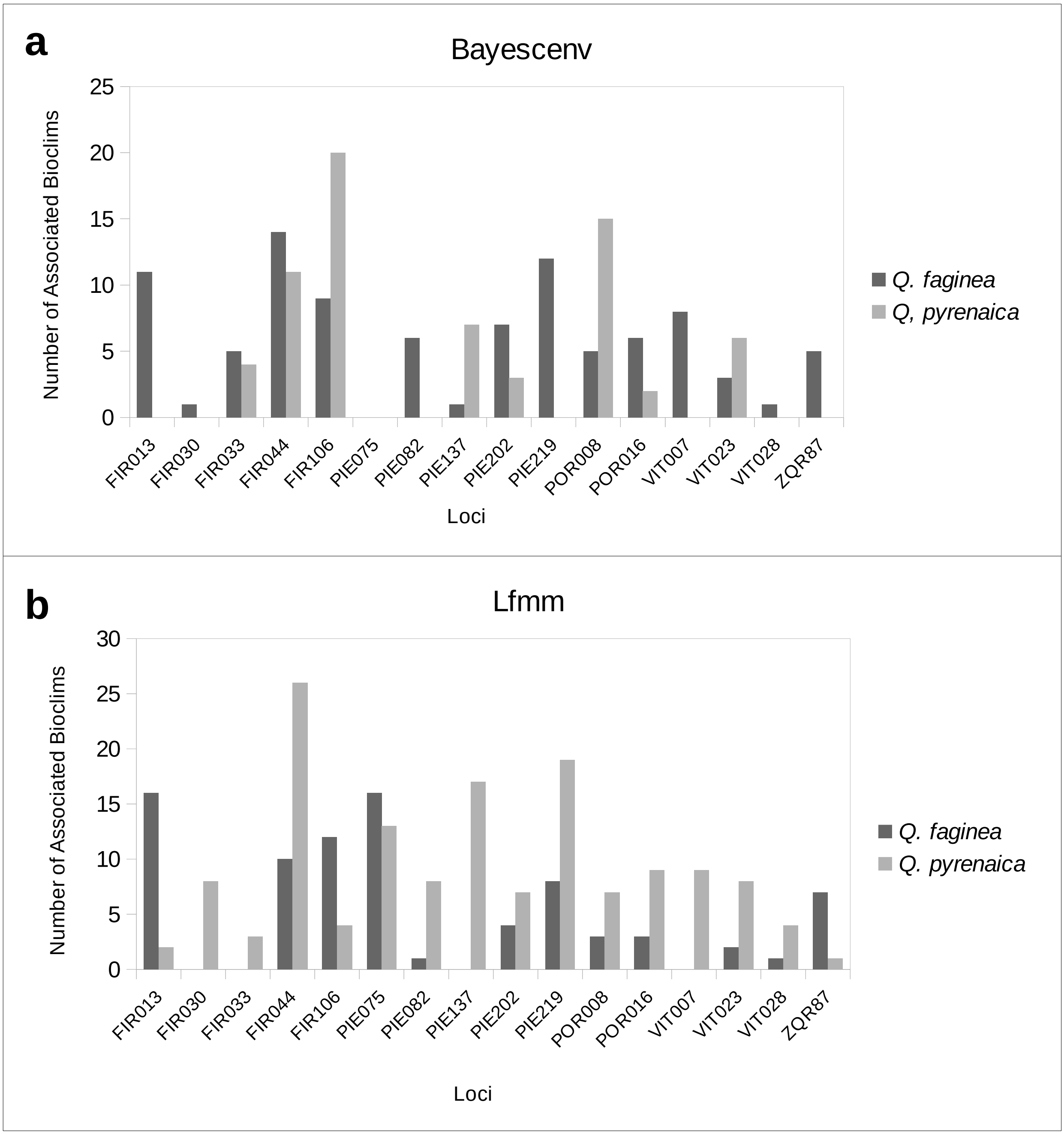
a
Bayescenv
Number of Associated Bioclims
25
20
15
10
5
0
FIR013
FIR030
FIR033
FIR044
FIR106
PIE075
PIE082
PIE137
PIE202
PIE219
POR008
POR016
VIT007
VIT023
VIT028
ZQR87
Loci
Q. faginea
Q, pyrenaica
b
Lfmm
Number of Associated Bioclims
30
25
20
15
10
5
0
FIR013
FIR030
FIR033
FIR044
FIR106
PIE075
PIE082
PIE137
PIE202
PIE219
POR008
POR016
VIT007
VIT023
VIT028
ZQR87
Loci
Q. faginea
Q. pyrenaica

**Supplemental. FILE 5:** ADAPTIVE INTROGRESSION

**Figure S5-1:** Allellic frequencies, for each population, of the 16 local adaptation candidates shared by the two oak speciess. Discontinuous ovals show that outliers alleles from each species belong to both populations of one inter-specific pair. Two introgression events were inferred for locus FIR013, one in population CABpy and another in population IZKfg. FIR044 also showed evidences of two introgression events, but private alleles impeded to infer the donor and recipient species in these cases.2

**Figure S5-2:** Pairwise comparisons between bootstrapped values of the average square distances (ASD) between the outliers (cab, cja-sne, izk) and the 'neutral' loci (cabN, cja-sneN, izkN), in the three population pairs**.**

**Figure S5-3:** Introgressed alleles (arrows) from loci flanking the candidates. PIE155 showed one private allele in the two CAB populations, which does not allow to totally exclude incomplete lineage sorting.

**Figure S5-4:** Divergent selection tests using haplotypes. Tests were performed by comparing the haplotype frequencies of the outlier and hitchhiked loci in the two populations of the target inter-specific pair, to the haplotype frequencies in the remaining populations from each species. Dotted lines indicate significance limits.

**Figure S5-1:**

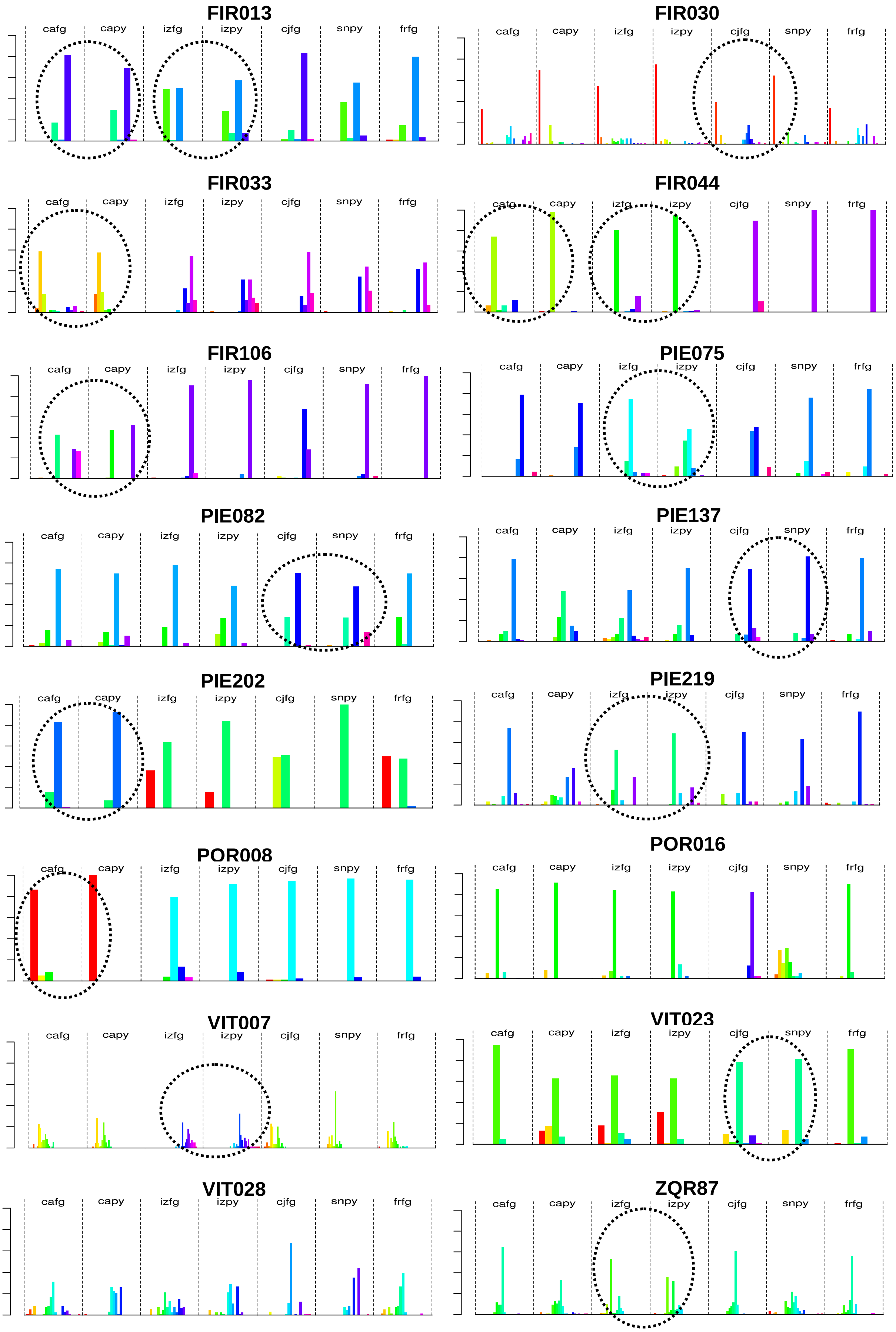
FIR013
cafg capy izfg izpy cjfg snpy frfg
FIR030
cafg capy izfg izpy cjfg snpy frfg
FIR033
cafg capy izfg izpy cjfg snpy frfg
FIR044
cafg capy izfg izpy cjfg snpy frfg
FIR106
cafg capy izfg izpy cjfg snpy frfg
PIE075
cafg capy izfg izpy cjfg snpy frfg
PIE082
cafg capy izfg izpy cjfg snpy frfg
PIE137
cafg capy izfg izpy cjfg snpy frfg
PIE202
cafg capy izfg izpy cjfg snpy frfg
PIE219
cafg capy izfg izpy cjfg snpy frfg
POR008
cafg capy izfg izpy cjfg snpy frfg
POR016
cafg capy izfg izpy cjfg snpy frfg
VIT007
cafg capy izfg izpy cjfg snpy frfg
VIT023
cafg capy izfg izpy cjfg snpy frfg
VIT028
cafg capy izfg izpy cjfg snpy frfg
ZQR87
cafg capy izfg izpy cjfg snpy frfg

**Figure S5-2**

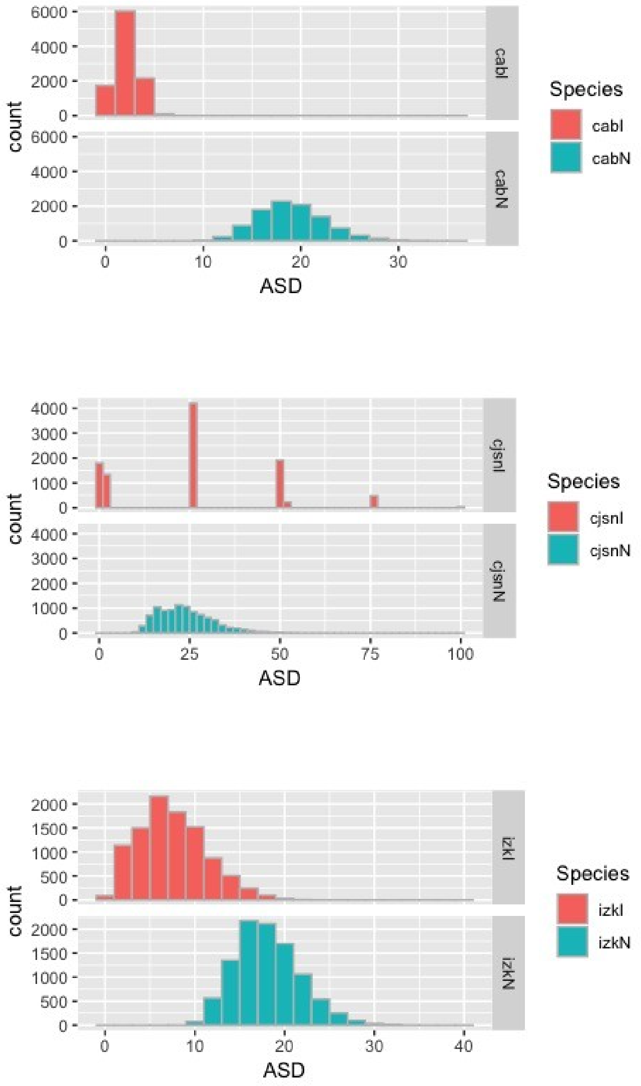
count
cabI
cabN
ASD
Species
cabI
cabN
cjsnI
cjsnN
Species
cjsnI
cjsnN
izkI
izkN
Species
izkI
izkN

**Figure S5-3:**

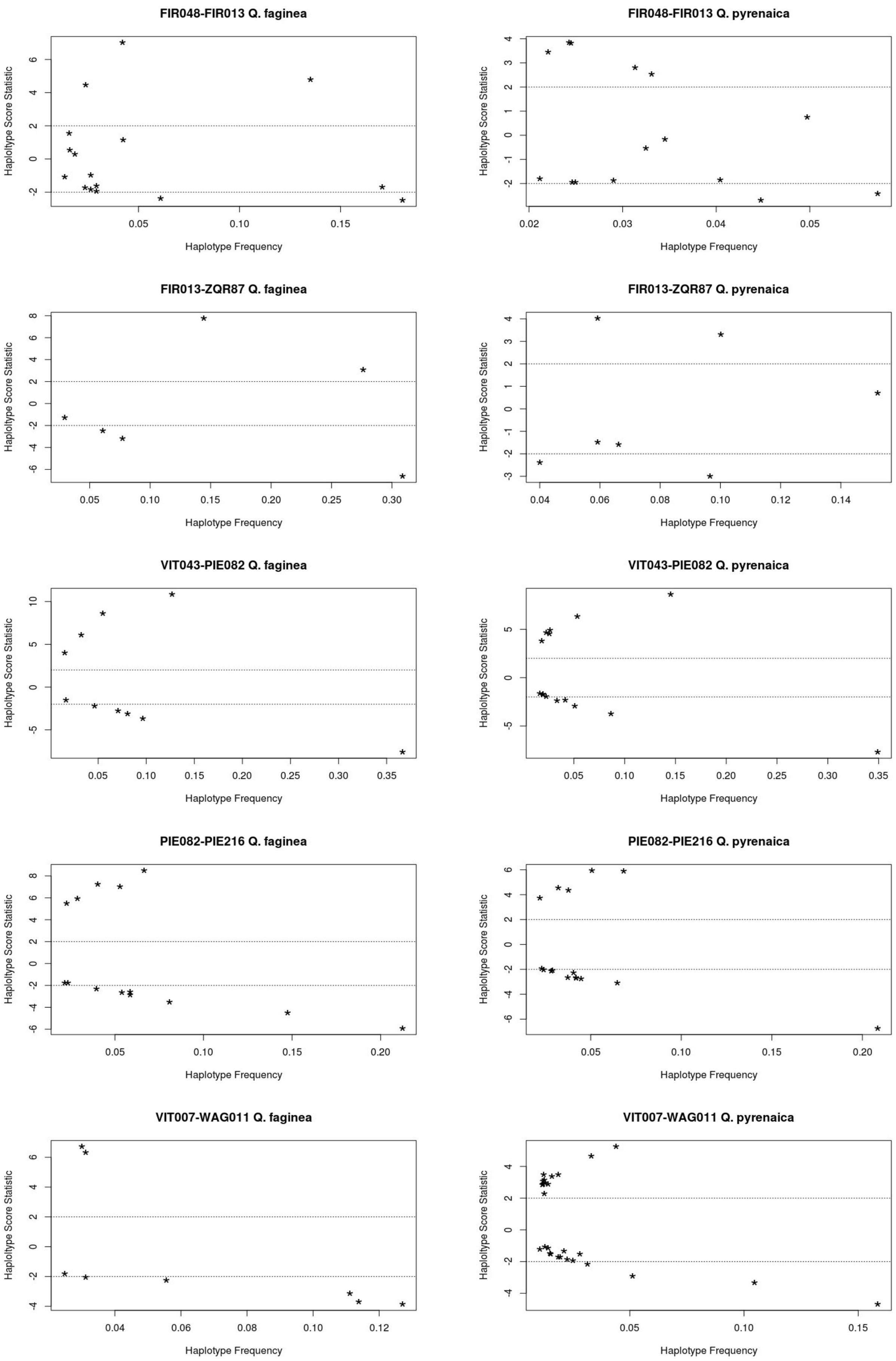
FIR048-FIR013 Q. faginea
FIR048-FIR013 Q. pyrenaica
FIR013-ZQR87 Q. faginea
FIR013-ZQR87 Q. pyrenaica
VIT043-PIE082 Q. faginea
VIT043-PIE082 Q. pyrenaica
PIE082-PIE216 Q. faginea
PIE082-PIE216 Q. pyrenaica
VIT007-WAG011 Q. faginea
VIT007-WAG011 Q. pyrenaica
Haplotype Score Statistic
Haplotype Frequency

**Supplemental FILE 6:** ABC ESTIMATES OF DIVERGENCE WITH GENE FLOW

**Table S6-1:** Model choice statistics for the three inter-specific population pairs. Sequential model choice was performed first among the three coalescent models without central populations, second among the three models with central populations and finally between the best two models for each population pair. Selected models in bold characters.

**Figure S6-1:** Coalescent models of speciation-with-gene-flow in marginal populations under source-sink dynamics with central populations from each species. IMc: continuos gene flow model, AMc: ancient admixture model, SCc: secondary contact model. Tsp: time to speciation, Tpop: time to populations-within-species differentiation, Tam: time to ancient admixture finalization, Tsc: time to secondary contact start; mij: gene flow rates; Nek: population sizes. Thick vertical lines indicate time periods without gene flow between the marginal population pairs. The ghost central populations are represented with dashed background.

**Figure S6-2:** Cross-validation of parameter estimates obtained from the retained simulations (1000), for the CAB populations.

**Figure S6-3:** Posteriors biases for IZK parameters estimated from the 1000 retained simulations. The hypothesis that the posterior cumulative probabilities of the true parameter values are equally distributed over all cross-validation replicates were examined with Kolmogorvo-Smirnov tests (K-S).

**Table S6-1:**

| Populations | AM | | | IM | | | SC | | |
|---|---|---|---|---|---|---|---|---|---|
| | Marginal Density (*p*-value) | Tukey Depth (*p*-value) | Bayes Factors | Marginal Density (*p*-value) | Tukey Depth (*p*-value) | Bayes Factors | Marginal Density (*p*-value) | Tukey Depth (*p*-value) | Bayes Factors |
| CAB-CAB | 6.02E-97 (1.00) | 0.04 (0.39) | 1.67E-96 | 2.13E-091 (1.00) | 0.21 (0.90) | 5.89E-91 | 0.36 (0.42) | 0.10 (0.62) | **1.70E+90** |
| CJA-SNE | 0.14 (0.83) | 0.16 (0.87) | 6.45E-05 | 0.00 (0.38) | 0.12 (0.77) | 1.12E-06 | 2244.75 (0.45) | 0.09 (0.65) | **1.52E+04** |
| IZK-IZK | 8.31 (0.11) | 0.04 (0.38) | **14.57** | 0.10 (0.49) | 0.12 (0.71) | 1.16E-02 | 0.47 (0.46) | 0.13 (0.73) | 5.57E-02 |

| Populations | $AM_C$ | | | $IM_C$ | | | $SC_C$ | | |
|---|---|---|---|---|---|---|---|---|---|
| | Marginal Density (*p*-value) | Tukey Depth (*p*-value) | Bayes Factors | Marginal Density (*p*-value) | Tukey Depth (*p*-value) | Bayes Factors | Marginal Density (*p*-value) | Tukey Depth (*p*-value) | Bayes Factors |
| CAB-CAB | 2.16 (0.01) | 0.09 (0.67) | 2.00E-02 | 107.83 (0.21) | 0.22 (0.93) | **12.04** | 6.80 (0.00) | 0.12 (0.69) | 6.00E-02 |
| CJA-SNE | 7.62 (0.83) | 0.17 (0.91) | 3.30E-01 | 9.58 (0.38) | 0.09 (0.68) | 4.60E-01 | 13.31 (0.20) | 0.06 (0.53) | **0.77** |
| IZK-IZK | 0.03 (0.31) | 0.00 (0.00) | 1.00E-03 | 3.67E-96 (0.73) | 0.14 (0.78) | 5.94E-99 | 616.72 (0.84) | 0.15 (0.75) | **997.99** |

| Populations | Models | Bayes Factors | |
|---|---|---|---|
| CAB-CAB | (SC *vs*. $IM_C$) | 0.02 | **41.67** |
| CJA-SNE | (SC *vs*. $SC_C$) | **23.59** | 0.04 |
| IZK-IZK | (AM *vs*. $SC_C$) | 1.52E-03 | **655.52** |

**Figure S6-1:**

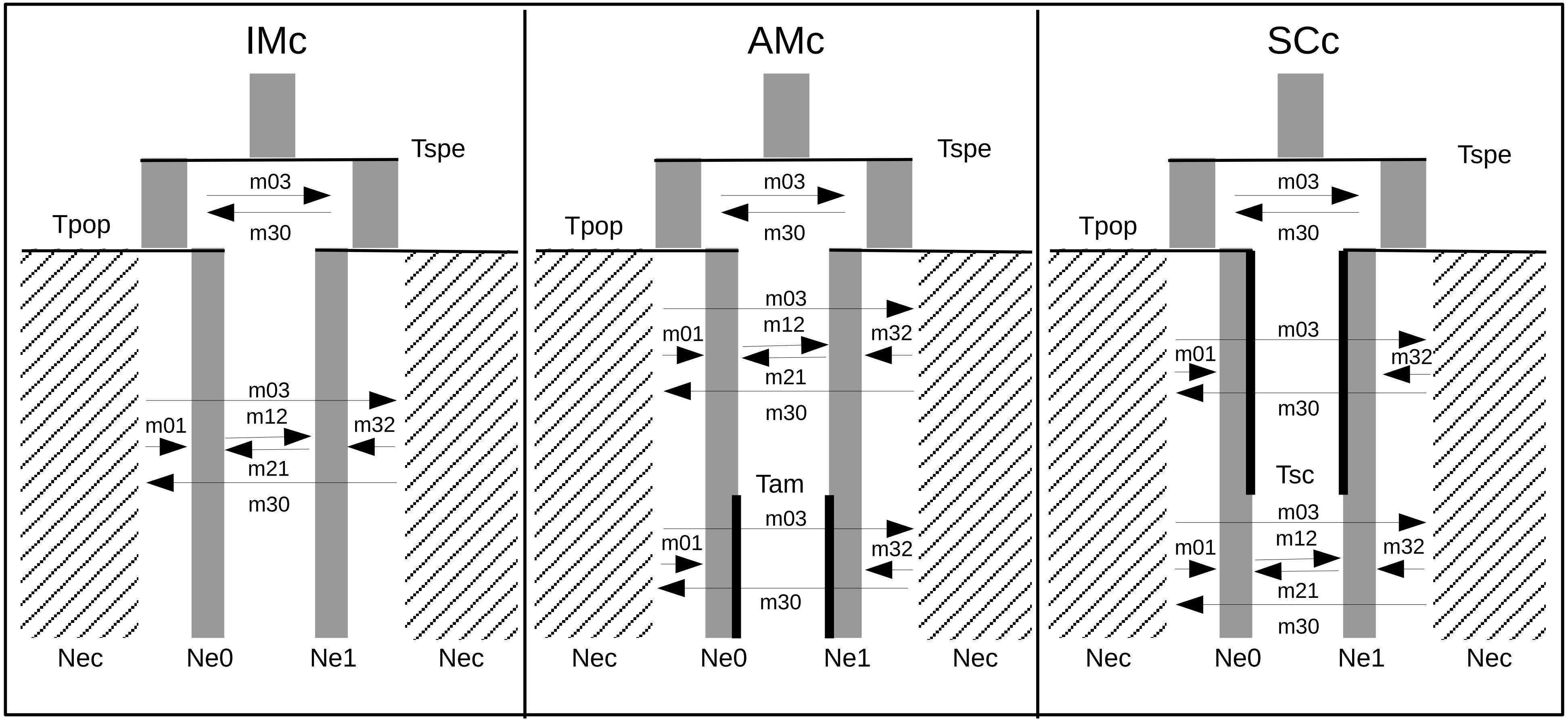
IMc
Tspe
m03
m30
Tpop
m03
m12
m01
m32
m21
m30
Nec
Ne0
Ne1
Nec
AMc
Tspe
m03
m30
Tpop
m03
m12
m01
m32
m21
m30
Tam
m03
m01
m32
m30
Nec
Ne0
Ne1
Nec
SCc
Tspe
m03
m30
Tpop
m03
m01
m32
m30
Tsc
m03
m12
m01
m32
m21
m30
Nec
Ne0
Ne1
Nec

**Figure S6-2:**

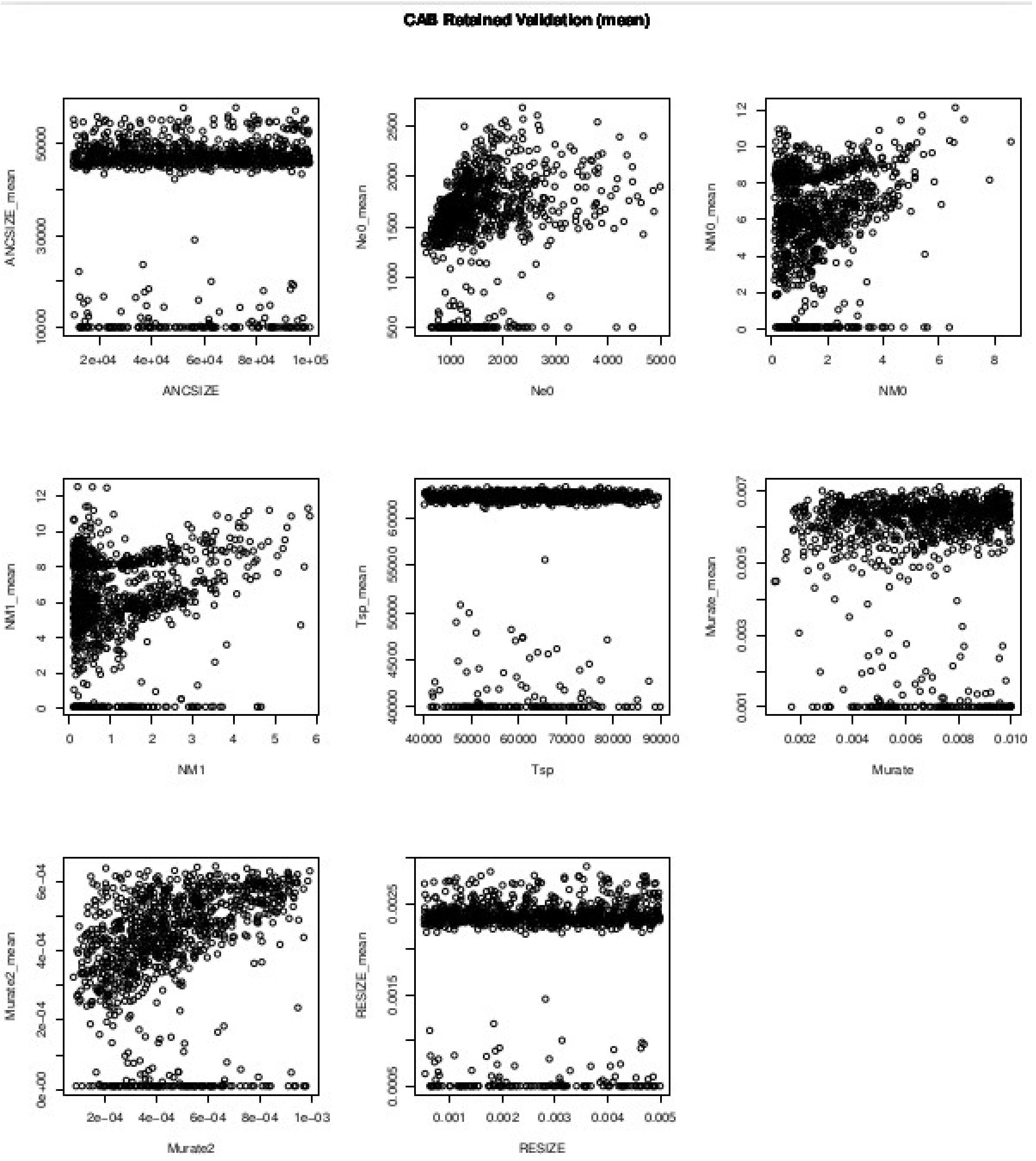
CAB Retained Validation (mean)
ANCSIZE_mean
ANCSIZE
Ne0_mean
Ne0
NM0_mean
NM0
NM1_mean
NM1
Tsp_mean
Tsp
Murate_mean
Murate
Murate2_mean
Murate2
RESIZE_mean
RESIZE

**Figure S6-3:**

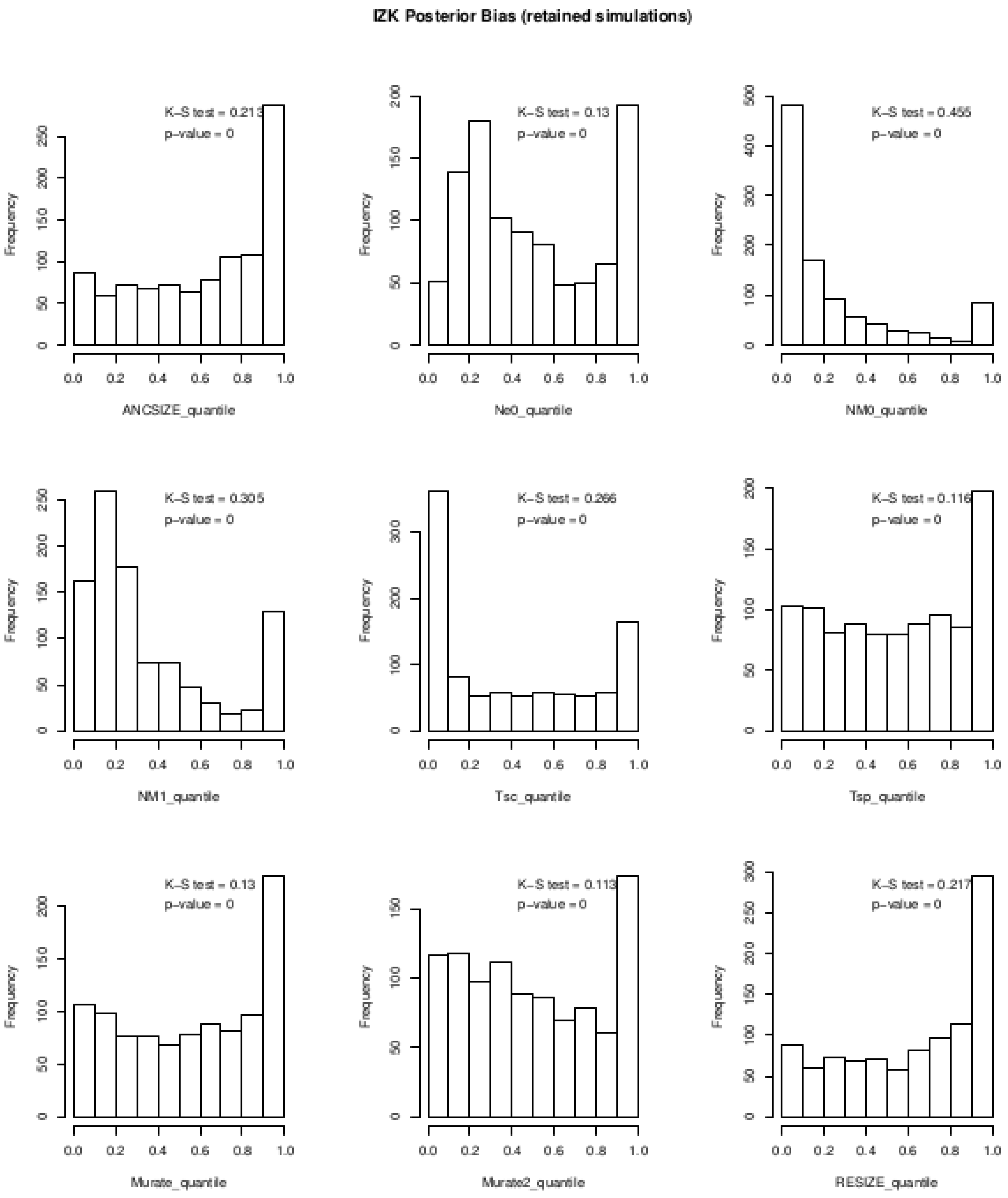
IZK Posterior Bias (retained simulations)
K–S test = 0.213
p–value = 0
ANCSIZE_quantile
K–S test = 0.13
p–value = 0
Ne0_quantile
K–S test = 0.455
p–value = 0
NM0_quantile
K–S test = 0.305
p–value = 0
NM1_quantile
K–S test = 0.266
p–value = 0
Tsc_quantile
K–S test = 0.116
p–value = 0
Tsp_quantile
K–S test = 0.13
p–value = 0
Murate_quantile
K–S test = 0.113
p–value = 0
Murate2_quantile
K–S test = 0.217
p–value = 0
RESIZE_quantile
Frequency
